# Arylsulfamates inhibit colonic *Bacteroides* growth through lipid kinases and not sulfatases

**DOI:** 10.1101/2024.04.27.591440

**Authors:** Conor J Crawford, Charles Tomlinson, Christian Gunawan, Zongjia Chen, Dominic P Byrne, Cosette Darby, Martina L. G. Conti, Tony Larson, Ana S Luis, Stefano Elli, Edwin A Yates, David N Bolam, Sjoerd van der Post, Spencer J Williams, Alan Cartmell

## Abstract

Excessive degradation of the colonic mucin layer by *Bacteroides* within the human gut microbiota drives inflammatory bowel disease in mice. Bacterial carbohydrate sulfatases are key enzymes in gut colonization, as they are elevated in human inflammatory bowel disease and correlate with disease severity. Selective inhibitors of carbohydrate sulfatases could function as sulfatase-selective drugs, allowing precise control of sulfatase activity while preserving these otherwise beneficial bacteria. Arylsulfamates are covalent inhibitors that target a catalytic formylglycine residue of steroid sulfatases, a residue that is also conserved in carbohydrate sulfatases. Here, we find that a library of aryl- and carbohydrate sulfamates is ineffective against *Bacteroides* carbohydrate sulfatases, yet can inhibit human gut microbiota species grown on sulfated glycans. Leveraging thermal proteome profiling, we identify a lipid kinase as the target responsible for these effects. This work highlights the imperative for developing specific inhibitors targeting carbohydrate sulfatases and reveals the adverse effects that arylsulfamates have on *Bacteroides* species of the human gut microbiota.

**Significance statement:** Arylsulfamates are currently the only effective class of sulfatase inhibitors available and offer a potential strategy to treat inflammatory bowel disease driven by gut microbiota carbohydrate sulfatases. Although arylsulfamates inhibit the growth of microbiota *Bacteroides* species on sulfated glycans, this is not mediated through carbohydrate sulfatases but, via a conserved lipid kinase. Carbohydrate sulfatases are resistant to arylsulfamates whilst steroid sulfatases are susceptible despite a conserved active site. Finally, selected complex plant glycans confer a resistant/protective phenotype against the harmful effects of arylsulfamates. These data guide the future development of targeted carbohydrate sulfatase inhibitors and potential drug-prebiotic pairings.

## Introduction

The human gut microbiota (HGM) is a microbial community found throughout the gastrointestinal tract but is densest in the distal colon where it is composed of trillions of bacteria. This community is critical to human health, providing essential vitamins^1,2^, such as soluble B vitamins, calories through short chain fatty acid production^3^, immune system regulation^4,5^, and production of metabolites that influence the gut-brain axis^6,7^. Fermentation of complex carbohydrates underpins all these processes; the complex carbohydrates being derived either from dietary fibre (plant glycans) or the host. In the latter case this involves mainly sulfated glycans from colonic mucin and glycosaminoglycans.

The colonic mucin layer is the most abundant host glycan in the colon. It is composed of a gel forming glycoprotein, MUC2, is 80% glycan by mass and is heavily sulfated^8^. This layer has many biological functions, and provides a colonisable niche and food source for colonic bacteria, whilst simultaneously acting as largely impenetrable barrier protecting the colonic epithelium^9,10^. The Bacteroidota phylum are the major glycan degraders present in the HGM and are enriched in carbohydrate sulfatases, which are essential enzymes for utilisation of sulfated host glycans. In host symbiosis, the model Bacteroidota, *Bacteroides thetaiotaomicron* VPI-5482 (*B. theta*), grazes on colonic mucin *O*-glycans in a carbohydrate sulfatase dependent process^11^. However, in dysbiosis, excessive degradation of the sulfated colonic mucin drives inflammatory bowel disease (IBD) and carbohydrate sulfatases are the enzymatic drivers of this effect in a ‘friend turned foe’ scenario^12–14^. Excessive inflammation caused by loss of the mucin barrier is also a risk factor for colon cancer^15^, the second leading cause of cancer deaths. Thus, effective inhibitors of HGM carbohydrate sulfatases are of high medical relevance.

Sulfatases are classified into four families, S1, S2, S3, and S4^16^. Of the over 160,000 sequences catalogued in the SulfAtlas database, the majority (over 90%, or >145,000 sequences) belong to the S1 family^17^. The S1 family occurs across all domains of life and is the only family that contains sulfatases able to desulfate carbohydrates^16,18,19^. Members of the S1 family of sulfatases have a conserved alkaline phosphatase-like fold characterised by a larger N-terminal domain housing a central mixed β sheet flanked by α helices. This domain abuts a smaller C-terminal subdomain composed of a 4-stranded antiparallel β sheet and a single α helix^20^ **(Figure 1a)**. Within this structural framework, the S1 family employs a catalytic formylglycine (FGly) residue, which is generated co-translationally from a Cys or Ser within the consensus sequence **C**/**S**-X-P/A-X-R, and is situated within an invariant sulfate binding (S) site^18^ **(Figure 1a)**.

**Figure 1.**
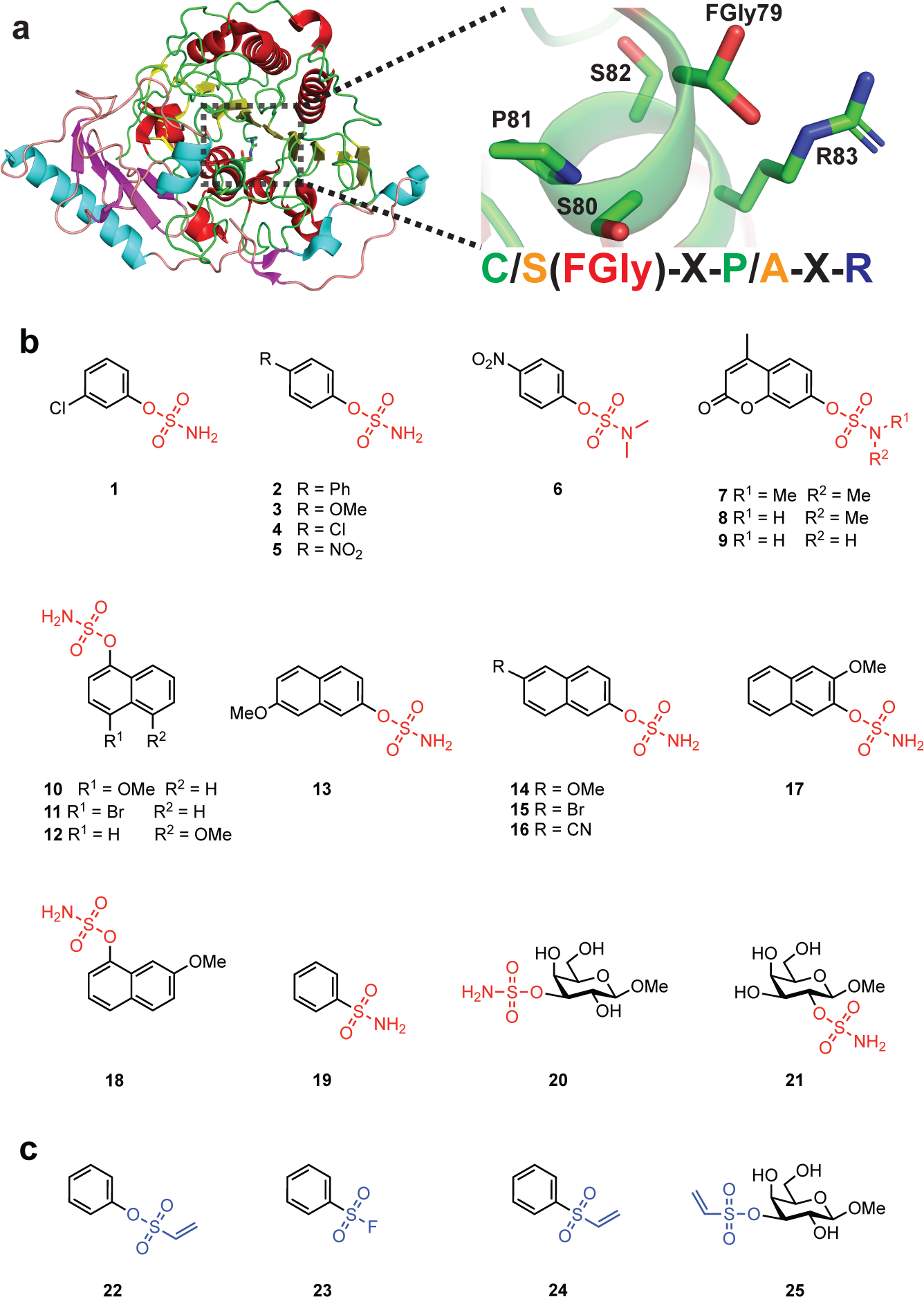
General example of the conserved S1 sulfatase fold, consensus active site sequence, and tested library of aryl- and carbohydrate sulfamates. **a.** 3-D structure of human lysosomal sulfatase GALNS (PDB:4FDJ). The N-terminal domain is coloured red and yellow for alpha helices and beta sheets, with the C-terminal subdomain coloured cyan and magenta for alpha helices and beta sheets. Zoom shows the consensus sequence of GALNS with a formylglycine residue, and with the general consensus pictured to the right. **b.** Structures of aryl- and carbohydrate sulfamate inhibitors used in this study. **c.** Structures of phenylvinyl sulfone, phenylsulfonyl fluoride, phenylvinyl sulfonate, and carbohydrate vinyl sulfone.

The major class of sulfatase inhibitors are arylsulfamates, with the general formula Ar-O-SO_2_NH_2_. These inhibitors, developed by replacing the sulfate group of assorted aryl sulfate substrates with a sulfamate group, are irreversible, active-site directed inhibitors that are broadly effective against steroid sulfatases of both bacterial and eukaryotic origin^21^. Clinical development of arylsulfamates targeting human steroid sulfatase (STS) for breast cancer treatment has advanced to phase II clinical trials, with positive outcomes^22–24^. Arylsulfamates work by targeting the unusual catalytic FGly found in the invariant S site of S1 family of sulfatases, and are believed to be pan-inhibitors of the family^25^ but this not been systematically tested.

Here, we investigate the ability of a panel of aryl- and carbohydrate sulfamates/sulfonates, as well as vinyl sulfones and fluorosulfates **(Figure 1b,c)**, to inhibit the growth of S1 sulfatase-containing Bacteroidota bacteria on sulfated glycans, and to directly inhibit recombinant and lysate derived carbohydrate sulfatases from these species, with the goal of developing novel, targeted strategies to treat IBD. Our data revealed that sulfamate based inhibitors inhibit the growth of *Bacteroides* species on sulfated glycans but, unexpectedly, do not inhibit any of the carbohydrate sulfatases from these organisms. Leveraging thermal proteome profiling we identify lipid kinases, and not sulfatases, as the targets through which arylsulfamates mediate their effects. We also show that arylsulfamate inhibition of growth can be overcome through the utilisation of select, non-sulfated, plant complex glycans. Our data suggest that arylsulfamates may have potential for the treatment of IBD but at the expense of *Bacteroides* species, whilst also uncovering potential prebiotic strategies to protect *Bacteroides* species of the HGM from arylsulfamate drugs when they are delivered orally to treat cancer, or IBD.

## Results

### Arylsulfamates inhibit growth of Bacteroides thetaiotaomicron

*Bacteroides* species of the HGM encode large numbers of S1 sulfatases, which are induced during growth on sulfated glycans. The model organism *Bacteroides thetaiotaomicron* VPI-5482 (*B. theta*) encodes 28 S1 sulfatases, 16 of which have been characterised as carbohydrate sulfatases. We hypothesized that growth of *B. theta* would be affected by the action of arylsulfamates when grown on sulfated glycans, but not on unsulfated glycans. Therefore, we cultivated *B. theta* in rich brain heart infusion media, and in minimal media supplemented with sulfated (chondroitin sulfate C and heparin and unsulfated (D-glucose, larch arabinogalactan, and potato galactan) carbon sources, each in the presence of a panel of arylsulfamate compounds at 1 mM **(Figure 2)**. This panel includes 3 control compounds: *N-*methyl and *N,N*-dimethyl sulfamates **6-8**, which should not inhibit S1 sulfatases. DMSO was used at a final concentration of 1%, which did not affect growth of *B. theta* **(Figure S1)**.

**Figure 2.**
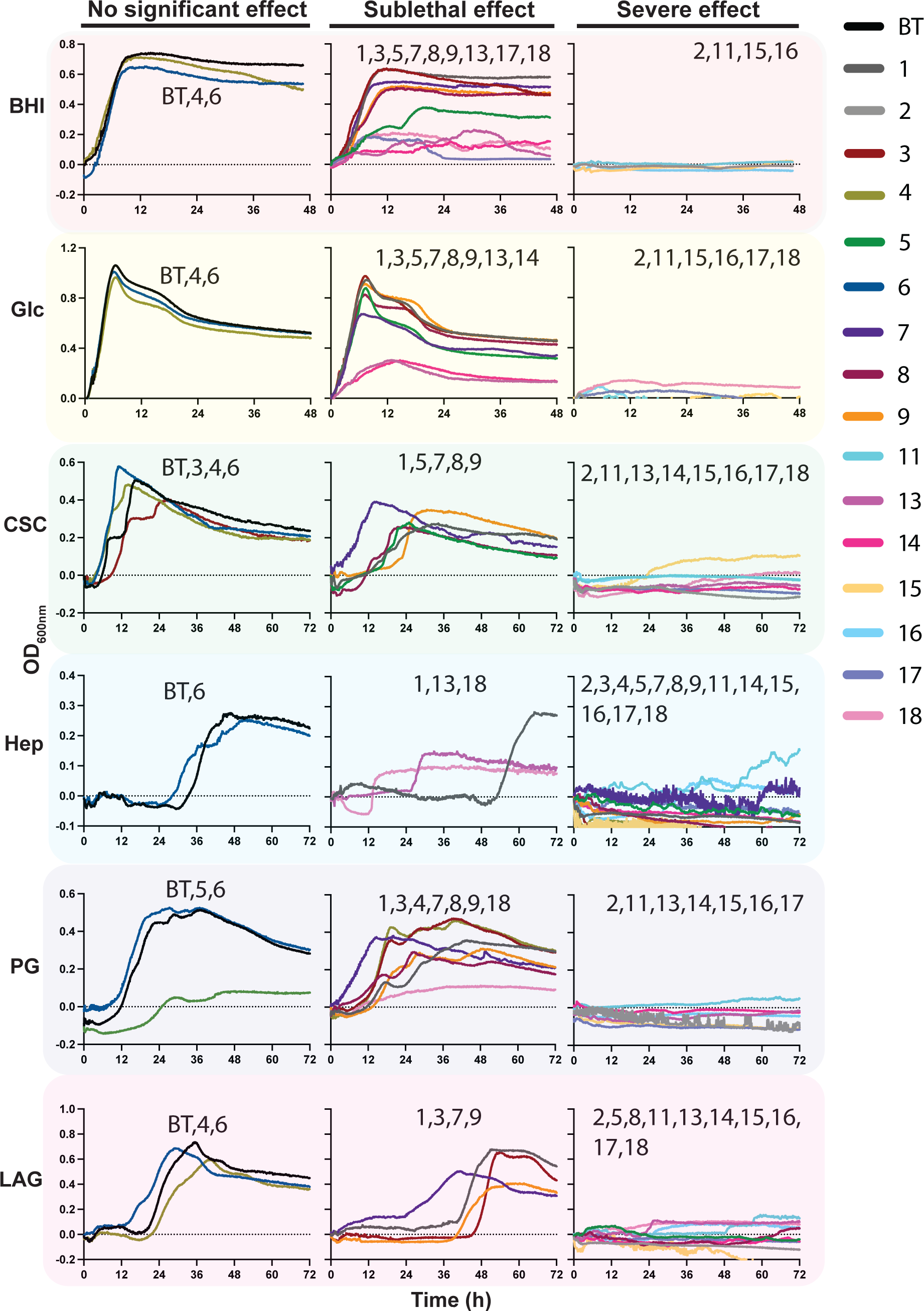
Growth of *Bacteroides thetaiotaomicron* on varied carbon sources in the presence of arylsulfamate inhibitors. *Bacteroides thetaiotaomicron* (*B. theta*) grown on varied carbon sources; two rich carbon sources, brain heart infusion media (BHI) and glucose (Glc); two sulfated host glycan carbon sources, chondroitin sulfate c (CSC) and Heparin (Hep), and two plant glycan carbon sources, potato galactan (PG) and larch arabinogalactan (LAG), in the presence of arylsulfamates inhibitors **1**-**18** (see Figure 1). Complex glycans and arylsulfamate inhibitors were used at a concentration of 5 mg/ml and 1 mM, respectively. All carbon sources were utilised in a minimal media except BHI, which was purchased from Sigma and dissolved in water. The traces are the mean of 3 independent growth experiments, and errors bars have been omitted for clarity; individual graphs with error bars are presented in supplemental figures **S2-S7**. Severe effects were judged qualitatively on the basis no classically identifiable features of a bacterial growth were able to be measured: a lag and mid-exponential growth phase leading to robust increases in OD_600nm_. For sublethal growth curves that displayed the aforementioned classical features, and were measurable, a two-tailed unpaired t-test was performed to check for significance at a threshold of p<0.05 (Table S1 and S2). A difference was called significant if it met this threshold in any one of three criteria: lag phase, growth rate, and max O.D. reached.

We observed variable growth rates for *B. theta* with different arylsulfamate and carbon source combinations **(Figure 2 and Figure S2-S7)**. Arylsulfamates based on a naphthyl scaffold (compounds 11-17) caused severe growth defects on all substrates. In contrast, the effect of arylsulfamates based on a substituted phenyl group (**1**, **3-6**) varied substantially. *B. theta* displayed mild growth defects in the presence of these compounds when grown on brain heart infusion media and glucose, while growth on the sulfated host glycan chondroitin sulfate C, and the unsulfated plant glycan potato galactan, was strongly perturbed by **1** and **5**. Compound **3** induced an increase in lag phase on chondroitin sulfate C, and compound **4** caused complete inhibition of growth on the sulfated host glycan heparin **(Figure 2)**. On the unsulfated plant glycan larch arabinogalactan, **1** and **3** caused extension of *B. theta* lag phase whilst **5** completely inhibited growth. Compound **2**, a 4-phenylphenyl sulfamate, inhibited growth on all substrates. For the coumate scaffold, compounds **7-9** had only mild effects when *B. theta* was grown on brain heart infusion media and glucose. However, they increased the lag phase and reduced on chondroitin sulfate C, and completely inhibited growth on heparin. Compounds **3** and **7** increased lag phase and reduced growth on the plant glycan larch arabinogalactan, whilst compound **8** completely inhibited growth. When potato galactan was used as the carbon source **3** caused a mild growth defect whilst **7** reduced the lag phase; **8** caused a significant defect but still allowed growth.

The variable impact of arylsulfamates on the growth of *B. theta*, observed for both sulfated and unsulfated glycan substrates, suggests that these compounds do not target S1 carbohydrate sulfatases. Additionally, the fact that the 3 negative control *N*-methylsulfamates (compounds **6-8**) also influenced growth of *B. theta* provides further support that the observed growth defects are not mediated through an S1 sulfatase dependent mechanism. A significant finding is that the same arylsulfamate structure has a different effect depending on the carbon source utilised. Brain heart infusion media, and the non-sulfated substrates potato galactan and glucose in a minimal media context, appear to switch *B. theta* to a more resistant/protective metabolic state evinced by milder phenotypes and fewer severe effects.

### Arylsulfamates do not inhibit HGM B. theta S1 carbohydrate sulfatases

With the observation that the arylsulfamates, including the negative control *N*-methylsulfamates, affected growth of *B. theta* on unsulfated and sulfated polysaccharides we next wanted to investigate the effect of our panel of arylsulfamates on purified S1 sulfatases. We first confirmed the potency of our panel of arylsulfamates against two S1 steroid sulfatases: *Pa*AstA from *Pseudomonas aeruginosa*, belonging to S1 subfamily 4 (S1_4), and commercially available snail sulfatase *Hp*Sulf from *Helix pomatia*, belonging to S1_2. The inhibitory activity of each compound was determined by measuring the sulfatase activity of each enzyme incubated in the presence of inhibitor, or following ∼24 h pre-incubation and jump-dilution ‘washout’^26^. At a concentration of 1 mM, all arylsulfamates were broadly equivalent in their ability to completely inhibit both enzymes; only minor differences being observed **(Figure 3, Figure S8-S11, Table S3 and S4).** Against *Pa*Asta, partial inhibition was observed for **3** and **4** before complete inactivation **(Figure 3a, left panel and Figure S8)**, whilst pre-incubation with the two compounds resulted in complete inhibition **(Figure 3a, right panel)**. Competitive inhibition of *Pa*Asta was observed with **7** in the in assay experiments but not in the enzyme incubation and jump dilution assays **(Figure 3a, b)**. Similarly, for *Hp*Sulf, residual activity observed when treated with **10** and **12** was completely lost in the jump dilution experiment **(Figure 3a and Figure S11)**. These findings suggest different inactivation rates between the enzymes and are consistent with the irreversible, covalent, nature of the inactivation process. Importantly, near control levels of activity were observed upon treatment with the *N-*methyl and *N,N*-dimethyl sulfamates **6-8 (Figure 3a, right panel)**.

**Figure 3.**
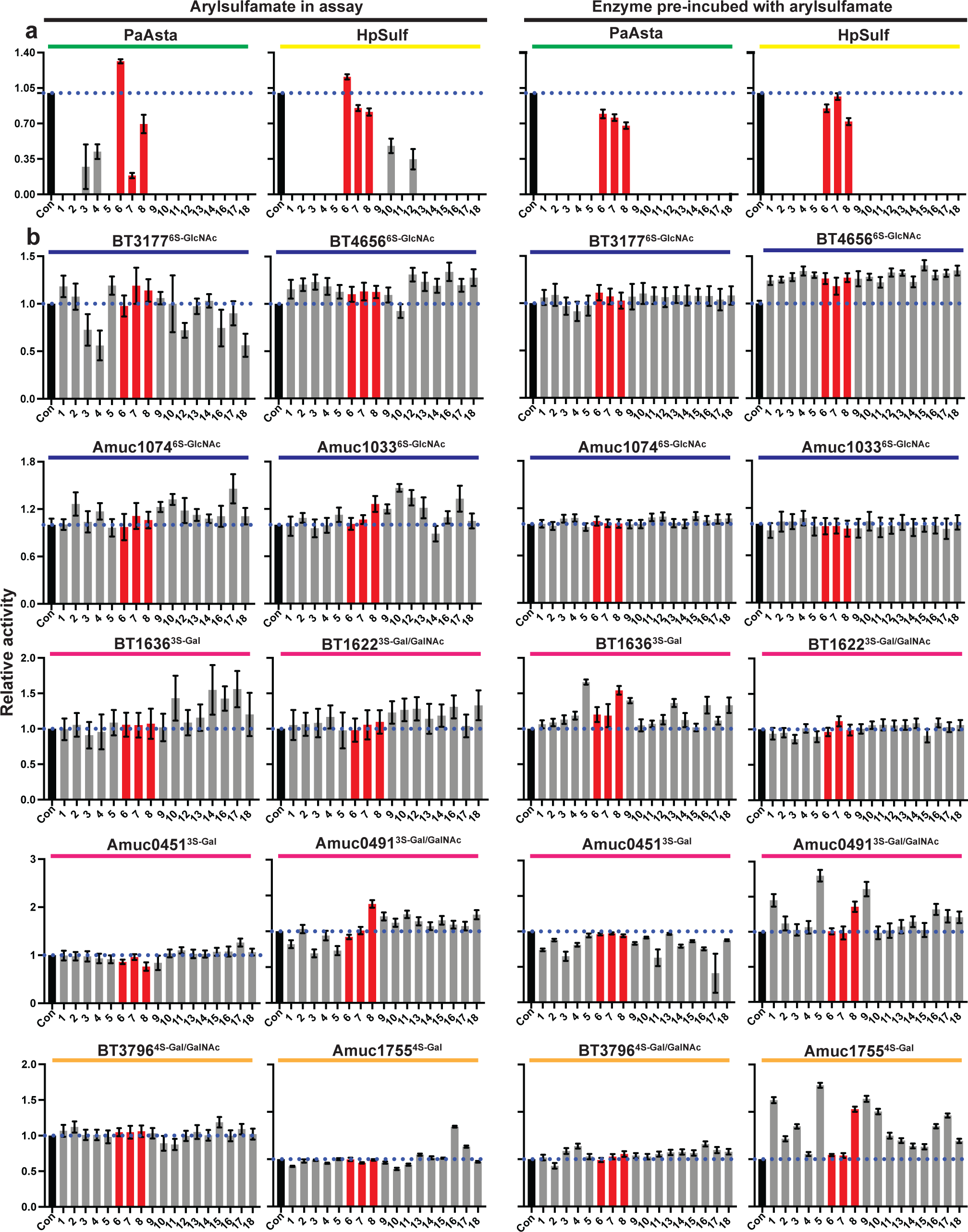
The effectiveness of arylsulfamate inhibitors against S1 steroid and carbohydrate sulfatases. **a.** The steroid sulfatases *Pa*Asta and *Hp*Sulf, belonging to families S1_4 (green bar) and S1_2 (yellow bar), respectively, assayed against the panel of arylsulfamates (compounds **1**-**18** listed in Figure 1). **b.** Activities of family S1_11 sulfatases (BT3177^6S-GlcNAc^, BT4656^6S-GlcNAc^, Amuc_1033^6S-GlcNAc^, Amuc_1074^6S-GlcNAc^; blue bar above), S1_20 sulfatases (BT1622^3S-Gal/GalNAc^, BT1636^3S-Gal^, Amuc_0451^3S-Gal^, Amuc_0491^3S-Gal^; pink bar above), and the S1_16 sulfatases (BT3796^4S-Gal/GalNAc^ and BT1755^4S-Gal^; orange bar above) against the panel of arylsulfamates. For both a and b the right two panels display activity with 1 mM arylsulfamate compound in the assay relative to untreated enzyme, whilst the left two panels are overnight preincubation of the enzyme with selected 1 mM arylsulfamates then dilution of this preincubation mix into the reaction. Con indicates enzyme control without arylsulfamate. Kinetic curves from which the data are derived are shown in supplemental figures S8-S31. The blue dotted line indicates 100% activity level and the red bars indicate control compounds.

Having confirmed the activities of the arylsulfamates against two S1 steroid sulfatases, we next examined their activity towards 10 bacterial S1 carbohydrate sulfatases from the HGM members *B. theta* and *Akkermansia muciniphila ATCC BAA-835 (A. muc*). These enzymes target common linkages found in host glycans: four from S1_11 (BT3177^6S-GlcNAc^, BT4656^6S-GlcNAc^, Amuc1033^6S-GlcNAc^, and Amuc1074^6S-GlcNAc^) target *O*6 sulfated *N*-acetyl-D-glucosamine (6S-GlcNAc), four from S1_20 (BT1636^3S-Gal^, BT1622^3S-Gal/GalNAc^, Amuc0451^3S-Gal^, and Amuc0491^3S-Gal/GalNAc^) target *O*3 sulfated D-galactose/*N*-acetyl-D-galactosamine (3S-Gal/GalNAc), and two from S1_16 (BT3796^4S-Gal/GalNAc^ and Amuc1755^4S-Gal^) target *O*4 sulfated D-galactose/*N*-acetyl-D-galactosamine (4S-Gal/GalNAc). No significant inhibition of any of the enzymes was observed when assaying their activity in the presence of 1 mM arylsulfamate, even after pre-incubation **(Figure 3b, Supplementary Figure S12-S31, Table S3 and S4)**. Thus, while the panel of arylsulfamates inhibit bacterial and eukaryotic S1 steroid sulfatases, they do not inhibit S1 HGM carbohydrate sulfatases.

We also explored the ability of two potent growth inhibitory sulfamates, **2** and **17 (Supplementary Figure S32 and Figure S33)**, to inhibit S1 carbohydrate sulfatases in cell lysates. Cell lysates derived from *B. theta* grown to mid-exponential growth phase on chondroitin sulfate A (CSA) or Hep (to stimulate production of native S1 GAG carbohydrate sulfatases) were incubated with arylsulfamates **2** and **17** for 1 hour, then sulfated mono- or disaccharide substrates were added. Compounds **2** and **17** did not impact the production of the final desulfated products (GlcNAc for Hep substrates and GalNAc for CSA substrates) as judged by TLC and HPAEC **(Supplementary Figure S32b,c)**. Thus, compounds **2** and **17** are not effective inhibitors of natively produced S1 GAG carbohydrate sulfatases, and the effects observed on bacterial growth are likely to be due to effects on a non-sulfatase target.

### Carbohydrate sulfamates and vinylsulfonates do not inhibit the S1_20 sulfatase BT1636^3S-Gal^

The refractory nature of S1 carbohydrate sulfatases to arylsulfamate inhibition may arise from the inability to bind at the active site of these enzymes, which has evolved to bind sulfated glycans. Notably, S1 carbohydrate sulfatases exhibit low activity towards 4-nitrophenyl sulfate (4NP-SO_3_), which in contrast is rapidly hydrolysed by the family S1 steroid sulfatase *Pa*Asta **(Supplementary Figure S34)**. We selected the well-studied S1_20 sulfatase BT1636^3S-Gal^, which desulfates 3*O* sulfated D-galactose (3S-Gal) and is essential for *B. theta* to utilise colonic mucin *O*-glycans, and to competitively colonise the colon of mice^11^. We prepared two substrate analogues, D-galactose-3-*O*-sulfamate (**20**) and 3-*O*-vinylsulfonyl D-galactose (**25**), the isomeric compound D-galactose-2-*O*-sulfamate (**21**), aryl vinyl sulfone analogues **22** and **24**, aryl fluorosulfate (**23**), and aryl sulfamate (**19**) **(Figure 4a)**. Of these compounds, the only activity towards BT1636^3S-Gal^ was for compounds **22**, **23** and **24 but only** after pre-incubation; **22** showed a ∼3 fold increase in activity while **23** and **24** showed just over 50 % inhibition **(Figure 4b, Supplementary Figure S35, S36, and Table S5).** We assessed whether any of these compounds could bind by studying the inactive mutant Cys77Ser BT1636^3S-Gal^ using a thermal shift assay (TSA).). No change in melting temperature was observed for compounds **20**, **21**, **23**, or **25** at concentrations up to 1.0 mM, whilst compounds **19, 22,** and **24** reduced the melting temperature (Tm), suggesting binding causes protein destabilisation (**Figure 4c, Supplementary Figure Figure S37, and Table S6**). In contrast, a substrate of BT1636^3S-Gal^, 3S-Gal, stabilised the protein (ΔTm = 3.0 °C) at a concentration of 1 mM **(Figure 4c, Supplementary Figure S37 and Table S6)**. Collectively, these data demonstrate that the carbohydrate sulfamate **20**, and its vinylsulfonyl analogue **25**, are not inhibitors of BT1636^3S-Gal^ and do not bind the enzyme.

**Figure 4.**
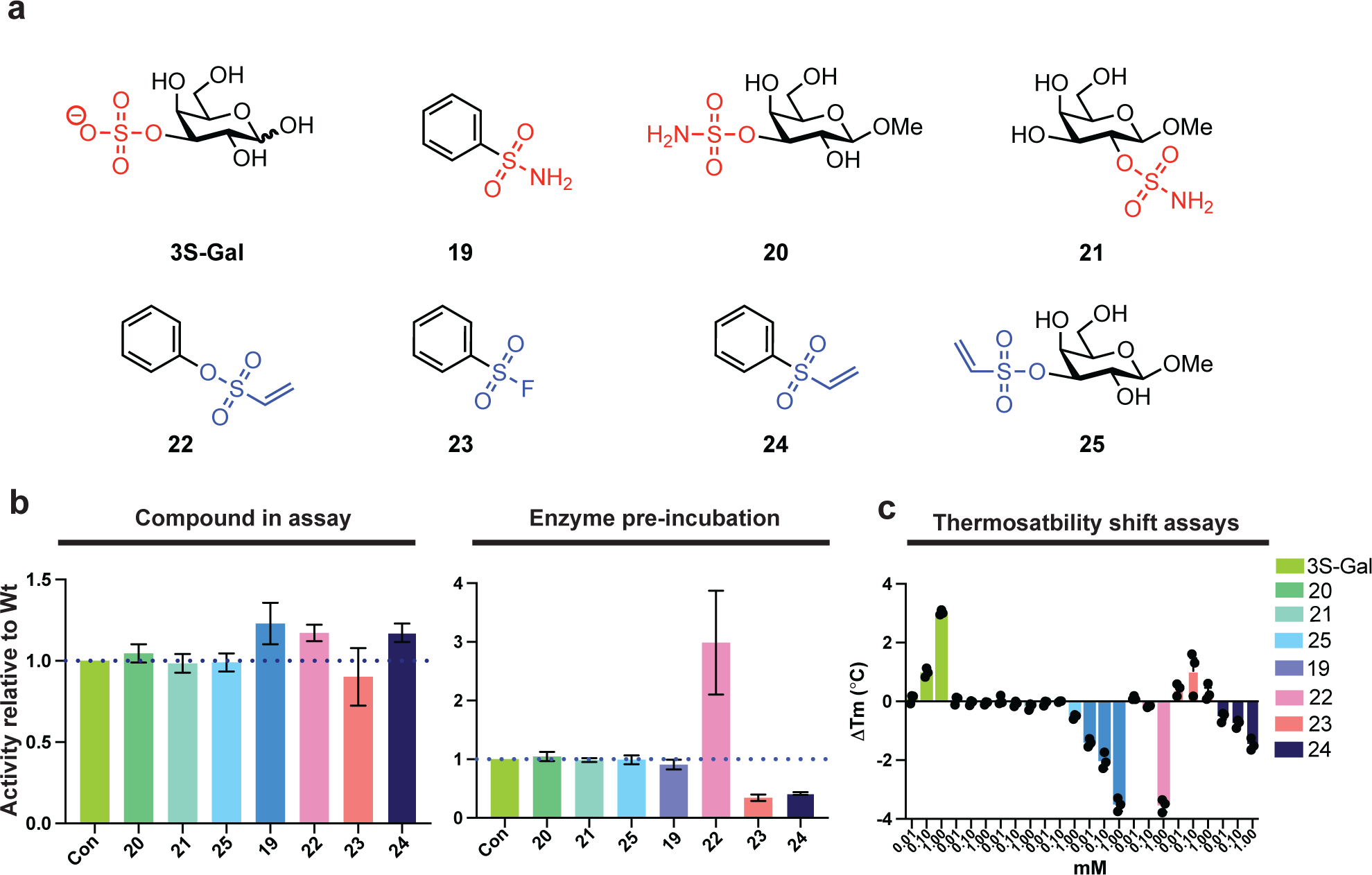
The carbohydrate sulfamate substrate mimics are ineffective against the S1 carbohydrate sulfatase BT1636^3S-Gal^. **a.** Structure of the carbohydrate sulfonate/sulfamates and their aryl equivalents**. b.** Plots show the activities of carbohydrate sulfamates, carbohydrate sulfovinyl, and arylsulfonates against BT1636^3S-Gal^. Left plot shows activity of enzyme treated with the compounds relative to untreated enzyme, whilst right plot show the relative activity after the enzyme had been preincubated with the compounds overnight then diluted into the reaction mixture. **c.** Thermostability shift assays (TSA) using inactive BT1636^3S-Gal^ against the substrate 3S-Gal, and the carbohydrate sulfamates, carbohydrate vinylsulfonate, and arylsulfonates. All compounds in **b** were used at a concentration of 1 mM.

### Growth inhibition by arylsulfamates extends to multiple HGM Bacteroidota species

We selected nine additional HGM *Bacteroides* species to see if the growth effects observed for *B. theta* in the presence of arylsulfamate compounds **2** and **17 (Figure 2)** applied more broadly. We also included the arylsulfamate anticancer drugs Irosustat and estradiol sulfamate. No growth was observed for any species in the presence of 1 mM of arylsulfamate **2 (Figure 5 and Supplementary Figure S38)**. Arylsulfamate **17** exhibited growth defects across all species; in particular, *B. cellulosilyticus* for which no growth could be detected. Irosustat potently inhibited growth with all but one species, *B. fragilis*, showing limited or no growth **(Figure 5 and Supplementary Figure S39)**. Only very mild growth defects were observed for estradiol sulfamate **(Figure 5 and Supplementary Figure S39)**.

**Figure 5.**
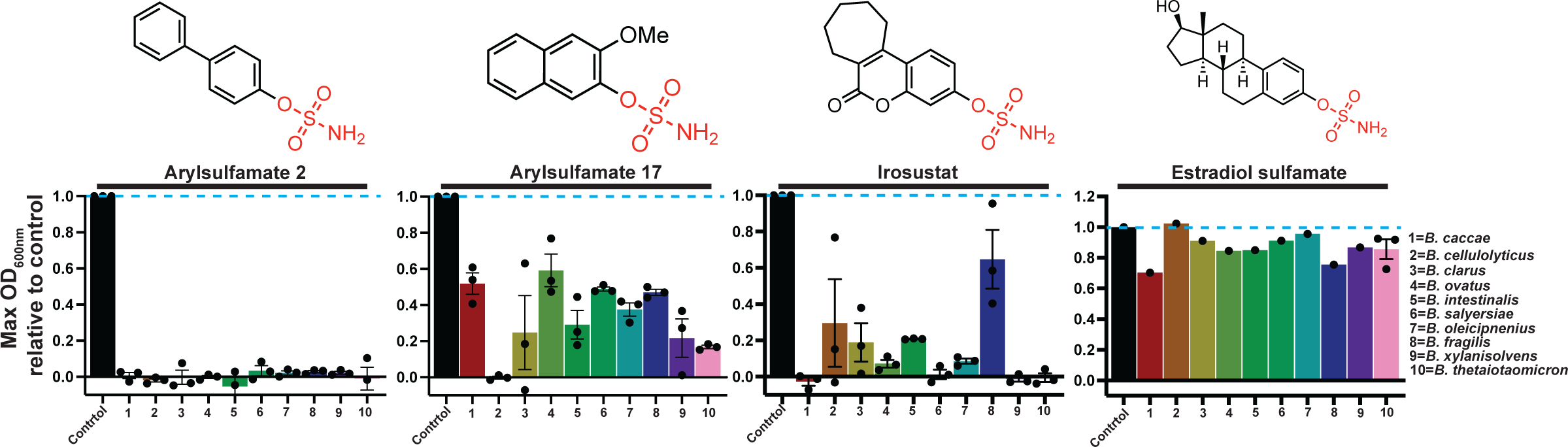
Arylsulfamates broadly affect growth of *Bacteroides* spp. of the HGM. *Bacteroides* species were grown in BHI media in the presence, or absence, of 1 mM of the listed arylsulfamate. The data plotted are the maximal OD_600nm_ observed for arylsulfamate treated growths relative to the untreated cells, which were each run in triplicate and randomly paired, then averages derived. For all the estradiol growth experiments limited material meant this shows a single experiment, except for *B. theta* which was run in triplicate.

### Thermal proteome profiling identifies a lipid kinase target for arylsulfamates

We next applied thermal proteome profiling (TPP) to determine the targets of arylsulfamate compounds **2** and **17**. TPP is a non-targeted mass spectrometry technique that can identify protein-compound interactions based on changes in a protein’s thermal stability. Increased, or decreased, thermal stability is quantified as an increase, or decrease, in the relative abundance of peptides originating from the soluble protein fraction over a temperature gradient following incubation with a ligand^27^.

Cell lysates were derived from *B. theta* grown on chondroitin sulfate A. *B. theta* lysates were treated with inhibitors **2** or **17**, or DMSO control and then heated at 10 different temperatures between 45–72 °C, and the soluble proteome subjected to trypsinisation, and analysed by liquid chromatography-mass spectrometry. The *B. theta* lysate proteome showed a relatively narrow melting temperature ranging between 50 and 58 °C (90% range) **(Supplementary Figure S40a)**. A total of 2550 proteins were identified in the combined analysis, from which 11 S1 sulfatases were identified, including the three upregulated in response to chondroitin sulfate A (BT1596, BT3333, and BT3349)^28^ **(Supplementary Figure S40b,c)**. The melting temperature (Tm) was established for 2112 proteins **(Supplementary Figure S40d)** based on the detection of at least two unique peptides in a minimum of two replicate analyses (see Methods). Melting curves were determined for 8 out of the 11 identified S1 sulfatases **(Supplementary Figure S41)**, none of which was significantly affected by treatment with compounds **2** or **17**, indicating no direct interaction.

TPP identified 7 proteins for which arylsulfamate **2** induced significant changes in Tm, and 4 proteins for arylsulfamate **17**, indicating a potential arylsulfamate-protein interaction **(Figure 6a, b, c)**. Some of these hits can be discounted as causing the defective growth phenotype: disruption of BT0600, BT2504, and BT3464 through transposon mutagenesis results in no phenotype when grown on supplemented BHI media^29^. BT0133 is a homologue of the *E. coli* protein MurQ (having 45% identity), a protein that produces GlcNAc-6-phosphate from 6*O* phosphorylated *N*-acetyl-D-muramic acid (MurNAc). Loss of MurQ in *E. coli* does not cause a no growth phenotype^30^ and no MurNAc was detected in our experiments. This leaves a total 7 candidate proteins (BT0218, BT1443, BT1889, BT4322, BT4335, BT4346, and BT4480), of which only one was identified in TPP experiments with arylsulfamates **2** and **17**, the uncharacterised protein BT4322 **(Figure 6b,c)**.

**Figure 6.**
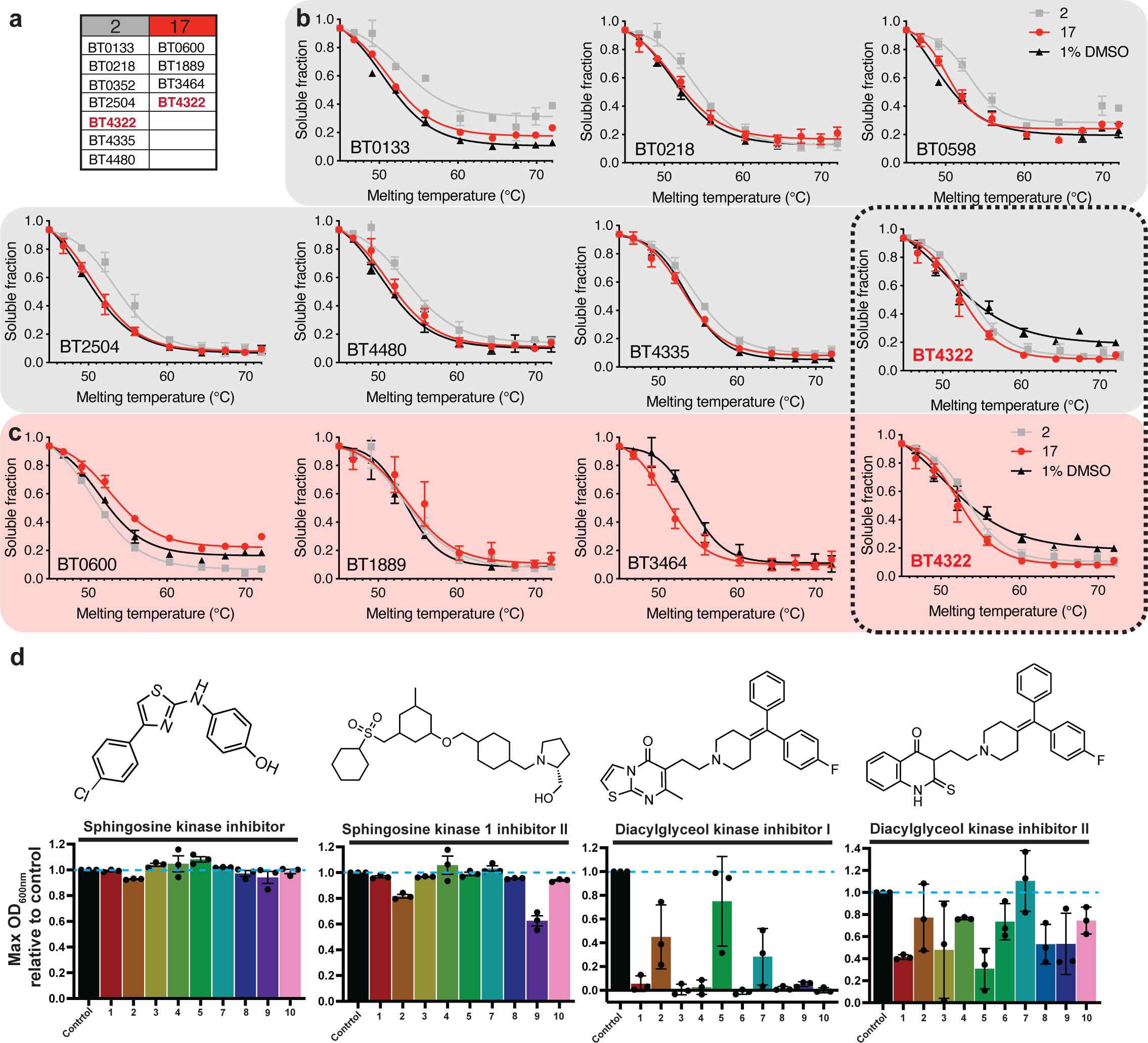
Global protein thermal stability reveals off target interactions of arylsulfamate inhibitors 2 and 17. **a.** Table of proteins that display an altered Tm in the presence of arylsulfamates **2** and **17**. **b.** Proteins identified with a significant change (see methods and Dataset 1) in melting point upon treatment with inhibitor **2** (n=2). **c.** Proteins identified with a significant change in melting point upon treatment with inhibitor **17** (n=2). The dashed box highlights BT4322 as the only protein common to both groups. **d.** *Bacteroides* species were grown in BHI media in the presence, or absence, of 0.125 mM or 0.0625 mM diacylglycerol kinase inhibitor I and II, respectively. The data plotted are the maximal OD_600nm_ observed for inhibitor treated growths relative to the untreated controls, which were each run in triplicate and randomly paired, then averages derived.

BT4322 is homologous to the cytoplasmic diacylglycerol kinases (DGK) YerQ and DgkB, from the gram-positive bacteria *Bacillus subtilis*^31^ and *Staphylococcus aureus*^32^, respectively, and the human sphingosine kinase SphK1^33^. YerQ is essential for growth of *B. subtilis* and is involved in production of lipoteichoic acid (LTA), a cell wall glycophospholipid^31^. BT4322 is predicted to have the same fold and catalytic apparatus as both DgkB (PDB:2QV7) and SphK1 (PDB:3VZB), and similar ATP binding site motifs and lipid binding pockets **(Supplementary Figure S42)**. BT4322 is therefore likely to be a *B. theta* lipid kinase, but the nature of the lipid substrate is yet to be determined. LTA is specific to gram positive bacteria, and so BT4322 may act on another membrane lipid in *B. theta*, or another as yet unknown target. BLASTp using BT4322 against the nine other arylsulfamate-sensitive *Bacteroides* bacteria returned a single orthologue from each species, with 100% query coverage, and a minimum of 84% identity suggesting a conserved role for these proteins across these organisms **(Supplementary Figure S43 and Table S7**).

### Diacylglycerol kinase inhibitors significantly inhibit the growth of Bacteroides species

We next investigated whether we could recapitulate the arylsulfamate induced growth defects assigned to inhibition of the putative lipid kinase BT4322 using commercially available sphingosine and diacylglycerol kinase inhibitors. Diacylglcerol kinase inhibitor (DAGKI)-I was effective at inhibiting the growth of all species with only *B. cellulosilyticus*, *B. intestinalis*, and *B. oleicipnenius* showing limited, albeit highly variable, growth with extended lag phases **(Figure 6d and Supplementary Figure S44)**. Bacteria treated with DAGKI-II exhibited a milder phenotype, but this compound still inhibited the growth of all species (except for *B. oleicipnenius*), lowering the final OD and extending the lag phase **(Figure 6d and Supplementary Figure S44)**. No growth defect could was observed with sphingosine kinase inhibitor (SKI), whilst sphingosine kinase 1 inhibitor (SK1I)-II caused mild phenotypes in four species, manifesting as an elongated lag phase, except for *B. xylanisolvens* which showed severe defects **(Figure 6d and Supplementary Figure S45)**. These data support the view that the observed arylsulfamate growth defects are mediated through inhibition of BT4322, which functions as a diacylglycerol kinase, and not a sphingosine kinase.

### Destabilising inhibitor interactions with BT4322 correlate with severe phenotypes

BT4322 was expressed recombinantly in *E. coli* and its interaction with inhibitors and ATP assessed by DSF. Arylsulfamates **2** and **17** show a concentration dependent destabilisation up to 1 mM **(Figure 7)** which is consistent with the TPP data **(Figure 6b,c)**. The DAGK-I inhibitor also caused concentration-dependent destabilisation, similar to arylsulfamates **2** and **17**, and correlates with a no growth phenotype for *B. theta* grown in the presence of DAGK-I **(Figure 6d)**. By contrast DAGK-II caused a concentration dependent stabilisation **(Figure 7)** and correlated with only a mild growth defect **(Figure 6d)**. The substrate ATP also stabilised BT4322, supporting its activity as a lipid kinase.

**Figure 7.**
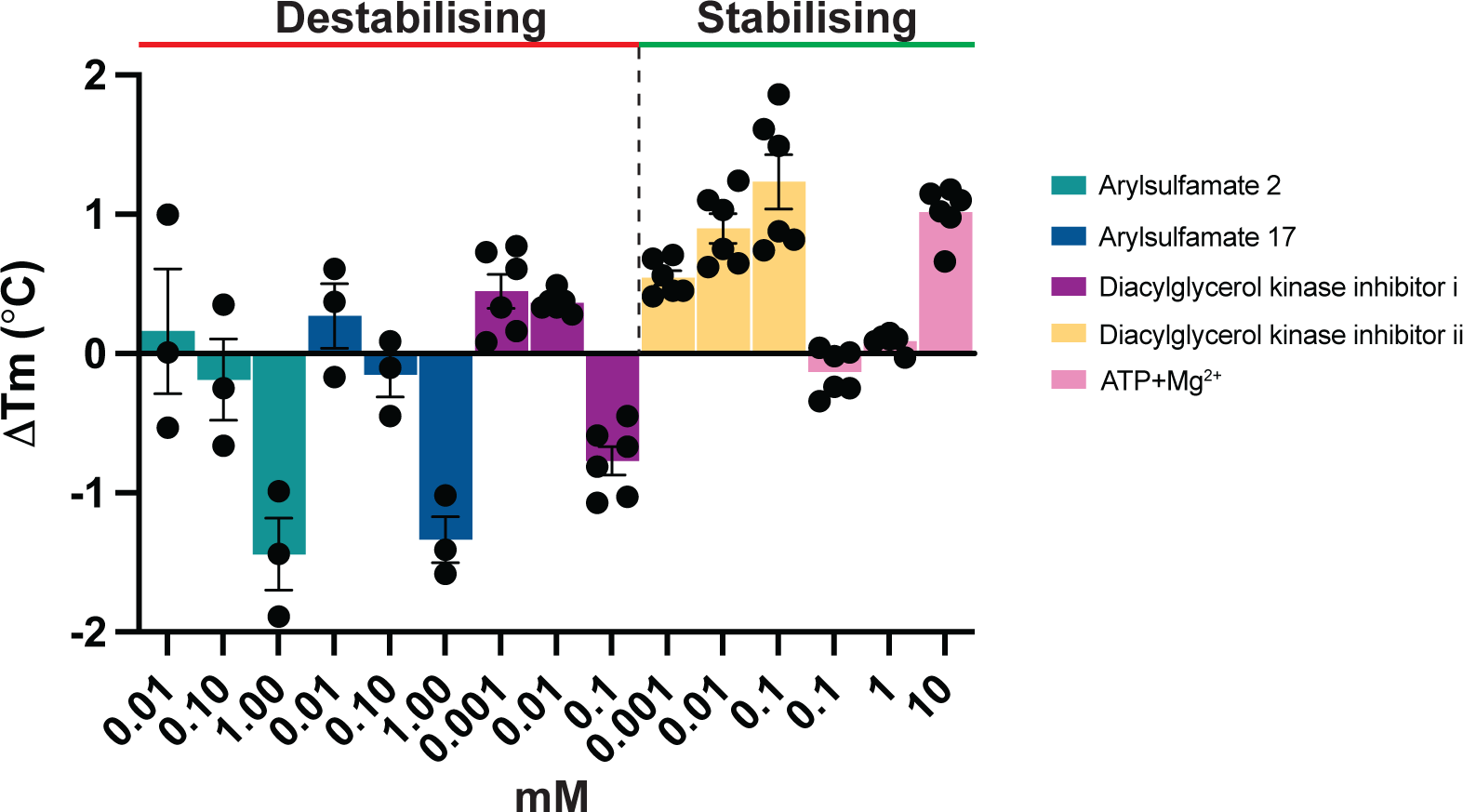
Recombinantly expressed BT4322 binds arylsulfamates, diacylglycerol kinase inhibitors, and ATP. Thermostability shift assays of BT4322 against arylsulfamate and diacylglycerol kinase inhibitors, and the kinase substrate ATP. A final protein concentration of 5 μM, in 100 mM Bis-Tris-Propane (BTP), pH 7.0, and 150 mM NaCl, supplemented with the appropriate ligand concentration and 5 % DMSO. A minimum of three, and a maximum of six, independent assays were performed for each protein and protein ligand combination and standard errors of the mean derived.

### Ligand docking of arylsulfamates into putative lipid kinase BT4322

To understand how arylsulfamates may interact with BT4322 we performed blind ligand docking (no pocket specified) with arylsulfamates **2**, **17** and Irosustat using the AlphaFold 2 predicted model of BT4322. The docking results for Irosustat with BT4322 predicted binding almost exclusively to the putative lipid binding pocket, with some minor solutions elsewhere. Similar results were obtained for arylsulfamate **2** and **17**, but with some models also predicting binding in the ATP site **(Supplementary Figure S46)**. This binding of the arylsulfamates in the lipid pocket is consistent with what has been observed for other lipid kinase inhibitors^32,34^.

### In vitro and in vivo effects of arylsulfamates on colonic bacteria

Growth of *B. theta* in the presence of sublethal doses of arylsulfamate **2**, and subsequent lipidomic analysis, showed a drastically altered membrane lipid composition whilst only causing a 15% reduction in total cell mass. These observations indicate arylsulfamate **2** affects lipid metabolism, supporting the TPP target identification of a lipid kinase as mediating the effects of arylsulfamates **(Figure 8a)**. The largest decrease (log^2^ FC ∼-5) was in the phosphorylated ceramides, suggesting ceramides as the potential lipid substrate for BT4322, whilst large increase (log^2^ FC >8) was in membrane unphosphorylated triglycerides were also observed **(Figure 8a)**.

**Figure 8.**
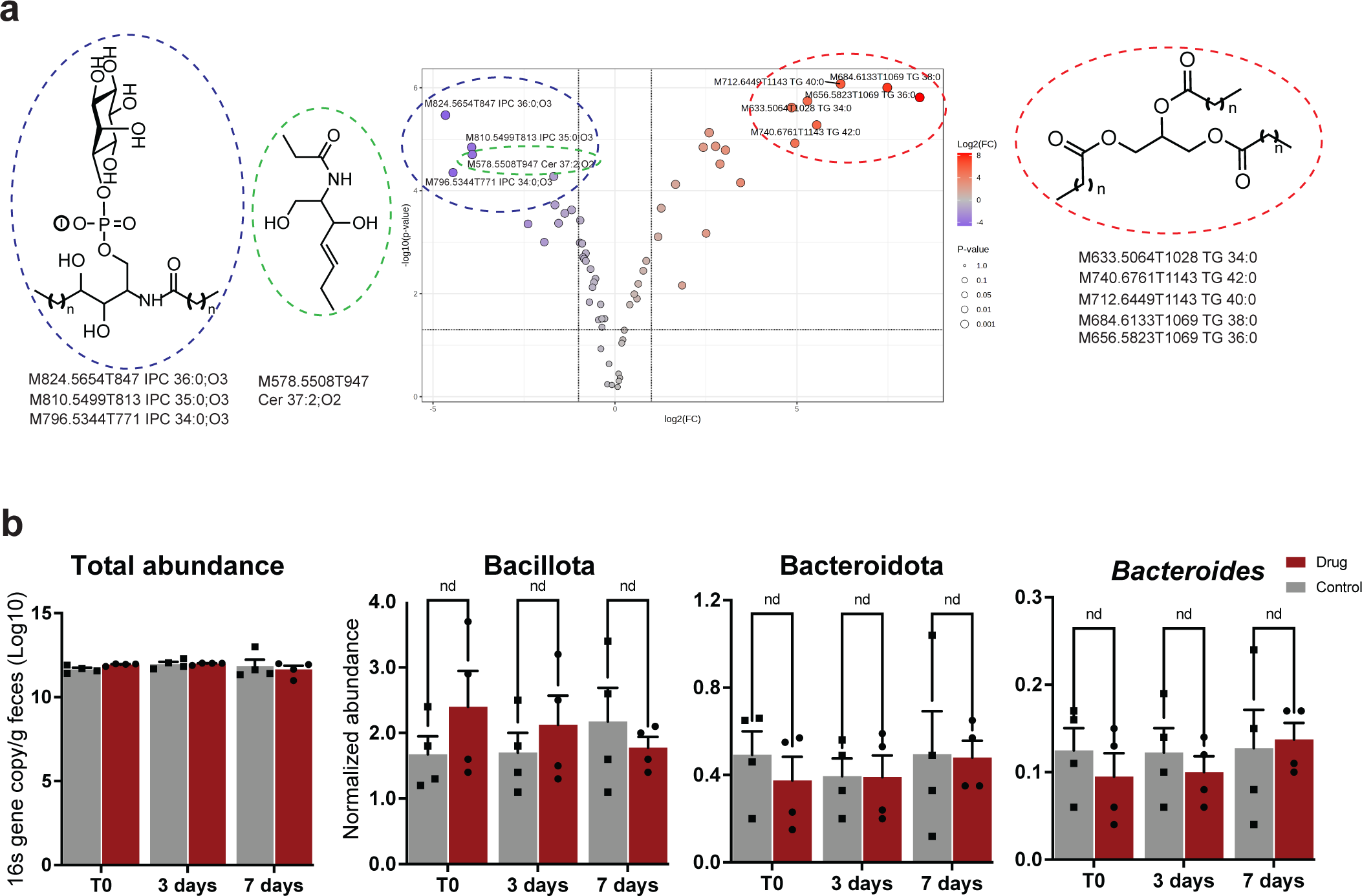
The effect of arylsulfamate on the growth of colonic bacteria *in vitro* and *in vivo*. **a.** a volcano plot showing the alterations of membrane lipids when *B. theta* is grown in brain heart infusion media in the presence of 0.2 mM arylsulfamate 2. The structures of the up and down regulated lipids are shown to the right and left of the plot, respectively. **b.** The changes in relative abundance of colonic bacteria in the mouse gut from animals that have been orally gavaged with the phase II arylsulfamate Irosustat; nd indicates no significant difference as determined from a two-tailed, unpaired, t-test with a threshold of p-value of ≤0.05.

In *in vivo* animal studies, where mice were gavaged with Irosustat, showed no significant alteration of Bacteroidota and Bacillota levels, or at the genus level for *Bacteroides* **(Figure 8b)**. However, these data lack the resolution to resolve at the species level where it is possible changes may have occurred. Furthermore, any changes in bacterial lipid composition *in vivo* were not assessed. Additionally, Irosustat may be absorbed in the small intestine and not reach the colon to exert an effect on colonic bacterial species.

## Discussion

Arylsulfamates are effective irreversible, active site directed, inhibitors of S1 steroid sulfatases^25^ and display relatively mild side effects when administered to humans to treat hormone dependent cancers in phase II clinical trials^24,35^. S1 carbohydrate sulfatases have been implicated in driving several disease states and it was presumed that arylsulfamates would be broadly effective across the S1 family. However, the work presented here shows that bacterial S1 carbohydrate sulfatases of the HGM, despite sharing the invariant sulfate binding subsite (S subsite) and a FGly nucleophile, are not inhibited by arylsulfamates *in vitro* and *in vivo*. Furthermore, substrate mimicking carbohydrate sulfamate not only failed to inhibit the target S1 carbohydrate sulfatase, but showed no evidence of binding **(Figure 4b,c)**. Collectively, these data demonstrate that aryl or carbohydrate sulfamates are not effective inhibitors of the HGM S1 carbohydrate sulfatases tested here. Similarly, the vinyl sulfone and fluorosulfate modifications are ineffective inhibitors. With the S subsite being invariant, the apparent differing susceptibilities of these enzyme classes must be due to the distinctive chemical environments and/or structural geometries of glycan versus steroid binding subsites. Steroid binding sites include a hydrophobic patch that can accommodate lipophilic molecules and may allow the sulfamate moiety to adopt an alternative binding conformation relative to when arylsulfamates bind to carbohydrate sulfatases. In contrast, carbohydrates are hydrophilic, and S1 carbohydrate sulfatases have polar substrate binding regions, which form extensive hydrogen bonding networks^11,20,36,37^. The polar, hydrogen bond-driven nature of the glycan binding site may impose a level of rigidity on substrate interactions that may be incompatible with alternative binding modes that may be necessary to accommodate sulfamate groups. Moreover, the aryloxy group in arylsulfamates is a good leaving group (p*K*_a_ ≤ 10, depending on structure), whereas a carbohydrate alcohol is a much poorer leaving group (p*K*_a_ value approximately 16).

Carbohydrate sulfamate inhibitors have been developed against the *endo*-acting human S1 carbohydrate sulfatase Sulf1 by replacing the sulfate in HS di-, tri, and tetrasaccharides with a sulfamate^38,39^. The most potent compound was the trisaccharide sulfamate, with an IC_50_ value of 0.5 μM (versus 70 μM for the disaccharide), but this compound operated through a competitive mechanism^39,38^ distinct from the time-dependent inactivation seen with arylsulfamates, with potency largely driven by the number of sugar residues. In contrast, TSA gave no evidence of the monosaccharide sulfamate **20** binding to the *exo*-acting BT1636^3S-Gal^, suggesting that the sulfamate cannot interact with this sulfatase.

TPP analysis highlighted BT4322 as the most likely target through which the arylsulfamates mediate their growth inhibiting effects on *B. theta* and lipidomic analyses of *B. theta*, grown in the presence of arylsulfamate **2**, showed large changes in membrane lipid composition supporting BT4322 as the target. Studies using lipid kinase inhibitors further support BT4322 as a lipid kinase. A single homologue of BT4322, with high sequence identity, is found in *B. theta* and the nine other species examined here, suggesting an important role for BT4322 in phospholipid biosynthesis and a lack of functional redundancy in *Bacteroides*. A homologue of BT4322 in *Bacillus subtilis* is implicated in peptidoglycan metabolism and cell membrane integrity, and the corresponding genetic knock out mutant displayed a no-growth phenotype^31^. *In vivo* mouse data upon oral dosing with Irosustat did not identify any significant effects on the colonic microbiota at the phylum or *Bacteroides* genus level. This may be as a result of rapid uptake of the drug in the stomach or small intestine. It will be important to identify sulfamates that can transit to the distal gut and study mice inoculated with HGM bacteria, which will allow studies of species level changes and alteration of membrane lipid composition. Interestingly, it has recently been shown that altered *B. theta* membrane lipid composition can lead to the transfer of bacterial lipids to the host cell membrane^40^.

This work reveals three findings with corresponding implications: (1) aryl and carbohydrate sulfamates and sulfonates are not effective inhibitors of *exo*-acting HGM S1 carbohydrate sulfatases. There are no known inhibitors of these enzymes, and this is an area where further research is needed. (2) Arylsulfamates have unexpected off-target effects with negative impacts on HGM *Bacteroides* species. Potentially, the removal of these mucin degrading bacteria could help treat IBD but the effects on the HGM may also lead to undesirable side-effects on the host. (3) Selected non-sulfated complex carbohydrates confer a resistant/protective phenotype on HGM *Bacteroides* species that can protect from the harmful effects of arylsulfamates. This could help to mitigate the effect of arylsulfamates (and potentially other drugs that transit it to the colon), opening the potential for paired drug-glycan co-treatment strategies.

## Acknowledgements

CJC was funded by MSCA grant MARINEGLYCAN (101029842) and acknowledges funding from the Max Planck Society. SVDP was supported by the Swedish Research Council (2020-02536). ASL was supported by Swedish Research Council grant (2021-01409) Swedish Society for Medical Research (Svenska Sällskapet för Medicinsk Forskning, grant S21-0026), Sahlgrenska Academy International Starting Grant (GU2021/1070), and Jeanssons Foundation grant. This project has received funding from the Academy of Medical Sciences/Wellcome Trust through the Springboard Grant (SBF005\1065 163470 awarded to AC), the Australian Research Council (DP210100235, DP240100126, DP250100819), the Royal Society (RGS\R2\212050), and the Wellcome Trust (career development award to AC, 225897/Z/22/Z).

## Author contributions

AC conceived the project. AC, CJC and SW wrote the manuscript. AC, CT, DPB, and ASL carried out enzymology experiments. AC, CT, and DPB analysed enzymological data. CJC, EAY, and CG and ZC synthesised compounds. SE carried out computational analyses. SVDP carried out and analysed proteomic experiments and data. AC, CT, CD, and DN carried out bacterial growth experiments. MC and TL carried out lipidomic analysis. All authors read and approved the manuscript.

## Data availability statement

Source Data for all experiments, along with corresponding statistical test values, where appropriate, are provided within the paper and in Supplementary information. The mass spectrometry data has been deposited to the ProteomeXchange consortium via PRIDE with the dataset identified PXDXXXXXX.

## Code availability statement

No new codes were developed or compiled in this study

## Methods

### Recombinant protein production

Catalytically-active recombinant S1 sulfatases (in which Ser, within the C/S-X/P/A-X-R consensus sequence, for *B*. *thetaiotaomicron* sulfatases was mutated to Cys, to permit conversion to the formylglycine (FGly) residue by *E. coli*^18^), or catalytically-inactive sulfatases (in which Cys in the consensus sequence is mutated to Ser, thus preventing formation of FGly) were expressed in the *E. coli* strain TUNER (Novagen). Cultures were grown to mid-exponential phase in LB media supplemented with 50 μg/mL kanamycin at 37 °C, in an orbital shaker set to 180 rpm. Cultures were then cooled to 16 °C, and recombinant gene expression was induced by the addition of 0.1 mM isopropyl β-D-1-thiogalactopyranoside for 16 h at 16 °C, and 180 rpm. Cells were collected by centrifugation at 5,000 × g and pellets were resuspended in 20 mM HEPES, pH 7.4, with 500 mM NaCl, then were sonicated on ice. Recombinant protein was then purified by immobilized metal ion affinity chromatography using a cobalt-based matrix (Talon, Clontech) and, after a wash with resuspension buffer, eluted with a step gradient of 10, 50, and then 100 mM imidazole in resuspension buffer. Proteins were then analysed by SDS-PAGE and appropriately pure fractions were pooled and dialysed into 10 mM HEPES pH 7.0 with 150 mM NaCl. Protein concentrations were determined by measuring absorbance at 280 nm using the molar extinction coefficient calculated by ProtParam on the ExPasy server (web.expasy.org/protparam/).

### Spectrophotometric based sulfatase assays

Production of *para*-nitrophenolate from *para*-nitrophenol sulfate by steroid sulfatases was monitored at *A*_400nm_ using a HIDEX sense or spectra max plus 384 (molecular devices) plate reader in a 96 well plate format. Reaction mixtures (100 μl) were monitored at ambient temperature (20-25 °C) and contained 1 mM substrate with 100 mM Bis-Tris-Propane, pH 7.0 with 5% (v/v) DMSO, 150 mM NaCl, and 5 mM CaCl_2_.

### Microfluidic-based de-sulfation assay

Reducing end BODIPY (maximal excitation/emission coefficient of ∼503/511 nm) labelled sulfated substrates were detected using the EZ Reader II platform (Ret biochem) *via* LED-induced fluorescence, as described previously^41^. Real-time kinetic evaluation of substrate desulfation was achieved using a non-radioactive microfluidic mobility shift carbohydrate sulfation assays were optimised in solution with a 12-sipper chip coated with CR8 reagent and performed using a PerkinElmer EZ Reader II system employing EDTA-based separation buffer. Pressure and voltage settings were adjusted manually (1.8 psi, upstream voltage: 2250 V, downstream voltage: 500 V) to afford optimal separation in the reaction mixture of the sulfated substrate and unsulfated glycan product, with a sample (sip) time of 0.2 s, and total assay times appropriate for the experiment. Individual de-sulfation assays were carried out at 28°C and were pre-assembled in a 384-well plate in a volume of 80 μl in the presence of substrate concentrations of 1 μM with 100 mM Bis-Tris-Propane, MES, or Tris, dependent on the pH optimum of the sulfatase being assayed, and 5% DMSO, 150 mM NaCl, 0.02% (v/v) Brij-35 and 5 mM CaCl_2_. The amount of de-sulfation was directly calculated in real-time using EZ Reader II software by measuring the sulfated carbohydrateLJ:LJunsulfated carbohydrate ratio at each time-point during the assay. The activity of sulfatase enzymes was quantified in ‘kinetic mode’ by monitoring the amount of unsulfated glycan generated over the assay time, relative to control assay with no enzyme; with desulfation of the substrate limited to ∼20% to prevent loss assay linearity via substrate depletion. *k*_cat_/*K*_M_ values, using the equation V_0_=(*k*_cat_/*K*_M_)/[E][S], were determined by linear regression analysis with GraphPad Prism software.

### Differential scanning fluorimetry

Thermal shift/stability assays (TSAs) were performed using a StepOnePlus Real-Time PCR system (LifeTechnologies) and SYPRO-Orange dye, at a 1:1000 dilution, (excitation 470 nm, emission maximum 570 nm, Invitrogen) with thermal ramping between 20 and 95°C in 0.3°C step intervals per data point to induce denaturation of purified, folded, inactive, wild-type (Ser77) BT1636^S1_20^ in the presence or absence of substrate or potential inhibitors. The melting temperature (Tm) corresponding to the midpoint for the protein unfolding transition was calculated by fitting the sigmoidal melt curve using the Boltzmann equation in GraphPad Prism, with R^2^ values of ≥0.99, as described in^41^. Data points after the fluorescence intensity maximum were excluded from the fitting. Changes in the unfolding transition temperature compared with the control curve (ΔT_m_) were calculated for each ligand. A positive ΔT_m_ value indicates that the ligand stabilises the protein from thermal denaturation, and confirms binding to the protein. All TSA experiments were conducted using a final protein concentration of 5 μM in 100 mM Bis-Tris-Propane (BTP), pH 7.0, and 150 mM NaCl, supplemented with the appropriate ligand concentration and 5 % DMSO. Three independent assays were performed for each protein and protein ligand combination.

### Bacterial growth experiments

Growth of various *Bacteroides* species was conducted in brain heart infusion media. BHI [Sigma 53286-500G], 37g per litre) was dissolved in 18.2Ω deionised water, and then sterilised by autoclaving. The media was then allowed to cool then haematin (1.2 mg/ml in 0.2 His-HCl pH 8.0) was added in a 1:1000 dilution. Five ml of the resulting BHI media was then inoculated with *Bacteroides* species and grown overnight, and diluted 1:20 into 96 well plate growths (10 μl in 200) to monitor growth. Growth of the *B. thetaiotaomicron* VPI-5482 on specified glycan sources was achieved by mixing 1:1 the 0.22 micron filter-sterilised polysaccharides (10.2 mg/ml in H_2_O) with 0.22 micron filter-sterilised 2 x minimal media (per 50 ml was: 0.1g of ammonium sulfate and sodium carbonate, 0.05g cysteine, 10 ml 1 M potassium phosphate pH 7.2, 0.1 ml 1 mg.ml^−1^ vitamin K, 1 ml of 0.4 mg.ml^−1^ iron sulfate, 0.4 ml of 0.25 mg.ml resazurin, 0.05 ml of 0.01 mg.ml^−1^ vitamin B12, 5 ml of mineral salts for defined media, and 100 μl of 1.2 mg/ml haematin in 0.2 M His-HCl pH8.0); this gives a final glycan concentration of 5 mg/ml in a 1 x minimal media. All growth experiments were monitored continuously in 96-well plates using a cerillo stratus plate reader within a Don whitely VA-500, A85, or A35 cabinet configured for anaerobic conditions (80% N_2_, 10% CO_2_, 10% H_2_). Where appropriate, either 1% DMSO (control growths) or 1% DMSO and 1 mM arylsulfamates was added to growth conditions. For sphingosine kinase inhibitor (SKI, 567731) and sphingosine kinase 1 inhibitor II (SK1-II, 567741) a maximum of 0.1 and 0.5 mM, respectively, could be dissolved in BHI with 1% DMSO. For diacylglycerol kinase inhibitors i (D5794-5MG) and ii (D5919-5MG) a maximum concentration of 125 μM and 62.5 μM, respectively, could be dissolved in BHI with 1% DMSO but were hazy; this did not affect measurement readings at OD_600nm_. Overnight growth cultures were diluted 1:20 into 96 well plate growths (10 μl in 200 μl). Growth curves presented are averages of three technical replicates.

### Thin layer chromatography (TLC)

*B. thetaiotaomicron* VPI-5482 was grown in 10 ml minimal media with 10 mg/ml (w/v) appropriate GAG (either CSA or Heparin) as the sole carbon source to mid-exponential phase in glass test tubes. Cells were harvested by centrifugation at 5,000 × g for 10 min at room temperature and washed 2 x with 5 ml PBS (pH 7.2) before being resuspended in 1 mL PBS. Cells washed in PBS were lysed and the lysates assayed against 5 mM of the appropriate substrate at 37 °C for up to 24 h. Assays were analysed by TLC, and 2 μL each sample was spotted onto silica plates and resolved in butanol:acetic acid:water (2:1:1) mobile phase. The plates were dried, and the sugars were visualized using diphenylamine stain (DPA: 1 ml of 37.5% HCl, 2 ml of aniline, 10 ml of 85% H_3_PO_3_, 100 ml of ethyl acetate, and 2 g of diphenylamine) by heating using a heat gun set to 450 °C on medium flow. Arylsulfates and their pNP products were visualised using 1 M NaOH and then heated as described for DPA stains.

### High performance anion exchange chromatography (HPAEC)

Sonicated bacterial cell lysates grown on the glycosaminoglycans CSA and heparin were treated with 1 mM of arylsulfamates 3 and 12 for 1 hour then challenged with CSA or heparin mono- and disaccharides by mixing 1:1, to a final concentration of 5 mM, to see if sulfatase activity was still present. Analysis of the sugar products was performed by HPAEC using an ICS-6000 with pulsed amperometic detection using a gold work electrode and pDH reference electrode with a standard quad carbohydrate waveform. Separation was done using a Dionex PA-200 analytical column preceded by a PA-200 guard column. Samples were resolved isocratically in 100 mM NaOH for 20 mins, then the column cleaned with 500 mM NaOH for 10 mins, and finally ran back into 100 mM NaOH for 5 mins, with a flow rate of 0.25 ml/min.

### Synthesis of chemical compounds

Full detail of the synthetic procedures can be found in the supporting information.

### Thermal proteome profiling

A 300 ml culture of minimal media (see ‘Bacterial growth experiments’ section) supplemented with 5 mg/ml chondroitin sulfate A (CSA), stored in the anaerobic chamber overnight, and inoculated with 10 ml of *B. thetaiotaomicron* VPI-5482 was grown overnight in BHI media. This culture was grown to a target OD_600nm_ of ∼0.5-0.6 at which point cells were centrifuged at 5000 x g and washed with PBS, this process was repeated two more times. Cells were then treated as in *Mateus et al*^27^; briefly, cells were then resuspended in lysis buffer (50μg/ml lysozyme, 1×protease inhibitor(Roche), 250 U/ml benzonase, and 1 mM MgCl_2_ in PBS) to give an OD_600nm_ of 50. Samples were then sonicated on ice for 15 seconds three times. The lysate was then split into three: one sample was a control (1% (v/v) DMSO) and experimental samples contained 1 mM arylsulfamate **3** or 1 mM arylsulfamate **12** (1% (v/v) DMSO). Next 20 μl of each condition was aliquoted into a 96 well PCR plate across 10 wells in quadruplicate. The plates were then subjected to a temperature gradient from 45 – 72 °C (45, 46.8, 49.1, 52.1, 55.9, 60.3, 64.4, 67.4, 69.9, 72°C) for 3 min, followed by 3 min at room temperature. NP-40 (nonyl phenoxypolyethoxylethanol) was then added to samples to a final concentration of 0.8% (v/v). Samples were transferred to a 0.22 μm 96 well filter plate (Millipore: MSGVS2210) and centrifuged at 500 x g for 5 min to remove protein aggregates. Following filtration, the flowthrough was combined 1:1 with 400 mM Tris pH 8.0 and 4% SDS (w/v). The protein concentration for each sample was determined by Bradford assay, and a volume corresponding to 10 µg for the lowest-temperature condition was used as input for all other samples in a set for analysis. Protein digestion was performed using a modified solid-phase-enhanced sample preparation (SP3) based method adapted to a 96 well format in 0.45µM filter plates (Millipore, MSRPN04)^42,43^. Premixed magnetic SpeedBeads (Cytiva, (1:1, GE45152105050250 and GE 65152105050250)) were added to the samples, combined with two volumes of ethanol and incubated for 10 min while mixing at 150 RPM. Samples were transferred to the filter plates and washed four times with 200 µL of 70% ethanol (v/v) with centrifugation for 2 min at 2000 x g between each wash. Beads were resuspended in 40 µL 0.1 mM HEPES pH 8.5 containing trypsin/Lys-c (Thermo Scientific), 1.25 mM TCEP and 5 mM chloroacetamide (Sigma-Aldrich) and incubated at room temperature overnight while mixing 500 rpm. Peptides were collected by centrifugation, followed by a second elution with 2% DMSO. TMT 10-plex labelling (Thermo Scientific) was performed at a 2:1 ratio for 1 hr while mixing at 500 RPM. The labelling reaction was quenched with 0.4% hydroxylamine (v/v) for 15 min before each sample was combined and dried under vacuum. Peptides were resolved in 0.1% trifluoroacetic acid (TFA) and fractionated using 0.1% triethylamine with increasing concentrations of acetonitrile by high pH reverse phase into 8 fractions using C18 desalting columns (Pierce, 89852) pooled into 4 fractions and dried under a vacuum.

Lyophilized samples were redissolved in 0.1% TFA and analysed by LC-MS/MS using a nano HPLC system (EASY-nLC 1200, Thermo Scientific) coupled to a Q-Exactive HF-X mass spectrometer (Thermo Scientific). Peptides were separated using in-house packed columns (150 x 0.075 mm) packed with Reprosil-Pur C18-AQ 3 μm particles (Dr. Maisch). Elution was performed with a 5 to 45% gradient (A: 0.1% formic acid, B: 0.1% formic acid, 80% acetonitrile) in 135 minutes at 250 nl. Full mass spectra were acquired over a mass range of minimum 400 m/z and maximum 1600 m/z, with a resolution of 60,000 at 200 m/z. The isolation window for fragmentation spectra collection was set to 1Da and a fixed first mass of 100 m/z with a resolution of 30,000 at 200 m/z. The top 15 most intense peaks with a charge state ≥2 – 5 were selected for fragmentation by HCD with a normalized collision energy of 32% and subsequent excluded for selection for 30 seconds. Thermo .RAW files were converted into MZml using msConvert (proteowizard) and FragPipe (v20.0) configured with Msfragger (v3.8) and Philosopher (v5.0)^44,45^ was used for protein identification and TMT quantification. Database searches were performed against the *Bacteroides thetaiotaomicron* reference proteome (4782 entries, UP000001414, 2023_10) with the following settings, parent ion mass and fragment ion mass accuracy after recalibration was set to 20 ppm. Enzyme specificity was set to trypsin, fixed modifications were set for cysteine carbamidomethylation and TMT modified lysine, variable modifications considered were methionine oxidation, protein n-terminal acetylation and TMT modification on peptide n-terminal or serine. The identified proteins and peptides were both filtered at a 1% and only identifications based on ≥2 unique peptides with a minimum ion purity of 0.75 were considered for quantification and thermal proteome profiling. Further data analysis was performed in R environment to combine the search outcome, normalize reporter ion channels. Meltome curve fitting analysis and statistical analysis were executed using the TPP-TR workflow in the TPP package^46^.Briefly, protein candidates with altered melting temperature were considered based on the following criteria. P-values for the two replicate experiments are <0.05 and <0.1, the melting point shifts in the vehicle versus treatment experiments have the same direction. Both melting point differences in the two pairs of control versus treatment experiments are greater than the melting point difference between the two vehicle controls and the minimum slope in each of the control versus treatment experiments was <0.06. In addition, proteins were only considered for melting curve analysis if the identification was based on a minimum of two unique peptide identification; see Dataset 1.

### Ligand docking experiments

All docking experiments were conducted using SwissDock^47^ on the Expasy server using default parameter; no binding site was specified. The Alphafold model of BT4322 (Q89ZQ4) and the crystal structure of SphK1 (3VZB) were used, and where appropriate heteroatoms removed, and uploaded to the SwissDock server as PDB files. Small molecules to be docked were built in Jligand^48^, the generated PDB or CIF files converted to MOL2 files using the openbabel server, and uploaded to SwissDock as MOL2 files.

### Lipidomic analysis

All solvents used were HPLC or LC-MS grade. To the cell pellet, 997 μL of 2:1 methanol : chloroform (MeOH:CHCl_3_) and 208 μL of 0.005 N HCl were added. The tubes were vortexed briefly and incubated on ice for 5 minutes. Following centrifugation at 18,000 xg for 5 minutes at 4°C the supernatant was transferred to a fresh tube. CHCl_3_ (332 μL)) and H_2_O (332 μL) were added and briefly vortexed to form two phases. After centrifugation at 3600 xg for 1 minute at 4°C, the lower organic phase was transferred to a LC-MS tapered vial and dried under a gentle stream of nitrogen. The samples were resuspended in 80 μL of 7:3 acetonitrile : isopropanol (ACN:IPA) for LC-MS analysis.

Untargeted lipid analysis was performed on bacterial pellets and data processed using R scripts as previously described^49^, except that data was collected on a Thermo Orbitrap Exploris 480 mass spectrometer with MS1 and HCD ddMS2 scans collected at 120,000 and 15,000 resolution (FWHM), respectively. The following parameters were used to acquire MS data: spray voltage 3500 V (positive mode), 2500 V (negative mode), sheath gas 50, auxiliary gas 10, sweep gas 1, ion transfer tube temperature 280°C, vaporizer temperature 200°C, RF lens 30, scan range m/z 270-2000. For MS2 parameters, mass selection range m/z 400-2000, ramped HCD collision energies 30, 50, 70%. MS1 m/z values were searched against candidate ions from positive and negative ionization mode adduct databases generated from the LipidMAPS structural database (https://www.lipidmaps.org/) and the list of Bacteroides lipids described in Barone *et al* 2024^40^. Candidate lipids were additionally annotated by searching MS2 spectra using LipidMatch^50^, LipidBlast^51^ and SIRIUS (version 6.1.0)^52^.

Annotations were manually curated using either Barone’s names, or the sum composition notation described in LipidMAPS. To distinguish between multiple isomers, annotations were prepended with a unique label indicating MS1 m/z values (“M”) and retention time in s (“T”). The feature list was filtered to remove background contaminants (cutoff = 3 standard deviations above average background feature area), and any feature where relative standard deviation exceeded 30%. Features that could not be annotated as lipids were removed and where there were redundant measurements across positive and negative ionization modes or adduct type, only the most abundant representative was kept. This resulted in a final feature list of 63 lipids, covering ceramides (CE), menaquinones, triglycerides (TG), phosphatidylethanolamines (PE), and various glycolipids. Feature area data was further processed through MetaboAnalyst 6.0 (https://www.metaboanalyst.ca) using the Statistical Analysis (one factor) workflow. Data was sample normalized by median before calculating fold-changes and log10 transformed before performing t-tests to calculate p-values.

### Mice experiments

All experimental procedures involving animals were approved by the Swedish Laboratory Animal Ethical Committee at the University of Gothenburg. 8-week-old C57BL/6N mice were maintained under standardized specific pathogen free conditions with ad libitum access to food and water. Mice were treated at day 0, 1 and 2 by gavage with 3.2 mM of Irosustat dissolved in PEG 400:water:DMSO (6:3:1). The control group was gavaged with the vehicle only. Fecal samples for all animals were collected at day 0, 3 and 7. The weight of the mice was recorded at the days the mice were gavaged or during fecal collection. Bacterial genomic DNA was extracted with QIAamp PowerFecalPro DNA kit (Qiagen^®^) and quantification by qPCR as described previously^53^. The following primers were used for relative quantification of phylum Bacteroidetes (Fw 5’-CRAACAGGATTAGATACCT; Rv 5’-GGTAAGGTTCCTCGCGTAT), phylum Firmicutes (Fw 5’-TGAAACTYAAAGGAATTGAGG; Rv 5’-ACCATGCACCACCTGTC) and genus Bacteroides (Fw 5’-GAAGGTCCCCCACATTG; Rv 5’-CGCKACTTGGCTGGTTCAG). The relative quantification normalized of specific taxons was determined for each sample using the 16S rRNA gene as reference.

