## Supplemental information for "Arylsulfamates inhibit colonic *Bacteroides* growth through lipid kinases and not sulfatases"

PNAS

**This PDF file includes:**

General procedure and analysis for chemical syntheses of compounds not previously reported
Pages 3-15

Figures S1 to S46
Pages33-80

Tables S1 to S7
Pages 81-89

Legends for Datasets S1 and S2, and SI References
Pages

### General procedures

All chemicals used were reagent grade and used as supplied unless otherwise noted. Compounds **19** and **22-24** were purchased from commercial suppliers. Analytical thin-layer chromatography (TLC) was performed on Merck silica gel 60 F254 plates (0.25 mm). Compounds were visualized by UV irradiation or dipping the plate in a 5% $H_2SO_4$ ethanol solution. Flash column chromatography was carried out on automated Grace flash chromatography system. Analysis and purification by normal and reverse phase HPLC were performed by using an Agilent 1200 series. Products were lyophilized using a Christ Alpha 2-4 LD plus freeze dryer. $^1H$, $^{13}C$ and HSQC NMR spectra were recorded on a Varian 400MR (400 MHz), Varian 600MR (600 MHz), or Bruker Biospin AVANCE700 (700 MHz) spectrometer. Signals are reported in terms of chemical shift [δ in parts per million (ppm)] relative to tetramethylsilane (TMS) or in $D_2O$ using the solvent as the internal standard in $^1H$ NMR (D2O: 4.79 ppm $^1H$). NMR data is presented as follows: Chemical shift, multiplicity (s = singlet, d = doublet, t = triplet, dd = doublet of doublet, m = multiplet and/or multiple resonances), coupling constant in Hertz (Hz), integration. All NMR signals were assigned on the basis of $^1H$ NMR, $^{13}C$ NMR, COSY, and HSQC experiments. High resolution mass spectra were obtained using a 6210 ESI-TOF mass spectrometer (Agilent) and a MALDI-TOF autoflexTM (Bruker). MALDI and ESI mass spectra were run on IonSpec Ultima instruments.

**a**

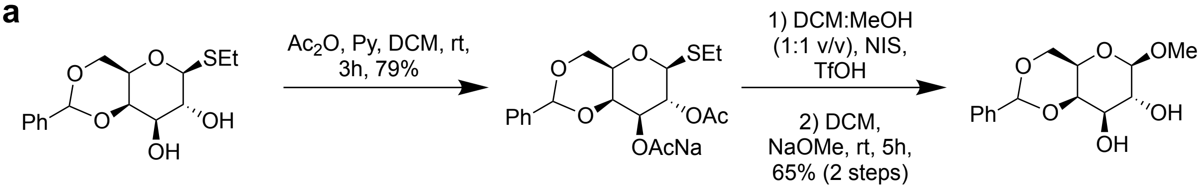


**b**

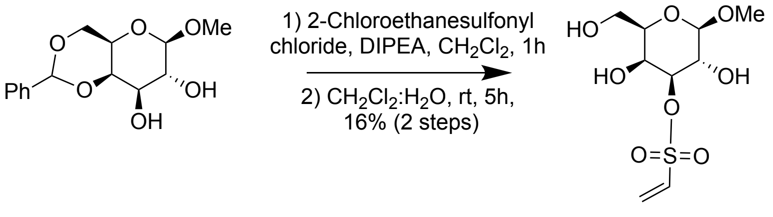


**c**

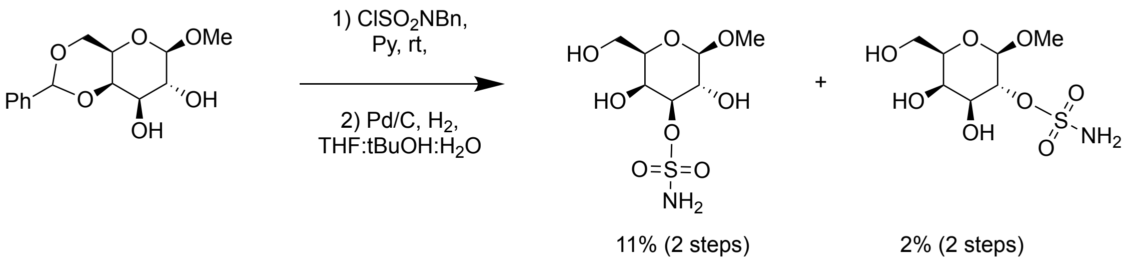


**SI Scheme 1. Synthesis of galactosides equipped with sulfonate and vinyl sulfones groups.** **a** synthesis of common ethyl 4,6-O-benzylidene-1-thio-β-D-galactoside intermediate**.** **b** synthesis of methyl 3-*O*-ethene-1-sulfonate-1-β-D-galactoside. **c** synthesis of methyl sulfonate galactosides.

**Synthesis of aryl sulfamates**

The following aryl sulfamates were prepared as described previously: 3-chlorophenyl sulfamate **1**,[1] *N*,*N*-dimethyl-4-nitrophenyl sulfamate **2**,[3] 4-phenylphenyl sulfamate **3**,[2] 4-nitrophenyl sulfamate **4**,[1] *N*,*N*-dimethyl-coumate **5**,[4] coumate **7**,[5] 4-methoxyphenyl sulfamate **17**,[1] 4-chlorophenyl sulfamate **18**,[1]

**General procedure for the synthesis of aryl sulfamates**

Chlorosulfonyl isocyanate (CSI, 1 equiv.) was cautiously added to a stirred, heated solution of the parent phenol (1 equiv.) in toluene (5-10 ml) at reflux. The solution was stirred at reflux overnight, then cooled to 0 °C. Water was added in a dropwise manner to the stirred, cooled toluene solution until evolution of gas ceased, resulting in the formation of a precipitate. Unless otherwise described, the precipitate was collected by filtration, washed (toluene), dried, and recrystallised (toluene) to afford the desired aryl sulfamate.

**General procedure for the synthesis of naphthyl sulfamates**

A mixture of formic acid (1.5 - 5 equiv.) and dimethylacetamide (DMA, 0.1 equiv.) was added to a stirred solution of CSI (1.5 - 5 equiv.) in $CH_2Cl_2$ at 40 °C. The mixture was refluxed for 15 min and cooled to r.t. then a solution of naphthol (1 equiv.) in DMA was added and stirred at r.t. for 17h. Water and EtOAc was added, the organic phase was collected and concentrated. Recrystallization or column chromatography afford the desired naphthyl sulfamate.

**HPLC purification**

**Method 1, Prep RP-HPLC**

Hypercarb column, 150 x 10 mm, 5 µm, flow rate of 3.5 mL/min with H2O (0.1% formic acid) as eluents [isocratic 100 % $H_2O$ (0.1% formic acid) (5 min), linear gradient to 100% ACN (30 min)].

**Method 2, analytical RP-HPLC**

Hypercarb column, 150 x 4.6 mm, 3 µm, flow rate of 0.7 mL/min with H2O (0.1% formic acid) as eluents [isocratic 100 % $H_2O$ (0.1% formic acid) (5 min), linear gradient to 100% ACN (30 min)].

### Compound Characterisation

### *N*-Methyl coumate (8)

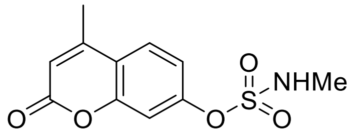

Freshly prepared *N*-methylsulfamoyl chloride (1.51 g, 11.65 mmol) was added to a stirred solution of 4-methyl-7-hydroxycoumarin (1.00 g, 4.66 mmol) in *N*,*N*-dimethylacetamide (10 mL) under an atmosphere of nitrogen. The solution was stirred under nitrogen (6 h). The reaction was quenched with water (100 mL), and the product extracted with ethyl acetate (3 × 30 mL), washed with water (3 × 30 mL) and dried ($MgSO_4$). The solvent was removed under reduced pressure, and the crude material subjected to flash chromatography (20% EtOAc: 80% pet. spirit). The product was repeatedly recrystallised (EtOAc: pet. spirit) due to the continual reappearance of an impurity to afford the *N*-methylsulfamate as cubic colourless crystals (794 mg; 59% yield). **m.p.** 171 °C. **$^1$H NMR** (500 MHz, $d_6$-DMSO) δ 2.44 (s, 3H), 2.75 (d, *J* = 4.7 Hz, 3H), 6.41 (s, 1H), 7.31 (dd, *J* = 8.7, 2.2 Hz, 1H), 7.35 (d, *J* = 2.2 Hz, 1H), 7.86 (d, *J* = 8.7 Hz, 1H), 8.45 (q, *J* = 4.4 Hz, 1H). **$^{13}$C NMR** (126 MHz, $d_6$-DMSO) δ 18.2, 29.2, 109.8, 114.1, 118.1, 118.2, 126.9, 152.0, 152.8, 153.6, 159.5.

### 4-Methoxynaphthalen-1-yl sulfamate (10)

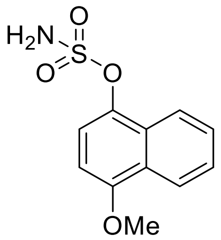


4-Methoxy-1-naphthol (390 mg, 2.24 mmol) and CSI (1.59 g, 11.2 mmol), column chromatography from 40% EtOAc/petrol (with 2% $Et_3N$) yielded the sulfamate (162 mg, 29%). **m.p.** 135 °C. **$^1$H NMR** (500 MHz, $d_6$-DMSO) δ 3.99 (s, 3H), 6.99 (d, *J* = 8.5 Hz, 1H), 7.43 (d, *J* = 8.4 Hz, 1H), 7.57 (ddd, *J* = 8.2, 6.9, 1.3 Hz, 1H), 7.63 (ddd, *J* = 8.3, 6.9, 1.3 Hz, 1H), 8.06 (s, 2H), 8.09 (d, *J* = 8.0 Hz, 1H), 8.18 (d, *J* = 8.2 Hz, 1H). **$^{13}$C NMR** (126 MHz, $d_6$-DMSO) δ 55.9, 103.6, 118.6, 121.6, 122.1, 125.4, 126.1, 127.1, 127.8, 139.4, 153.1.

### 7-Methoxynaphthalen-2-yl sulfamate (13)

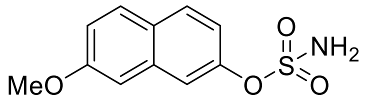


7-Methoxy-2-naphthol (390 mg, 2.24 mmol) and CSI (1.59 g, 11.2 mmol), column chromatography from 40% EtOAc/petrol (with 2% $Et_3N$) yielded the sulfamate (509 mg, 90%). **m.p.** 149 °C. **$^1$H NMR** (500 MHz, $d_6$-DMSO) δ 3.88 (s, 3H), 7.18 (dd, *J* = 8.9, 2.4 Hz, 1H), 7.26 (dd, *J* = 8.8, 2.2 Hz, 1H), 7.37 (d, *J* = 2.2 Hz, 1H), 7.71 (d, *J* = 2.0 Hz, 1H), 7.87 (d, *J* = 9.0 Hz, 1H), 7.92 (d, *J* = 8.8 Hz, 1H), 8.04 (s, 2H). **$^{13}$C NMR** (126 MHz, $d_6$-DMSO) δ 55.3, 106.0, 118.2, 118.8, 119.0, 126.8, 129.2, 129.4, 134.8, 148.4, 158.0.

### 6-Methoxynaphthalen-2-yl sulfamate (14)

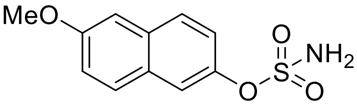

6-Methoxy-2-naphthol (300 mg, 1.72 mmol) and CSI (1.21 g, 8.61 mmol), column chromatography from 40% EtOAc/petrol (with 2% $Et_3N$) yielded the sulfamate (250 mg, 57%). **m.p.** 153–154 °C. **$^1$H NMR** (500 MHz, $d_6$-DMSO) δ 3.88 (s, 3H), 7.22 (dd, *J* = 8.9, 2.4 Hz, 1H), 7.44–7.35 (m, 2H), 7.74 (d, *J* = 1.9 Hz, 1H), 7.88 (dd, *J* = 11.7, 9.1 Hz, 2H), 7.99 (s, 2H). **$^{13}$C NMR** (126 MHz, $d_6$-DMSO) δ 55.3, 106.0, 119.2, 119.5, 122.0, 128.3, 129.2, 132.7, 146.1, 157.4.

### 4-Bromonaphthalen-1-yl sulfamate (11)

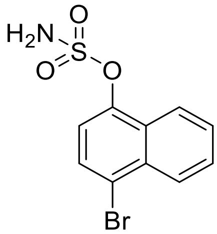

4-Bromo-1-naphthol (250 mg, 1.12 mmol) and CSI (793 mg, 5.6 mmol), column chromatography from 40% EtOAc/petrol (with 2% $Et_3N$) yielded the sulfamate (78 mg, 23%). **m.p.** 157 °C. **$^1$H NMR** (500 MHz, $d_6$-DMSO) δ 7.47 (d, *J* = 8.2 Hz, 1H), 7.76 (dt, *J* = 15.1, 7.1 Hz, 2H), 7.97 (d, *J* = 8.2 Hz, 1H), 8.18 (d, *J* = 8.3 Hz, 1H), 8.22 (d, *J* = 8.3 Hz, 1H), 8.28 (s, 2H). **$^{13}$C NMR** (126 MHz, $d_6$-DMSO) δ 119.1, 119.3, 122.9, 126.6, 127.8, 128.3, 128.7, 129.7, 132.1, 146.0.

### 3-Methoxynaphthalen-2-yl sulfamate (17)

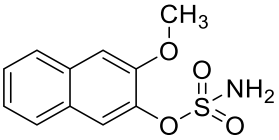

3-Methoxy-2-naphthol (390 mg, 2.24 mmol) and CSI (1.59 g, 11.2 mmol), column chromatography from 40% EtOAc/petrol (with 2% $Et_3N$) yielded the sulfamate (295 mg, 52%). **m.p.** 127–129 °C. **$^1$H NMR** (500 MHz, $d_6$-DMSO) δ 3.92 (s, 3H), 7.40 (t, *J* = 7.6 Hz, 1H), 7.51–7.47 (m, 2H), 7.87–7.81 (m, 3H), 8.03 (s, 2H). **$^{13}$C NMR** (126 MHz, $d_6$-DMSO) δ 55.9, 108.0, 120.4, 124.3, 126.4, 127.4, 127.6, 132.3, 139.4, 150.6.

### 6-Bromonaphthalen-2-yl sulfamate (15)

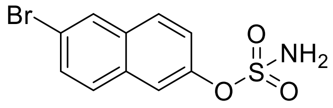

6-Bromo-2-naphthol (500 mg, 2.24 mmol) and CSI (475 mg, 3.36 mmol), recrystallization from $CHCl_3$/Petrol yielded the sulfamate (524 mg, 77%). **m.p.** 186 °C. **$^1$H NMR** (500 MHz, $d_6$-DMSO) δ 7.49 (dd, *J* = 8.9, 2.3 Hz, 1H), 7.70 (dd, *J* = 8.8, 1.9 Hz, 1H), 7.86 (d, *J* = 2.1 Hz, 1H), 7.96 (d, *J* = 8.8 Hz, 1H), 8.02 (d, *J* = 9.0 Hz, 1H), 8.10 (s, 2H), 8.28 (d, *J* = 1.4 Hz, 1H). **$^{13}$C NMR** (126 MHz, $d_6$-DMSO) δ 119.3, 119.3, 122.8, 129.1, 129.6, 129.8, 130.0, 131.9, 132.4, 148.2.

### 7-Methoxynaphthalen-1-yl sulfamate (12)

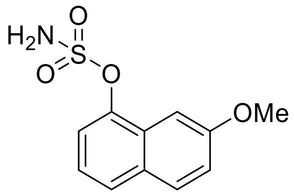

7-Methoxy-1-naphthol (150 mg, 0.86 mmol) and CSI (609 g, 4.3 mmol), column chromatography from 40% EtOAc/petrol (with 2% $Et_3N$) yielded the sulfamate (65 mg, 30%). **m.p.** 148–149 °C. **$^1$H NMR** (500 MHz, $d_6$-DMSO) δ 3.91 (s, 3H), 7.24 (dd, *J* = 8.9, 2.2 Hz, 1H), 7.38 (t, *J* = 7.9 Hz, 1H), 7.43 (d, *J* = 1.9 Hz, 1H), 7.48 (d, *J* = 7.6 Hz, 1H), 7.81 (d, *J* = 8.1 Hz, 1H), 7.91 (d, *J* = 9.0 Hz, 1H), 8.16 (s, 2H). **$^{13}$C NMR** (126 MHz, $d_6$-DMSO) δ 55.3, 100.4, 118.9, 119.3, 123.1, 126.1, 128.2, 129.6, 129.9, 145.3, 157.9.

### 6-Cyanonaphthalen-2-yl sulfamate (18)

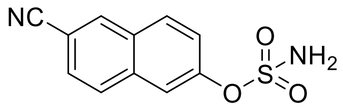

6-Cyano-2-naphthol (300 mg, 1.77 mmol) and CSI (1.25 g, 8.86 mmol), column chromatography from 30% EtOAc/Petrol (with 2% AcOH) yielded the sulfamate (210 mg, 48%). **m.p.** 198 °C. **$^1$H NMR** (500 MHz, $d_6$-DMSO) δ 7.60 (dd, *J* = 8.9, 1.5 Hz, 1H), 7.85 (d, *J* = 8.6 Hz, 1H), 7.97 (s, 1H), 8.23–8.15 (m, 4H), 8.64 (s, 1H). **$^{13}$C NMR** (126 MHz, $d_6$-DMSO) δ 108.55, 119.02, 119.30, 123.41, 127.24, 129.32, 130.15, 130.76, 134.26, 134.98, 150.10.

### Ethyl 4,6-*O*-benzylidene-1-thio-β-D-galactoside (26)

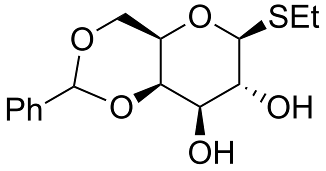

Purchased from GlycoUniverse.

**$^1$H NMR** (400 MHz, MeOD) δ 7.45 – 7.36 (m, 2H), 7.29 – 7.17 (m, 3H), 5.47 (s, 1H), 4.29 (d, *J* = 9.3 Hz, 1H), 4.12 (dd, *J* = 3.5, 1.1 Hz, 1H), 4.09 – 3.95 (m, 2H), 3.64 – 3.51 (m, 1H), 3.49 (dd, *J* = 9.4, 3.5 Hz, 1H), 3.44 (q, *J* = 1.5 Hz, 1H), 2.76 – 2.53 (m, 2H), 1.20 (t, *J* = 7.4 Hz, 3H). **$^{13}$C NMR** (101 MHz, MeOD) δ 139.8, 129.9, 129.0, 127.6, 102.4, 86.9, 77.8, 75.1, 71.4, 70.6, 70.4, 49.6, 49.5, 49.4, 49.3, 49.2, 49.1, 49.0, 48.9, 48.8, 48.6, 48.4, 24.5, 15.6.

### Ethyl 2,3-di-*O*-acetyl-4,6-*O*-benzylidene-1-thio-β-D-galactoside (27)

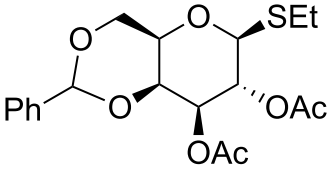

Ethyl 4,6-*O*-benzylidene-1-thio-β-D-galactoside **26** (5 g, 16.0 mmol) was suspended in $CH_2Cl_2$ (0.1M, 160 mL), pyridine (5.2 ml, 64.0 mmol, 4 eq.), and acetic acid (3.0 mL, 32.0 mmol, 2 eq.). Subsequently, a catalytic quantity of 4-(dimethylamino)pyridine (20 mg, 0.16 mmol, 0.01 eq.) was added. The mixture was left to stir at room temperature for 5 hours, and concentrated under vacuum to afford a yellow gel. Purification by flash chromatography ($SiO_2$, Hexane/Ethyl acetate) yielded title compound **27** (5.0 g, 79%) as a white amorphous solid. $R_f$ = 0.35 (Hexane/Ethyl acetate, 80:20, v/v).

**$^1$H NMR** (400 MHz, $CDCl_3$) δ 7.46 – 7.37 (m, 2H), 7.36 – 7.25 (m, 3H), 5.45 – 5.35 (m, 2H), 4.91 (dd, *J* = 10.0, 3.5 Hz, 1H), 4.39 (d, *J* = 9.9 Hz, 1H), 4.34 (df, *J* = 3.6, 1.0 Hz, 1H), 4.26 (dd, *J* = 12.5, 1.7 Hz, 1H), 3.94 (dd, *J* = 12.5, 1.7 Hz, 1H), 3.49 (q, *J* = 1.4 Hz, 1H), 2.89 – 2.74 (m, 1H), 2.66 (dq, *J* = 12.2, 7.5 Hz, 1H), 2.00 (d, *J* = 4.1 Hz, 6H), 1.27 – 1.16 (m, 3H). **$^{13}$C NMR** (101 MHz, $CDCl_3$) δ 170.8, 169.6, 137.6, 129.3, 128.3, 126.5, 101.3, 82.8, 73.7, 73.1, 69.8, 69.2, 66.6, 22.9, 21.0, 21.0, 14.9. **HRMS** calcd. $C_{19}H_{24}NaO_7S$ for $[M+Na]^+$ 419.1140, found 419.1151.

### Methyl 4,6-*O*-benzylidene-β-D-galactoside (28)

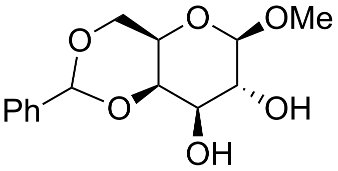


Ethyl 2,3-di-*O*-acetyl-4,6-*O*-benzylidene-1-thio-β-D-galactoside **27** (3.2 g, 8 mmol) was dissolved in anhydrous $CH_2Cl_2$ (0.05M, 160 mL), methanol (3.2 mL, 60 mmol, 10 eq) and cooled to 0°C. Next, NIS (3.6 g, 16 mmol, 10 eq) and TMSOTf (0.14 mL, 1.60 mmol, 0.2 eq) were added. Once complete, the reaction was quenched by the addition of triethylamine, filtered and washed with 10% aqueous $Na_2S_2O_3$, saturated aqueous $NaHCO_3$, and brine. The combined organic layer was dried over $MgSO_4$, filtered, concentrated, and purified by flash silica column chromatography ($SiO_2$, Hexane/EtOAc) to afford the intermediate glycoside. Next the methyl 2,3-di-*O*-acetyl-4,6-*O*-benzylidene-β-D-galactoside was dissolved in anhydrous $CH_2Cl_2$ (0.05M, 160 mL), cooled to 0°C, and sodium methoxide (0.5M) was added until the reaction reached pH 8. The temperature was allowed raised to room temperature and stirred complete (≈5 h). The reaction was tracked using thin layer chromatography (TLC) and once complete, it was quenched by the addition of Amberlite™ IR120(H), filtered, and concentrated. Purification by flash chromatography ($SiO_2$, $CH_2Cl_2$/Methanol) afforded **28** (1.6g, 65%) as an amorphous solid. $R_f$ = 0.3 ($CH_2Cl_2$/Methanol, 90:10, v/v).

**$^1$H NMR** $^1$H NMR (400 MHz, MeOD) δ 7.56 – 7.51 (m, 2H), 7.34 (d, *J* = 6.9 Hz, 2H), 5.60 (d, *J* = 1.5 Hz, 1H), 4.28 – 4.11 (m, 4H), 3.63 – 3.59 (m, 1H), 3.55 (s, 3H). **$^{13}$C NMR** (101 MHz, MeOD) δ 139.7, 129.9, 129.0, 129.0, 127.6, 127.5, 105.8, 102.5, 77.5, 73.7, 72.0, 70.2, 68.0, 57.4. **HRMS** calcd. $C_{14}H_{19}O_6$ for $[M+H]^+$ 283.1182, found 283.1210.

### Methyl 3-*O*-(ethenylsulfonyl)-β-D-galactoside (25)

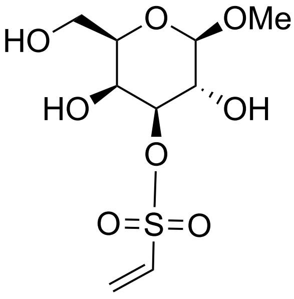


Methyl 4,6-*O*-benzylidene-β-D-galactoside (25 mg, 0.089 mmol) was dissolved in $CH_2Cl_2$ (0.1M, 1.0 mL) and 2-chloroethanesulfonyl chloride (10 µL, 0.095 mmol, 1.1 eq) and *N*,*N*-diisopropylethylamine

(45 µL, 0.267 mmol, 3 eq) were added. After 10 mins the reaction mixture was concentrated and the crude residue was purified by flash chromatography on silica gel (Hexane/ethyl acetate) to provide vinyl sulfonate intermediate. The crude material was dissolved in $CH_2Cl_2$ (25 mL) and 0.1M HCl (25 mL) was added. The reaction was stirred at room temperature until complete hydrolysis of the benzylidene acetal. The compound was then purified by HPLC (HyberCarb column, method 1) to isolate compound **25** (4 mg, 16 %)

**$^1$H NMR** (400 MHz, $D_2O$) δ 6.80 (dd, *J* = 16.6, 10.1 Hz, 1H), 6.48 (d, *J* = 16.6 Hz, 1H), 6.30 (d, *J* = 10.0 Hz, 1H), 4.53 (dd, *J* = 9.9, 3.4 Hz, 1H), 4.36 (d, *J* = 7.9 Hz, 1H), 4.17 (d, *J* = 3.4 Hz, 1H), 3.83 – 3.64 (m, 4H), 3.53 (s, 3H). **$^{13}$C NMR** (101 MHz, $D_2O$) δ 132.4, 131.4, 103.1, 83.1, 74.4, 68.3, 67.2, 60.5, 57.2. **HRMS** QTOF-MS: calcd. $C_9H_{16}NaO_8S$ for $[M+Na]^+$ 307.0464, found 307.0601.

### Methyl 3-*O*-sulfamoyl-β-D-galactoside (20)

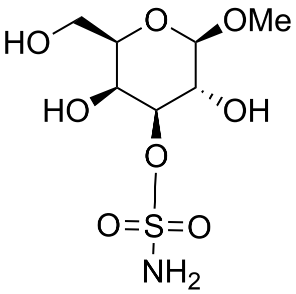


Methyl 4,6-O-benzylidene-β-D-galactoside (45 mg, 0.6 mmol) was dissolved in and pyridine (0.1M, 6 mL). Then *N*-benzyl sulfamoyl chloride (30 µL, 0.175 mmol, 1.1 eq) was added to the reaction.[6] Once complete, the reaction was quenched by the addition of methanol, concentrated and purified by column chromatography ($SiO_2$, Hexane/Ethyl acetate) to afford the 3-*O*-sulfonamide intermediate compound. At this stage also the 2-*O*-sulfonamide was separated. Next the 3-*O*-sulfonamide intermediate was dissolved in THF:t-BuOH:$H_2O$ (3 mL, 60:10:30, v/v/v) and 5% Pd/C (200 mg) was added. The reaction put under an atmosphere of hydrogen and left overnight. The reaction was then filtered through a pad of Celite®, and concentrated. The material was then purified by reverse-phase HPLC (HyberCarb column, method 1) to yield the title compound **23** (5 mg, 11%).

**$^1$H NMR** (400 MHz, $D_2O$) δ 4.51 (dd, *J* = 9.9, 3.4 Hz, 1H), 4.41 (d, *J* = 7.9 Hz, 1H), 4.28 (dd, *J* = 3.4, 0.9 Hz, 1H), 3.83 – 3.65 (m, 4H), 3.57 (s, 3H). **$^{13}$C NMR** (101 MHz, $D_2O$) δ 103.2, 82.1, 74.5, 68.5, 66.8, 60.6, 57.2. **HRMS** QTOF-MS: calcd. $C_7H_{15}NNaO_8S$ for $[M+Na]^+$ 296.0416, found 296.0431.

### Methyl 2-*O*-sulfamoyl-β-D-galactoside (21)

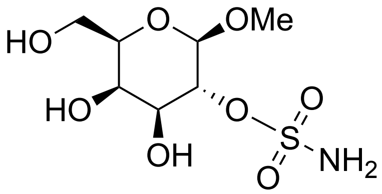


The 2-*O*-sulfamate intermediate was dissolved in THF:t-BuOH:$H_2O$ (3 mL, 60:10:30, v/v/v) and 5% Pd/C (200 mg) was added. The reaction left overnight under an atmosphere of hydrogen. The suspension was filtered through a pad of Celite®, and concentrated. The material was purified by reverse-phase HPLC (HyberCarb column, method 1) to yield the title compound **24** (1 mg, 2.3%).

**$^1$H NMR** (400 MHz, $D_2O$) δ 4.51 (d, *J* = 7.7 Hz, 1H), 4.31 (td, *J* = 9.8, 7.9 Hz, 1H), 3.96 (d, *J* = 3.5 Hz, 1H), 3.84 (dd, *J* = 9.8, 3.4 Hz, 1H), 3.79 – 3.65 (m, 3H), 3.54 (s, 4H).**$^{13}$C NMR** (101 MHz, $D_2O$) δ 101.3, 81.1, 75.1, 70.8, 68.9, 60.7, 57.2. **HRMS** calcd. $C_7H_{15}NNaO_8S$ for $[M+Na]^+$ 296.041, found 296.0429.

### NMR Spectra

#### 4-Methoxynaphthalen-1-yl sulfamate (10)

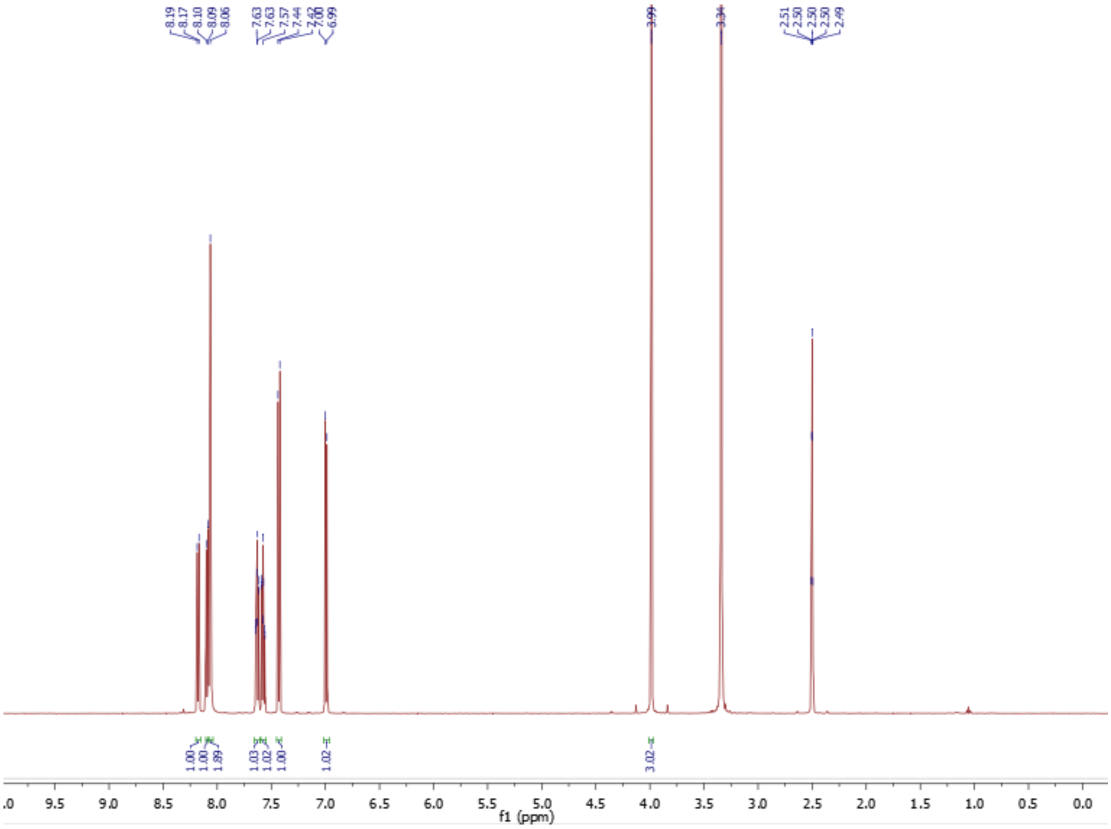


$^{1}H$ NMR

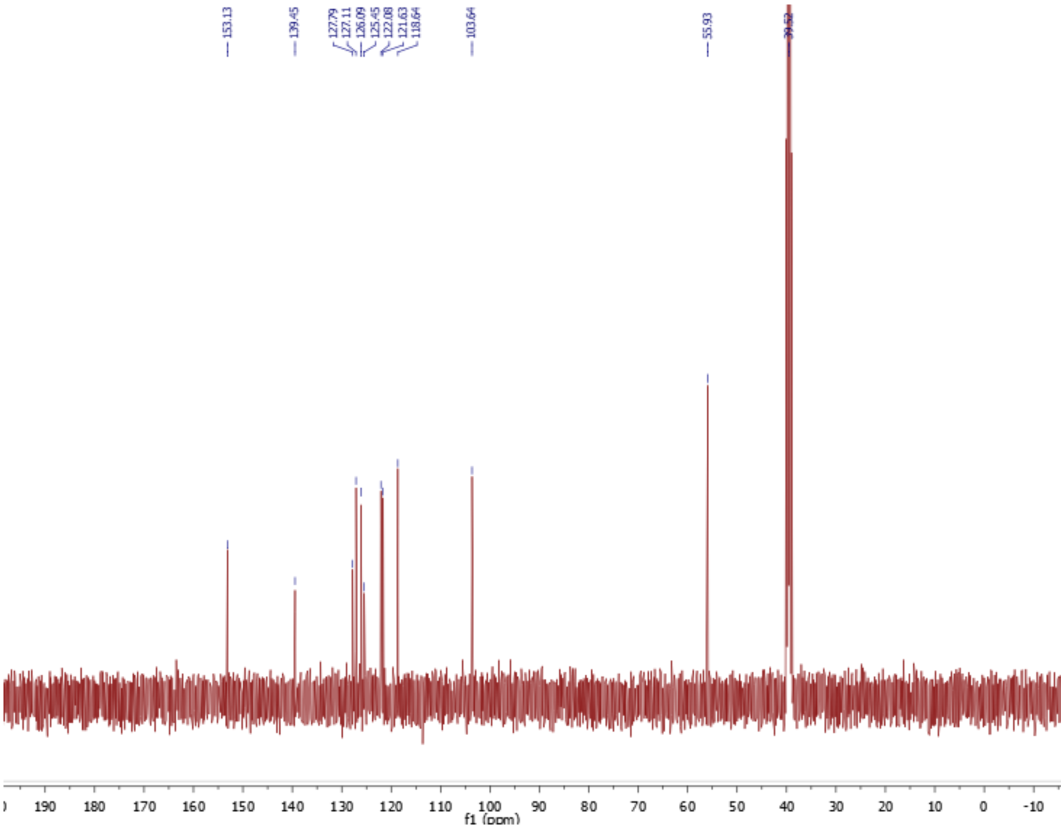


$^{13}C$ NMR

#### 4-Bromonaphthalen-1-yl sulfamate (11)

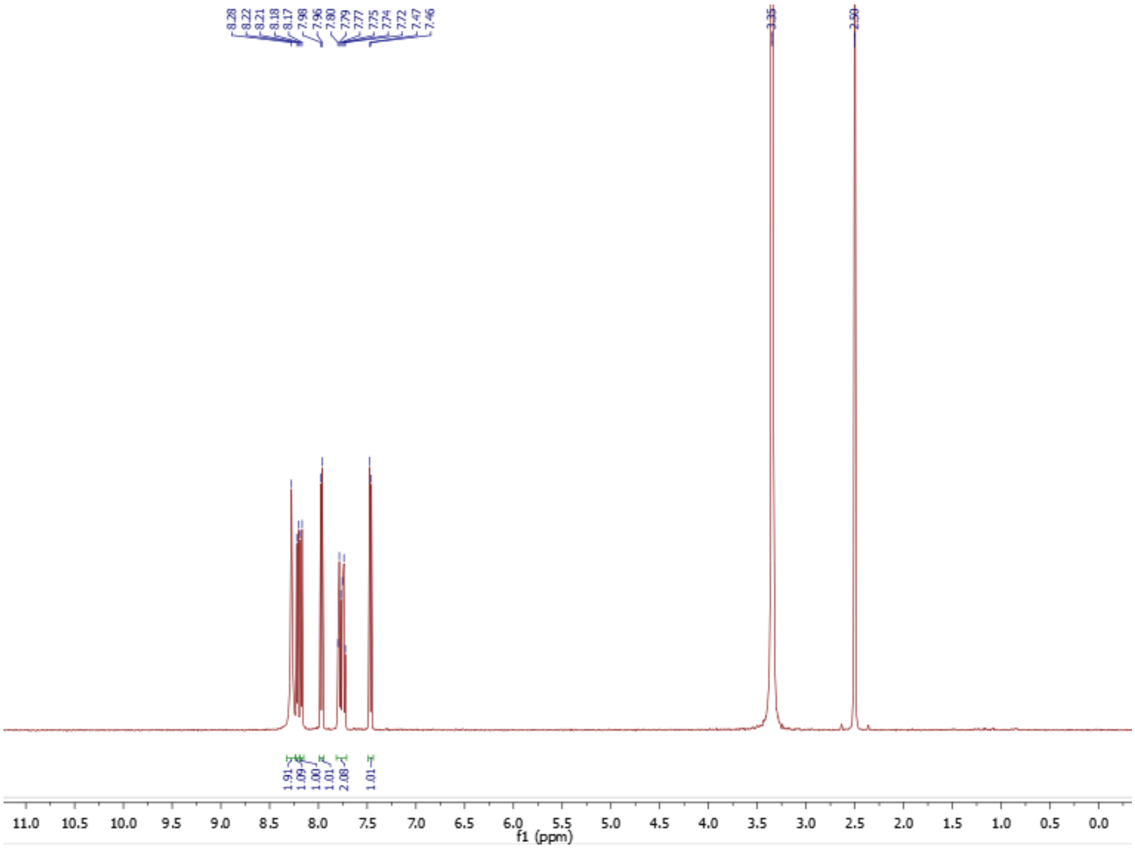


**[1]H NMR**

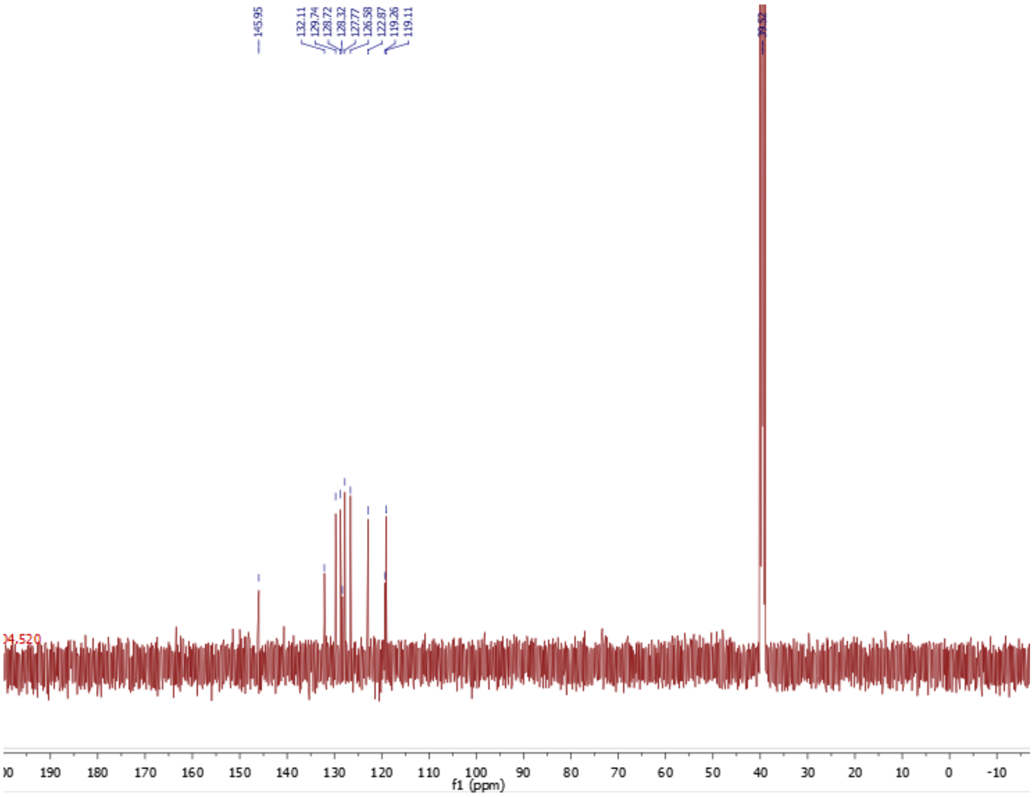


**[13]C NMR**

**7-Methoxynaphthalen-1-yl sulfamate (12)**

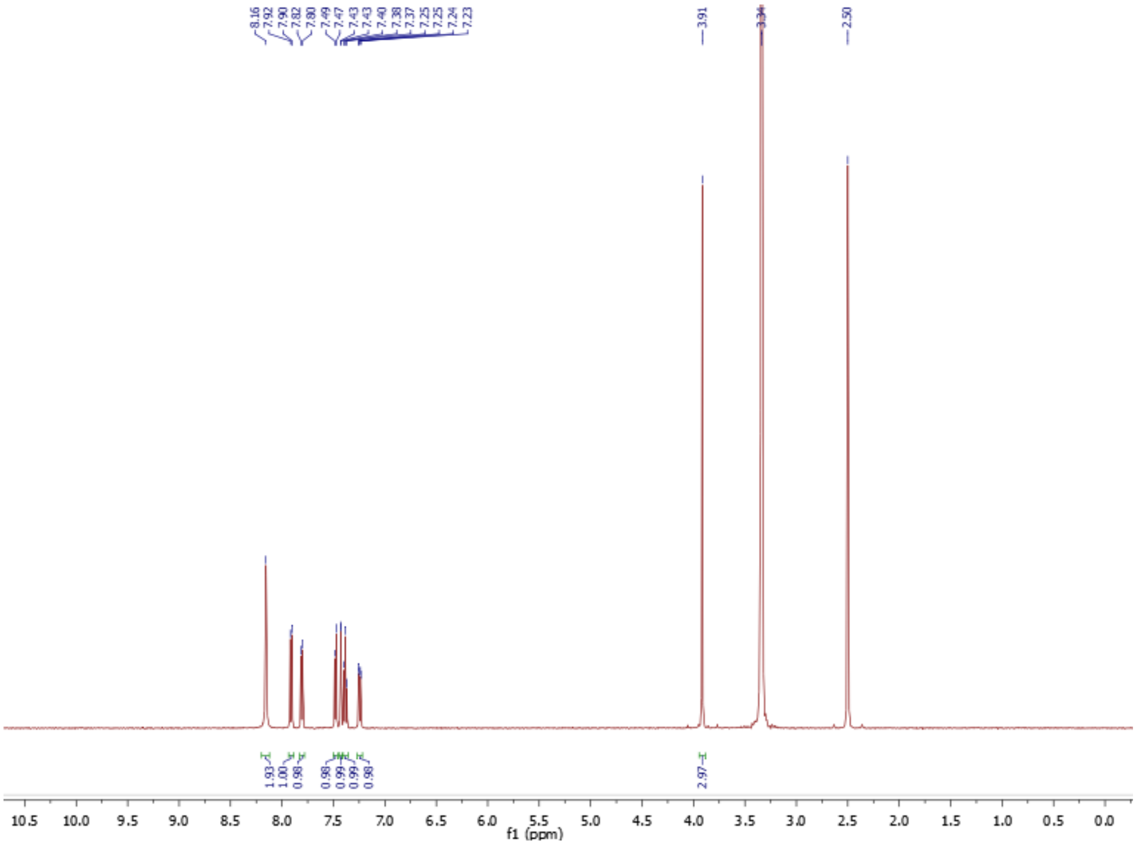


$^{1}$H NMR

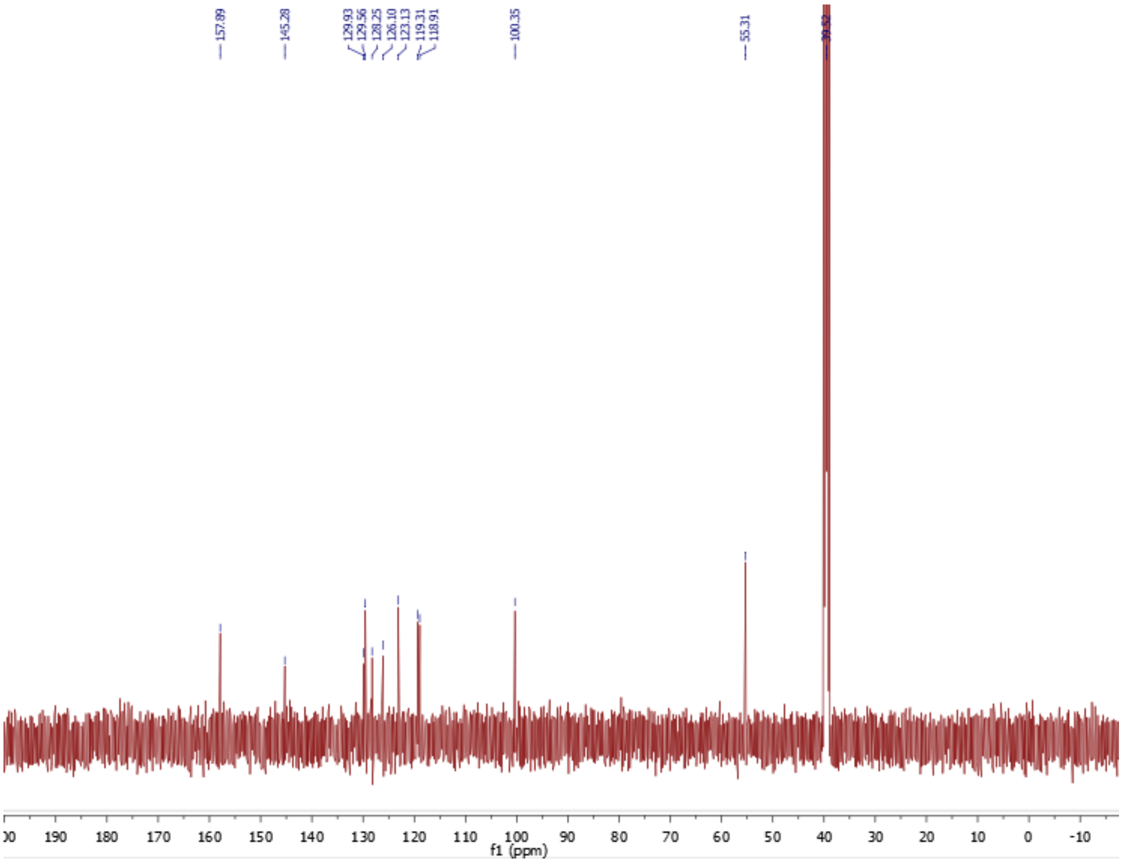


$^{13}$C NMR

**7-Methoxynaphthalen-2-yl sulfamate (13)**

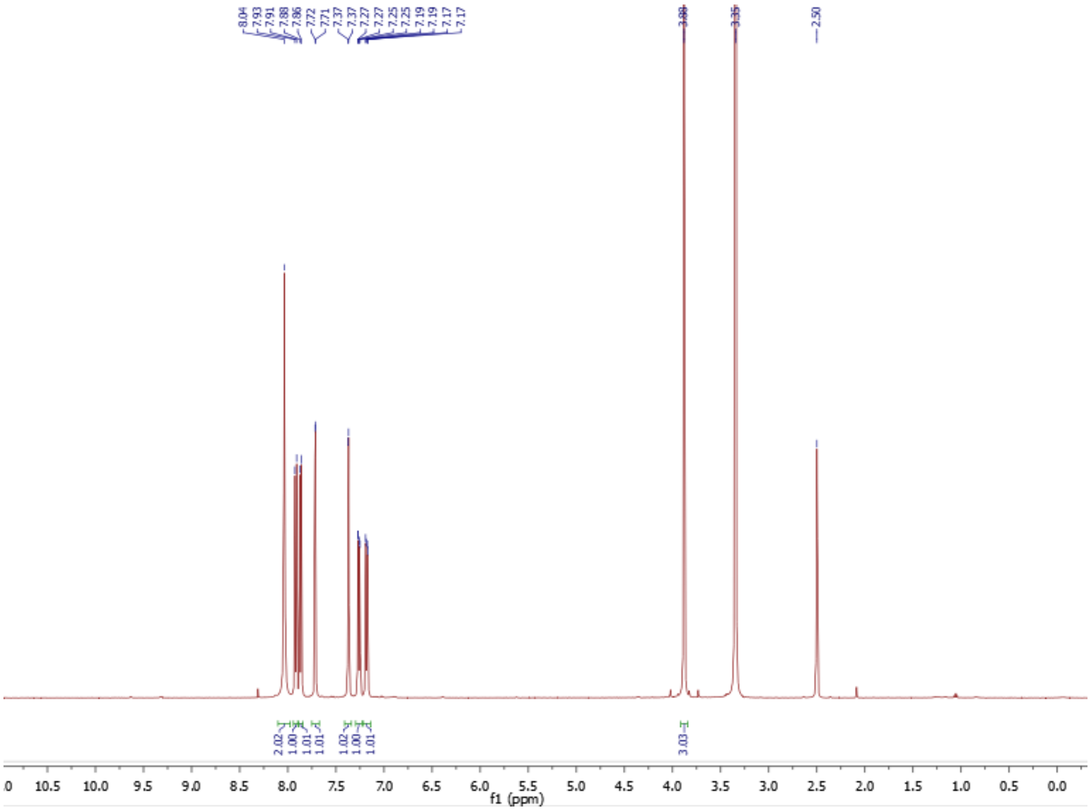

[1]H NMR

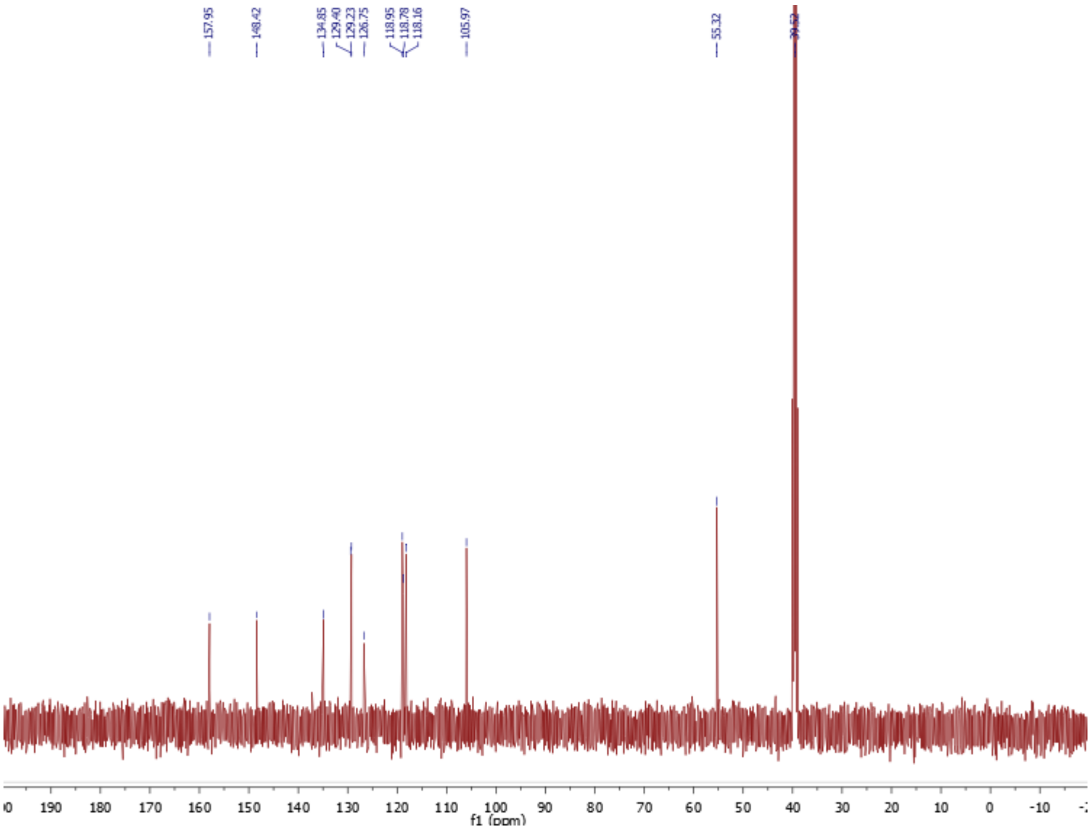

[13]C NMR

**6-Methoxynaphthalen-2-yl sulfamate (14)**

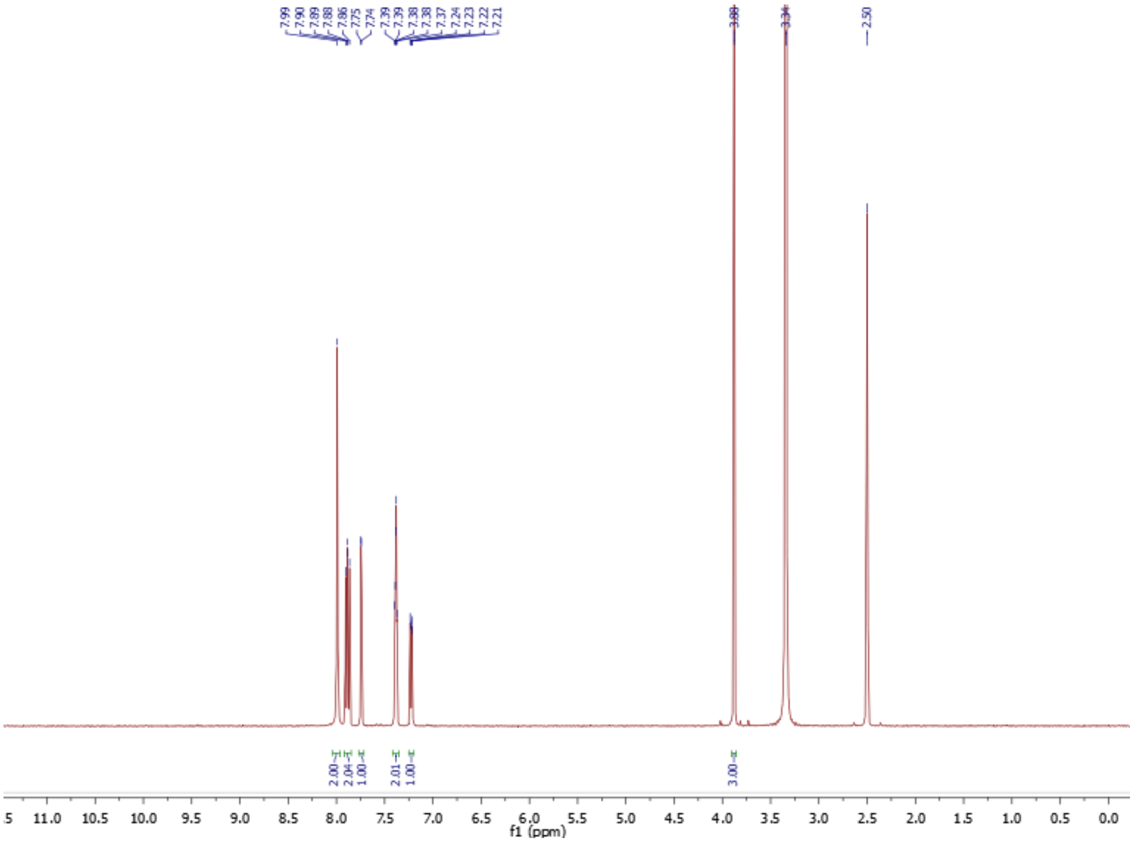


$^{1}H$ NMR

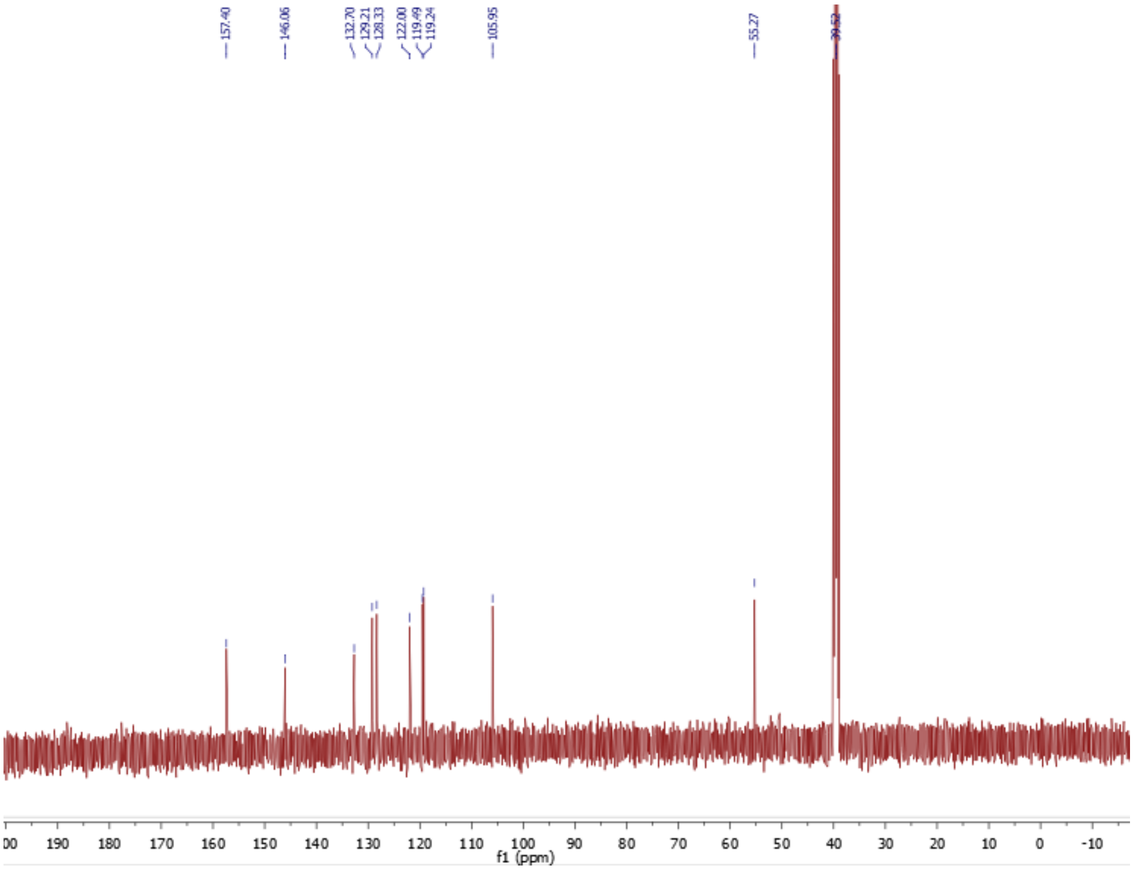


$^{13}C$ NMR

**3-Methoxynaphthalen-2-yl sulfamate (17)**

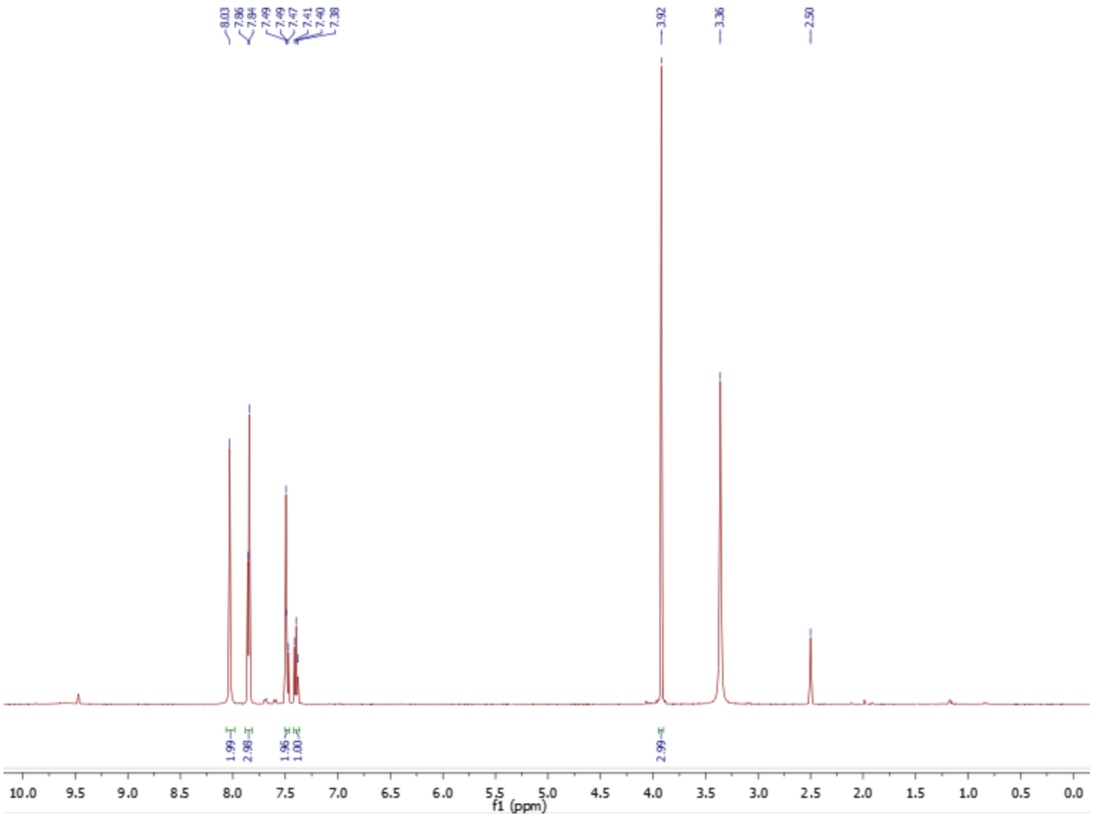


[1]H NMR

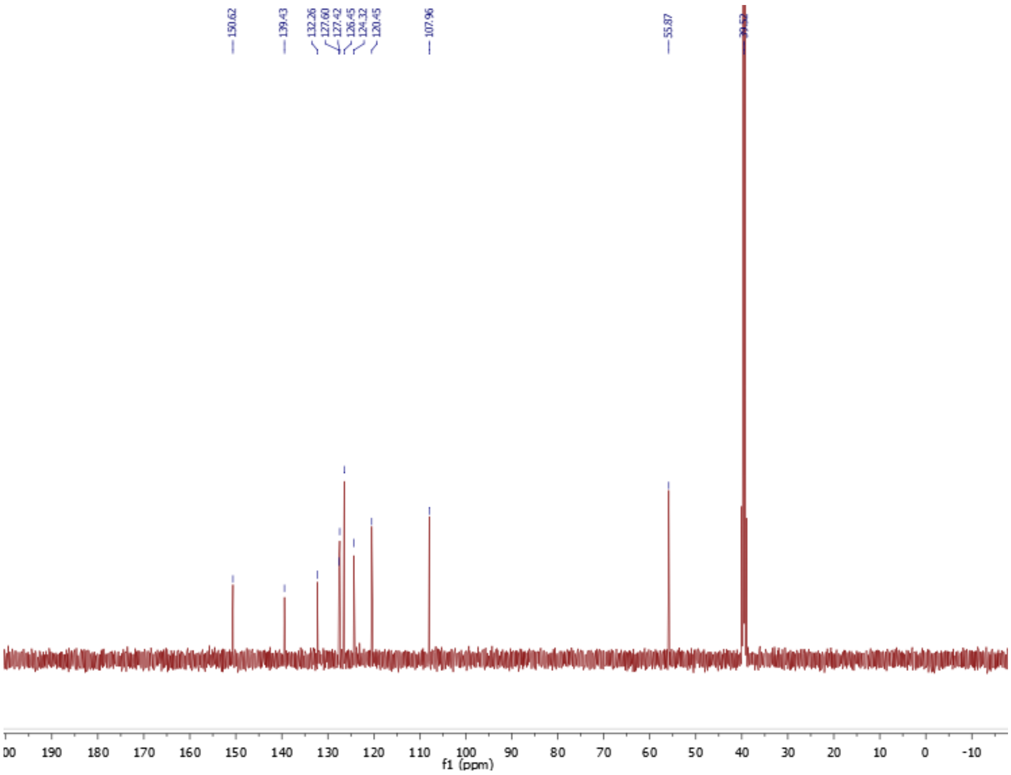


[13]C NMR

### 6-Bromonaphthalen-2-yl sulfamate (15)

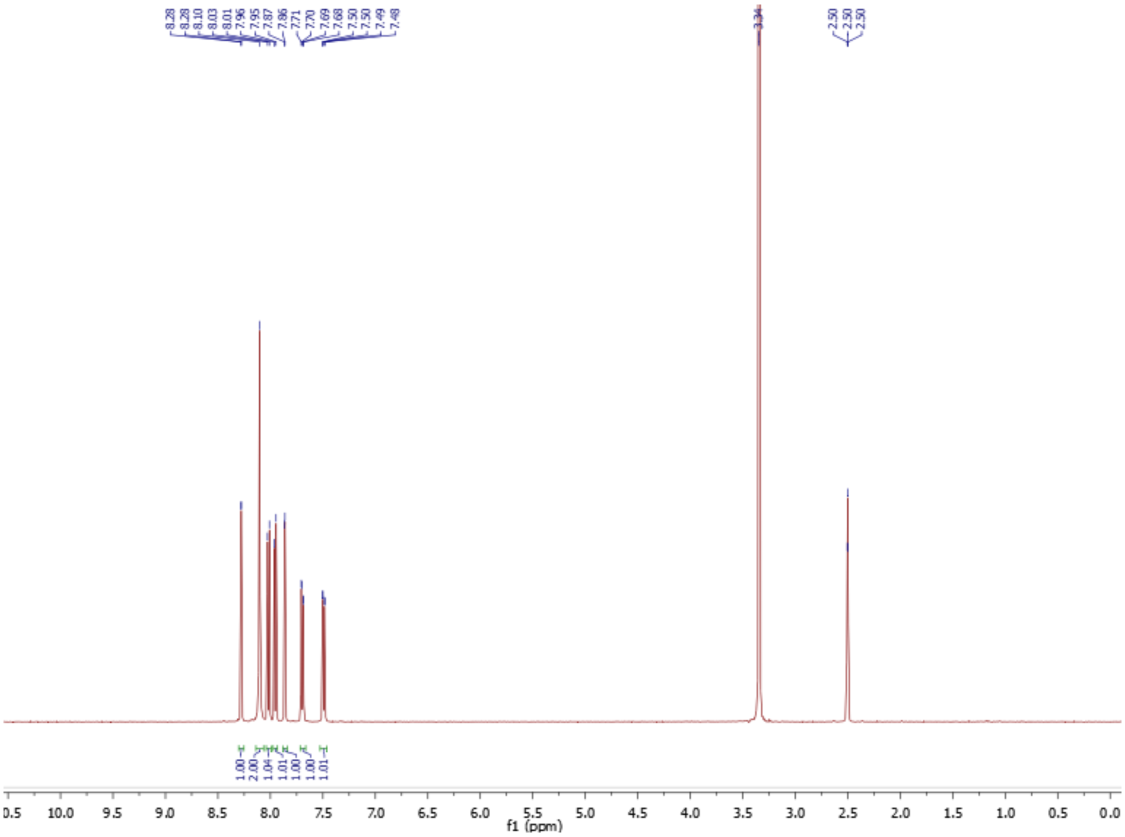

$^{1}$H NMR

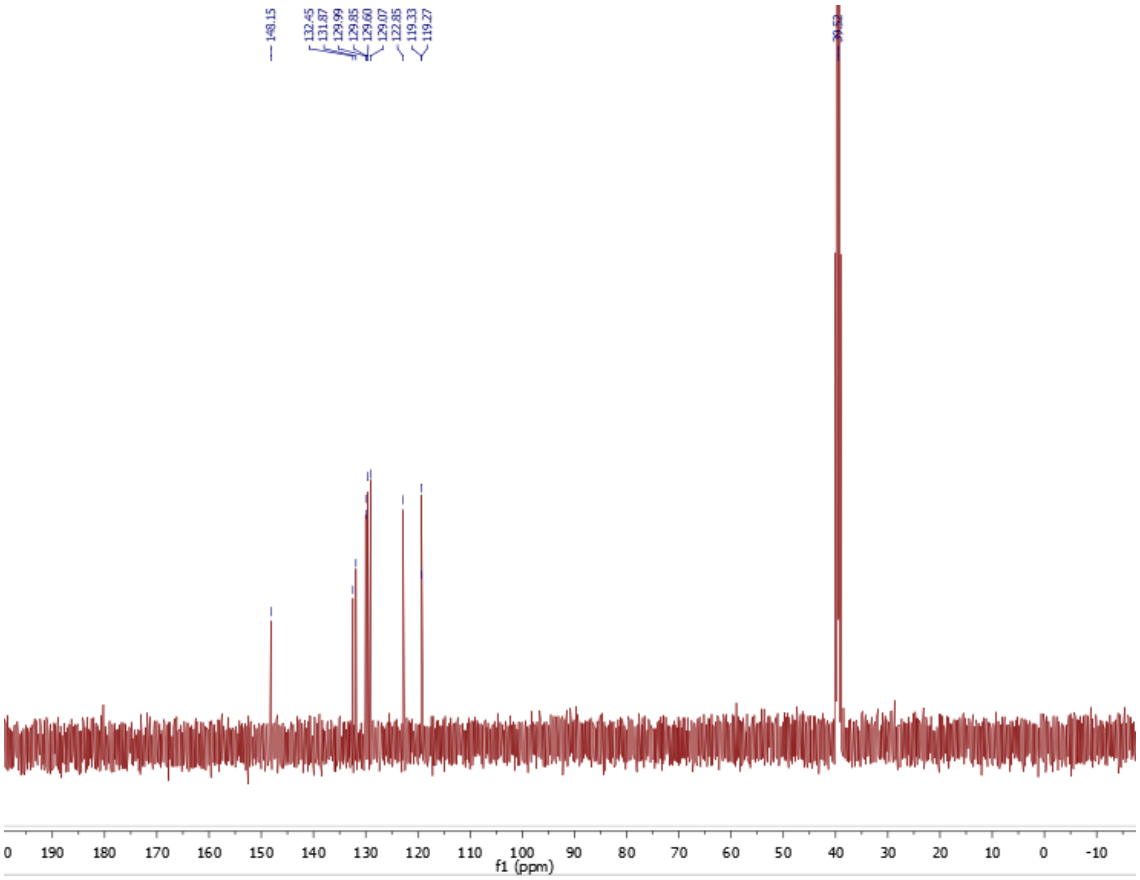

$^{13}$C NMR

### 6-Cyanonaphthalen-2-yl sulfamate (18)

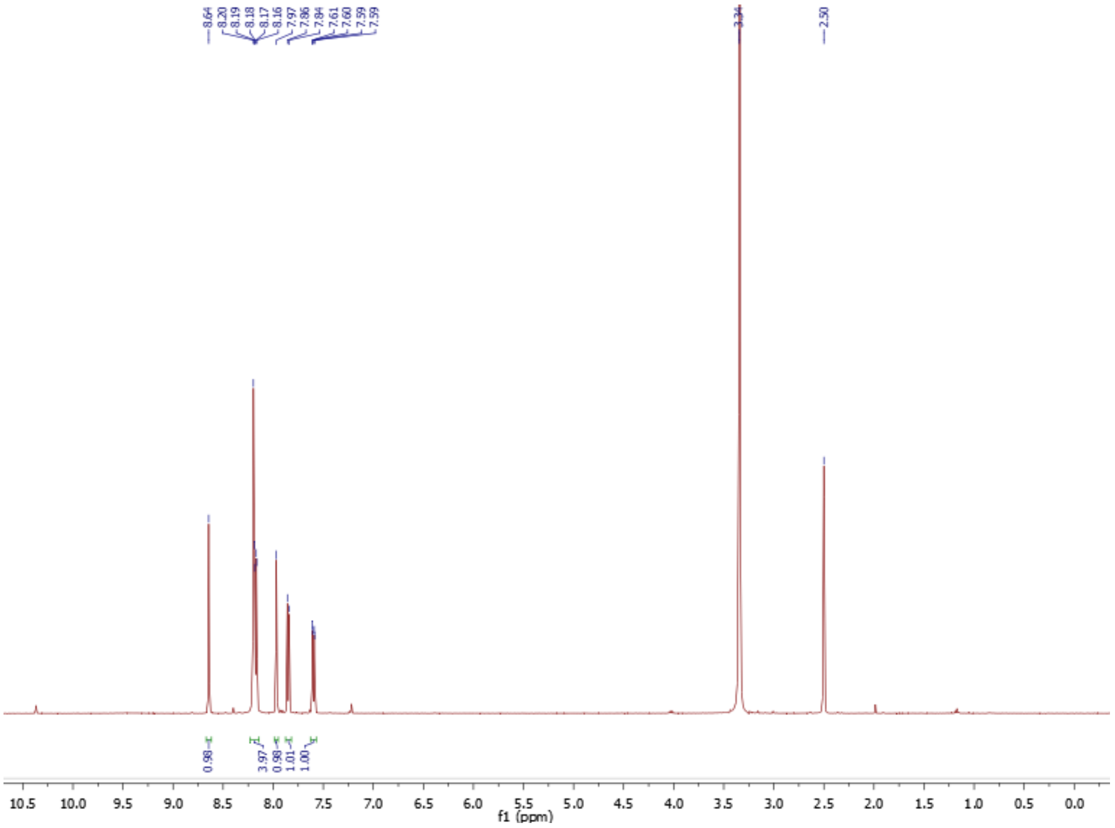


**[1]H NMR**

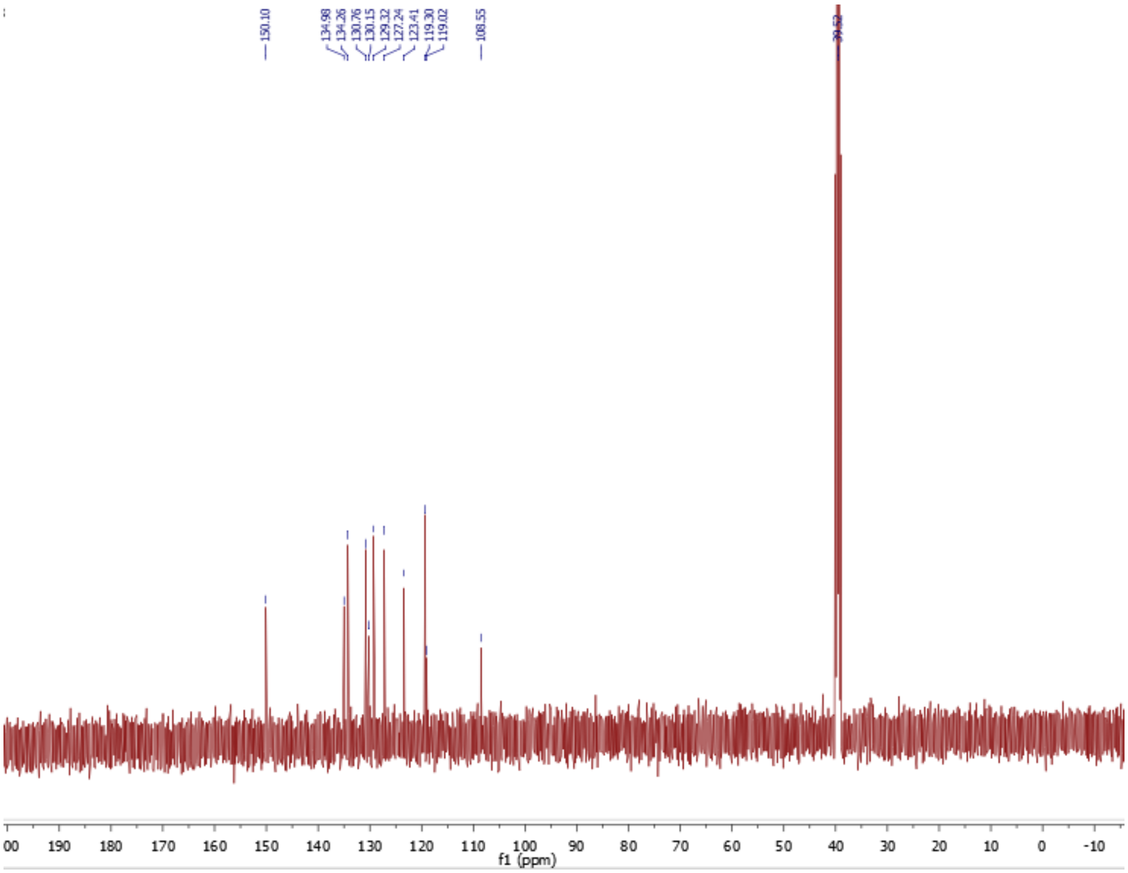


**[13]C NMR**

***N*-Methyl COUMATE (8)**

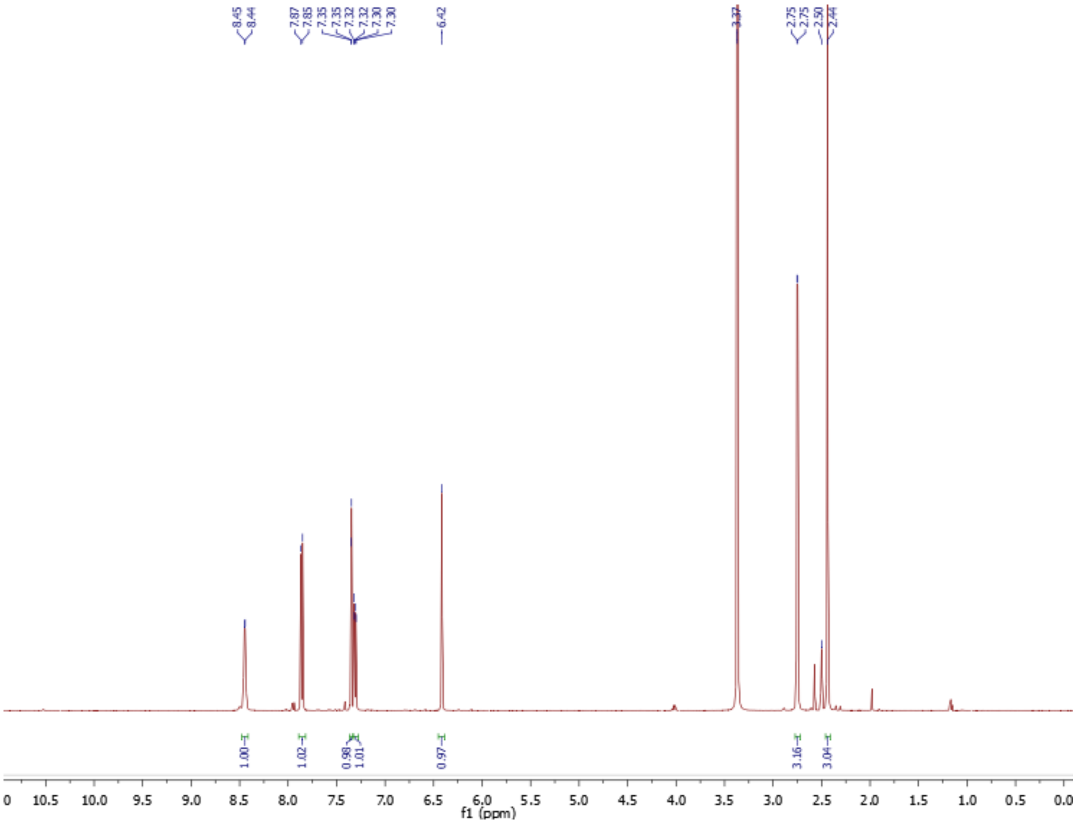

**$^1$H NMR**

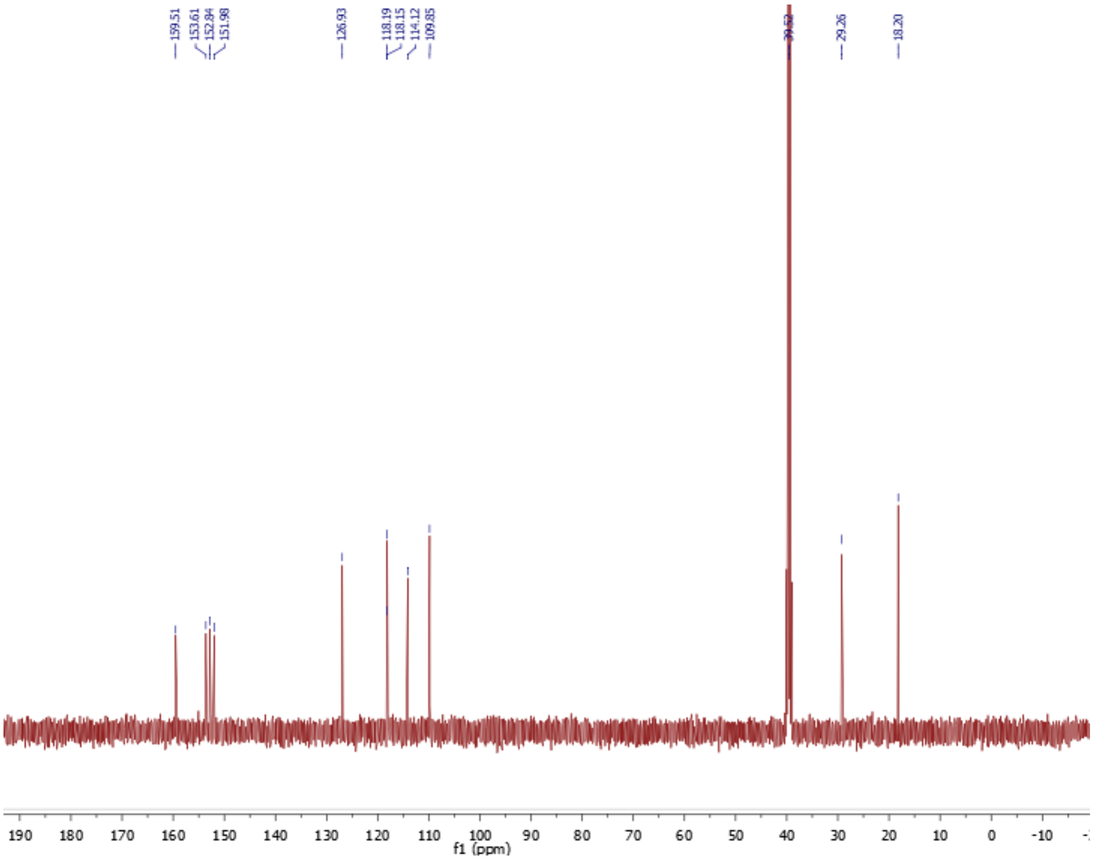

**$^{13}$C NMR**

**Methyl 3-*O*-sulfamoyl-β-D-galactoside (20)**

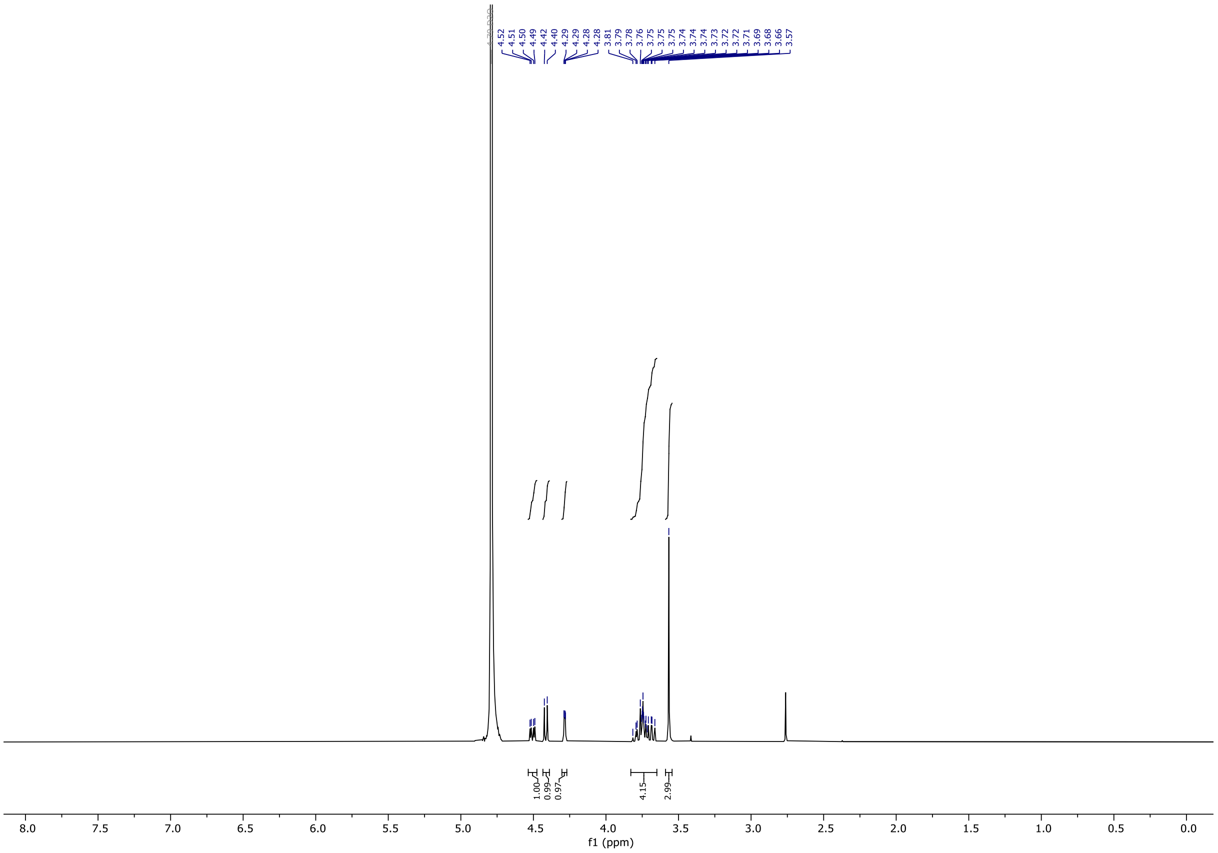

**$^{1}H$ NMR**

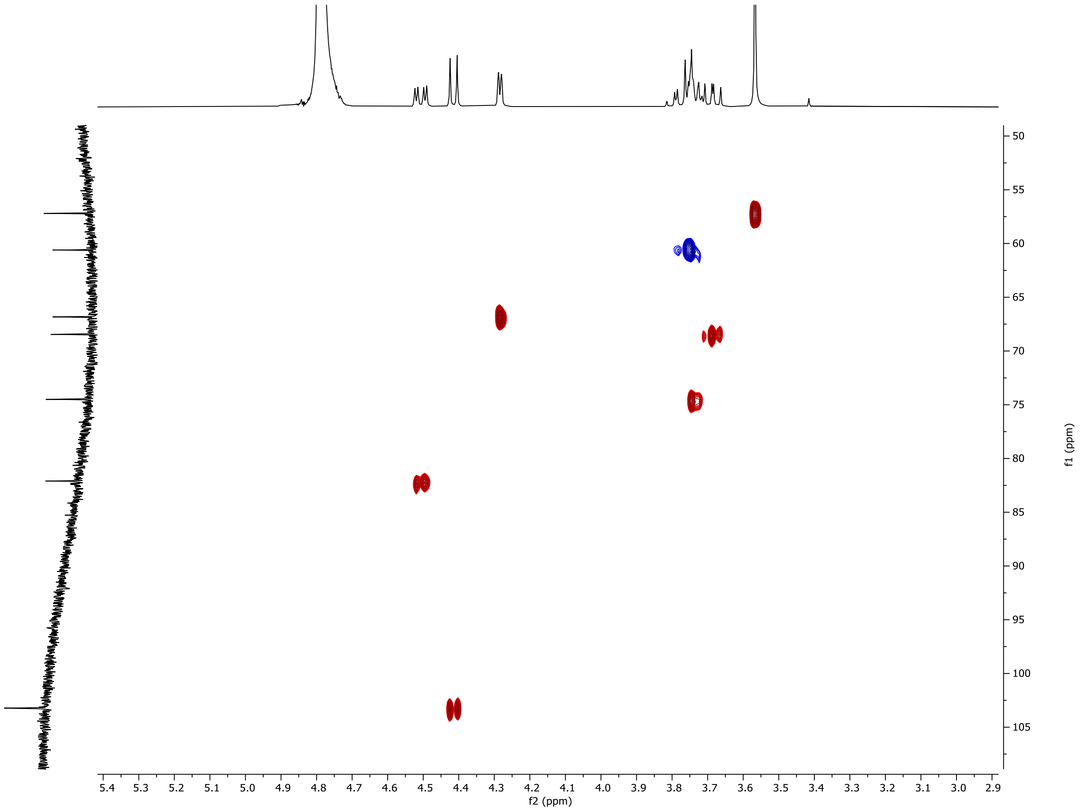

**$^{1}H$-$^{13}C$ NMR**

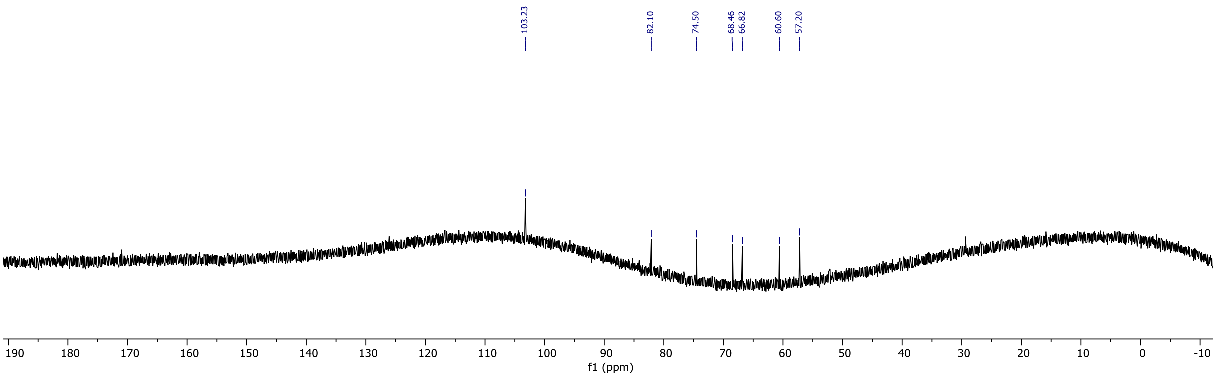

103.23
82.10
74.50
68.46
66.82
60.60
57.20
190 180 170 160 150 140 130 120 110 100 90 80 70 60 50 40 30 20 10 0 -10
f1 (ppm)


**$^{13}C$ NMR**

**Methyl 2-*O*-sulfamoyl-β-D-galactoside (21)**

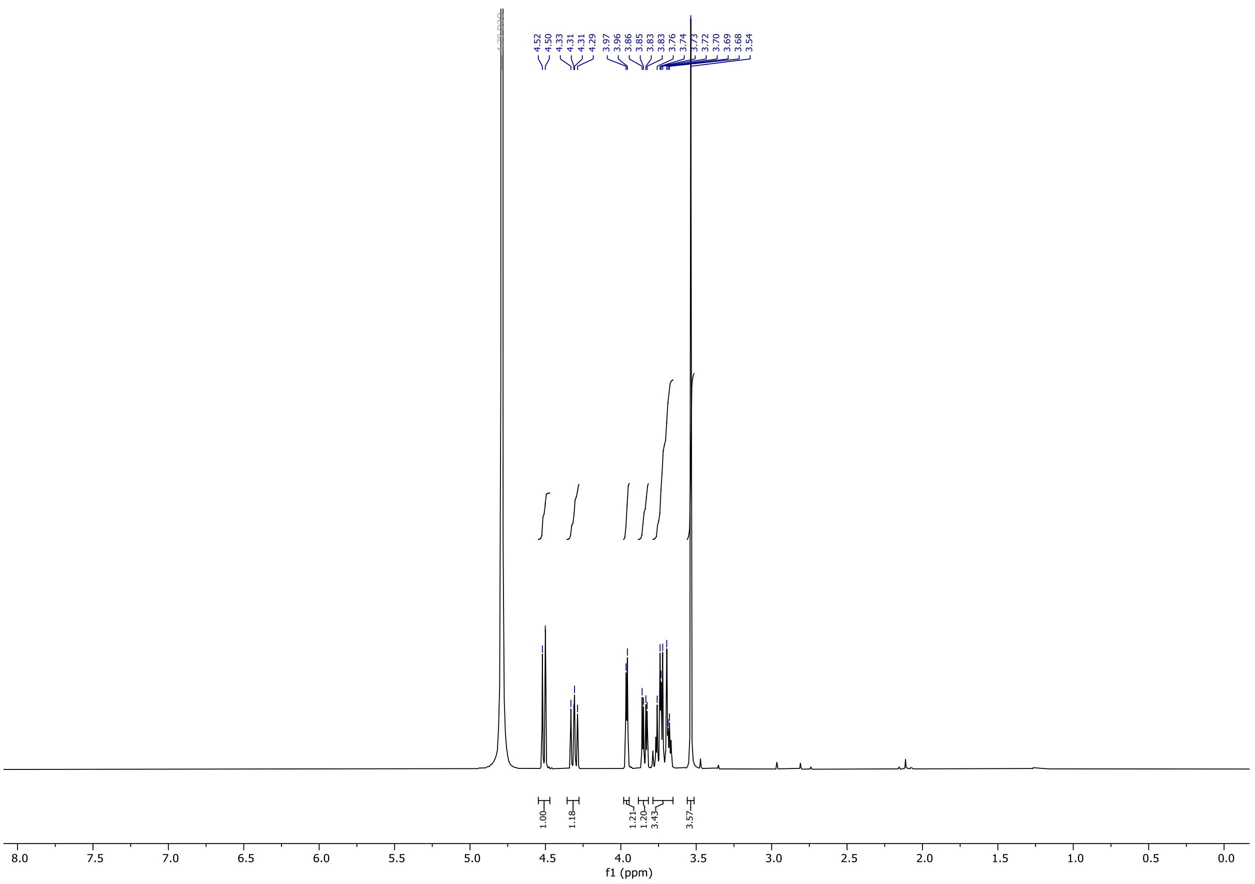

**[1]H NMR**

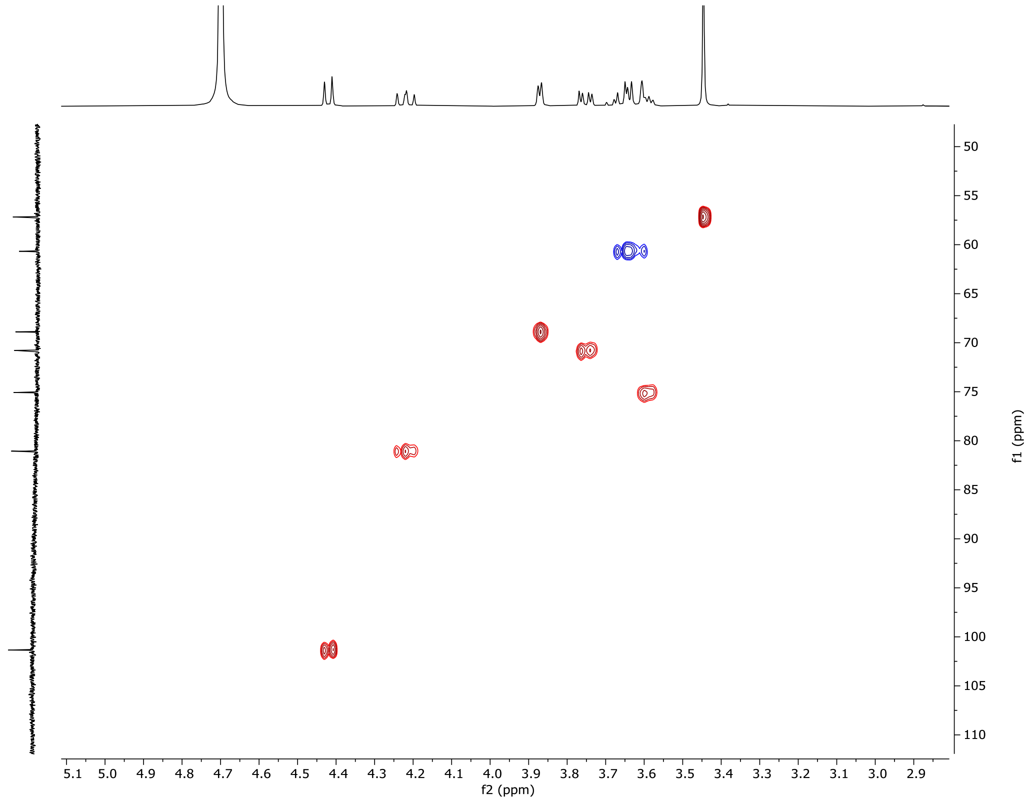

**[1]H- [13]C HSQC**

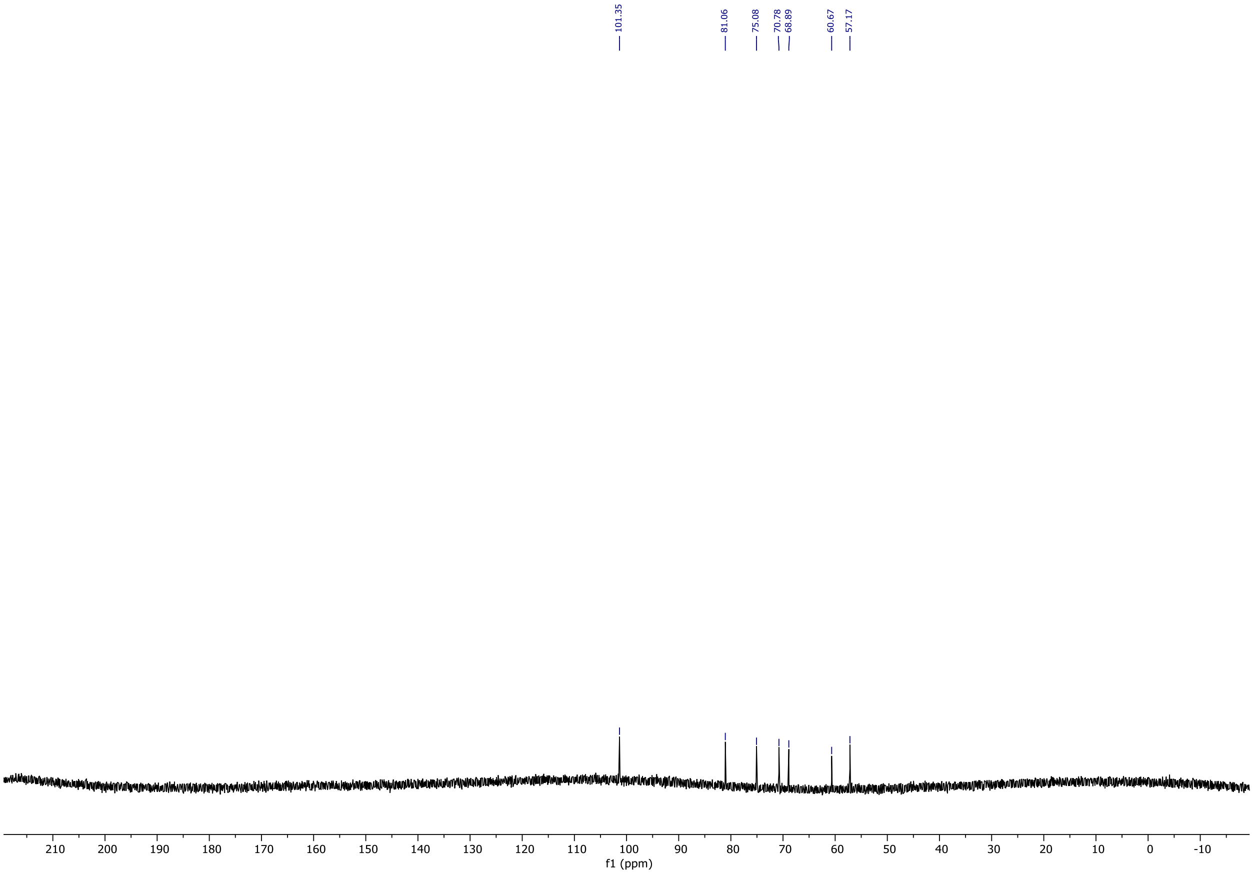
101.35
81.06
75.08
70.78
68.89
60.67
57.17
210 200 190 180 170 160 150 140 130 120 110 100 90 80 70 60 50 40 30 20 10 0 -10
f1 (ppm)

**$^{13}$C NMR**

### Methyl 3-*O*-(ethenylsulfonyl)-β-D-galactoside (25)

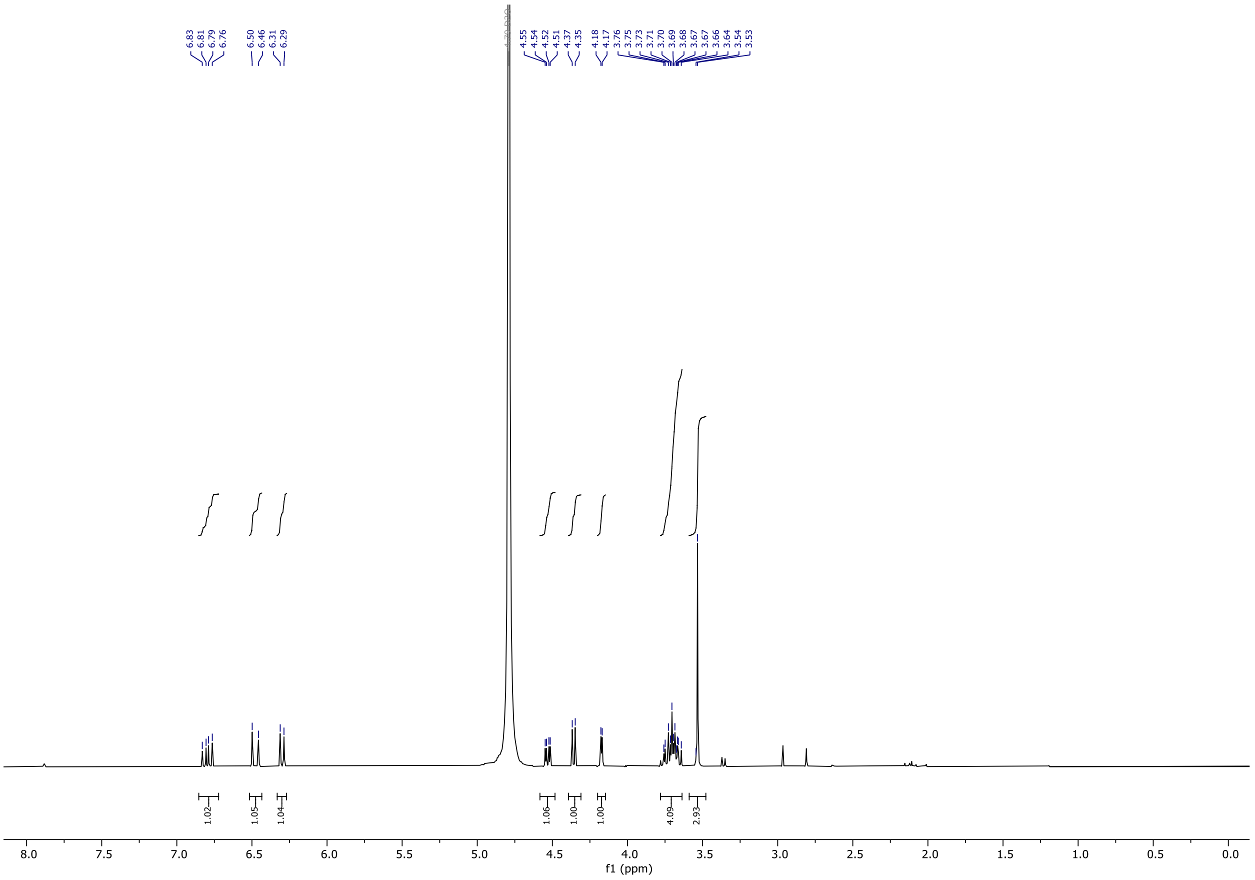

$^1$H NMR

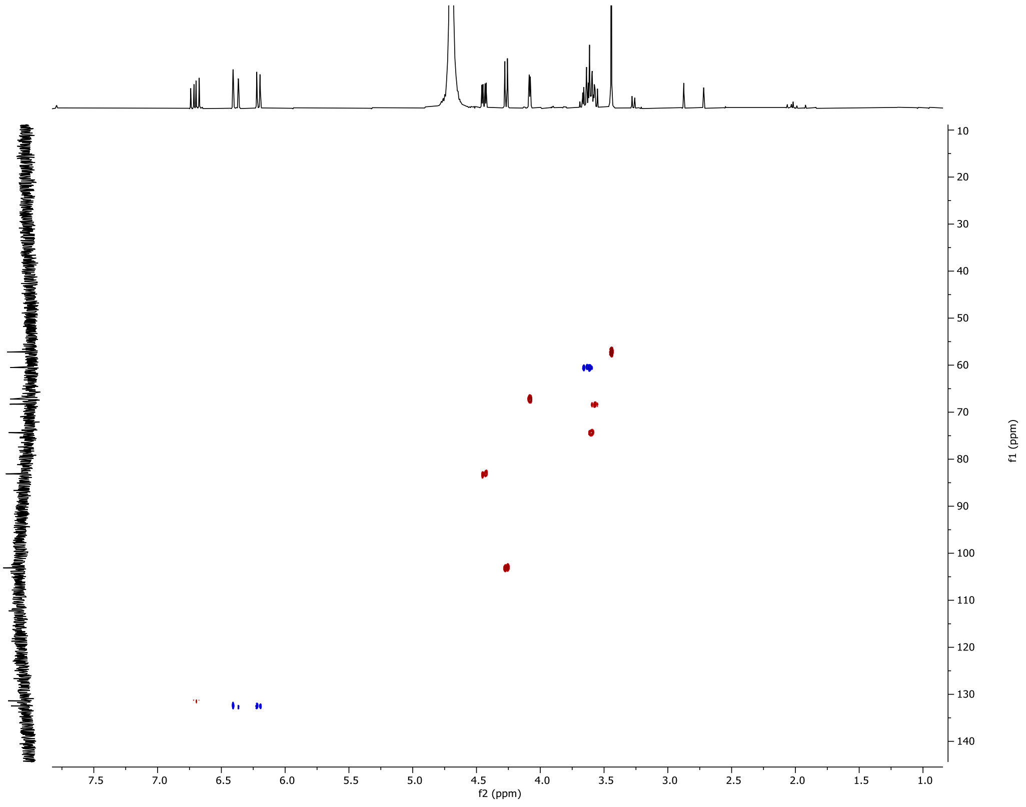

$^1$H-$^{13}$C HSQC NMR

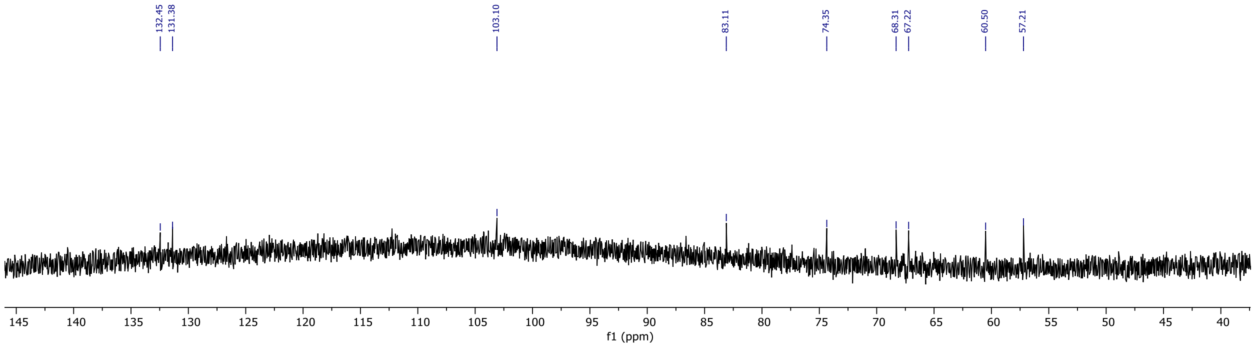
132.45
131.38
103.10
83.11
74.35
68.31
67.22
60.50
57.21
145 140 135 130 125 120 115 110 105 100 95 90 85 80 75 70 65 60 55 50 45 40
f1 (ppm)

**$^{13}$C NMR**

### Ethyl 4,6-*O*-benzylidene-1-thio-β-D-galactoside (26)

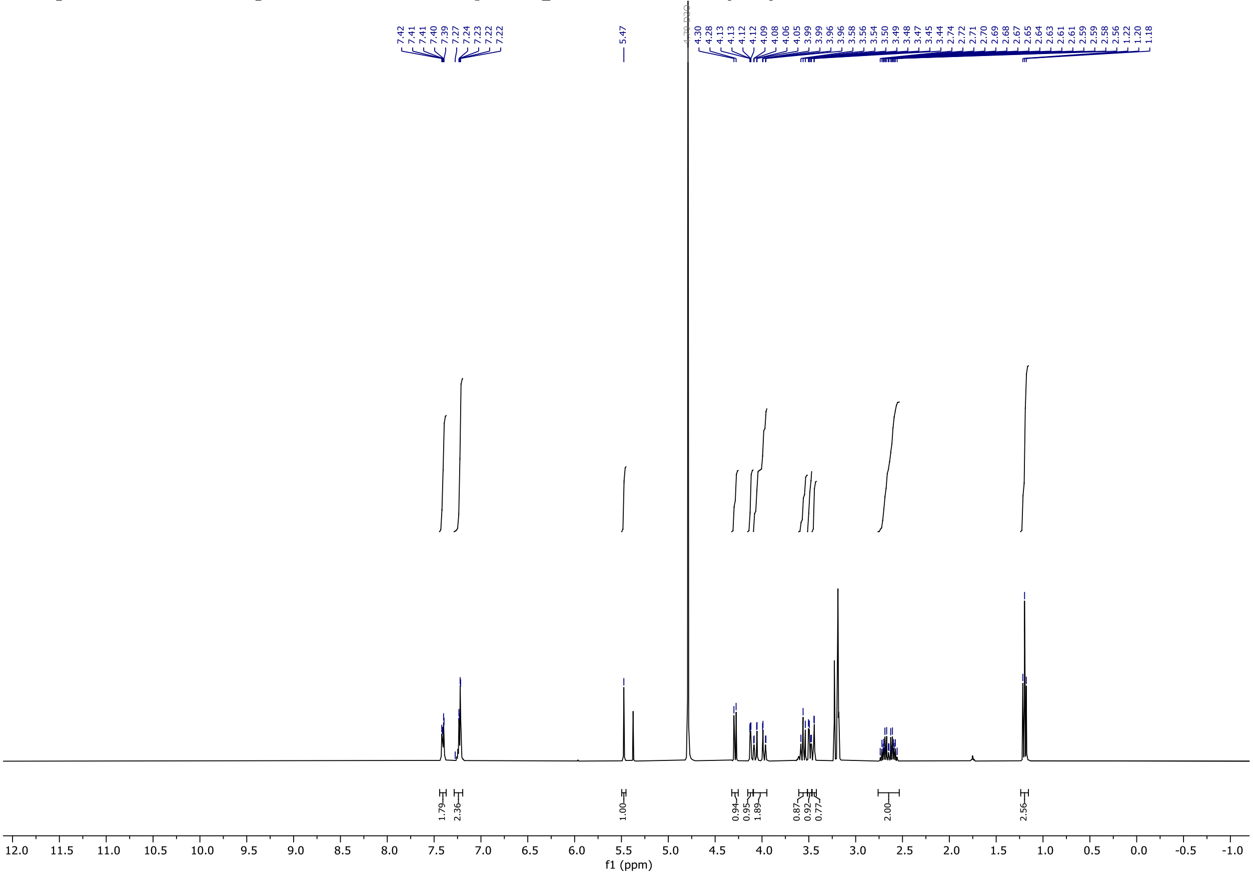

**$^{1}$H NMR**

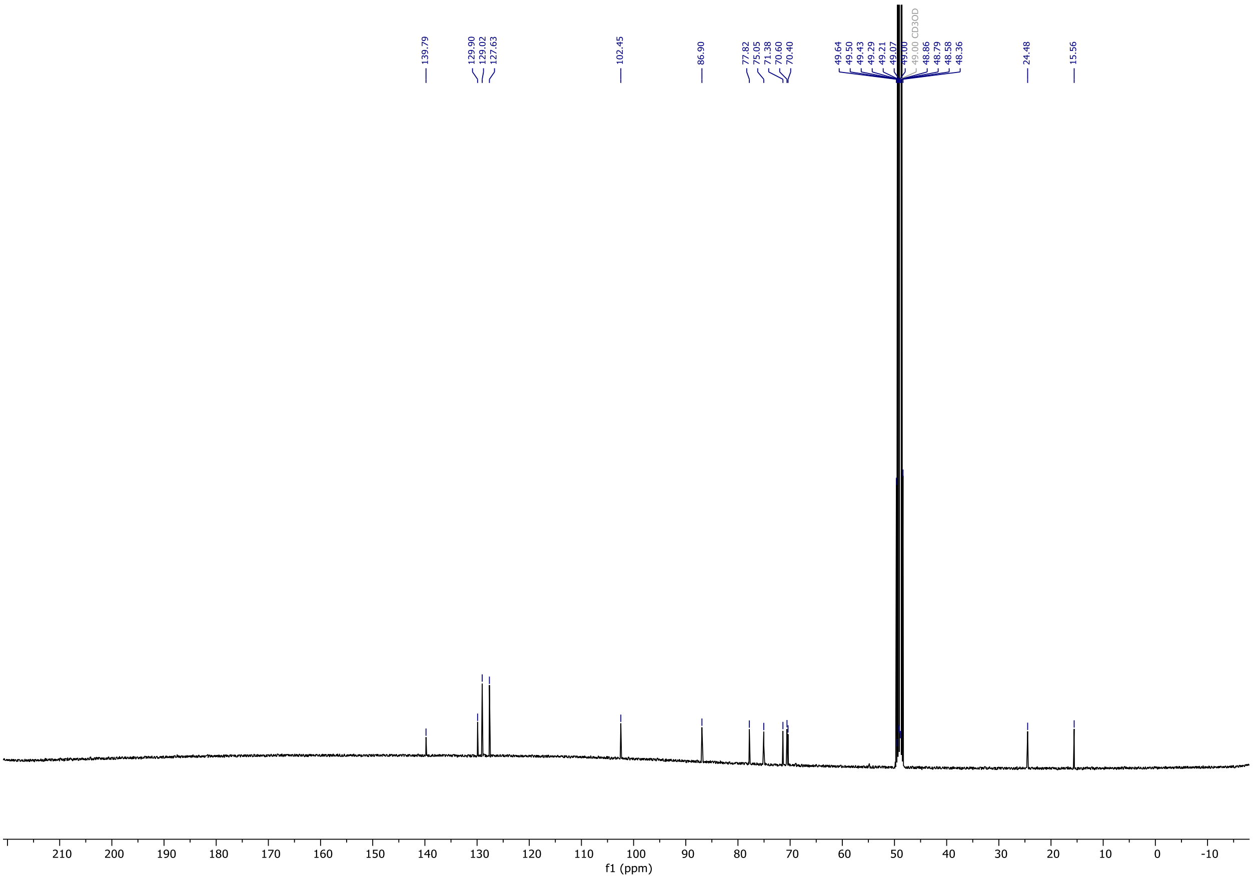

**$^{13}$C NMR**

### Ethyl 2,3-di-*O*-acetyl-4,6-*O*-benzylidene-1-thio-β-D-galactoside (27)

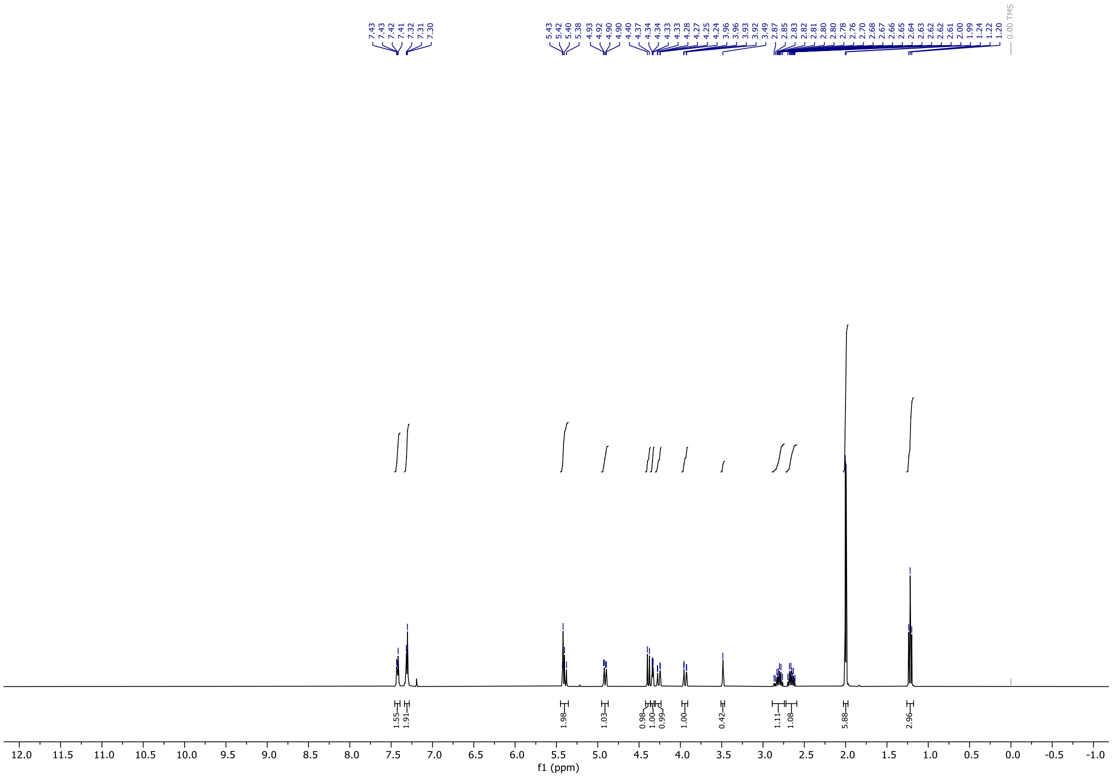


**$^{1}$H NMR**

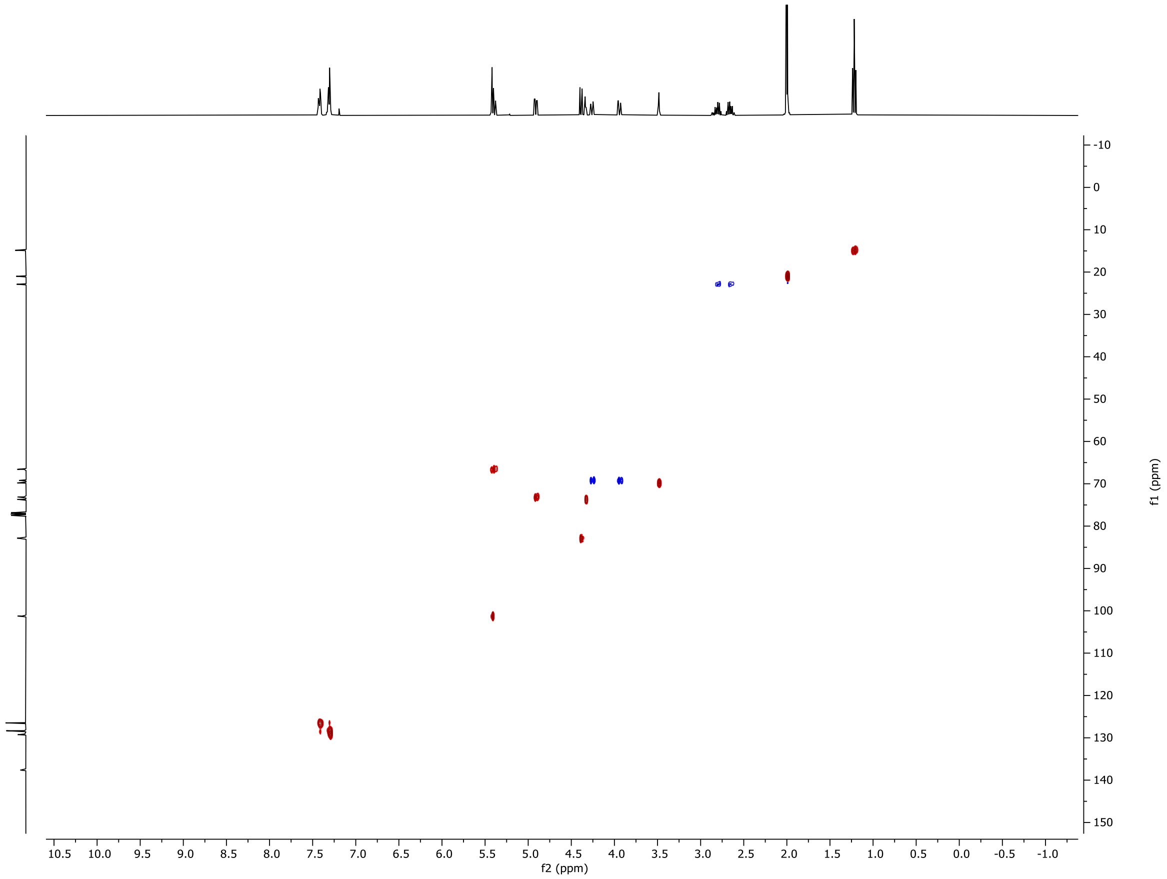


**$^{1}$H-$^{13}$C HSQC**

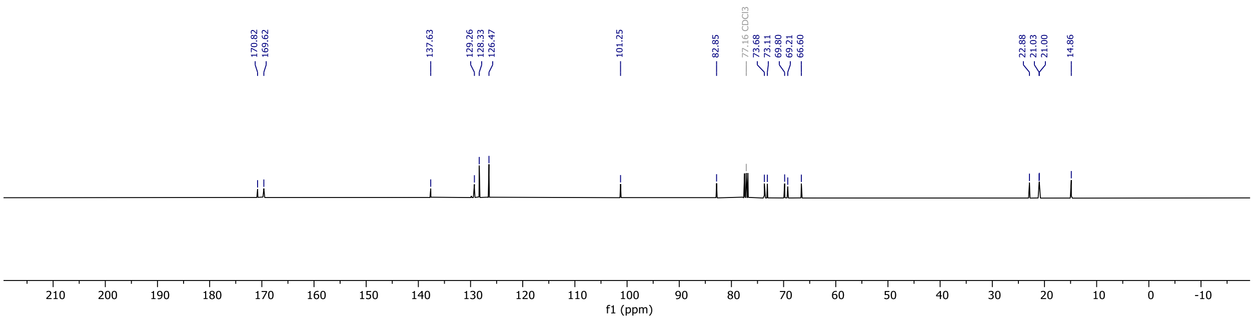

170.82
169.62
137.63
129.26
128.33
126.47
101.25
82.85
77.16 CDCl3
73.68
73.11
69.80
69.21
66.60
22.88
21.03
21.00
14.86
210
200
190
180
170
160
150
140
130
120
110
100
90
80
70
60
50
40
30
20
10
0
-10
f1 (ppm)


**$^{13}$C NMR**

### Methyl 4,6-*O*-benzylidene-β-D-galactoside (28)

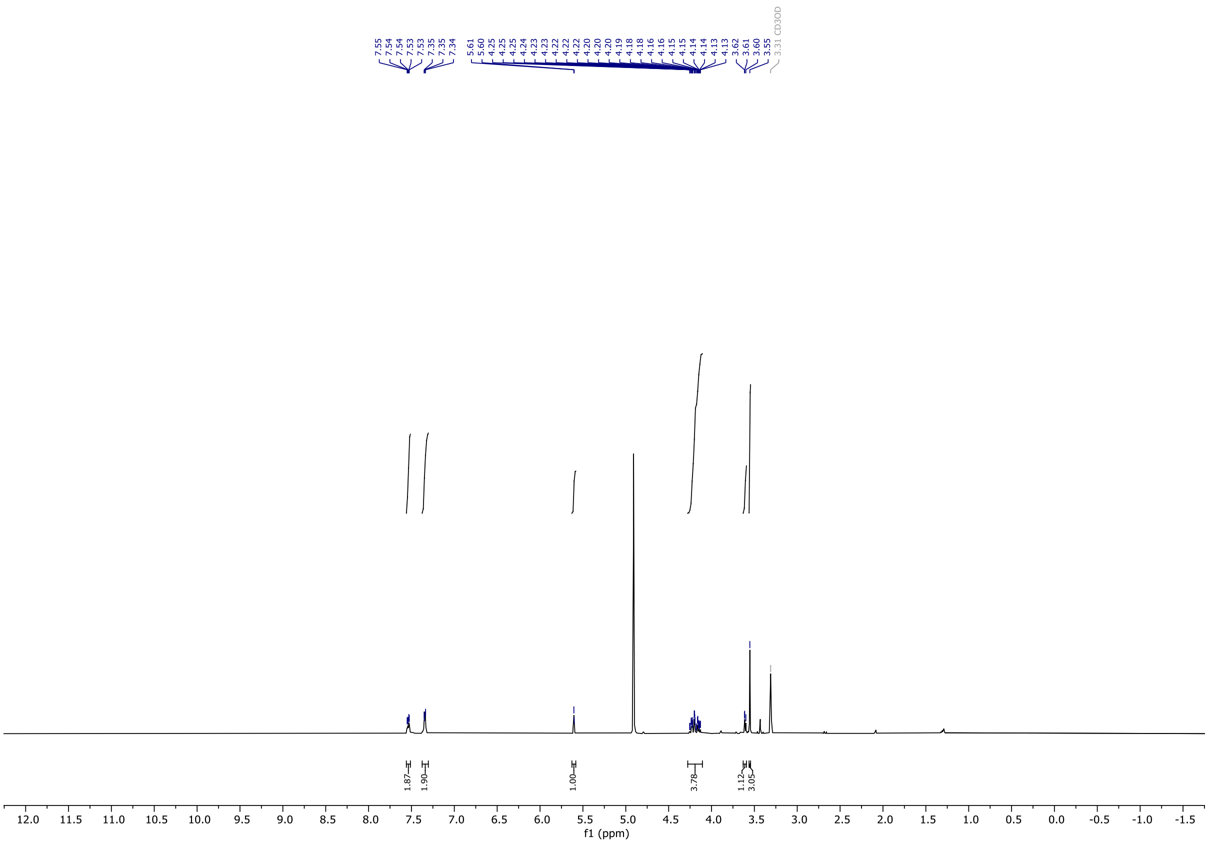

$^1$H NMR

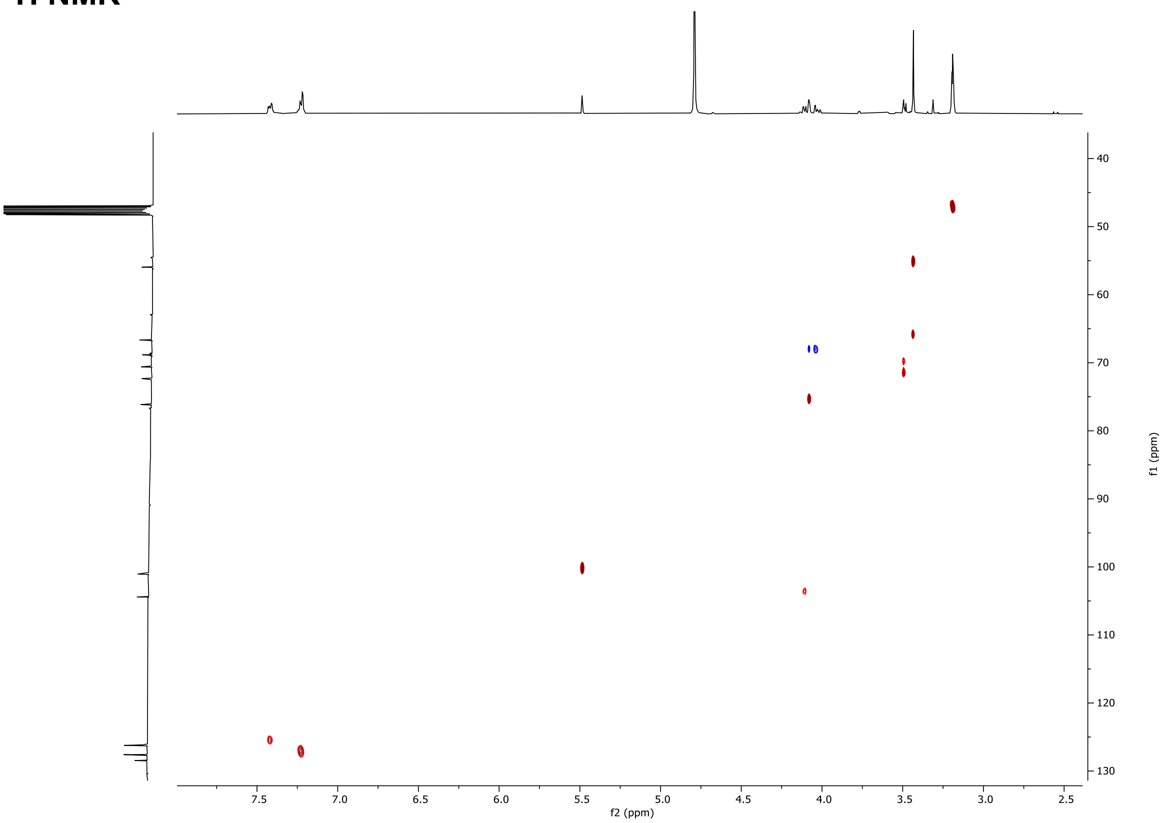

$^1$H-$^{13}$C HSQC NMR

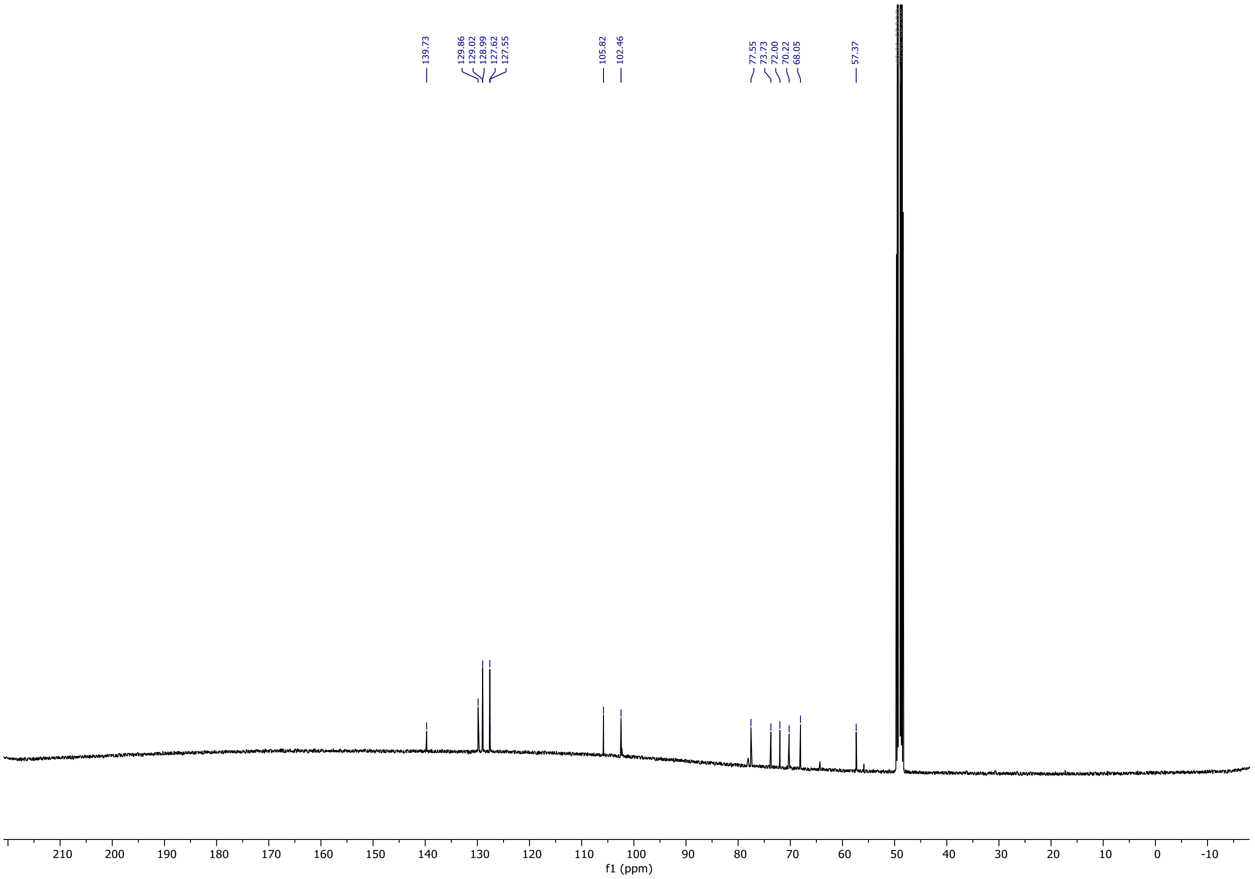
139.73
129.86
129.02
128.99
127.62
127.55
105.82
102.46
77.55
73.73
72.00
70.22
68.05
57.37
210
200
190
180
170
160
150
140
130
120
110
100
90
80
70
60
50
40
30
20
10
0
-10
f1 (ppm)

**$^{13}$C NMR**

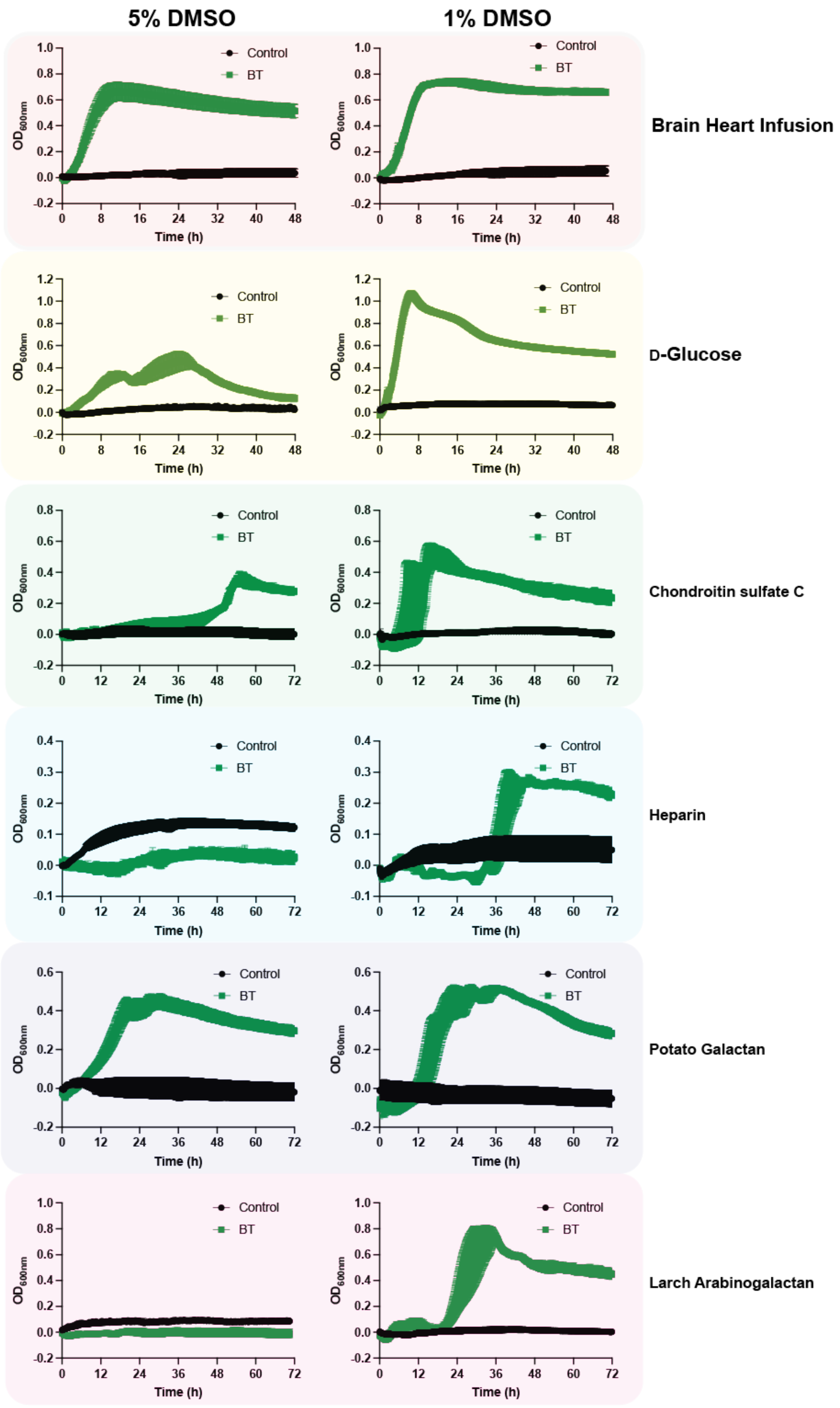


**Supplementary figure 1. The effects of DMSO on the growth of *Bacteroides thetaiotaomicron* VPI-5482**

*B. thetaiotaomicron* VPI-5482 was grown in rich media (BHI), or minimal media supplemented 5 mg/ml of an appropriate polysaccharide with either 5% or 1% DMSO. Data are technical triplicates with the standard error of the mean. Numbers indicate arylsulfamate compounds as in Figure 1

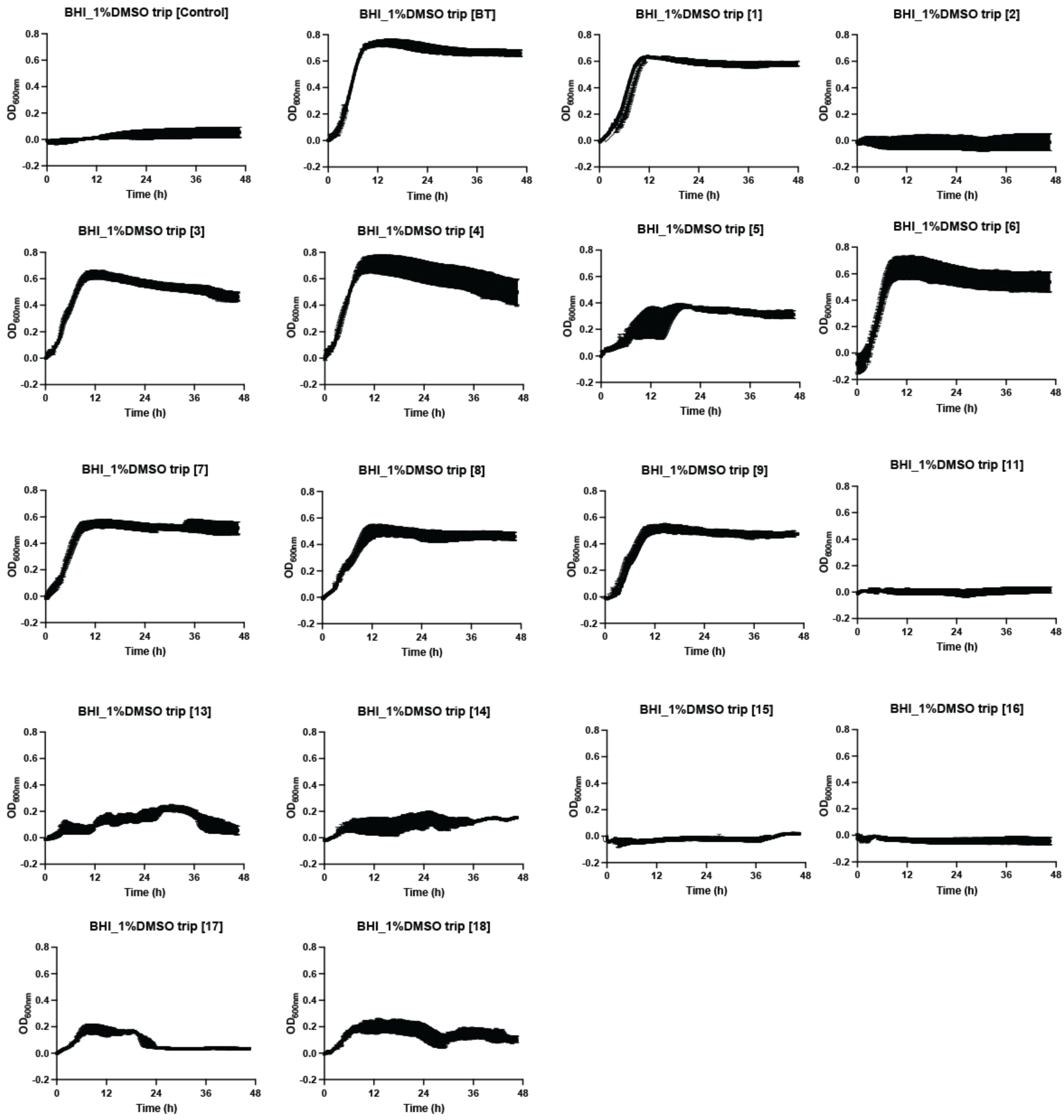


**Supplementary figure 2. The effects of arylsulfamates on the growth of *Bacteroides thetaiotaomicron* VPI-5482 grown in BHI media**

*B. thetaiotaomicron* VPI-5482 was grown in rich media (BHI), with 1% DMSO, and with or without 1 mM of the appropriate arylsulfamate. Data are technical triplicates with the standard error of the mean. Numbers indicate arylsulfamate compounds as in Figure 1.

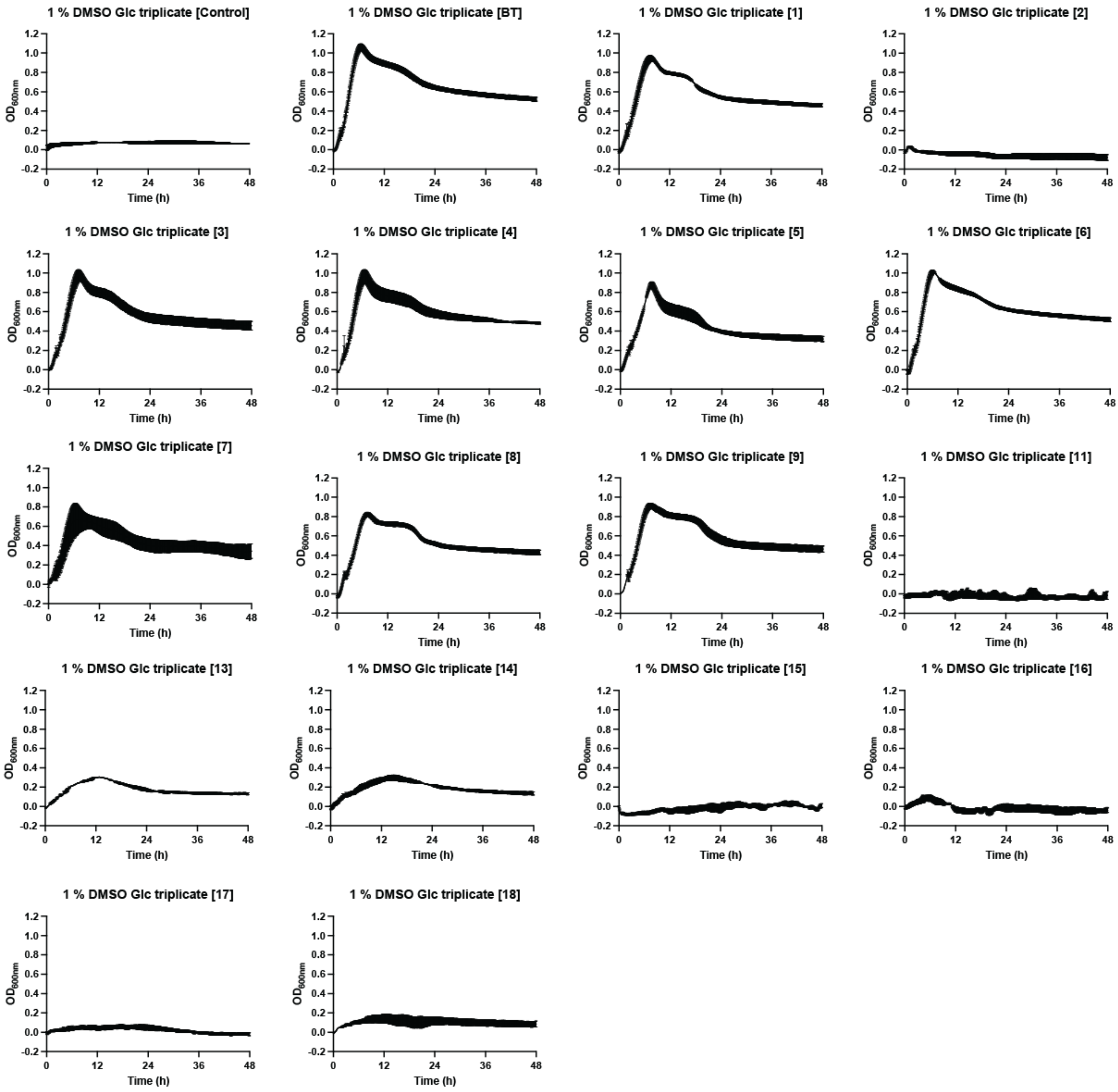


**Supplementary figure 3. The effects of arylsulfamates on the growth of *Bacteroides thetaiotaomicron* VPI-5482 grown in minimal media with D-glucose**

*B. thetaiotaomicron* VPI-5482 was grown in minimal media supplemented with 5 mg/ml D-glucose (Glc), 1% DMSO, and with or without 1 mM of the appropriate arylsulfamate. Data are technical triplicates with the standard error of the mean. Numbers indicate arylsulfamate compounds as in Figure 1.

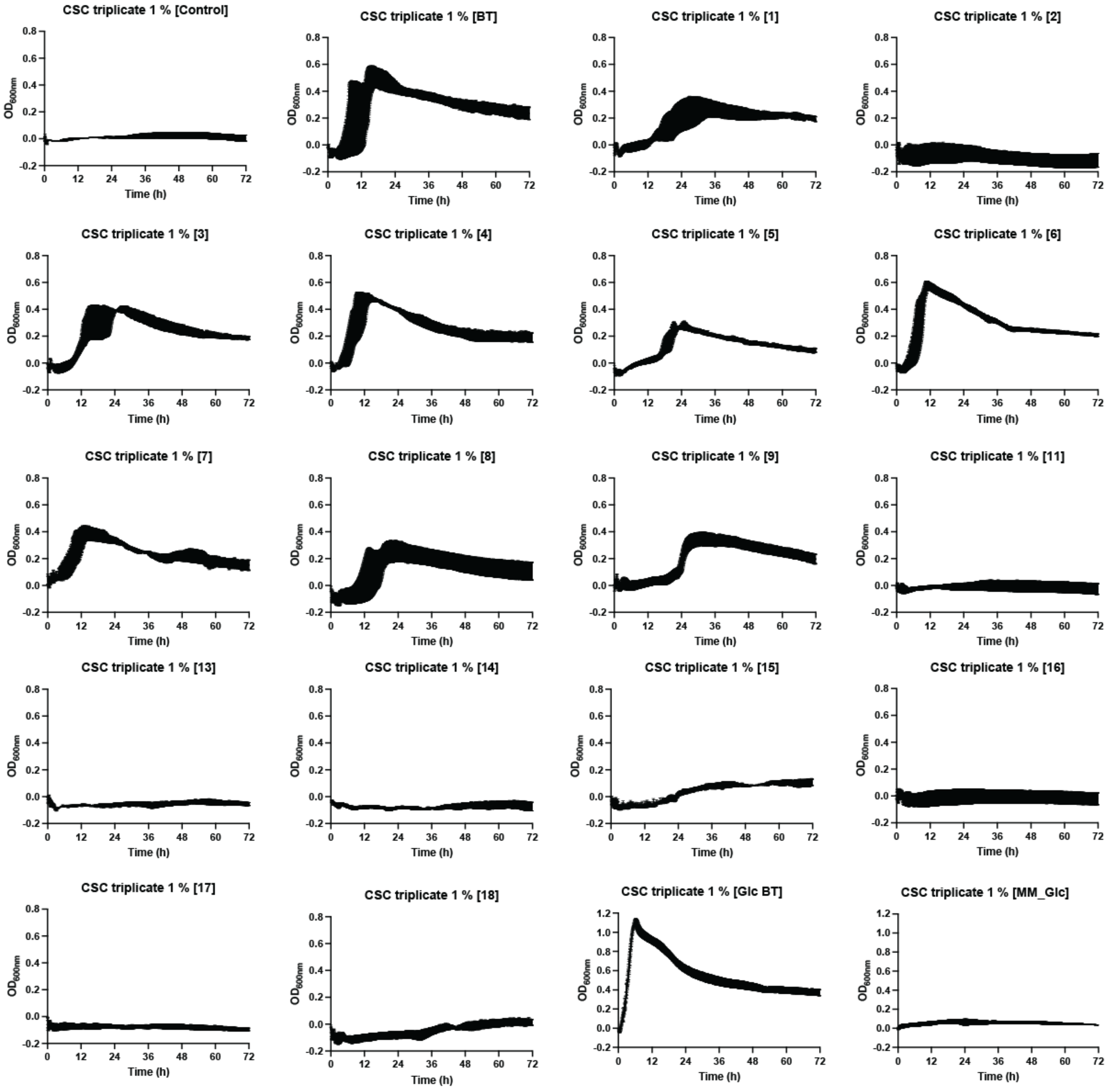


**Supplementary figure 4. The effects of arylsulfamates on the growth of *Bacteroides thetaiotaomicron* VPI-5482 grown in minimal media with Chondroitin sulfate C (CSC)**

*B. thetaiotaomicron* VPI-5482 was grown in minimal media supplemented with 5 mg/ml Chondroitin sulfate C (CSC), 1% DMSO, and with or without 1 mM of the appropriate arylsulfamate. Data are technical triplicates with the standard error of the mean. Numbers indicate arylsulfamate compounds as in Figure 1.

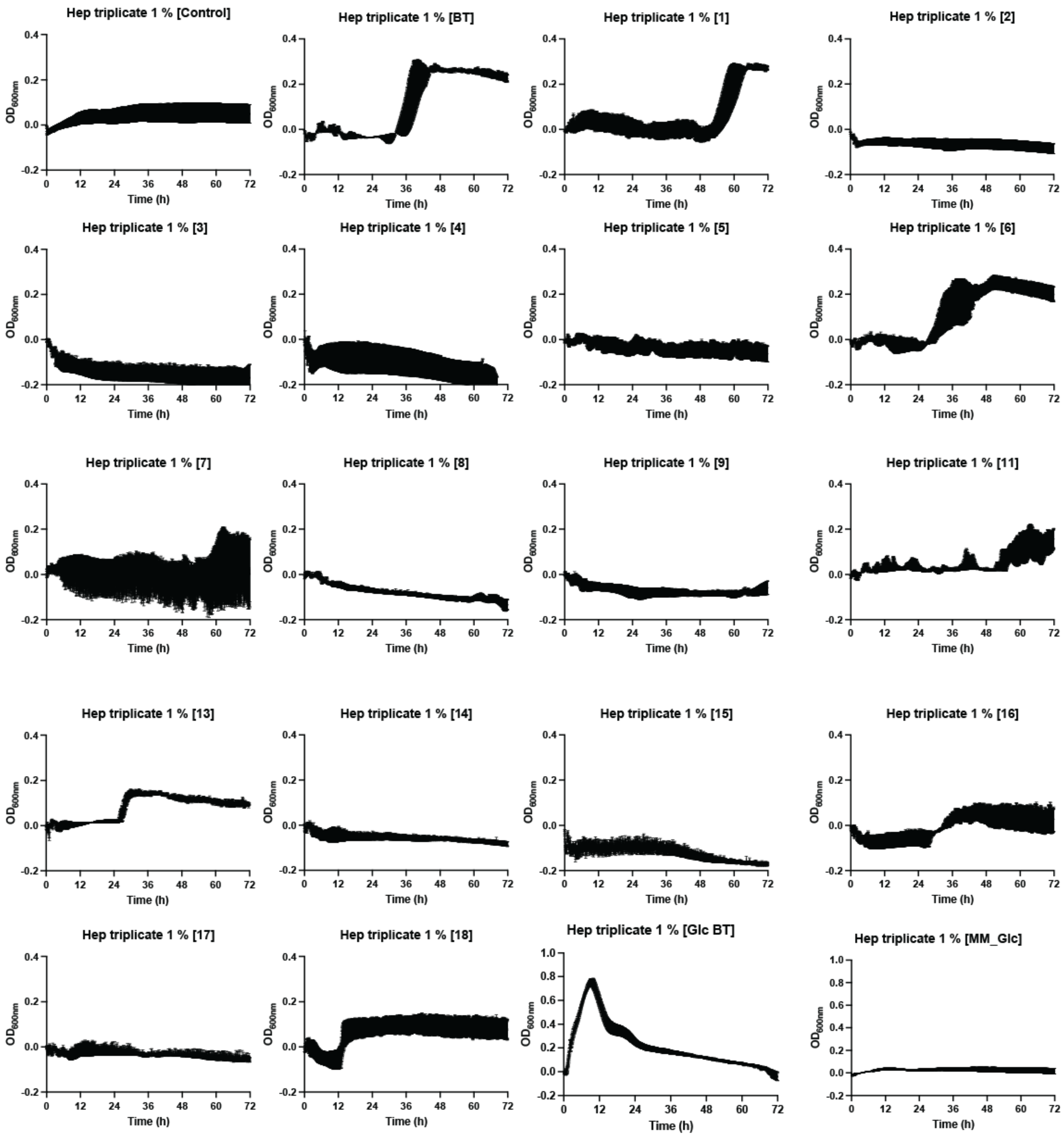


**Supplementary figure 5. The effects of arylsulfamates on the growth of *Bacteroides thetaiotaomicron* VPI-5482 grown in minimal media with Heparin**

*B. thetaiotaomicron* VPI-5482 was grown in minimal media supplemented with 5 mg/ml Heparin (Hep), 1% DMSO, and with or without 1 mM of the appropriate arylsulfamate. Data are technical triplicates with the standard error of the mean. Numbers indicate arylsulfamate compounds as in Figure 1.

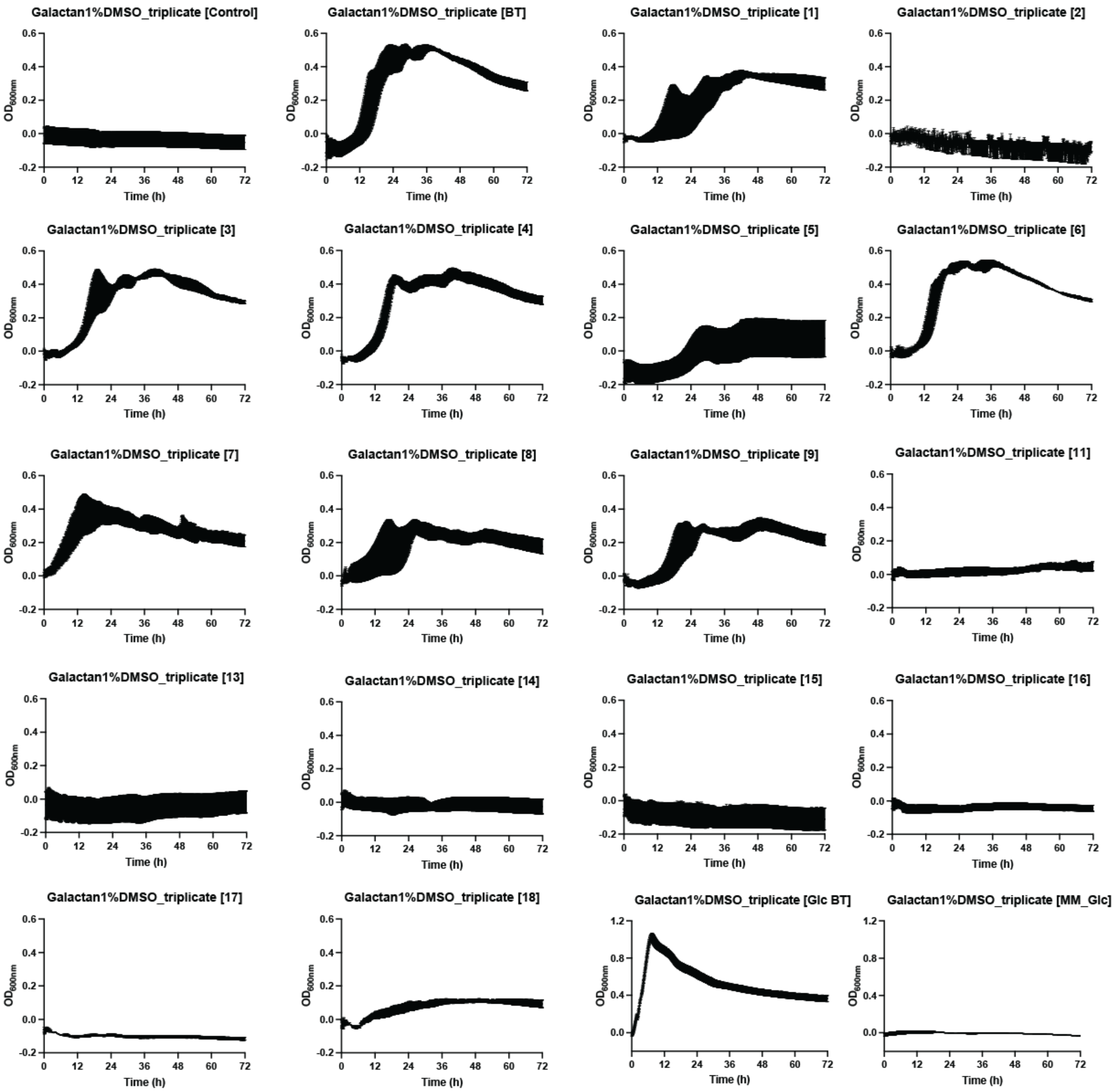


**Supplementary figure 6. The effects of arylsulfamates on the growth of *Bacteroides thetaiotaomicron* VPI-5482 grown in minimal media with Potato galactan**

*B. thetaiotaomicron* VPI-5482 was grown in minimal media supplemented with 5 mg/ml Potato galactan (PG), 1% DMSO, and with or without 1 mM of the appropriate arylsulfamate. Data are technical triplicates with the standard error of the mean. Numbers indicate arylsulfamate compounds as in Figure 1.

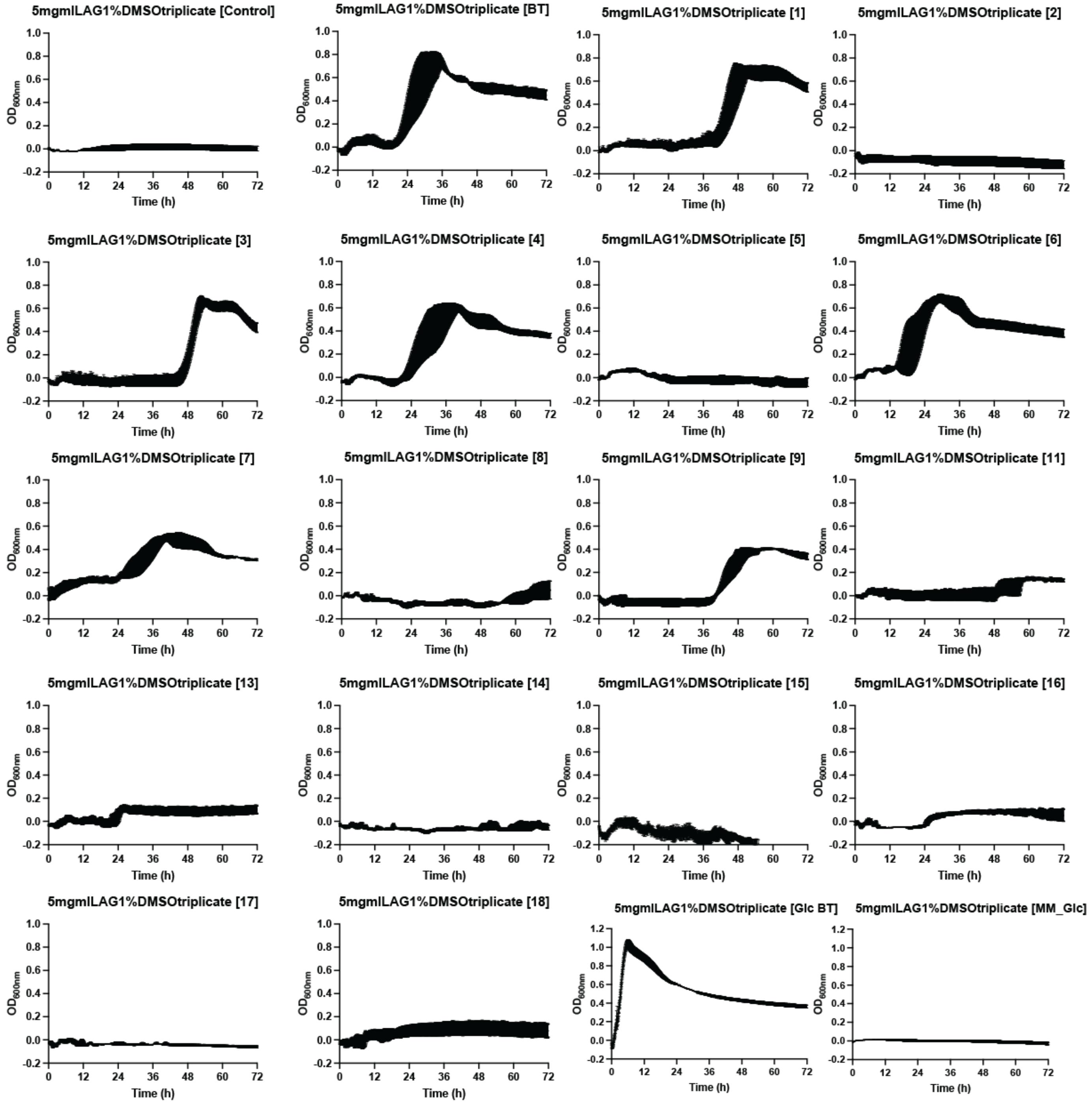


**Supplementary figure 7. The effects of arylsulfamates on the growth of *Bacteroides thetaiotaomicron* VPI-5482 grown in minimal media with Larch arabinogalactan**

*B. thetaiotaomicron* VPI-5482 was grown in minimal media supplemented with 5 mg/ml Larch arabinogalactan (LAG), 1% DMSO, and with or without 1 mM of the appropriate arylsulfamate. Data are technical triplicates with the standard error of the mean. Numbers indicate arylsulfamate compounds as in Figure 1.

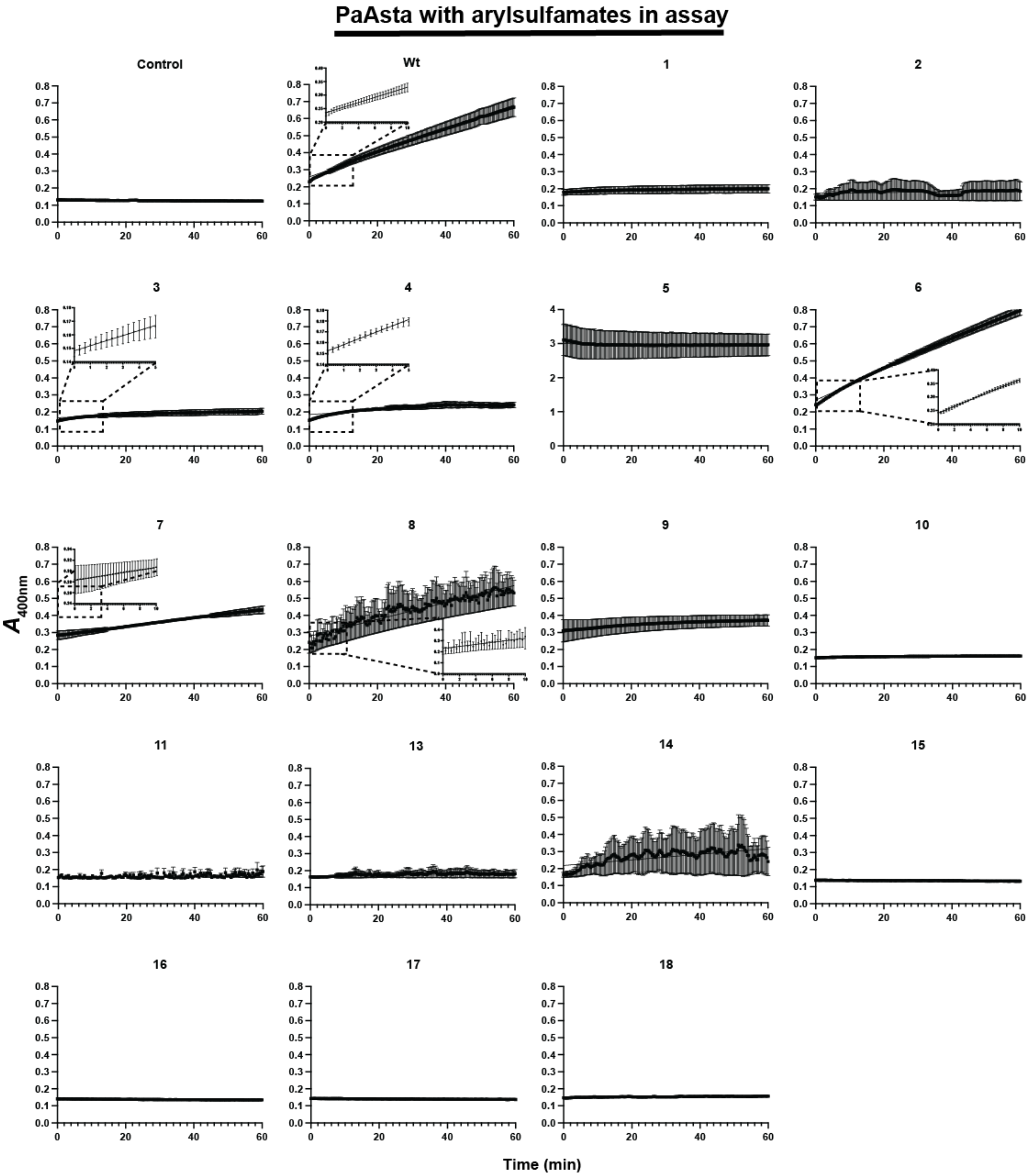


**Supplementary figure 8. Kinetic curves of *Pa*Asta assayed against *para*-nitrophenol sulfate with and without arylsulfamate inhibitors**

*Pa*Asta, at a concentration of 25 nM, was assayed against 1 mM *para*-nitrophenol with and without 1 mM of various arylsulfamate inhibitors included in the assay. The assay was performed in 100 mM of Bis-Tris-propane pH 7.0 with 5% DMSO, 150 mM NaCl, 0.02% (v/v) Brij-35 and 5 mM $CaCl_2$. Assays were performed in triplicate. Numbers indicate arylsulfamate compounds as in Figure 1.

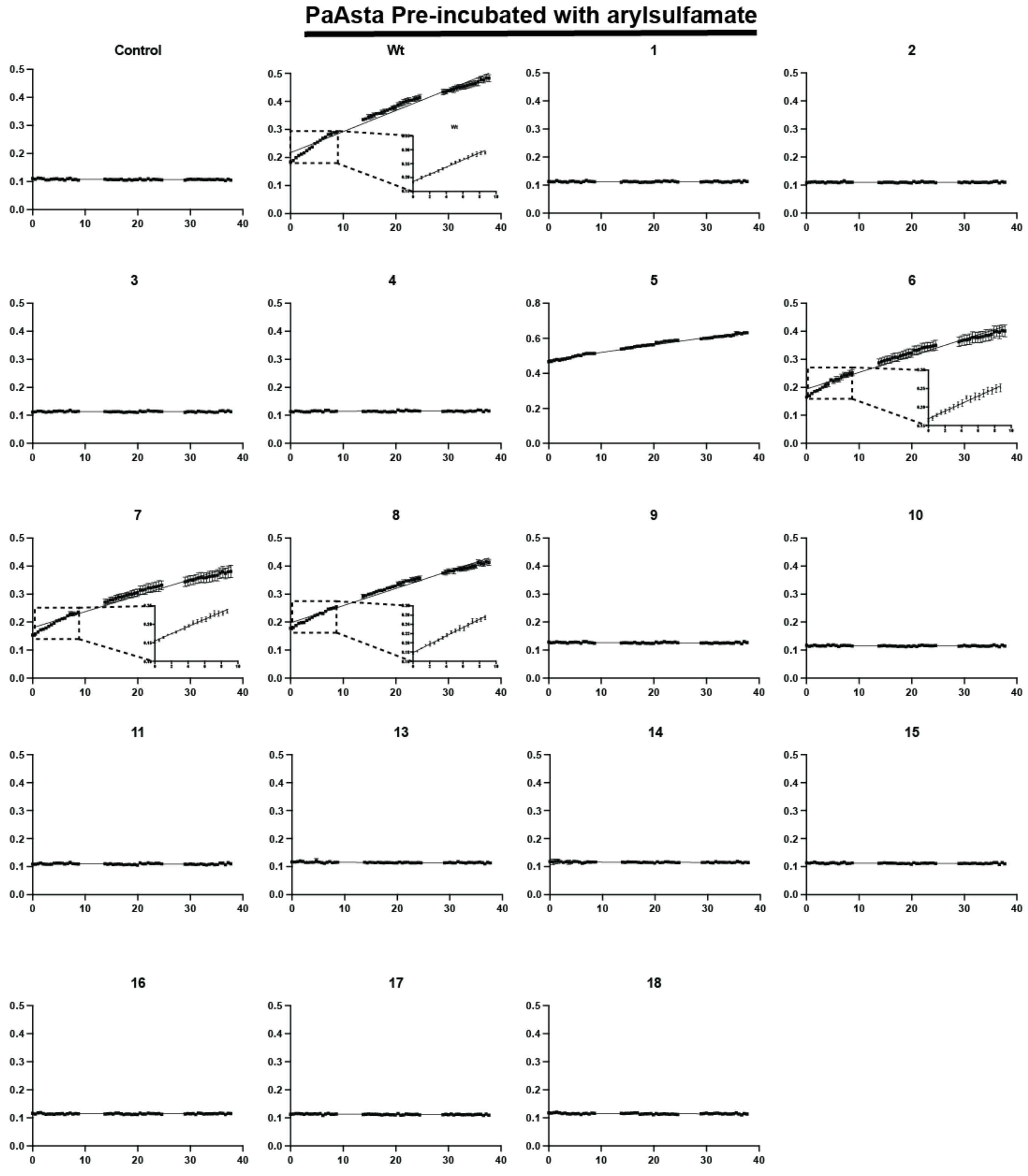


**Supplementary figure 9. Kinetic curves of *Pa*Asta assayed against *para*-nitrophenol sulfate after pre-incubation of the enzyme with arylsulfamate inhibitors**

*Pa*Asta, which had been incubated for ~24h with 1 mM of the appropriate arylsulfamate inhibitor, was assayed using a concentration of 25 nM against 1 mM para-nitrophenol. The assay was performed in 100 mM of Bis-Tris-propane pH 7.0 with 5% DMSO, 150 mM NaCl, 0.02% (v/v) Brij-35 and 5 mM $CaCl_2$. Assays were performed in triplicate. Numbers indicate arylsulfamate compounds as in Figure 1.

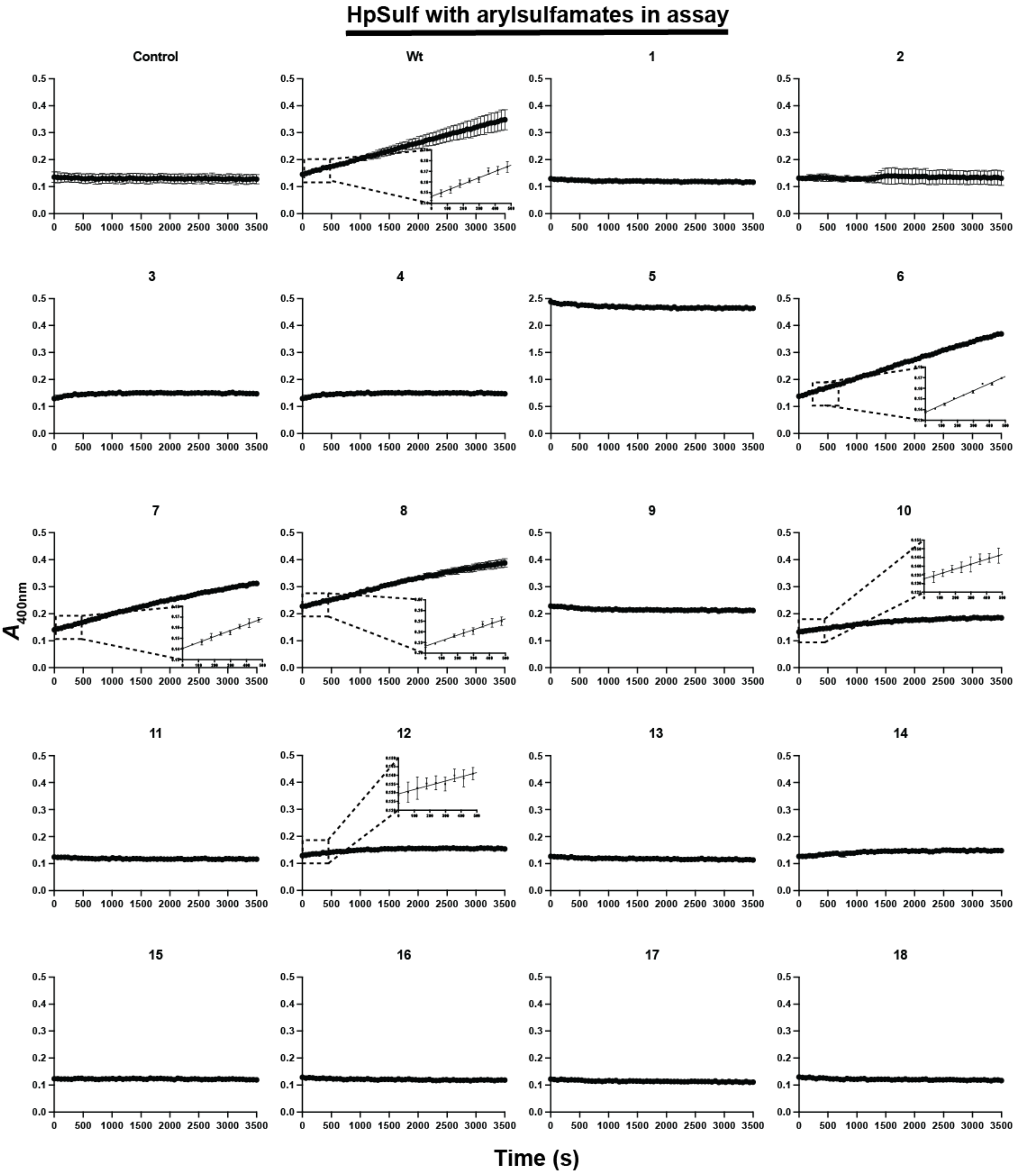


**Supplementary figure 10. Kinetic curves of *Hp*Sulf assayed against para-nitrophenol sulfate with and without arylsulfamate inhibitors**

*Hp*Sulf, at a concentration of 5 μg/ml, was assayed against 1 mM para-nitrophenol with and without 1 mM of various arylsulfamate inhibitors included in the assay. The assay was performed in 100 mM of Bis-Tris-propane pH 7.0 with 5% DMSO, 150 mM NaCl, 0.02% (v/v) Brij-35 and 5 mM $CaCl_2$. Assays were performed in triplicate. Numbers indicate arylsulfamate compounds as in Figure 1.

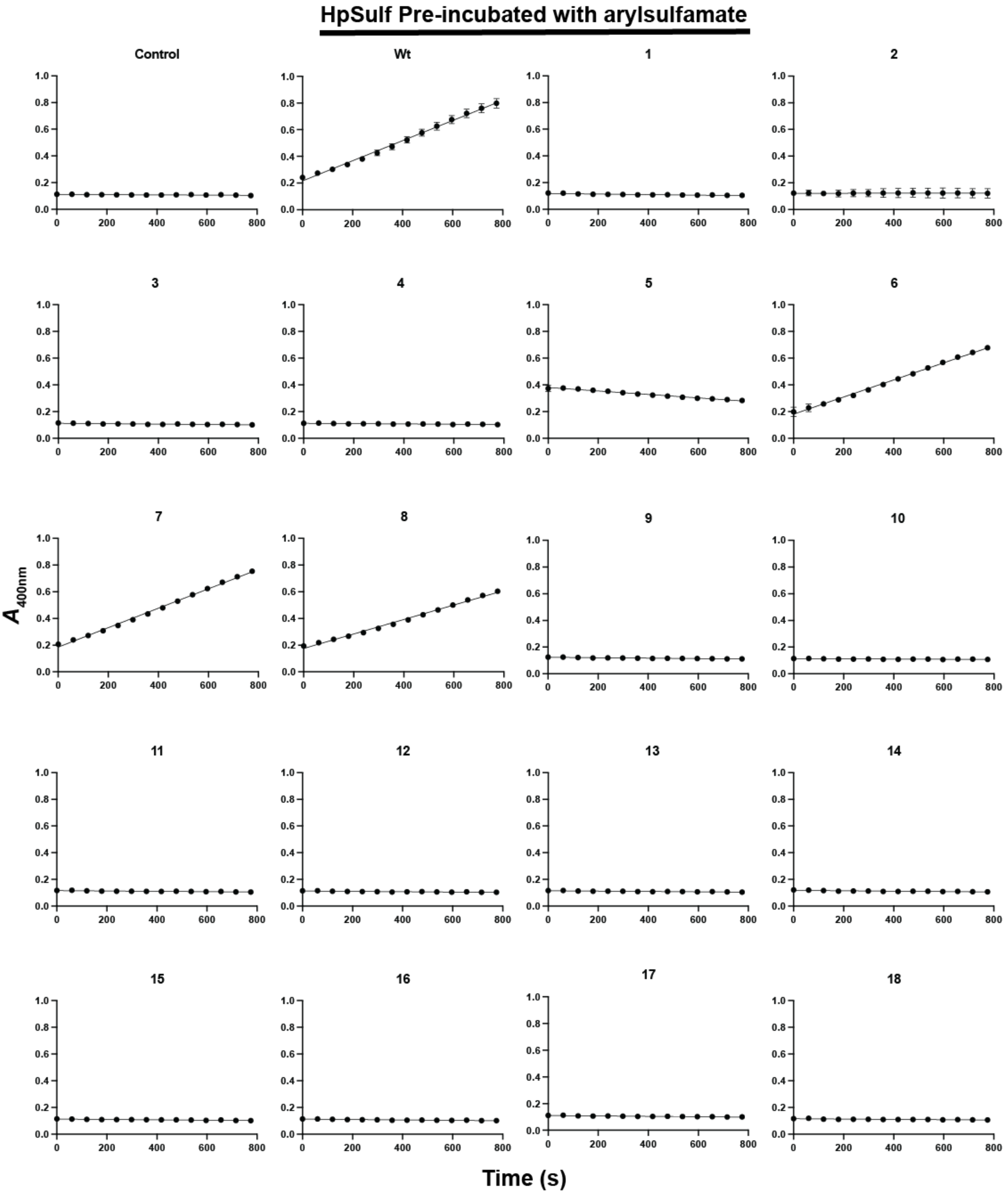


**Supplementary figure 11. Kinetic curves of *Hp*Sulf assayed against para-nitrophenol sulfate after pre-incubation of the enzyme with arylsulfamate inhibitors**

*Hp*Sulf, which had been incubated for ~24h with 1 mM of the appropriate arylsulfamate inhibitor, was assayed using a concentration of 50 μg/ml against 1 mM para-nitrophenol. The assay was performed in 100 mM of Bis-Tris-propane pH 7.0 with 5% DMSO, 150 mM NaCl, 0.02% (v/v) Brij-35 and 5 mM $CaCl_2$. Assays were performed in triplicate. Numbers indicate arylsulfamate compounds as in Figure 1.

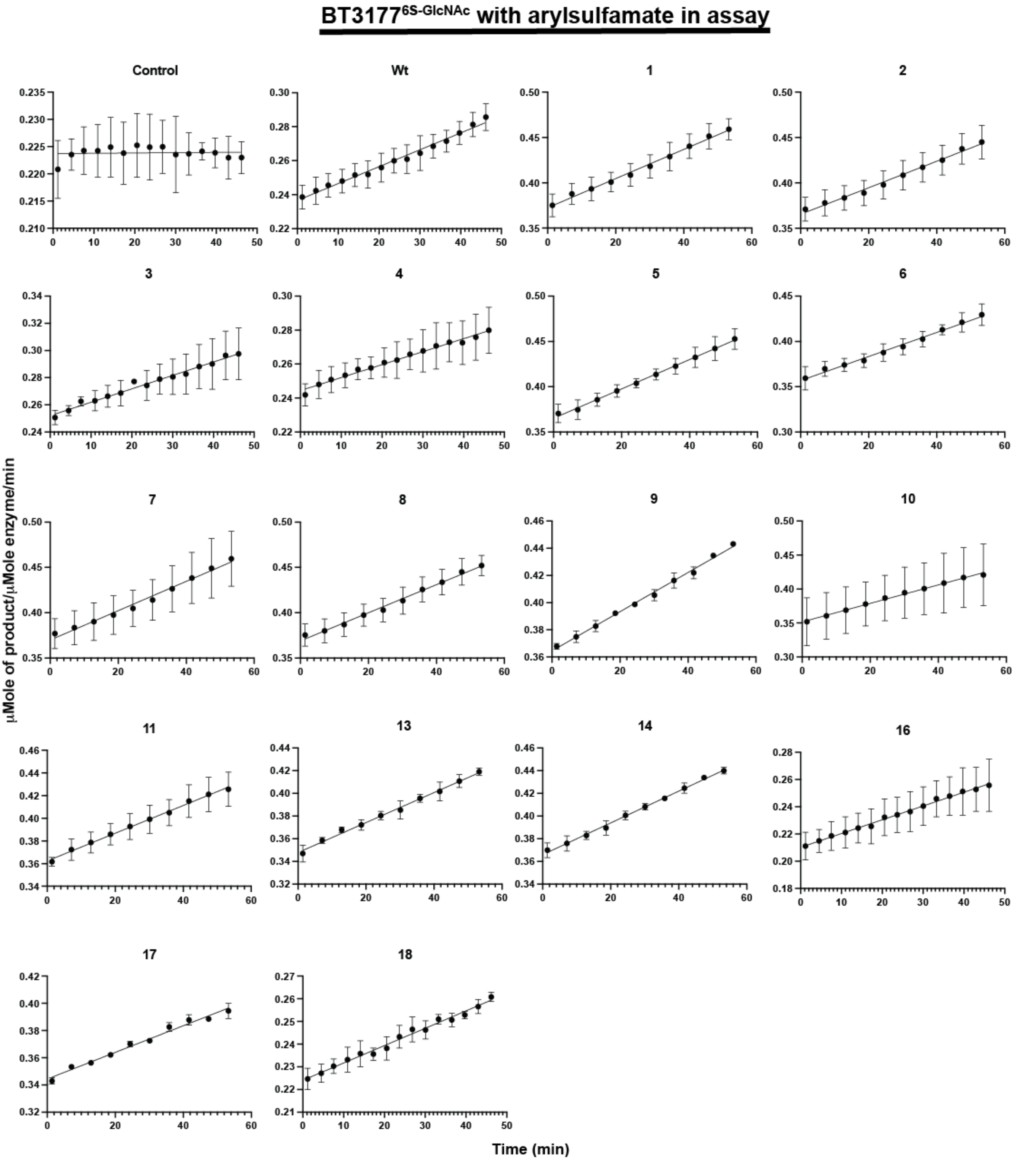


**Supplementary figure 12. Kinetic curves of BT3177$^{6S\text{-}GlcNAc}$ assayed against BODIPY labelled 6S-*N*-acetylglucosamine with and without arylsulfamate inhibitors**

BT3177$^{6S\text{-}GlcNAc}$, at a concentration of 446.5 nM, was assayed against 1 µM BODIPY labelled 6S-*N*-acetylglucosamine with and without 1 mM of various arylsulfamate inhibitors included in the assay. The assay was performed in 100 mM of Bis-Tris-propane pH 8.0 with 5% DMSO, 150 mM NaCl, 0.02% (v/v) Brij-35 and 5 mM $CaCl_2$. Assays were performed in triplicate. Numbers indicate arylsulfamate compounds as in Figure 1.

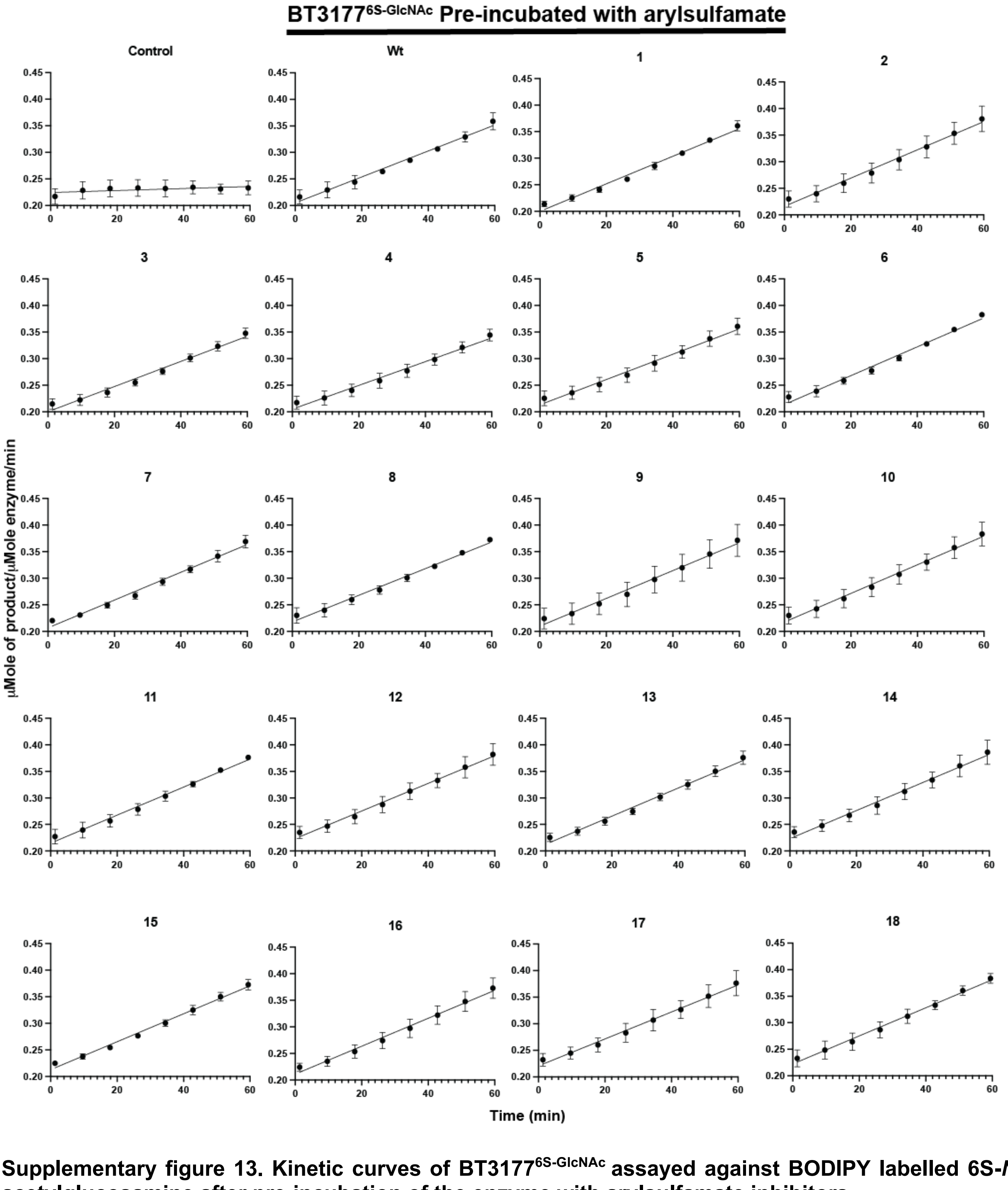


**Supplementary figure 13. Kinetic curves of BT3177$^{6S\text{-}GlcNAc}$ assayed against BODIPY labelled 6S-*N*-acetylglucosamine after pre-incubation of the enzyme with arylsulfamate inhibitors**
BT3177$^{6S\text{-}GlcNAc}$, which had been incubated for ~24h with 1 mM of the appropriate arylsulfamate inhibitor, was assayed using a concentration of 470 nM against 1 μM BODIPY labelled 6S-*N*-acetylglucosamine. The assay was performed in 100 mM of Bis-Tris-propane pH 8.0 with 5% DMSO, 150 mM NaCl, 0.02% (v/v) Brij-35 and 5 mM $CaCl_2$. Assays were performed in triplicate. Numbers indicate arylsulfamate compounds as in Figure 1.

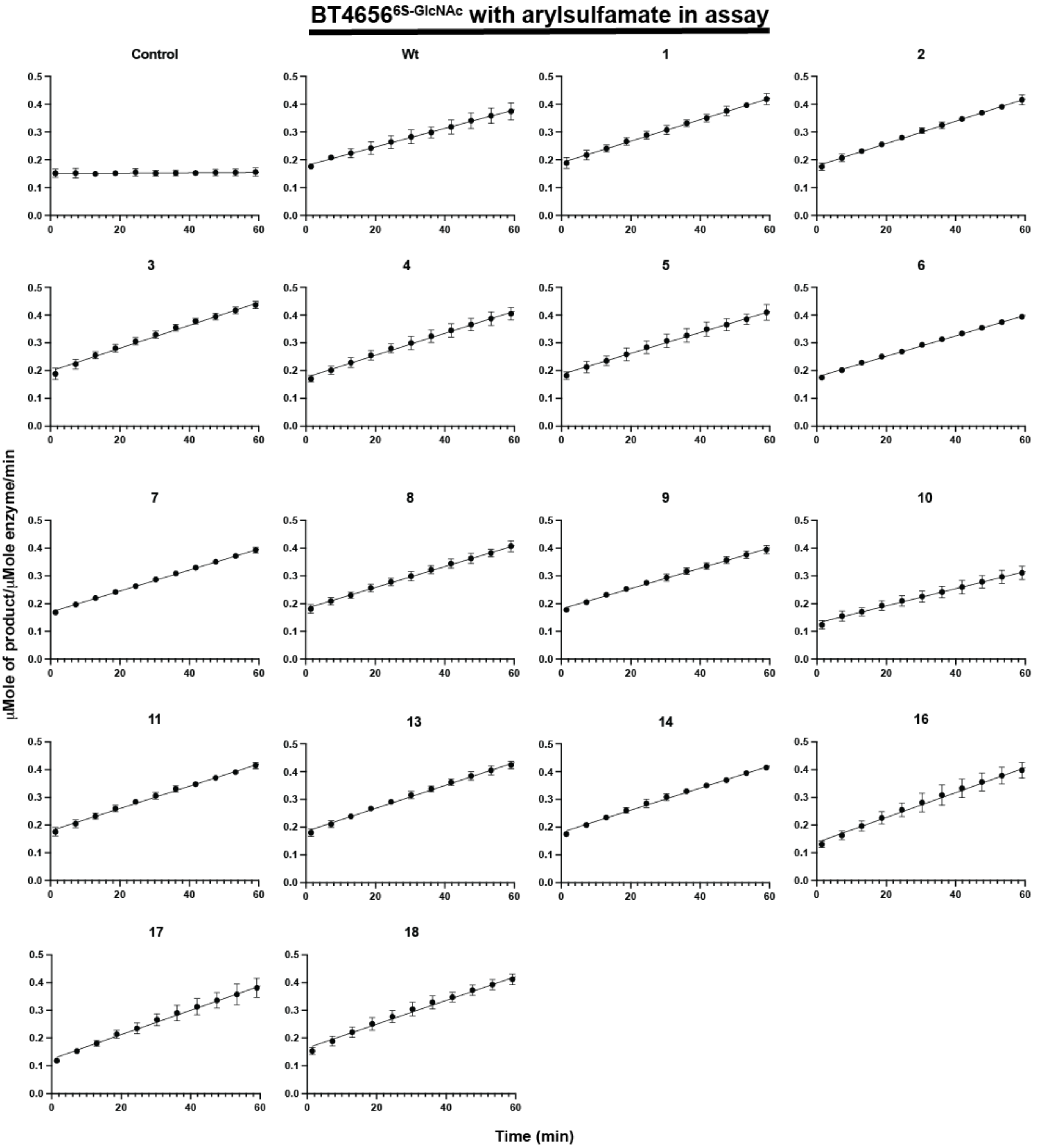


**Supplementary figure 14. Kinetic curves of BT4656$^{6S\text{-}GlcNAc}$ assayed against BODIPY labelled 6S-*N*-acetylglucosamine with and without arylsulfamate inhibitors**

BT4656$^{6S\text{-}GlcNAc}$, at a concentration of 200 nM, was assayed against 1 µM BODIPY labelled 6S-*N*-acetylglucosamine with and without 1 mM of various arylsulfamate inhibitors included in the assay. The assay was performed in 100 mM of MES pH 6.0 with 5% DMSO, 150 mM NaCl, 0.02% (v/v) Brij-35 and 5 mM $CaCl_2$. Assays were performed in triplicate. Numbers indicate arylsulfamate compounds as in Figure 1.

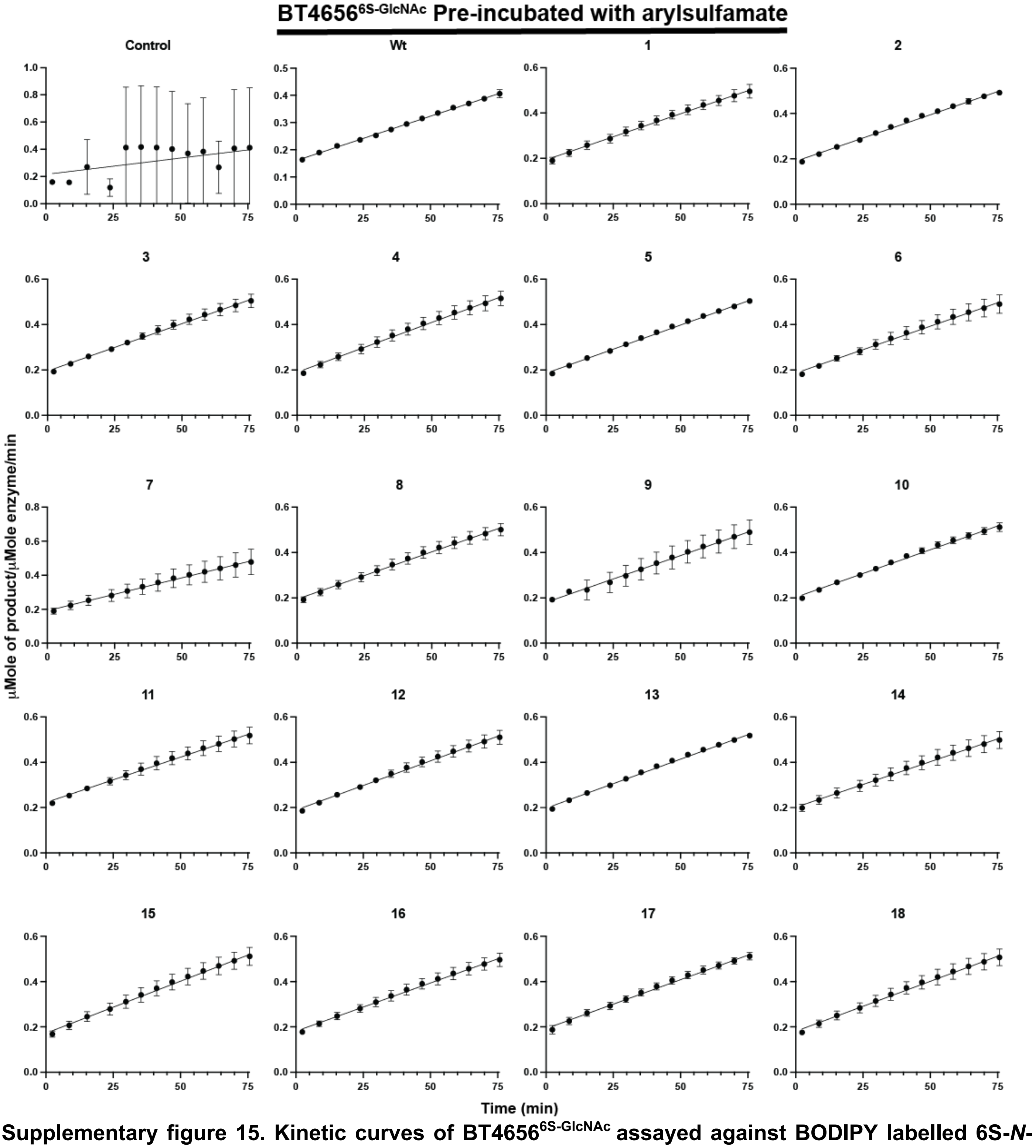


**Supplementary figure 15. Kinetic curves of BT4656$^{6S-GlcNAc}$ assayed against BODIPY labelled 6S-*N*-acetylglucosamine after pre-incubation of the enzyme with arylsulfamate inhibitors**

BT4656$^{6S-GlcNAc}$, which had been incubated for ~24h with 1 mM of the appropriate arylsulfamate inhibitor, was assayed using a concentration of 200 nM against 1 μM BODIPY labelled 6S-*N*-acetylglucosamine. The assay was performed in MES pH 6.0 with 5% DMSO, 150 mM NaCl, 0.02% (v/v) Brij-35 and 5 mM $CaCl_2$. Assays were performed in triplicate. Numbers indicate arylsulfamate compounds as in Figure 1.

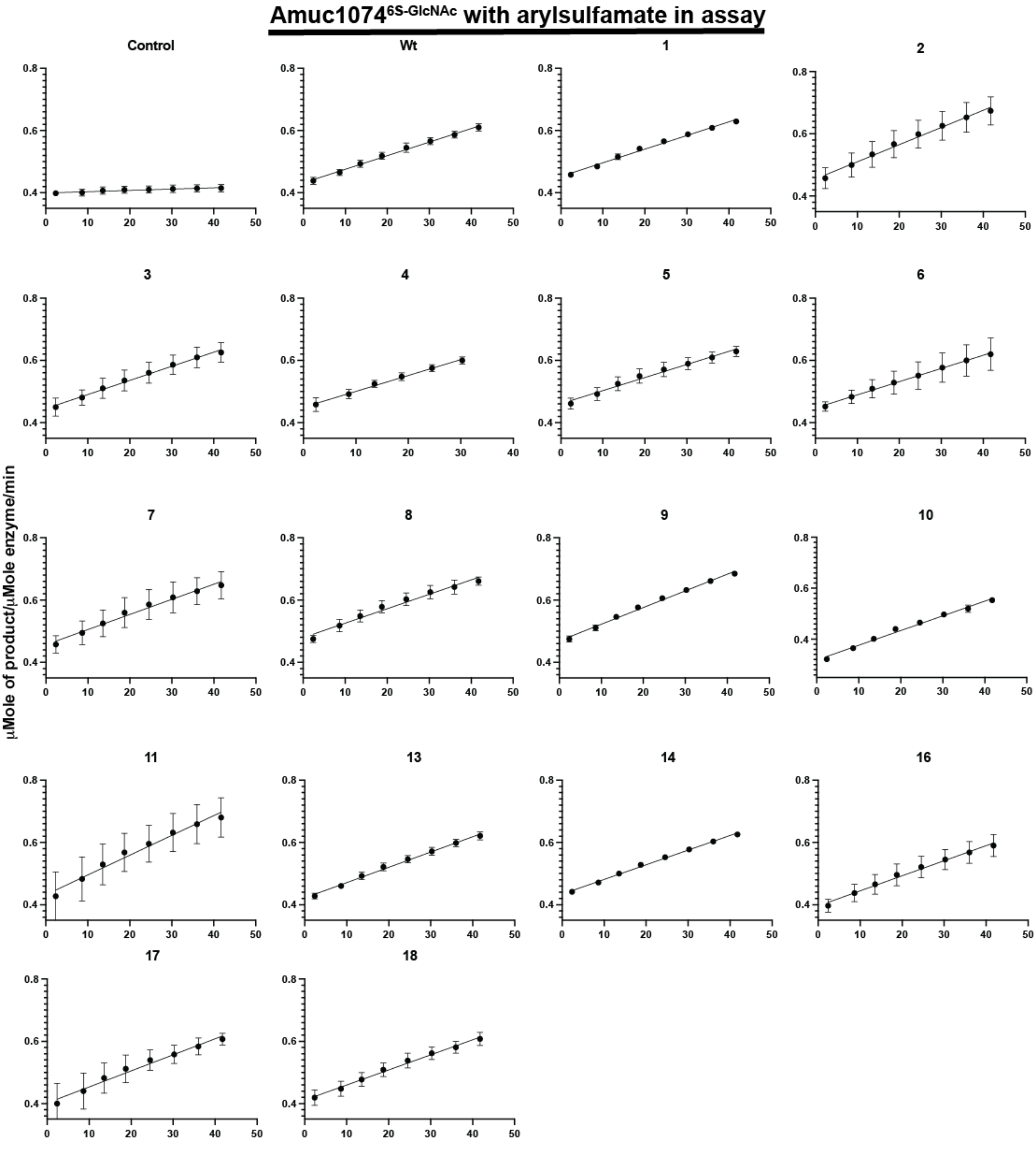


**Supplementary figure 16. Kinetic curves of Amuc1074$^{6S-GlcNAc}$ assayed against BODIPY labelled 6S-*N*-acetylglucosamine with and without arylsulfamate inhibitors**

Amuc1074$^{6S-GlcNAc}$, at a concentration of 6.4 µM, was assayed against 1 µM BODIPY labelled 6S-*N*-acetylglucosamine with and without 1 mM of various arylsulfamate inhibitors included in the assay. The assay was performed in 100 mM of MES pH 6.0 with 5% DMSO, 150 mM NaCl, 0.02% (v/v) Brij-35 and 5 mM $CaCl_2$. Assays were performed in triplicate. Numbers indicate arylsulfamate compounds as in Figure 1.

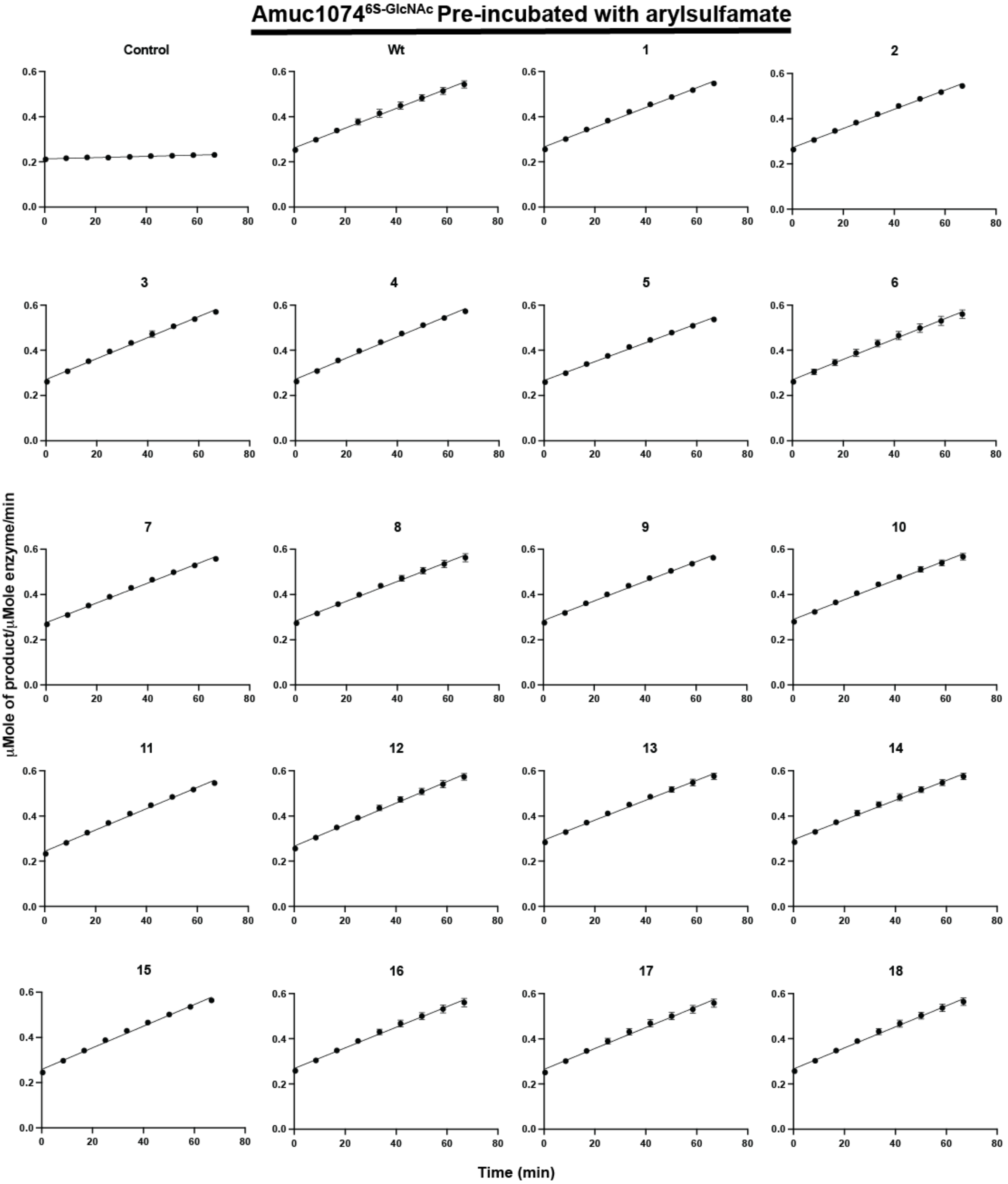


**Supplementary figure 17. Kinetic curves of Amuc1074$^{6S\text{-}GlcNAc}$ assayed against BODIPY labelled 6S-*N*-acetylglucosamine after pre-incubation of the enzyme with arylsulfamate inhibitors**

Amuc1074$^{6S\text{-}GlcNAc}$, which had been incubated for ~24h with 1 mM of the appropriate arylsulfamate inhibitor, was assayed using a concentration of 6.08 μM against 1 μM BODIPY labelled 6S-*N*-acetylglucosamine. The assay was performed in 100 mM of MES pH 6.0 with 5% DMSO, 150 mM NaCl, 0.02% (v/v) Brij-35 and 5 mM $CaCl_2$. Assays were performed in triplicate. Numbers indicate arylsulfamate compounds as in Figure 1.

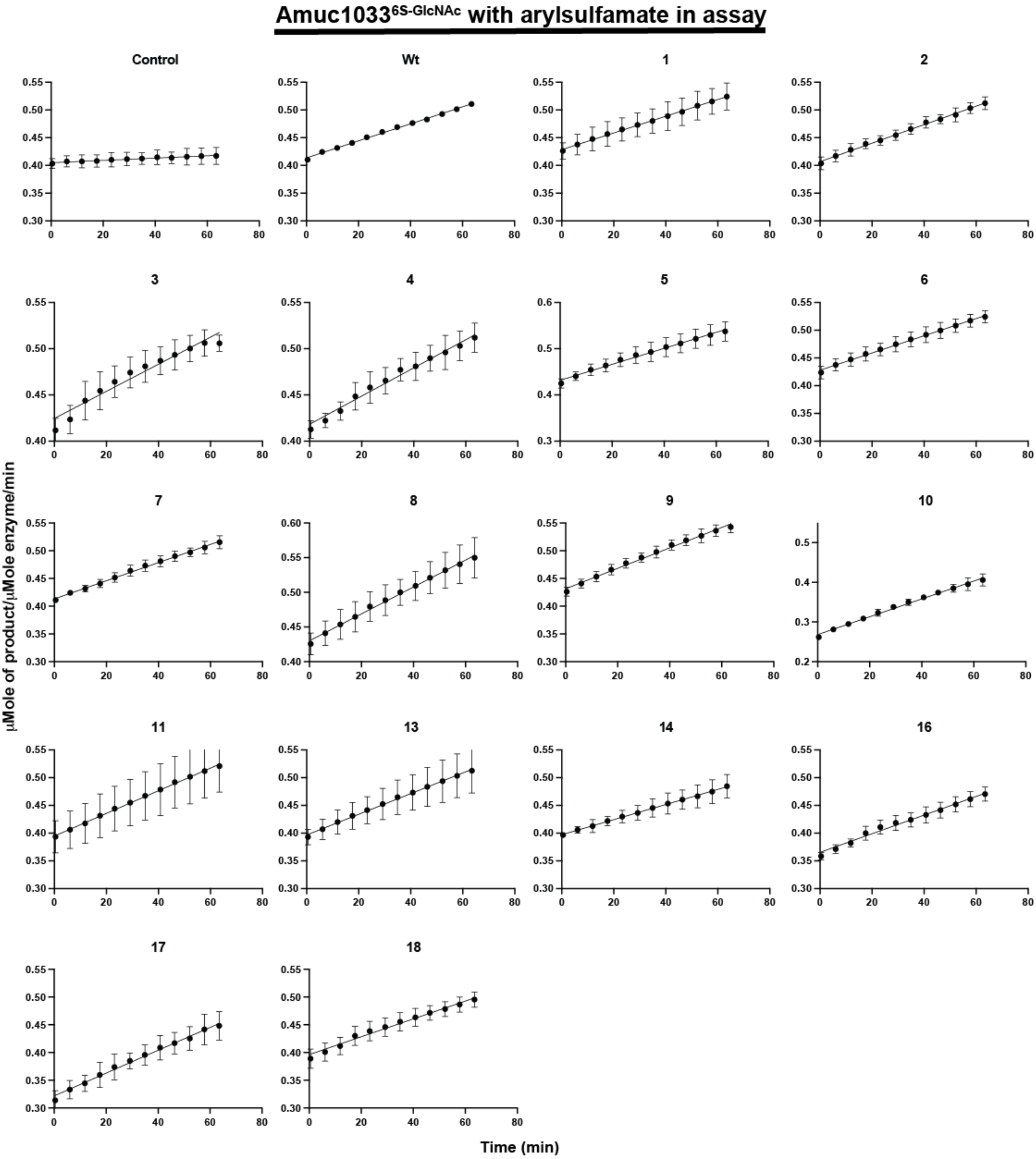


**Supplementary figure 18. Kinetic curves of Amuc1033$^{6S\text{-}GlcNAc}$ assayed against BODIPY labelled 6S-*N*-acetylglucosamine with and without arylsulfamate inhibitors**

Amuc1033$^{6S\text{-}GlcNAc}$, at a concentration of 4.6 μM, was assayed against 1 μM BODIPY labelled 6S-*N*-acetylglucosamine with and without 1 mM of various arylsulfamate inhibitors included in the assay. The assay was performed in 100 mM of MES pH 6.0 with 5% DMSO, 150 mM NaCl, 0.02% (v/v) Brij-35 and 5 mM $CaCl_2$. Assays were performed in triplicate. Numbers indicate arylsulfamate compounds as in Figure 1.

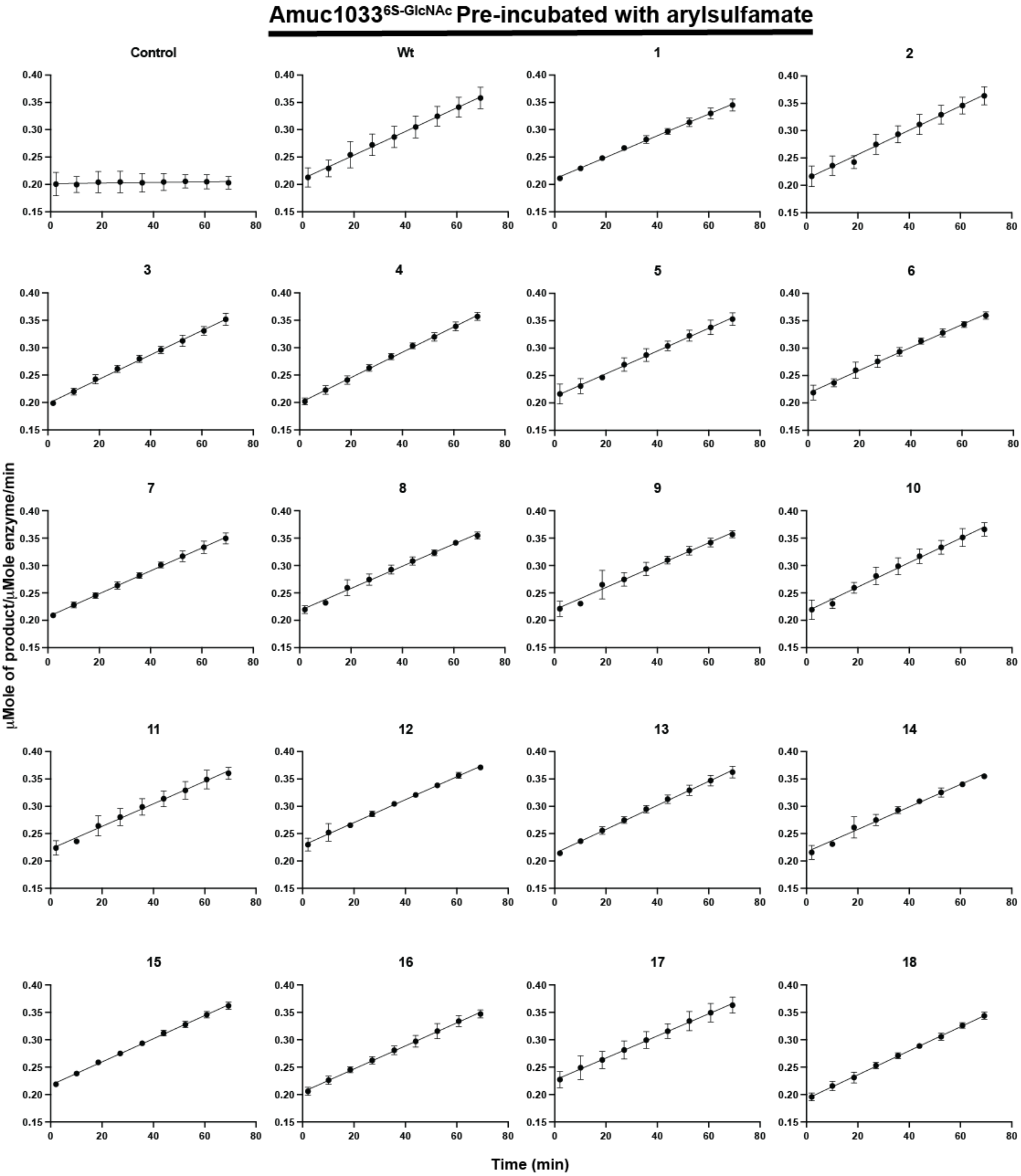


**Supplementary figure 19. Kinetic curves of Amuc1033$^{6S\text{-}GlcNAc}$ assayed against BODIPY labelled 6S-*N*-acetylglucosamine after pre-incubation of the enzyme with arylsulfamate inhibitors**

Amuc1033$^{6S\text{-}GlcNAc}$, which had been incubated for ~24h with 1 mM of the appropriate arylsulfamate inhibitor, was assayed using a concentration of 4.37 µM against 1 µM BODIPY labelled 6S-*N* acetylglucosamine. The assay was performed in 100 mM of MES pH 6.0 with 5% DMSO, 150 mM NaCl, 0.02% (v/v) Brij-35 and 5 mM $CaCl_2$. Assays were performed in triplicate. Numbers indicate arylsulfamate compounds as in Figure 1.

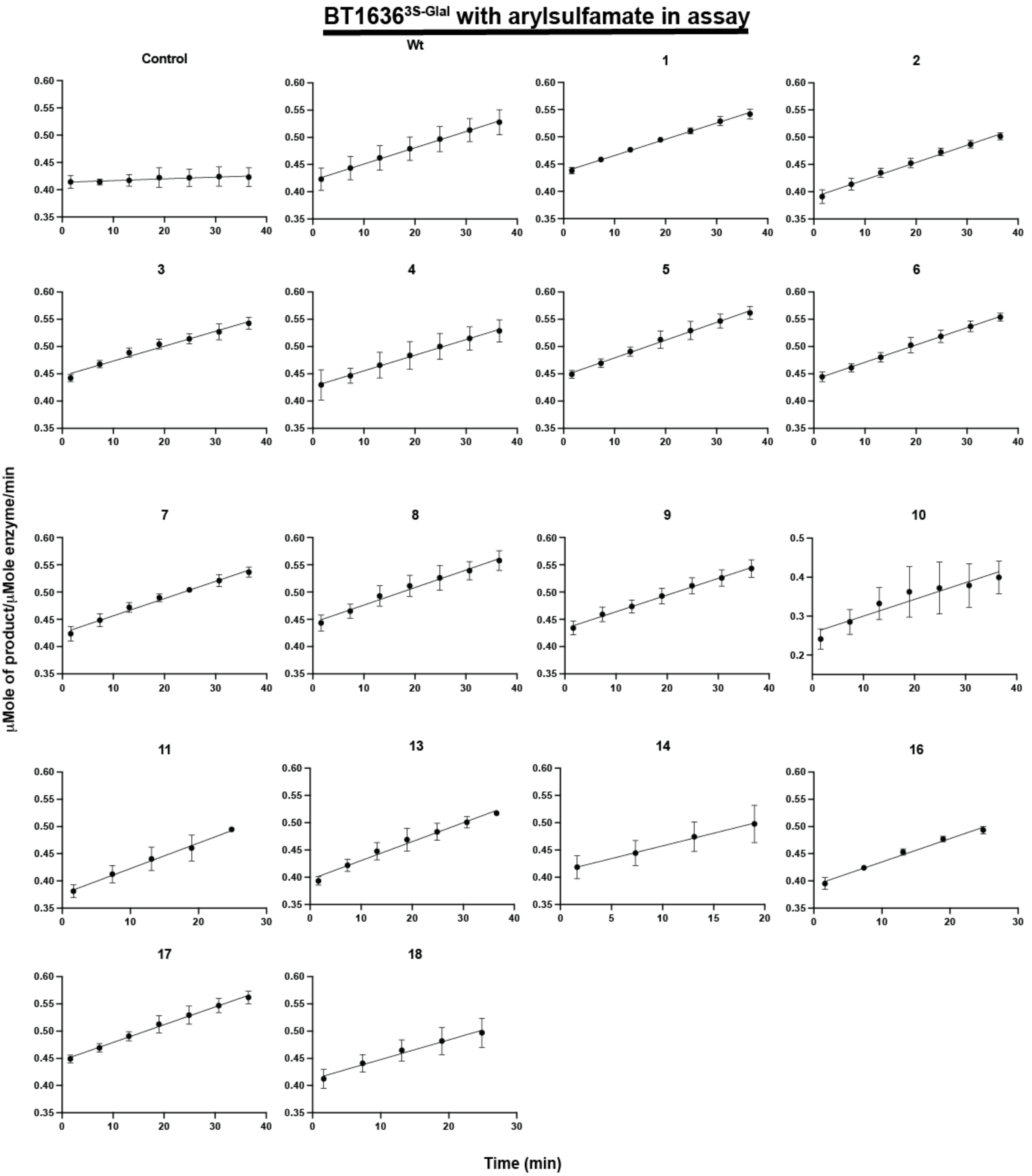


**Supplementary figure 20. Kinetic curves of BT1636$^{3S-Gal}$ assayed against BODIPY labelled 3S-galactose with and without arylsulfamate inhibitors**

BT1636$^{3S-Gal}$, at a concentration of 400 nM, was assayed against 1 µM BODIPY labelled 3S-galactose with and without 1 mM of various arylsulfamate inhibitors included in the assay. The assay was performed in 100 mM of MES pH 6.0 with 5% DMSO, 150 mM NaCl, 0.02% (v/v) Brij-35 and 5 mM $CaCl_2$. Assays were performed in triplicate. Numbers indicate arylsulfamate compounds as in Figure 1.

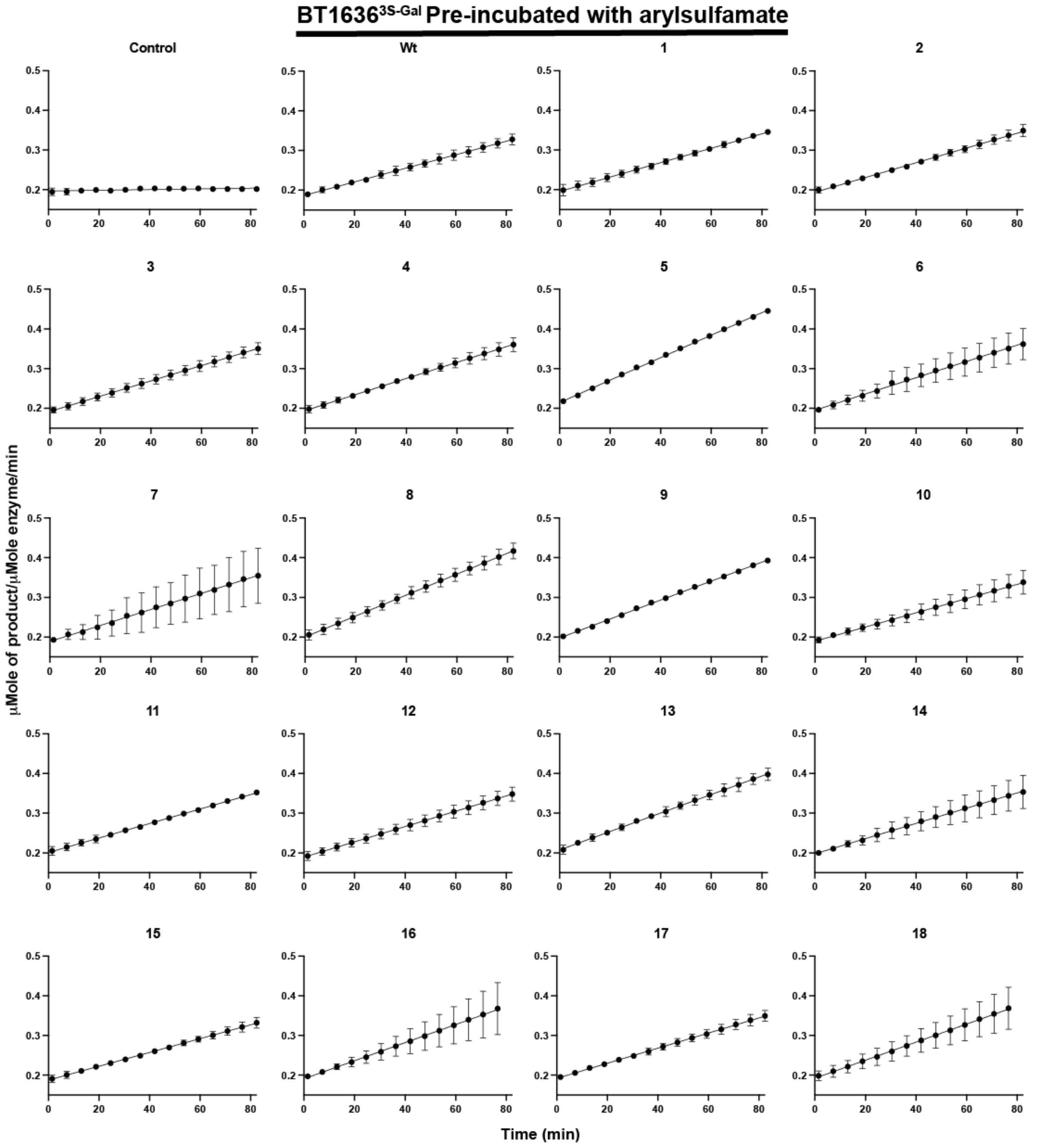


**Supplementary figure 21. Kinetic curves of BT1636$^{3S\text{-}Gal}$ assayed against BODIPY labelled 3S-galactose sulfate after pre-incubation of the enzyme with arylsulfamate inhibitors**

BT1636$^{3S\text{-}Gal}$, which had been incubated for ~24h with 1 mM of the appropriate arylsulfamate inhibitor, was assayed using a concentration of 200 nM against 1 μM BODIPY labelled 3S-galactose. The assay was performed in 100 mM of MES pH 6.0 with 5% DMSO, 150 mM NaCl, 0.02% (v/v) Brij-35 and 5 mM $CaCl_2$. Assays were performed in triplicate. Numbers indicate arylsulfamate compounds as in Figure 1.

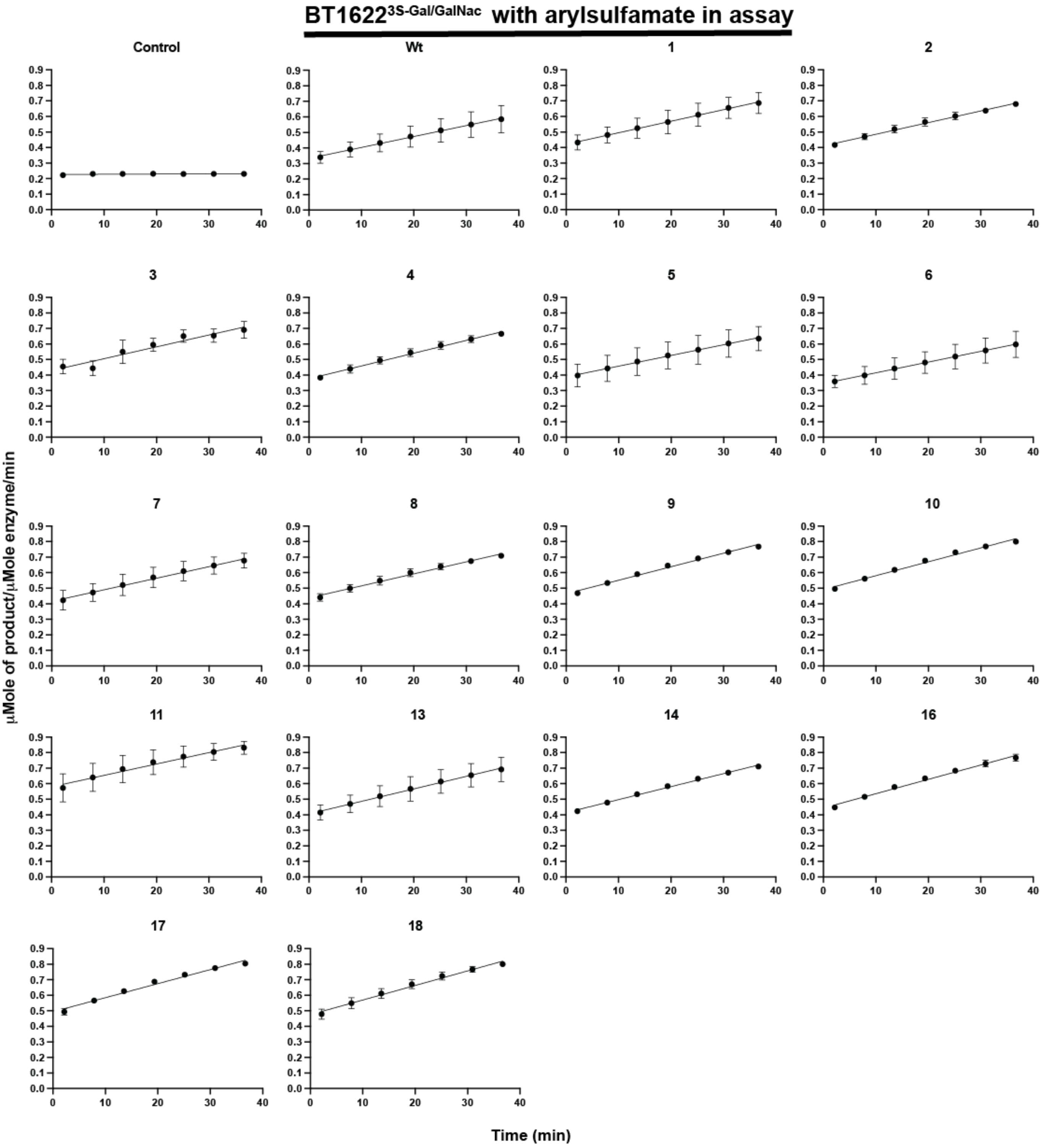


**Supplementary figure 22. Kinetic curves of BT1622$^{3S-Gal/GalNAc}$ assayed BODIPY labelled 3S-*N*-acetylgalactosamine with and without arylsulfamate inhibitors**

BT1622$^{3S-Gal/GalNAc}$, at a concentration of 6.5 µM, was assayed against 1 µM BODIPY labelled 3S-*N*-acetylgalactosamine with and without 1 mM of various arylsulfamate inhibitors included in the assay. The assay was performed in 100 mM of BTP pH 8.5 with 5% DMSO, 150 mM NaCl, 0.02% (v/v) Brij-35 and 5 mM $CaCl_2$. Assays were performed in triplicate. Numbers indicate arylsulfamate compounds as in Figure 1.

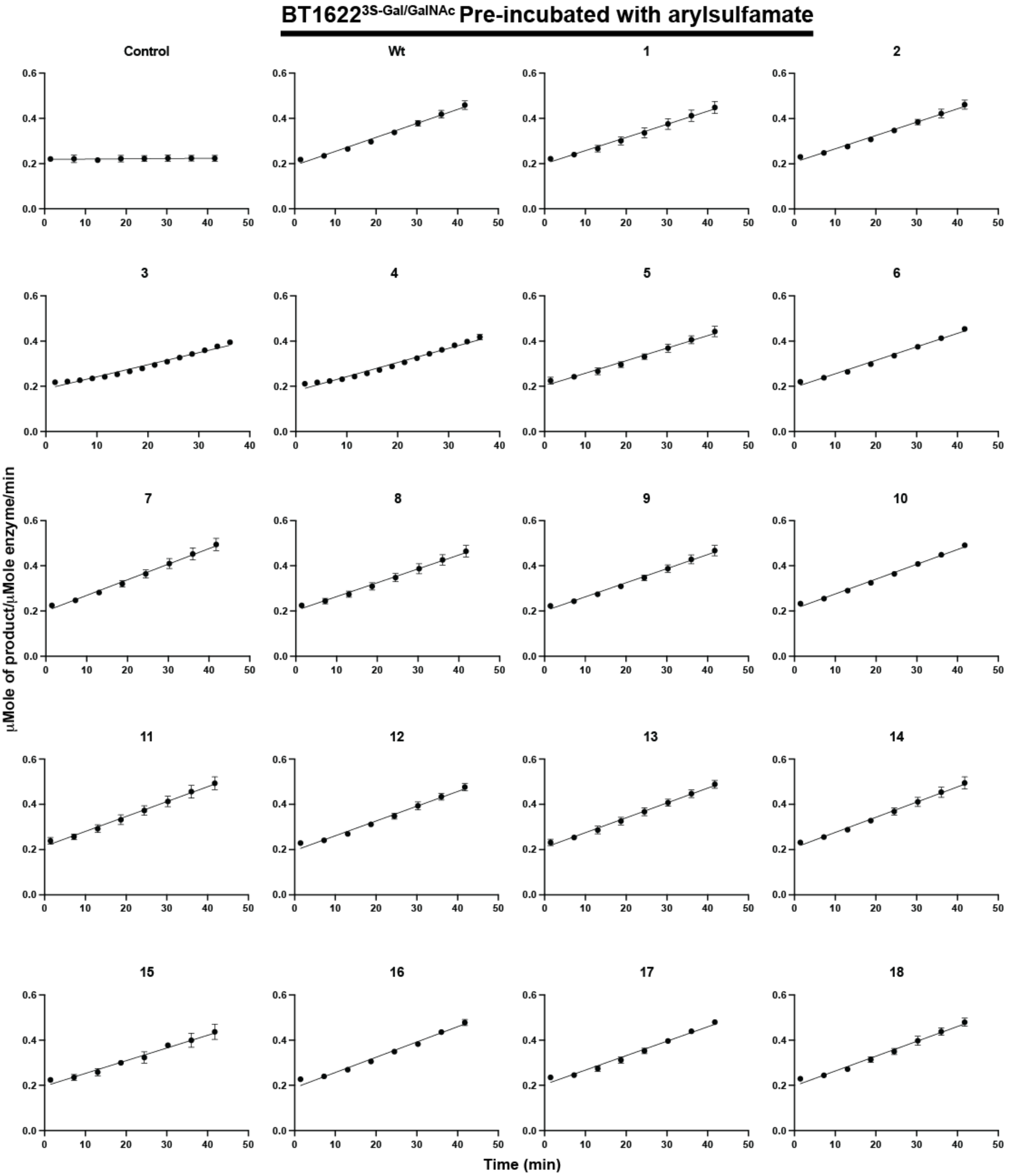


**Supplementary figure 23. Kinetic curves of BT1622$^{3S\text{-}Gal/GalNAc}$ assayed against BODIPY labelled 3S-*N*-acetylgalactosamine after pre-incubation of the enzyme with arylsulfamate inhibitors**
BT1622$^{3S\text{-}Gal/GalNAc}$, which had been incubated for ~24h with 1 mM of the appropriate arylsulfamate inhibitor, was assayed using a concentration of 2 μM against 1 μM BODIPY labelled 3S-*N*-acetylgalactosamine. The assay was performed in 100 mM of BTP pH 8.5 with 5% DMSO, 150 mM NaCl, 0.02% (v/v) Brij-35 and 5 mM $CaCl_2$. Assays were performed in triplicate. Numbers indicate arylsulfamate compounds as in Figure 1.

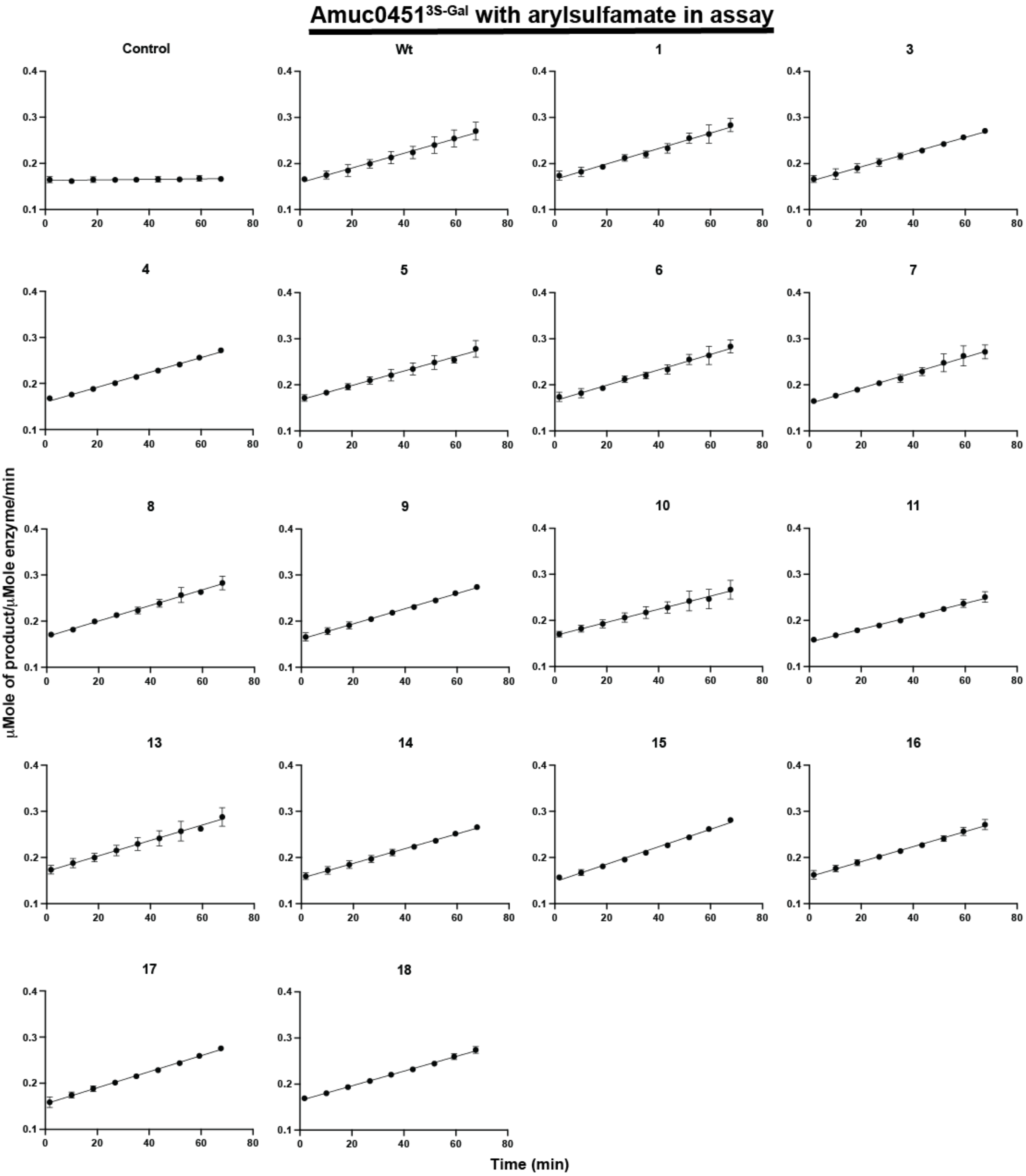


**Supplementary figure 24. Kinetic curves of Amuc0451$^{3S-Gal}$ assayed against BODIPY labelled 3S-galactose with and without arylsulfamate inhibitors**

Amuc0451$^{3S-Gal}$, at a concentration of 15.6 μM, was assayed against 1 μM BODIPY labelled 3S-galactose with and without 1 mM of various arylsulfamate inhibitors included in the assay. The assay was performed in 100 mM of MES pH 6.0 with 5% DMSO, 150 mM NaCl, 0.02% (v/v) Brij-35 and 5 mM $CaCl_2$. Assays were performed in triplicate. Numbers indicate arylsulfamate compounds as in Figure 1.

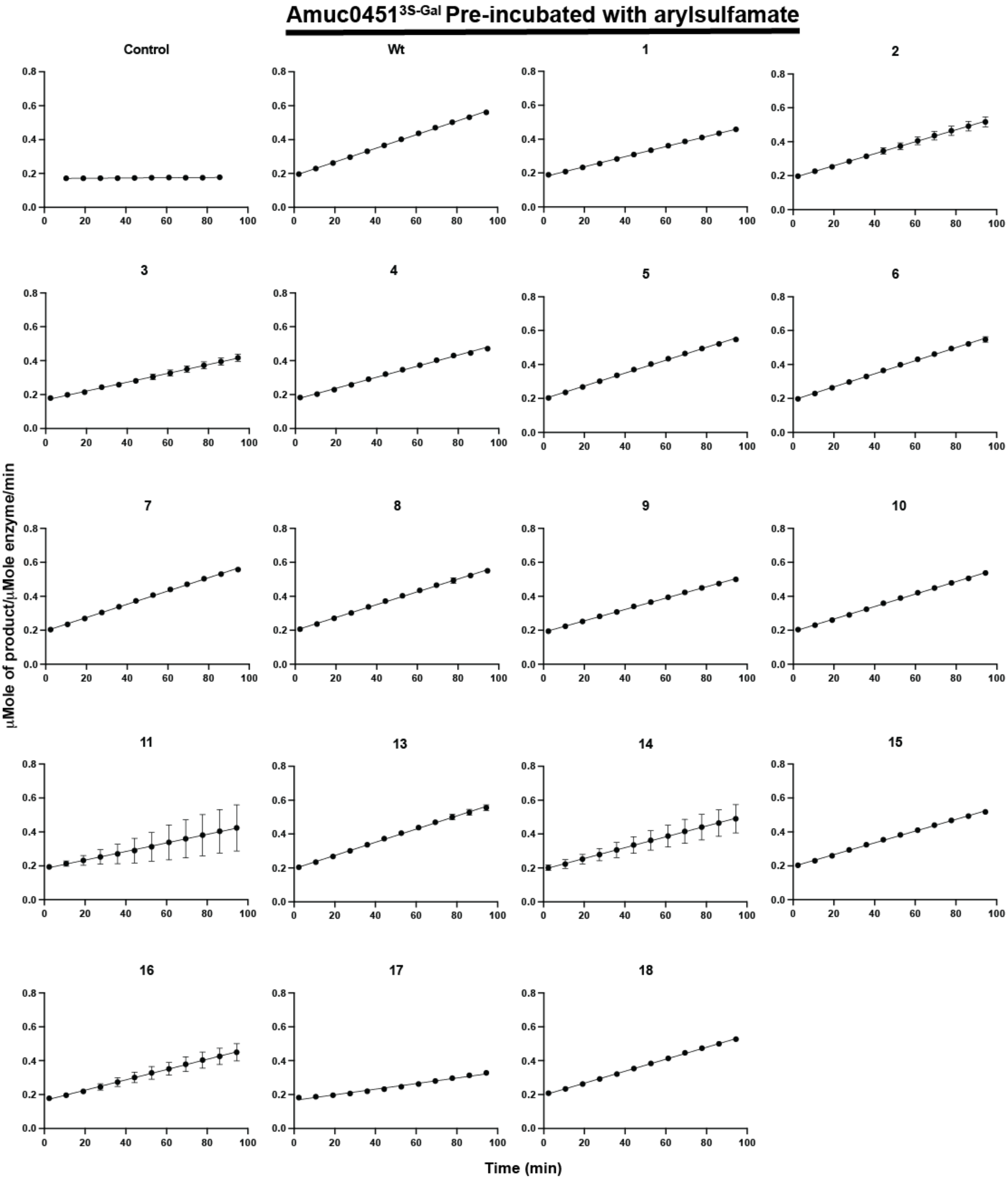


**Supplementary figure 25. Kinetic curves of Amuc0451[3S-Gal] assayed against BODIPY labelled 3S-galactose after pre-incubation of the enzyme with arylsulfamate inhibitors**

Amuc0451[3S-Gal], which had been incubated for ~24h with 1 mM of the appropriate arylsulfamate inhibitor, was assayed using a concentration of 15.6 µM against 1 µM BODIPY labelled 3S-galactose. The assay was performed in 100 mM of MES pH 6.0 with 5% DMSO, 150 mM NaCl, 0.02% (v/v) Brij-35 and 5 mM $CaCl_2$. Assays were performed in triplicate. Numbers indicate arylsulfamate compounds as in Figure 1.

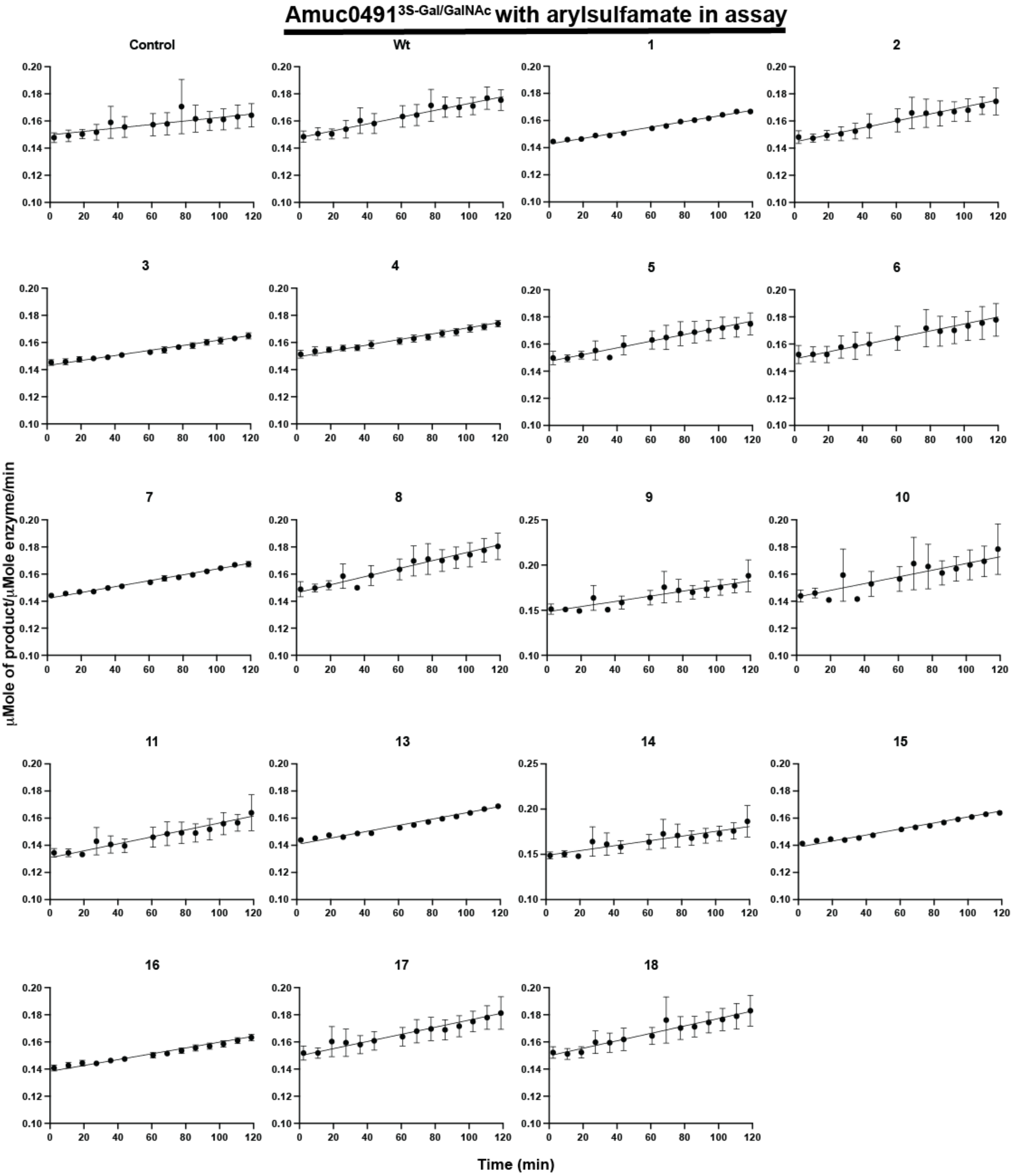


**Supplementary figure 26. Kinetic curves of Amuc0491$^{3S\text{-}Gal/GalNAc}$ assayed against BODIPY labelled 3S-*N*-acetylgalactosamine with and without arylsulfamate inhibitors**

Amuc0491$^{3S\text{-}Gal/GalNAc}$, at a concentration of 3.4 µM, was assayed against 1 µM BODIPY labelled 3S-*N*-acetylgalactosamine with and without 1 mM of various arylsulfamate inhibitors included in the assay. The assay was performed in 100 mM of MES pH 6.0 with 5% DMSO, 150 mM NaCl, 0.02% (v/v) Brij-35 and 5 mM $CaCl_2$. Assays were performed in triplicate. Numbers indicate arylsulfamate compounds as in Figure 1.

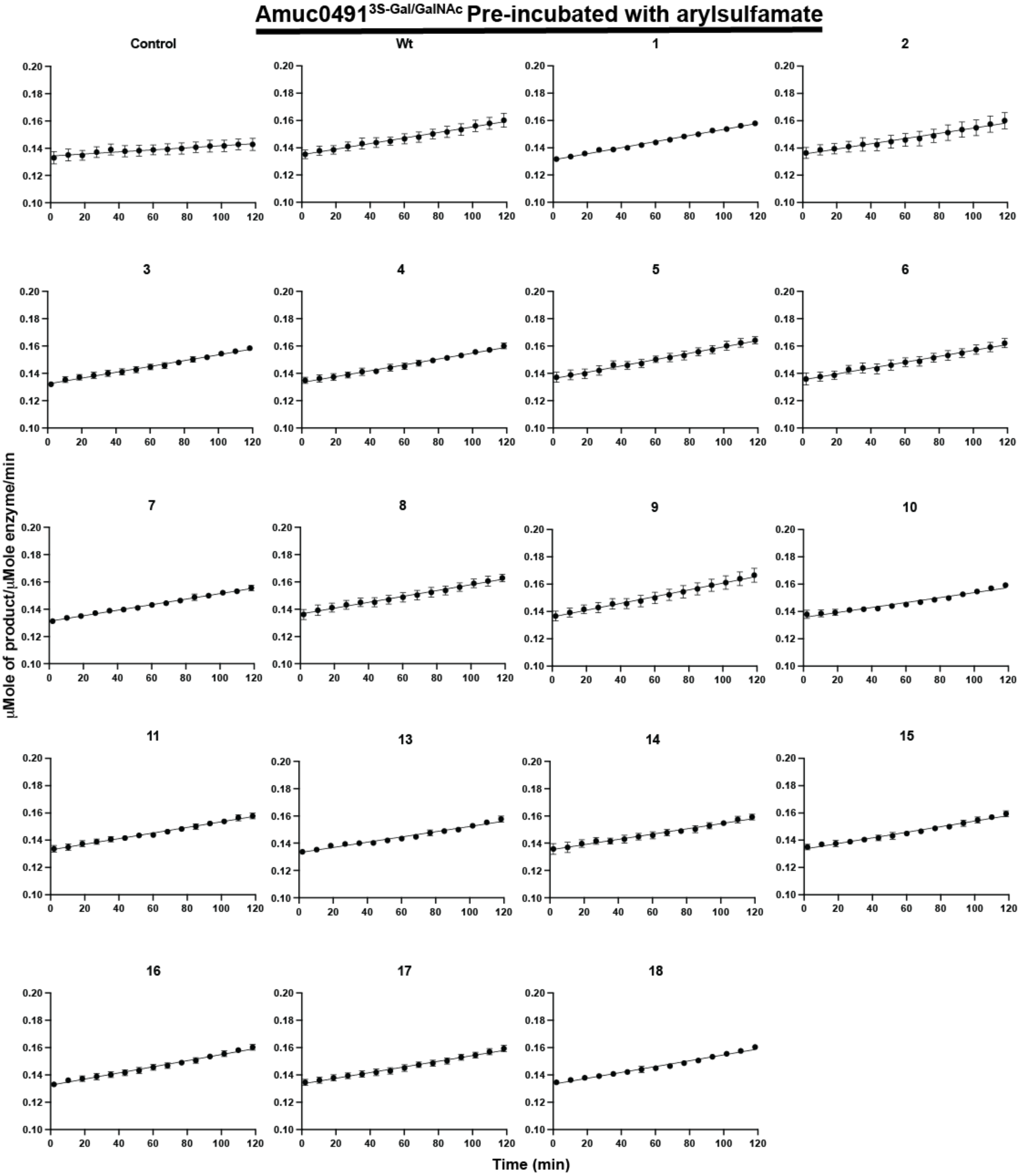


**Supplementary figure 27. Kinetic curves of Amuc0491$^{3S-Gal/GalNAc}$ assayed against BODIPY labelled 3S-*N*-acetylgalactosamine after pre-incubation of the enzyme with arylsulfamate inhibitors**
Amuc0491$^{3S-Gal/GalNAc}$, which had been incubated for ~24h with 1 mM of the appropriate arylsulfamate inhibitor, was assayed using a concentration of 3.2 μM against 1 μM BODIPY labelled 3S-*N*-acetylgalactosamine. The assay was performed in 100 mM of MES pH 6.0 with 5% DMSO, 150 mM NaCl, 0.02% (v/v) Brij-35 and 5 mM $CaCl_2$. Assays were performed in triplicate. Numbers indicate arylsulfamate compounds as in Figure 1.

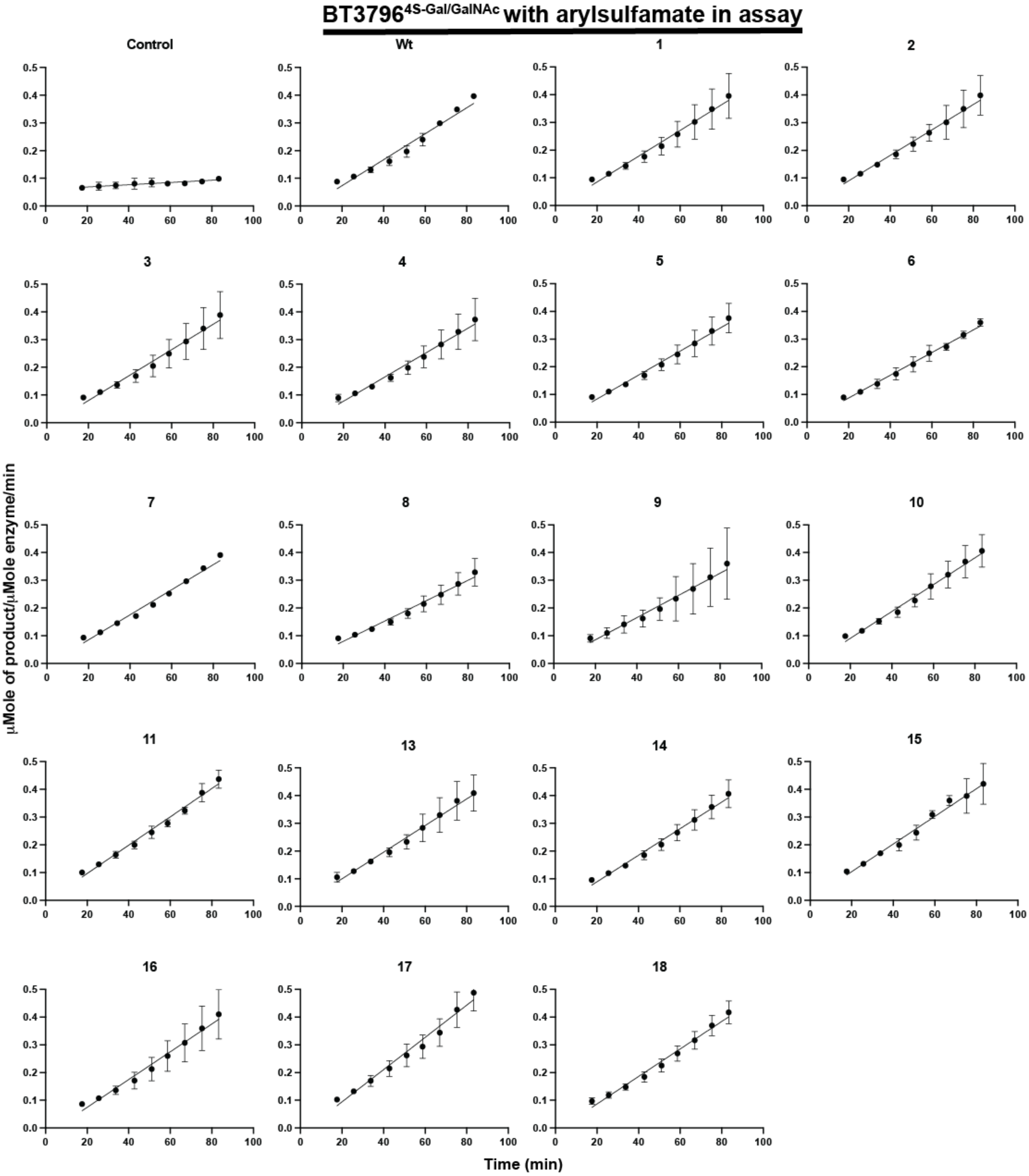


**Supplementary figure 28. Kinetic curves of BT3796$^{4S\text{-}Gal/GalNAc}$ assayed against BODIPY labelled 4S-galactose with and without arylsulfamate inhibitors**

BT3796$^{4S\text{-}Gal/GalNAc}$, at a concentration of 630 nM, was assayed against 1 μM BODIPY labelled 4S-galactose with and without 1 mM of various arylsulfamate inhibitors included in the assay. The assay was performed in 100 mM of BTP pH 8.5 with 5% DMSO, 150 mM NaCl, 0.02% (v/v) Brij-35 and 5 mM $CaCl_2$. Assays were performed in triplicate. Numbers indicate arylsulfamate compounds as in Figure 1.

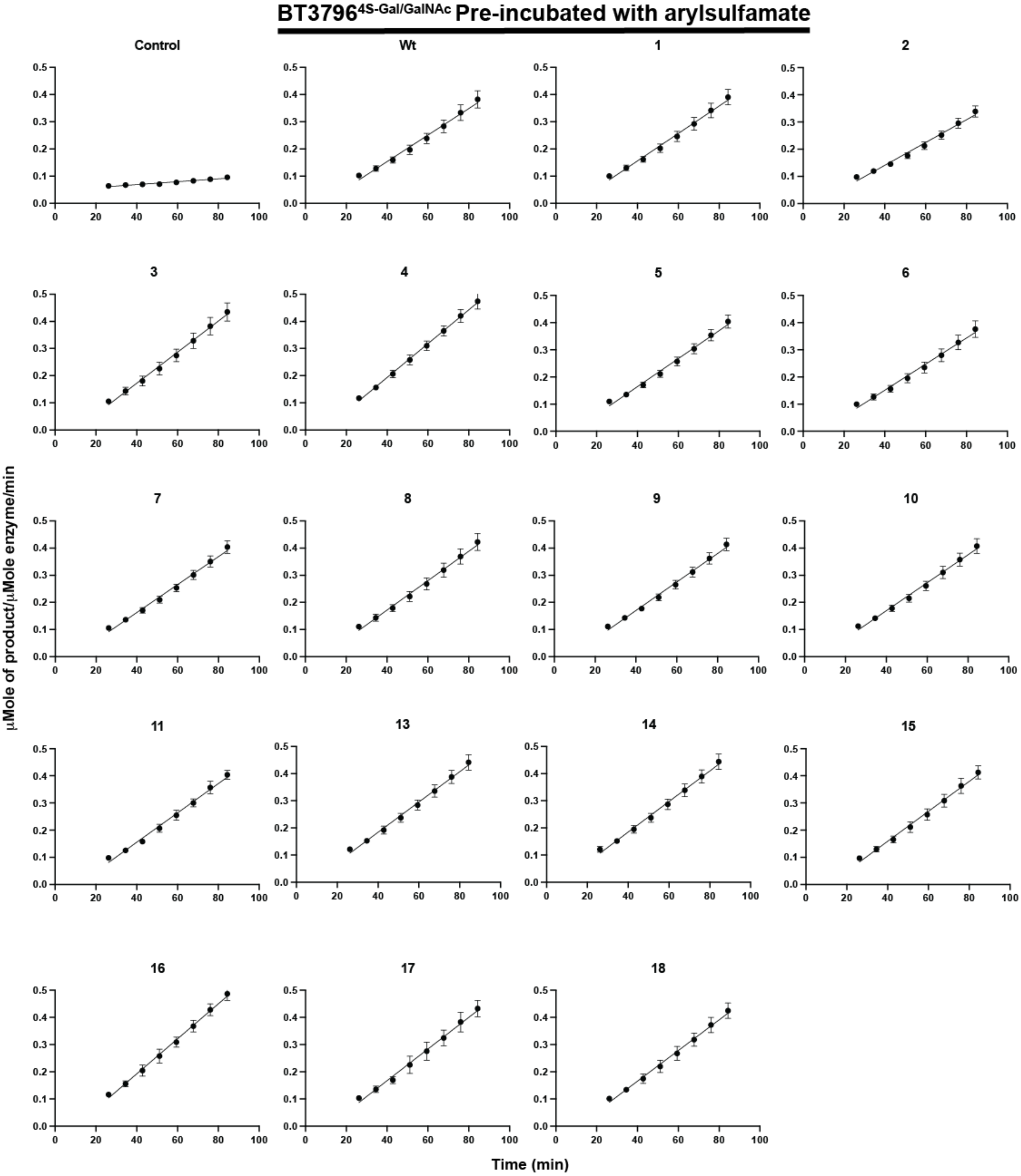


**Supplementary figure 29. Kinetic curves of BT3796$^{4S-Gal/GalNAc}$ assayed against BODIPY labelled 4S-galactose after pre-incubation of the enzyme with arylsulfamate inhibitors**

BT3796$^{4S-Gal/GalNAc}$, which had been incubated for ~24h with 1 mM of the appropriate arylsulfamate inhibitor, was assayed using a concentration of 630 nM against 1 µM BODIPY labelled 4S-galactose. The assay was performed in 100 mM of BTP pH 8.5 with 5% DMSO, 150 mM NaCl, 0.02% (v/v) Brij-35 and 5 mM $CaCl_2$. Assays were performed in triplicate. Numbers indicate arylsulfamate compounds as in Figure 1.

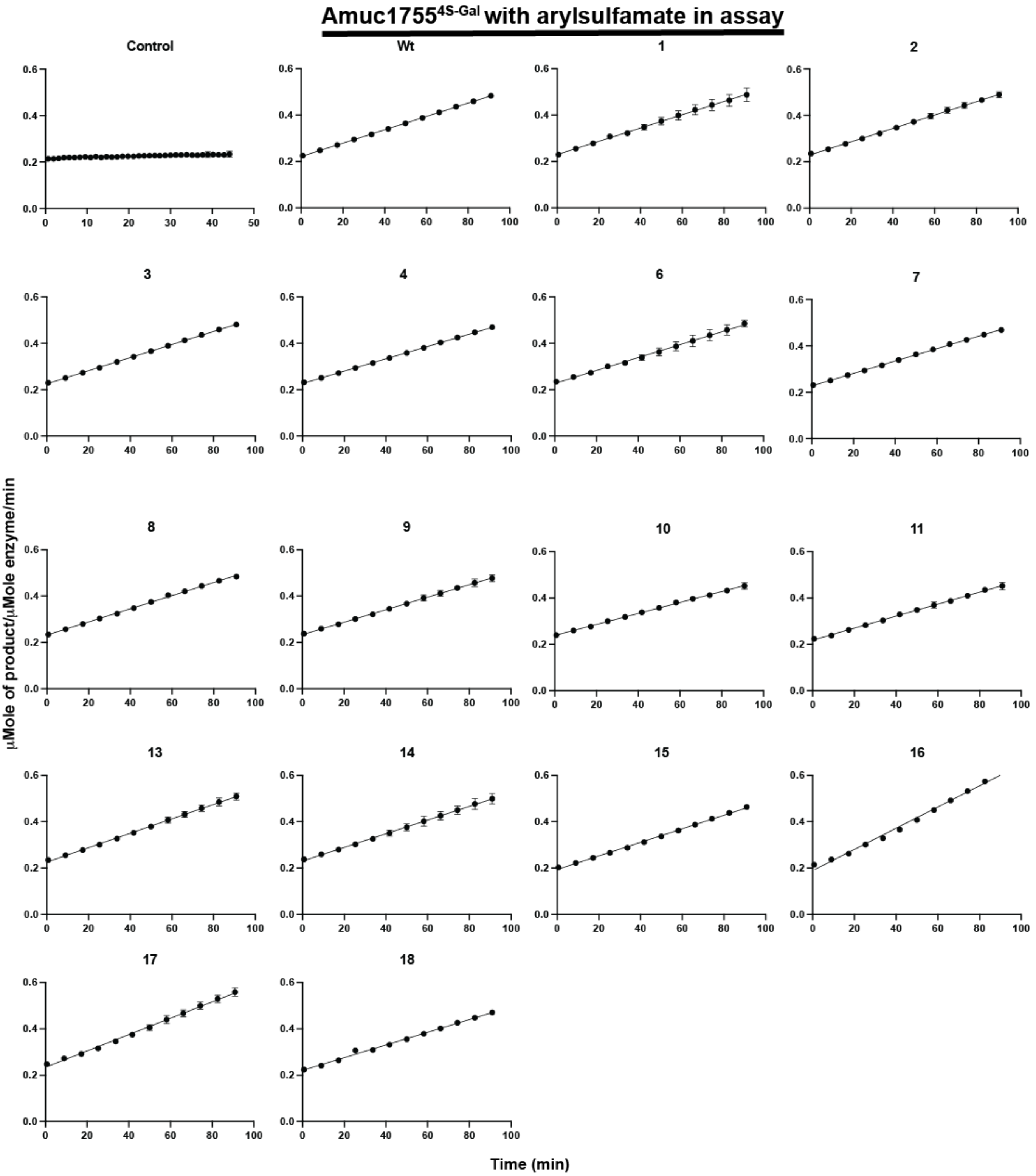


**Supplementary figure 30. Kinetic curves of Amuc1755$^{4S\text{-}Gal}$ assayed against BODIPY labelled 4S-galactose with and without arylsulfamate inhibitors**

Amuc1755$^{4S\text{-}Gal}$, at a concentration of 2.3 μM, was assayed against 1 μM BODIPY labelled 4S-galactose with and without 1 mM of various arylsulfamate inhibitors included in the assay. The assay was performed in 100 mM of MES pH 6.0 with 5% DMSO, 150 mM NaCl, 0.02% (v/v) Brij-35 and 5 mM $CaCl_2$. Assays were performed in triplicate. Numbers indicate arylsulfamate compounds as in Figure 1.

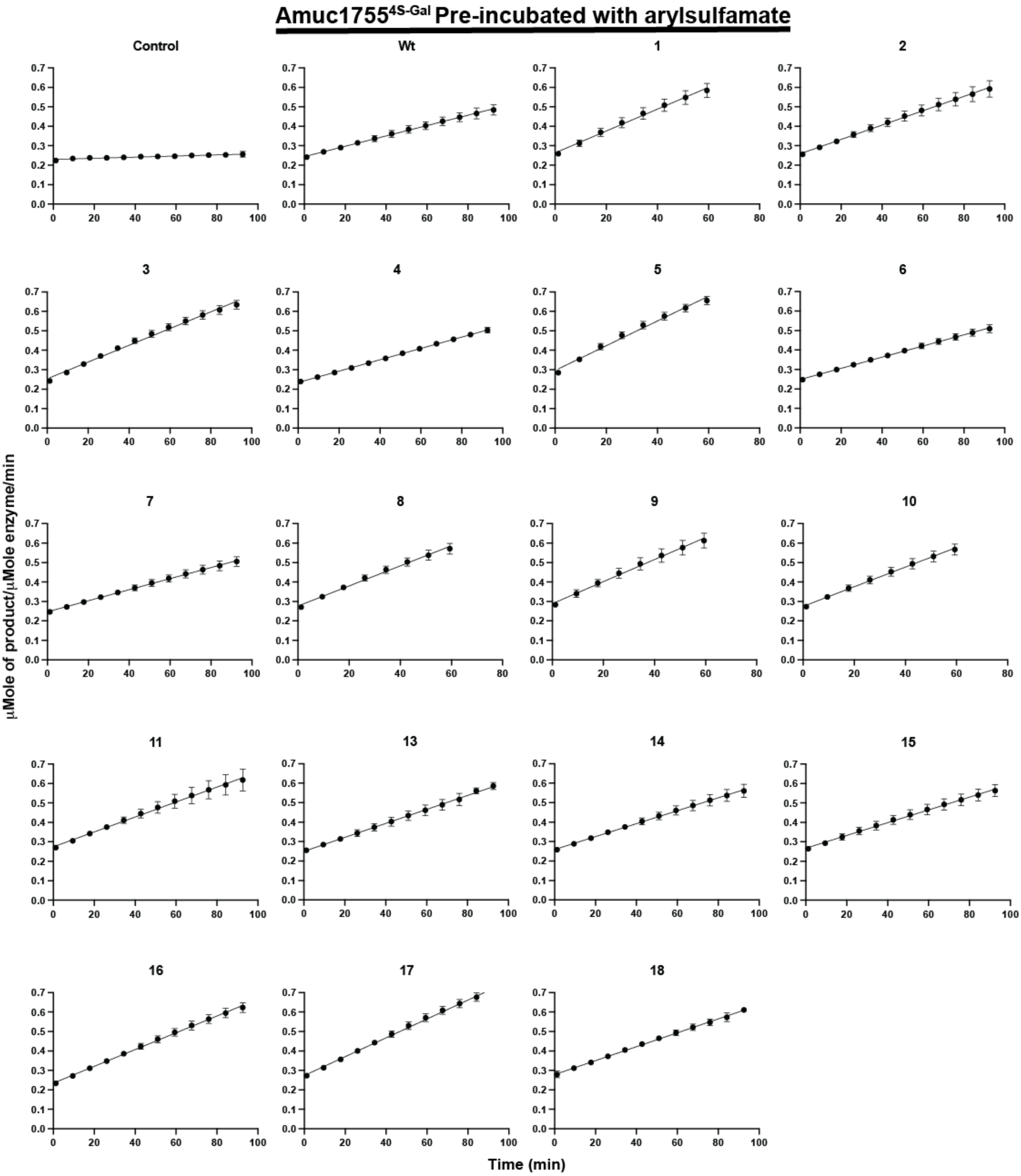


**Supplementary figure 31. Kinetic curves of Amuc1755$^{4S\text{-}Gal}$ assayed against BODIPY labelled 4S-galactose after pre-incubation of the enzyme with arylsulfamate inhibitors**

Amuc1755$^{4S\text{-}Gal}$, which had been incubated for ~24 h with 1 mM of the appropriate arylsulfamate inhibitor, was assayed using a concentration of 2.185 μM against 1 μM BODIPY labelled 4S-galactose. The assay was performed in 100 mM of MES pH 6.0 with 5% DMSO, 150 mM NaCl, 0.02% (v/v) Brij-35 and 5 mM $CaCl_2$. Assays were performed in triplicate. Numbers indicate arylsulfamate compounds as in Figure 1.

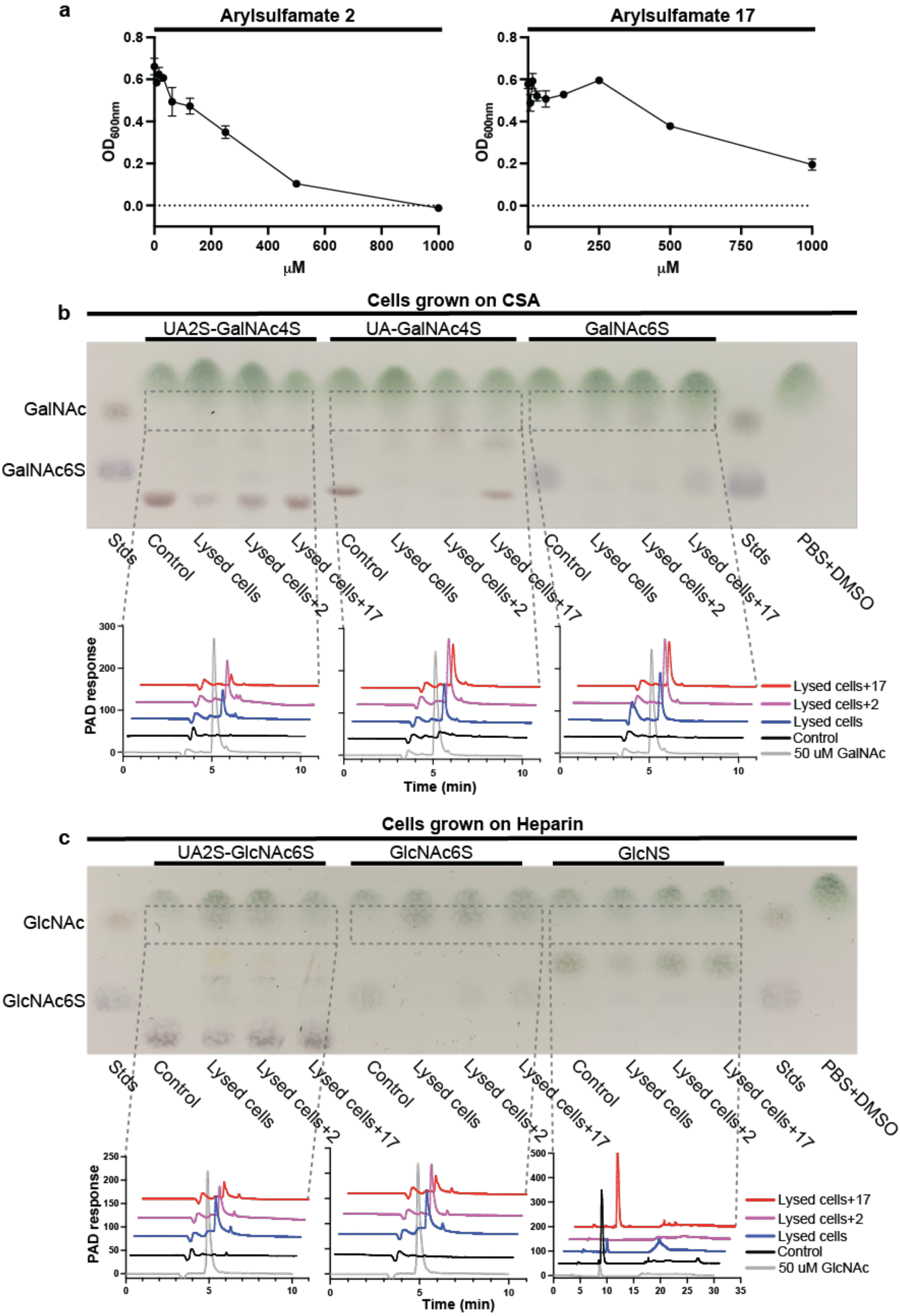
a
Arylsulfamate 2
Arylsulfamate 17
OD600nm
μM
b
Cells grown on CSA
UA2S-GalNAc4S
UA-GalNAc4S
GalNAc6S
GalNAc
GalNAc6S
Stds
Control
Lysed cells
Lysed cells+2
Lysed cells+17
PBS+DMSO
PAD response
Time (min)
Lysed cells+17
Lysed cells+2
Lysed cells
Control
50 uM GalNAc
c
Cells grown on Heparin
UA2S-GlcNAc6S
GlcNAc6S
GlcNS
GlcNAc
GlcNAc6S
Lysed cells+17
Lysed cells+2
Lysed cells
Control
50 uM GlcNAc

**Supplementary figure 32 Cell lysates of Bacteroides thetaiotaomicron incubated with two arylsulfamates inhibitors**

**a.** The effects of varied concentrations of arylsulfamate 2 and 17 on the growth of *B. theta* in BHI media. **b.** Thin layer chromatography (TLC) analysis of cell lysates from B.theta cells grown to mid-exponential phase on CSA as a carbon source, and incubated with arylsulfamates 2 and 17 (Figure 2), mixed 1:1 with the sulfated substrates: O2 sulfated uronic acid β1,3 linked to O4 sulfated N-acetylgalactosmaine (UA2S-GalNAc4S), uronic acid β1,3 linked to O4 sulfated N-acetylgalactosmaine (UA-GalNAc4S), and O6 sulfated N-acetylgalactosmaine (GalNAc6S). Below are high performance anion exchange chromatography (HPAEC) traces showing the production of GalNAc in all instances except the negative control; **c.** Thin layer chromatography (TLC) analysis of cell lysates from B.theta cells grown to mid-exponential phase on Heparin as a carbon source, and incubated with arylsulfamates 2 and 17 (Figure 2), mixed 1:1 with the sulfated substrates: O2 sulfated uronic acid α1,4 linked to O4 sulfated N-acetylgalactosmaine (UA2S-GalNAc4S), O6 sulfated N-acetylglucosmaine (GlcNAc6S), and N-sulfated glucosamine (GlcNS). Below are high performance anion exchange chromatography (HPAEC) traces showing the production of GlcNAc in all instances except the negative control. All reactions were in PBS, with 5 % DMSO, and incubated at 37ºC overnight.

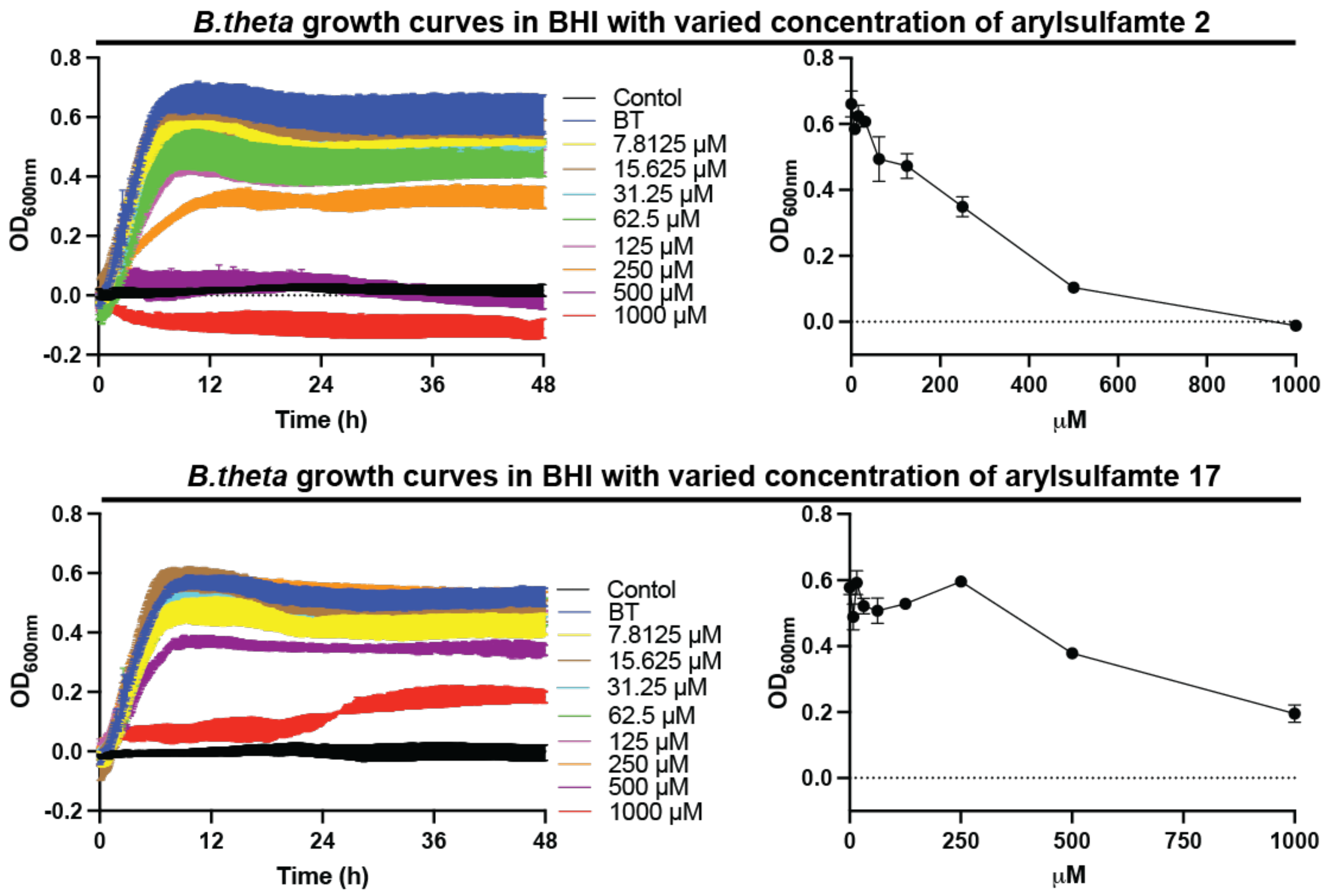


**Supplementary figure 33. The effects of varied concentration arylsulfamates 2 and 17 on the growth of *Bacteroides thetaiotaomicron* VPI-5482 grown in BHI media.**
*Bacteroides thetaiotaomicron* VPI-5482 was grown in BHI, 1 % DMSO, with varied concentrations of arylsulfamate 2 and 17. Data are technical triplicates with the standard error of the mean. Numbers indicate arylsulfamate compound as in Figure 1.

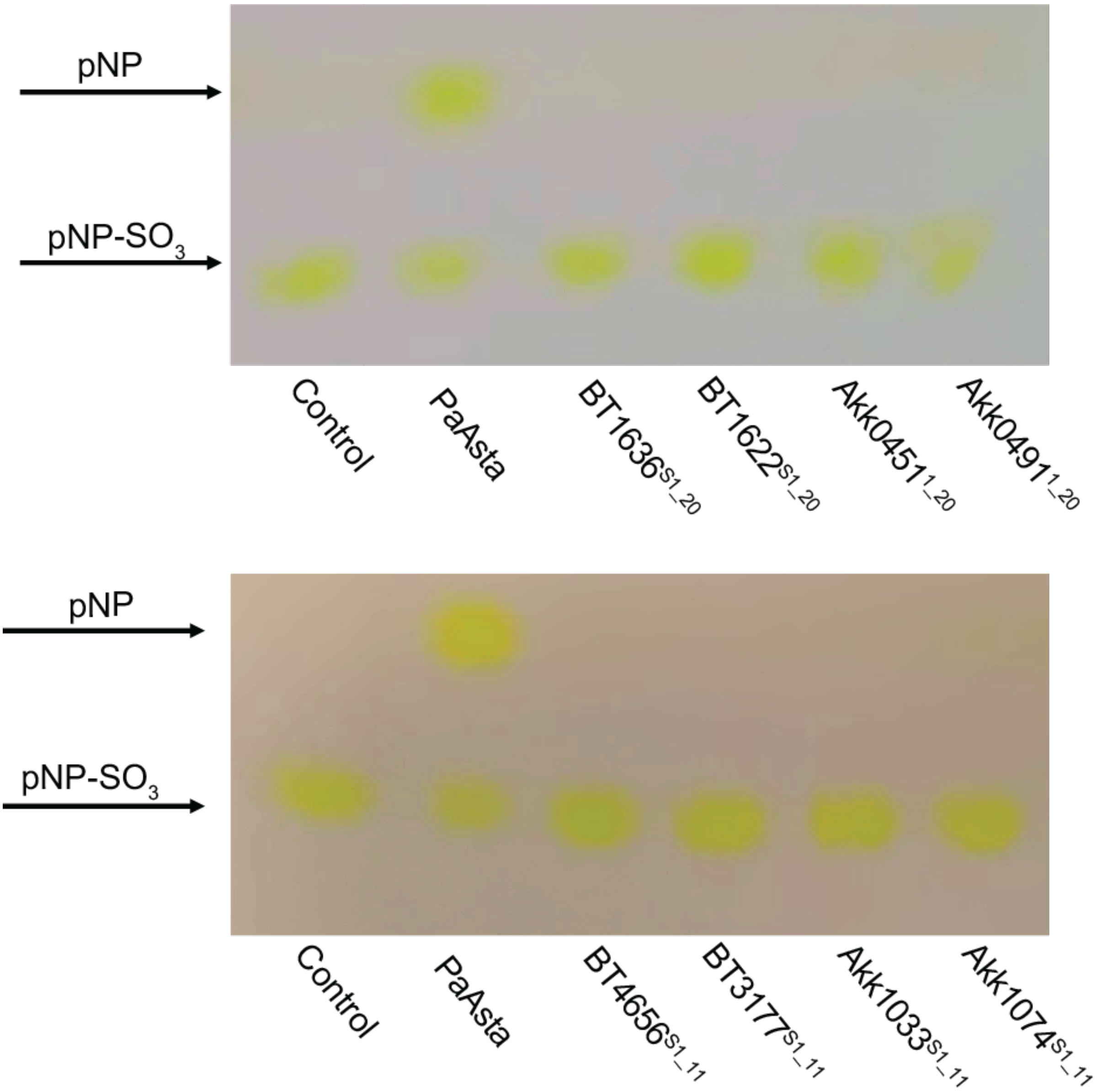


**Supplementary figure 34. Activity of carbohydrate sulfatases on para-nitrophenol sulfate.**

Sulfatases were incubated with 5 mM para-nitrophenol sulfate over night in 100 mM BTP pH 7.0 with 150 mM NaCl and 5 mM CaCl2. Reactions were then ran on silica based thin layer chromatography using a mobile phase of butanol:acetic acid:water (2:1:1) and develop by soaking the plates with 1 M NaOH and heating with a heat gun. It can be seen only the steroid sulfatase PaAsta shows the production of a para-nitrolphenol product.

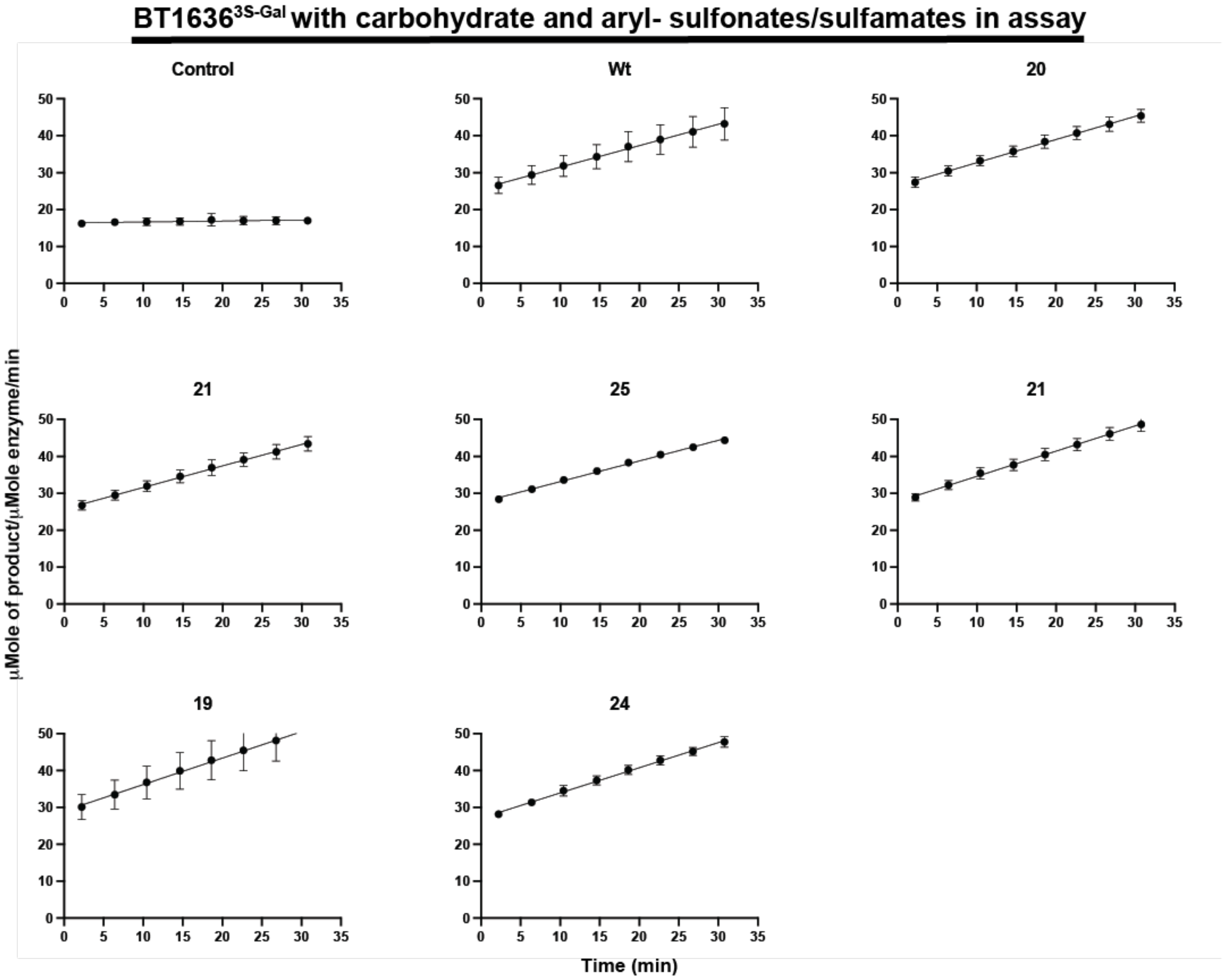


**Supplementary figure 35. Kinetic curves of BT1636$^{3S\text{-}Gal}$ assayed against BODIPY labelled 3S-galactose with and without carbohydrate and aryl- sulfonates/sulfamates**
BT1636$^{3S\text{-}Gal}$, at a concentration of 270 nM, was assayed against 1 μM BODIPY labelled 3S-galactose with and without 1 mM of various arylsulfamate inhibitors included in the assay. The assay was performed in 100 mM of MES pH 6.0 with 5% DMSO, 150 mM NaCl, 0.02% (v/v) Brij-35 and 5 mM $CaCl_2$. Assays were performed in triplicate. Numbers indicate arylsulfamate compounds as in Figure 1.

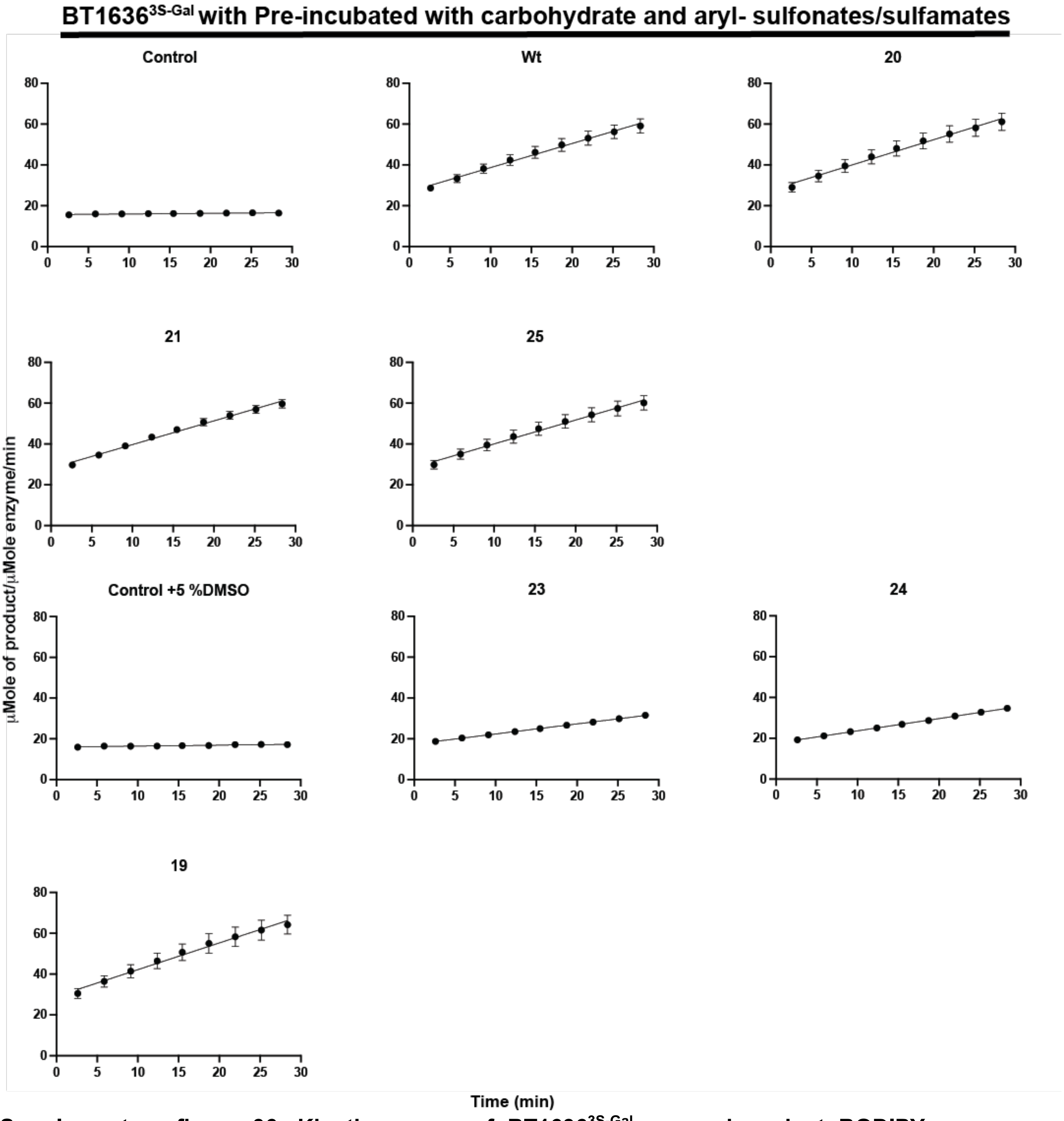


**Supplementary figure 36. Kinetic curves of BT1636$^{3S-Gal}$ assayed against BODIPY labelled 3S-galactose sulfate after pre-incubation of the enzyme with carbohydrate and aryl- sulfonates/sulfamates**

BT1636$^{3S-Gal}$, which had been incubated for ~24h with 1 mM of the appropriate arylsulfamate inhibitor, was assayed using a concentration of 270 nM against 1 μM BODIPY labelled 3S-galactose. The assay was performed in 100 mM of MES pH 6.0 with 5% DMSO, 150 mM NaCl, 0.02% (v/v) Brij-35 and 5 mM $CaCl_2$. Assays were performed in triplicate. Numbers indicate arylsulfamate compounds as in Figure 1.

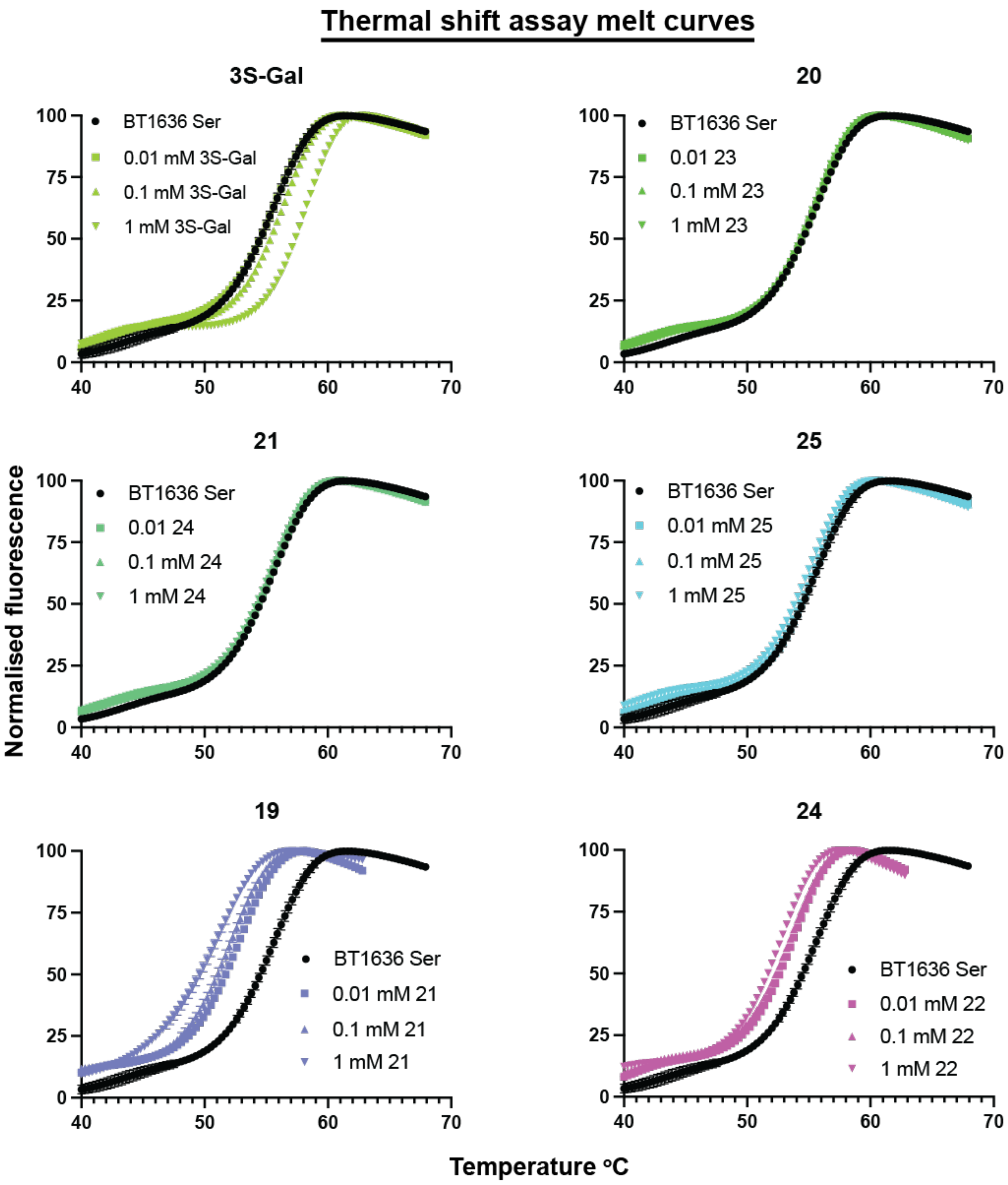


**Supplementary figure 37. Thermal melt curves of BT1636$^{3S\text{-}Gal}$ incubated substrate and various carbohydrate and aryl- sulfonates/sulfamates**

BT1636$^{3S\text{-}Gal}$, at a concentration of 5 μM, was incubated with no compound, and varying concentrations and the effect on its melting temperature monitored. A shift in melting temperature is indicative of an interaction. The assays were performed in 100 mM of BTP pH 7.0 with 5% DMSO, 150 mM NaCl. Assays were performed in triplicate.

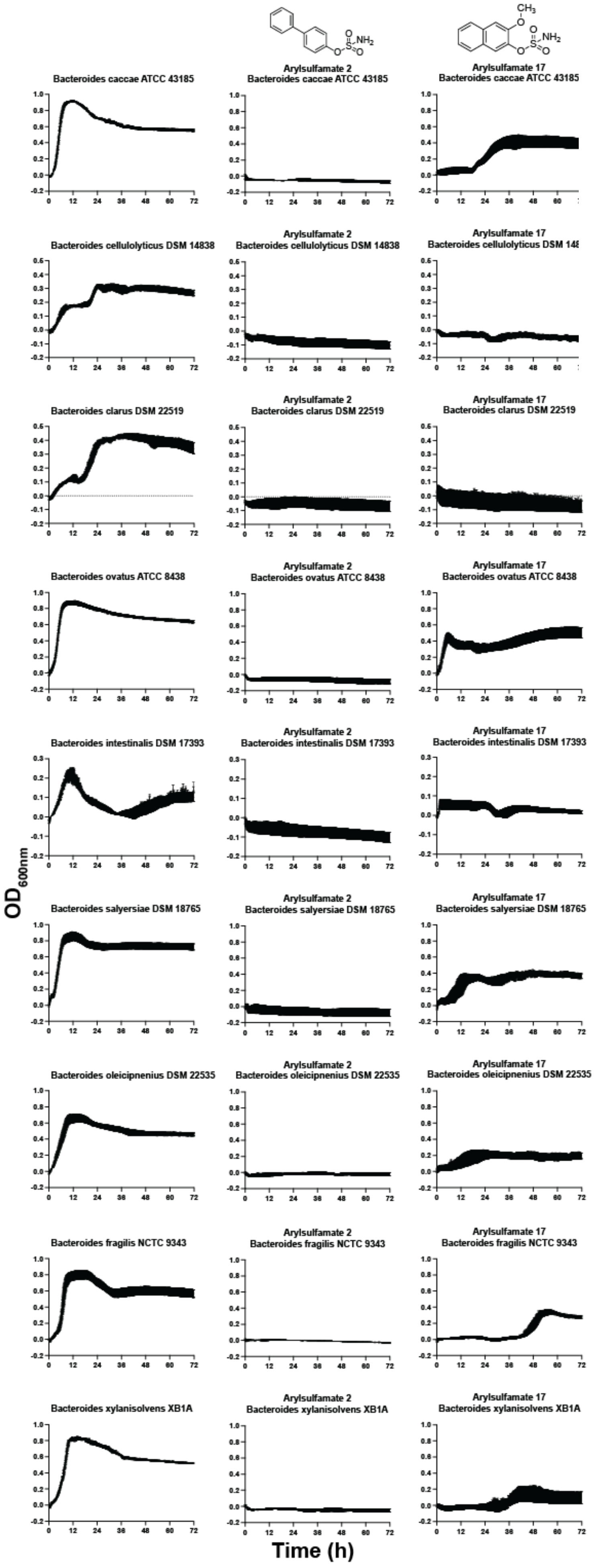


**Supplementary figure 38. The effects of arylsulfamates 2 and 17 on the growth of HGM Bacteroidota species**

Bacteroidota species were grown in BHI, 1% DMSO, with 1 mM of arylsulfamate 2 and 17 and the effects on growth observed. Data are technical triplicates with the standard error of the mean.

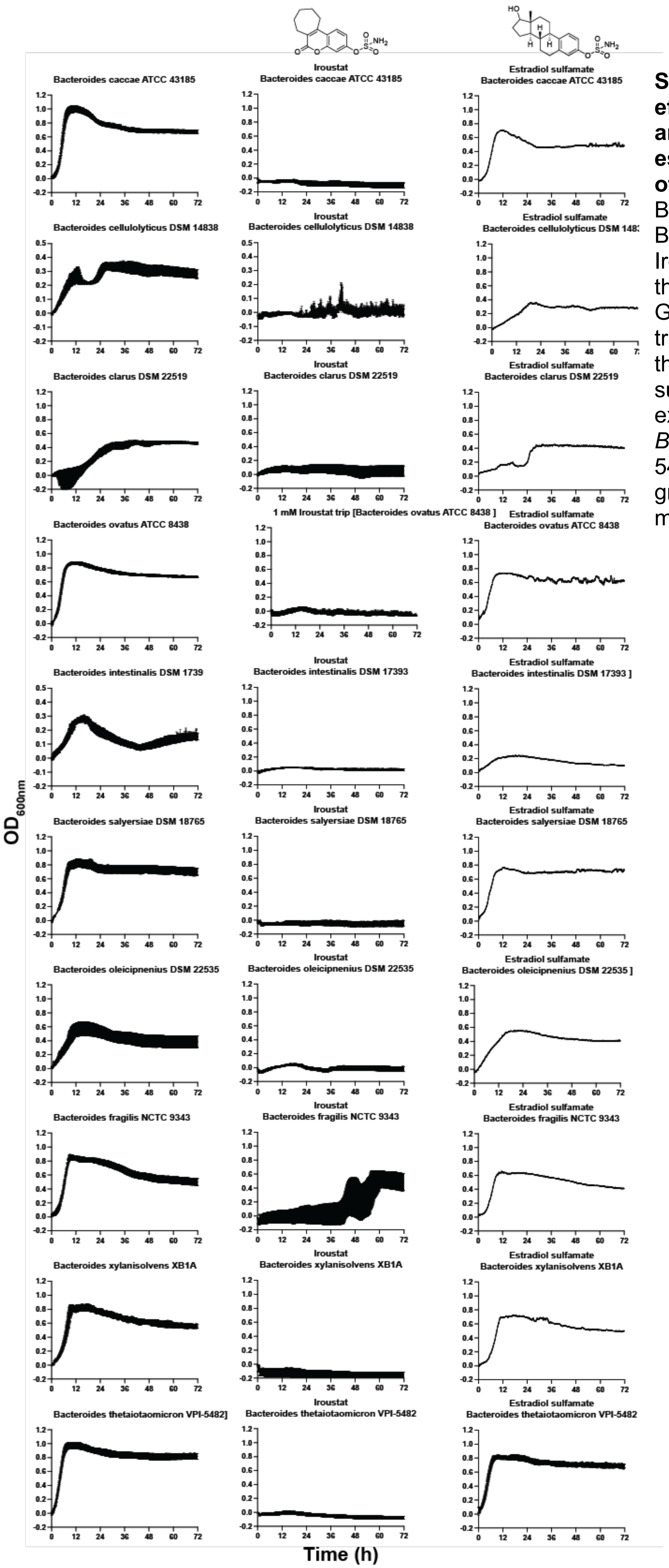


**Supplementary figure 39. The effects of the phase I/II arylsulfamate drugs Irosustat and estradiol sulfamate on the growth of Bacteroidota species**

Bacteroidota species were grown in BHI, 1% DMSO, with 1 mM of Irosustat and estradiol sulfamate and the effects on growth observed. Growths with Irosustat are technical triplicates with the standard error of the mean. Growths with estradiol sulfamate represent single growth experiments with the exception of *Bacteroides thetaiotaomicron* VPI-5482 where the data is from triplicate growths with standard error of the mean.

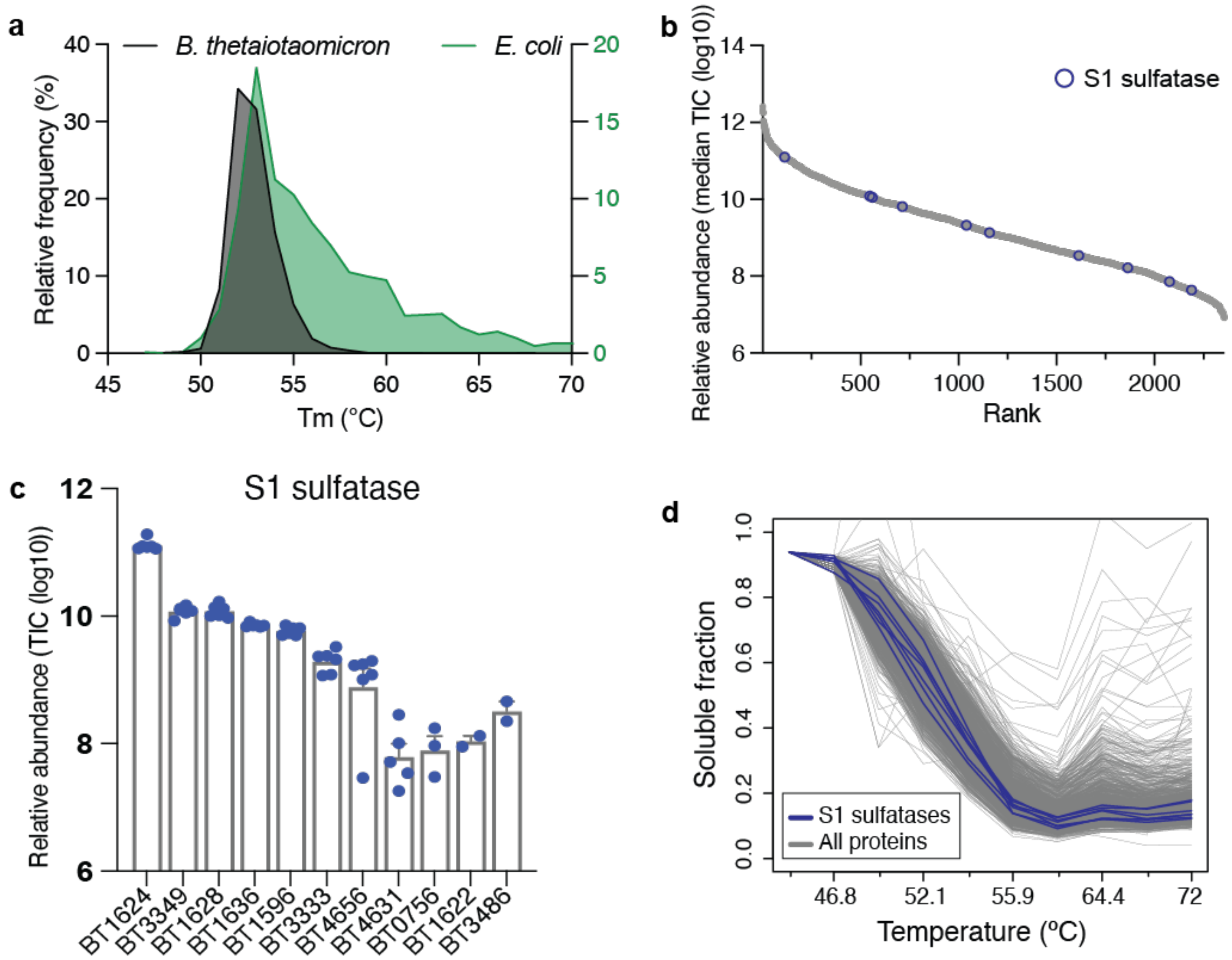


**Supplementary figure 40. Global features of the thermal proteome profiling data**
**a.** Distribution of the melting temperature (Tm) of Bacteroides thetaiotaomicron VPI-5482 (B. theta) lysate proteome and the whole cell E.coli proteome. **b.** Ranked protein abundance plot of all identified proteins in the 1%DMSO treated controls, S1 sulfatase are highlighted. **c.** Relative abundance of 11 S1 sulfatases identified by mass spectrometry for B. theta cultured in the presence of chondroitin sulfate A (n=6). (b) Melting curve profiles for all identified B. theta proteins in control (1%DMSO) treated lysate. The thermal stability profiles for each individual S1 sulfatases are highlighted in blue. **d.** Melting curve profiles for all identified B. theta proteins in control (1%DMSO) treated lysate. The thermal stability profiles for each individual S1 sulfatases are highlighted in blue.

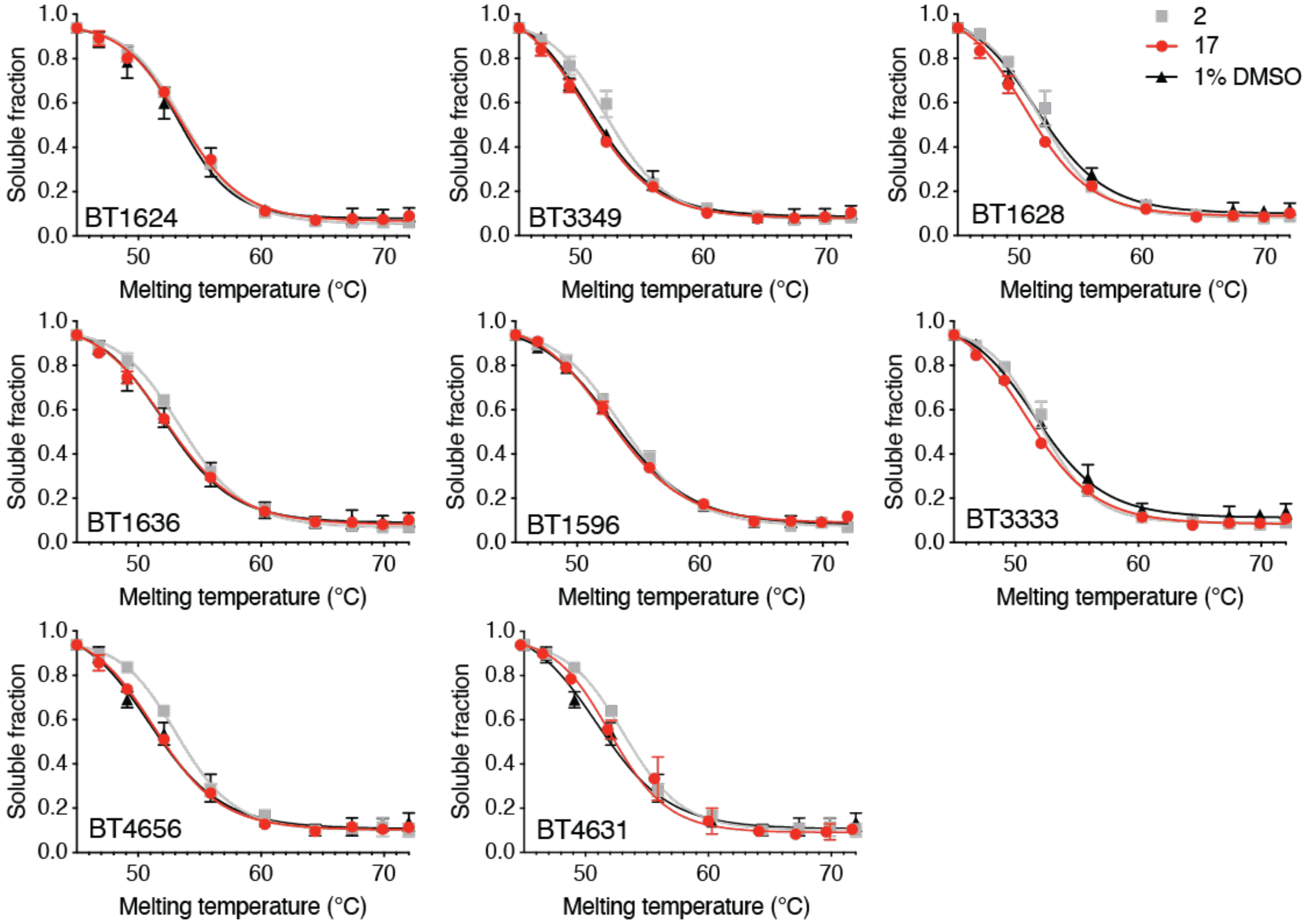


**Supplementary figure 41. Global protein thermal stability analysis confirms that arysulfamate inhibitors**
2 and 17 do not directly interact with S1 sulfatases. Individual melt curve for selected S1 sulfatase for control and arysulfamate inhibitors 2 and 17 treated lysates (n=2 per treatment group).

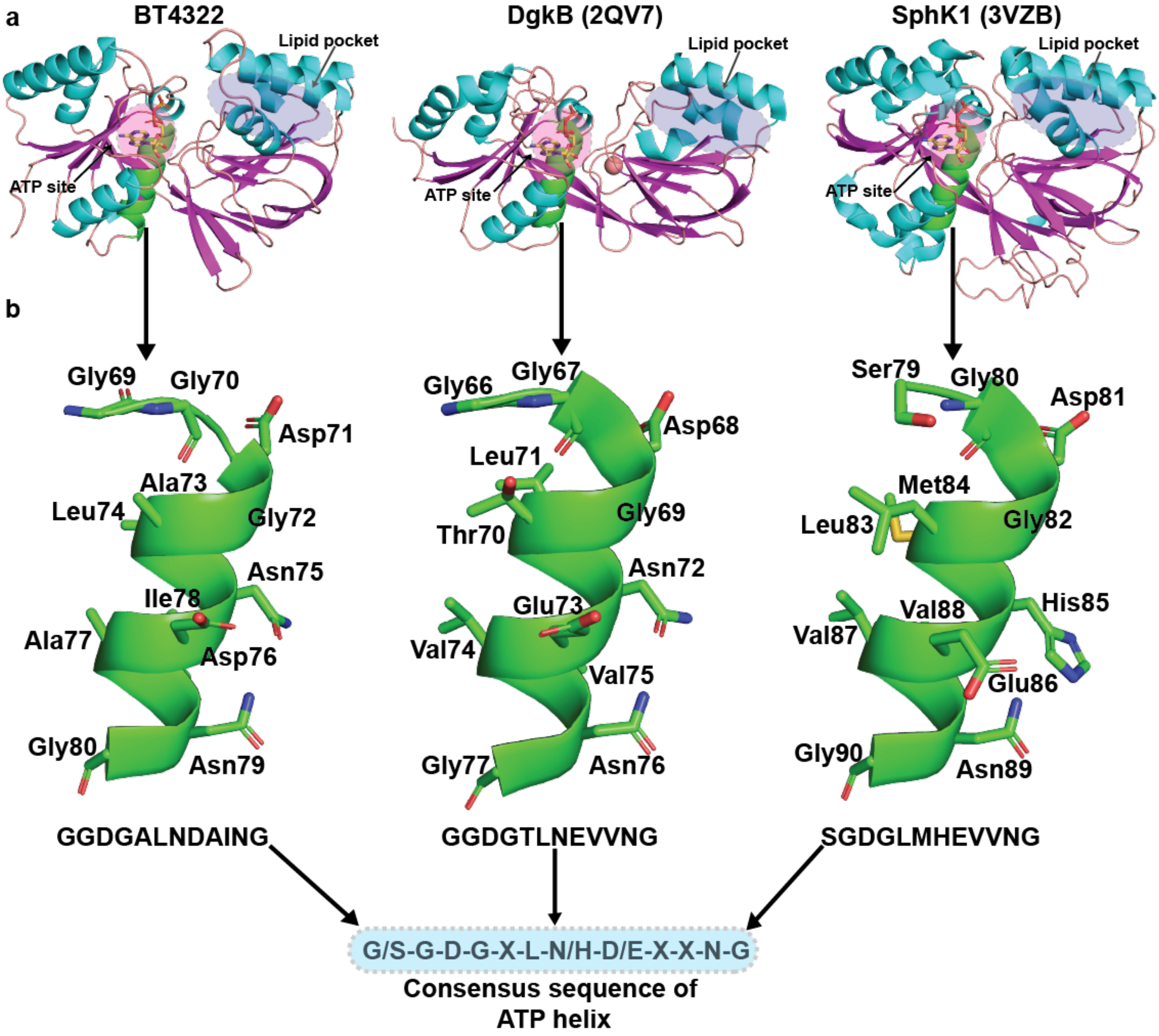


**Supplementary figure 42. Structure of BT4322, SaDgKB, and SphK1 and comparison of the ATP helix**

**a.** Tertiary structures of BT4322, SaDgKB, and SphK1 shown as with a helices, b sheets, and loops coloured cyan, magenta, and pink. The ‘ATP helix’ is coloured in green. **b.** An expanded view of the ATP helix and its sequence.

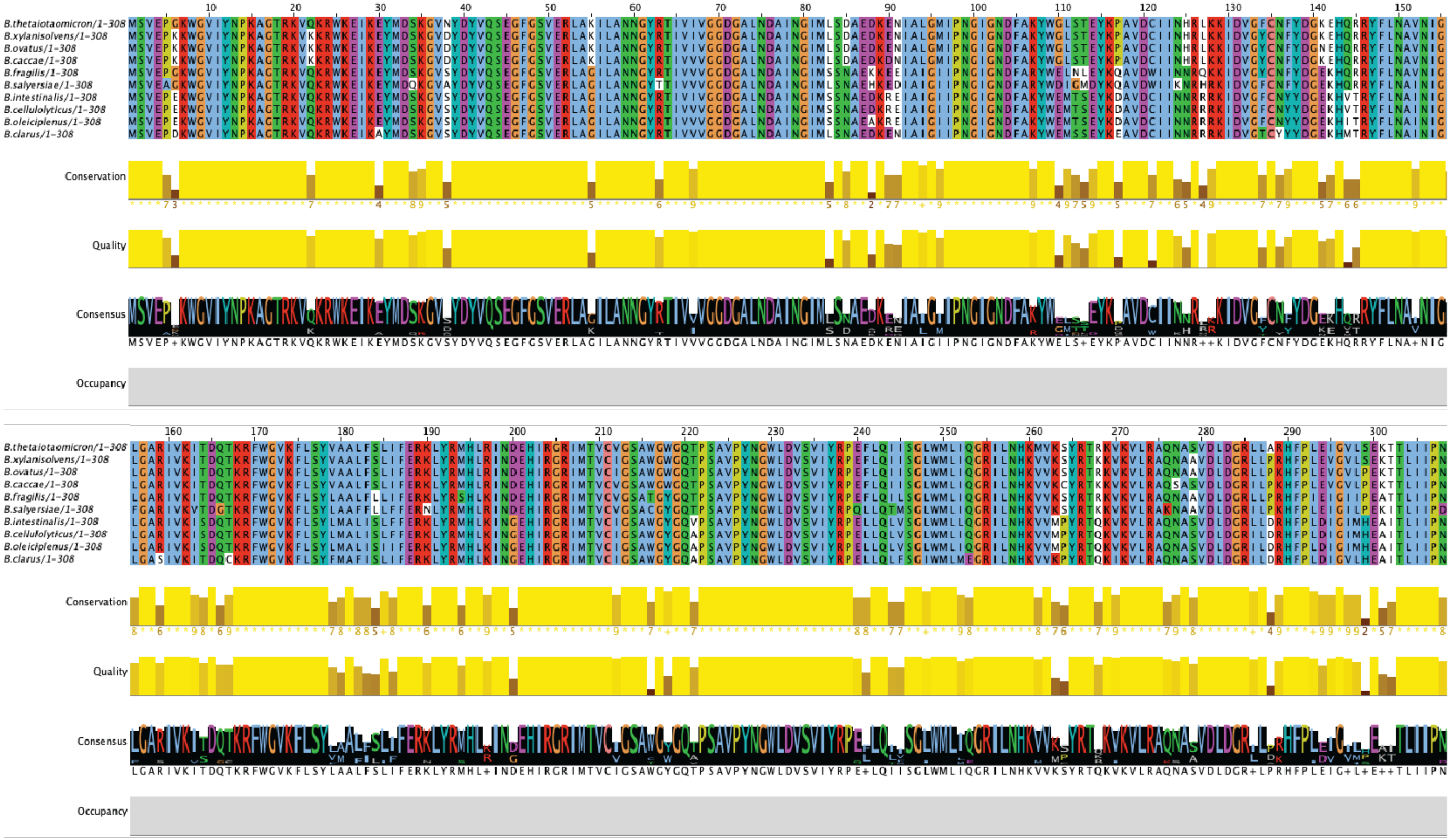

**Supplementary figure 43. Sequence alignment of putative DAGKs from select Bacteroides species.**
Sequences were aligned using the MAFFT online server and visualised using Jalview.

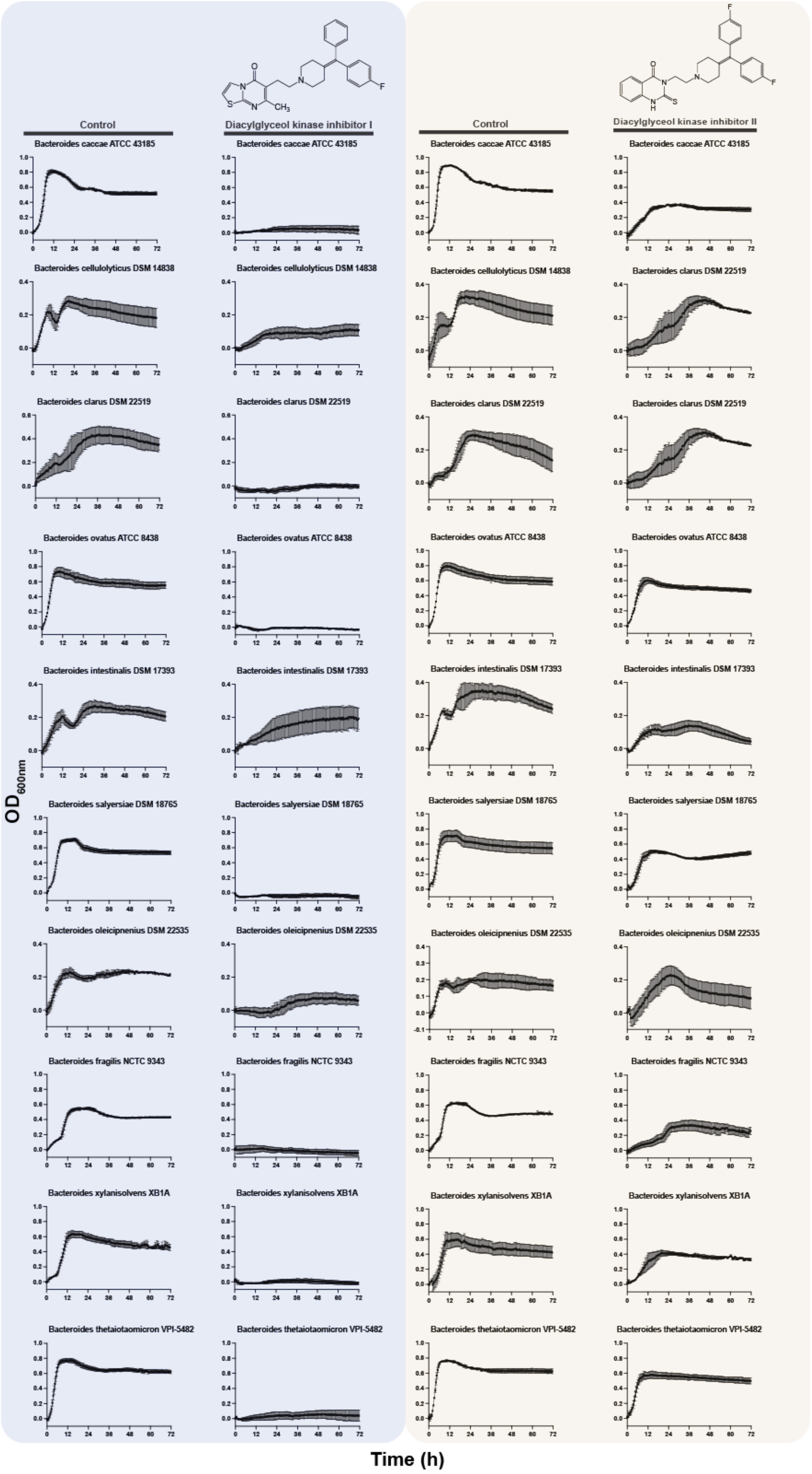

Control
Diacylglyceol kinase inhibitor I
Control
Diacylglyceol kinase inhibitor II
Bacteroides caccae ATCC 43185
Bacteroides cellulolyticus DSM 14838
Bacteroides clarus DSM 22519
Bacteroides ovatus ATCC 8438
Bacteroides intestinalis DSM 17393
Bacteroides salyersiae DSM 18765
Bacteroides oleicipnenius DSM 22535
Bacteroides fragilis NCTC 9343
Bacteroides xylanisolvens XB1A
Bacteroides thetaiotaomicron VPI-5482
$OD_{600nm}$
Time (h)

**Supplementary figure44. The effects of diacylglycerol kinase inhibitors on the growth of HGM Bacteroidota species.**
Bacteroidota species were grown in BHI, 1 % DMSO, with 0.125 mM of diacylglycerol kinase inhibitor I (DAGKI-i) and 0.0625 mM diacylglycerol kinase inhibitor II (DAGKI-ii) and the effects on growth observed. Data are technical triplicates with the standard error of the mean.

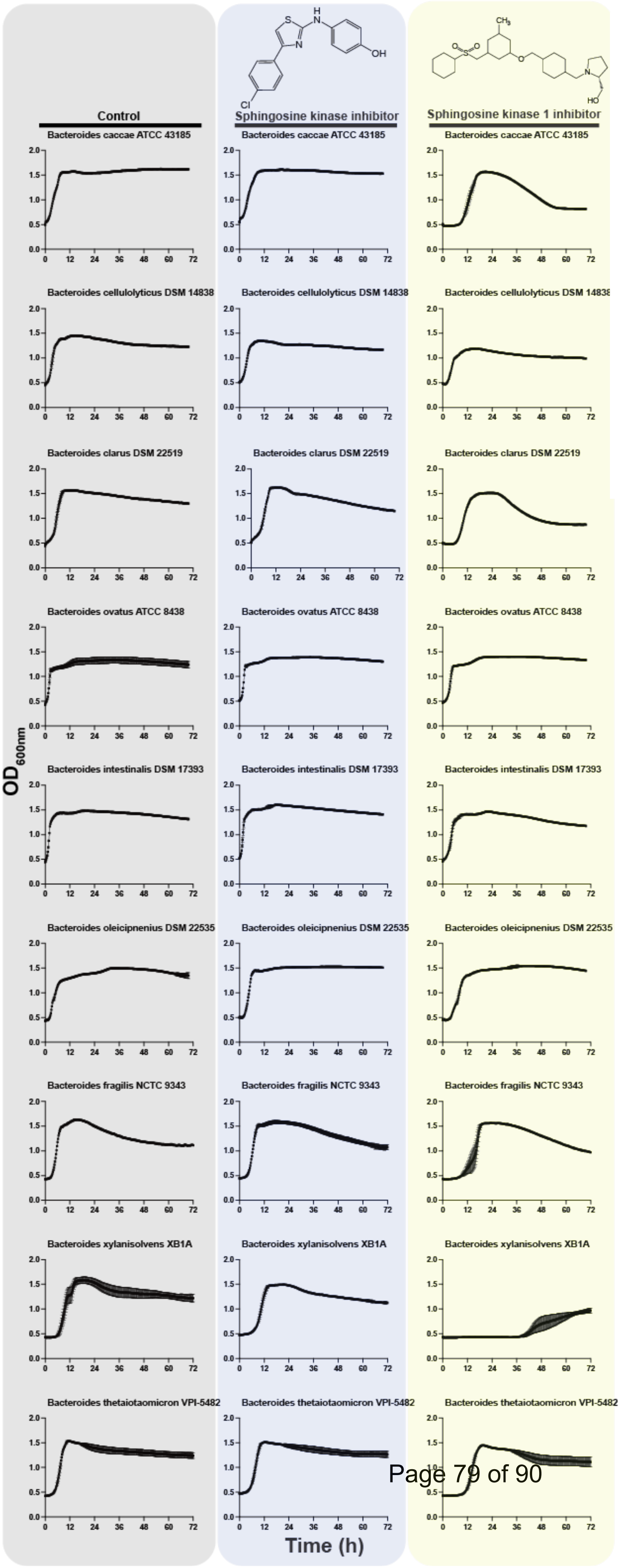


**Supplementary figure 45. The effects of sphigosine kinase inhibitors on the growth of HGM Bacteroidota species**
Bacteroidota species were grown in BHI, 1 % DMSO, with 0.1 mM of sphingosine kinase inhibitor (SKI) and 0.5 mM sphingosine kinase 1 inhibitor II and the effects on growth observed. Data are technical triplicates with the standard error of the mean.

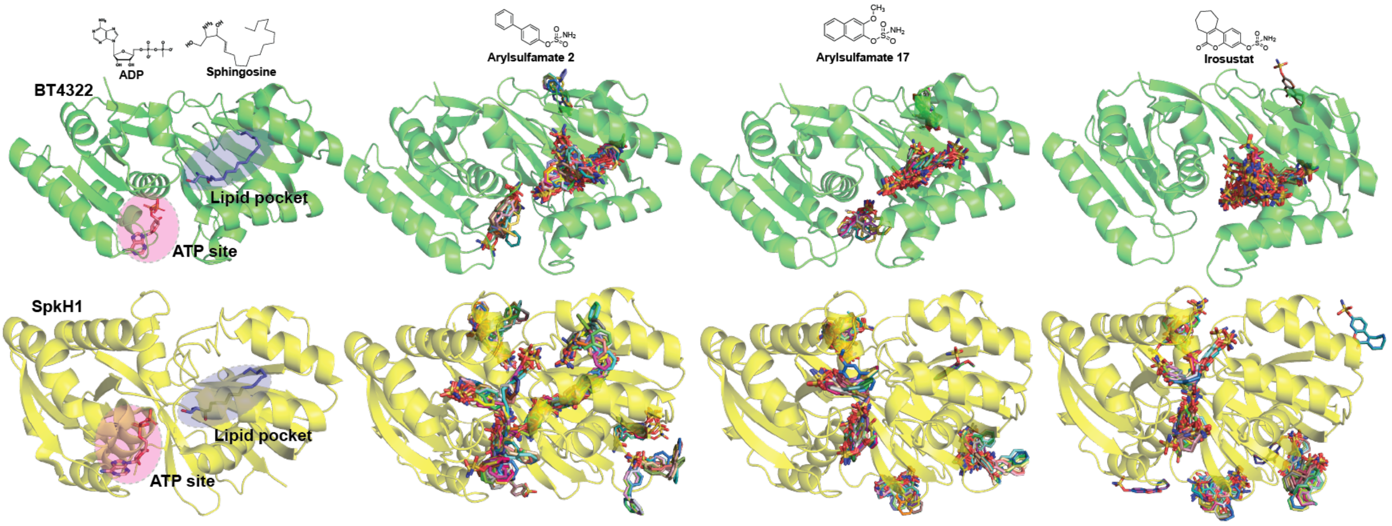


**Supplementary figure 46. Blind docking of arylsulfamates onto BT4322 and SpkH1.**
The results of blind docking (no site specified) arylsulfamate inhibitors into the alphafold2 model of BT4322 (Q89ZQ4) and the crystal structure of SpKH1 (3VZB). It can be observed that all arylsulfamates preferentially cluster in, or near, the lipid binding pocket. For SpKH1 there appears to be no single preferred clustering site.

| Compound | BHI | | | Glucose | | | Chondroitin sulfate C | | |
|---|---|---|---|---|---|---|---|---|---|
| | *Lag phase* | *Growth rate* | *Max OD* | *Lag phase* | *Growth rate* | *Max OD* | *Lag phase* | *Growth rate* | *Max OD* |
| **1** | 0.6139 | 0.0364 | 0.0444 | 0.7619 | 0.0098 | 0.0641 | 0.1456 | 0.0066 | 0.0255 |
| **2** | ND | ND | ND | ND | ND | ND | ND | ND | ND |
| **3** | 0.1482 | 0.0384 | 0.0503 | 0.8820 | 0.0890 | 0.2863 | 0.5727 | 0.0830 | 0.4548 |
| **4** | 0.1326 | 0.3574 | 0.6895 | 0.8541 | 0.2984 | 0.2756 | 0.4276 | 0.2872 | 0.4695 |
| **5** | 0.0803 | 0.0249 | 0.0012 | 0.0993 | 0.0005 | 0.0061 | 0.0616 | 0.0115 | 0.0117 |
| **6** | 0.5497 | 0.7241 | 0.2979 | 0.0651 | 0.2298 | 0.1024 | 0.2239 | 0.1954 | 0.9682 |
| **7** | 0.0245 | 0.0312 | 0.0174 | 0.2573 | 0.0366 | 0.0681 | 0.4966 | 0.0104 | 0.1935 |
| **8** | 0.0512 | 0.0028 | 0.0101 | 0.1649 | 0.0013 | 0.0033 | 0.4682 | 0.0339 | 0.0639 |
| **9** | 0.0947 | 0.0086 | 0.0109 | 0.2181 | 0.0181 | 0.0306 | 0.2210 | 0.0266 | 0.0037 |
| **10** | ND | ND | ND | ND | ND | ND | ND | ND | ND |
| **11** | ND | ND | ND | ND | ND | ND | ND | ND | ND |
| **12** | ND | ND | ND | ND | ND | ND | ND | ND | ND |
| **13** | ND | ND | 0.0020 | 0.0023 | 0.0001 | 0.0001 | ND | ND | ND |
| **14** | ND | ND | ND | 0.0167 | 0.0001 | 0.0001 | ND | ND | ND |
| **15** | ND | ND | ND | ND | ND | ND | ND | ND | ND |
| **16** | ND | ND | ND | ND | ND | ND | ND | ND | ND |
| **17** | 0.1466 | 0.0009 | 0.0001 | ND | ND | ND | ND | ND | ND |
| **18** | 0.7088 | 0.0081 | 0.0010 | ND | ND | ND | ND | ND | ND |

**Supplementary table 1. Significant tests for brain heart infusion, glucose and chondroitin sulfate C growths.**
For growth curves that displayed the classical features of a growth curve, and were measurable, a two-tailed unpaired t-test was performed to check for significance at a threshold of $p<0.5$. Green indicates a significant difference at the chosen threshold, orange no significant difference, and red indicates an analysis was not done due to the lack of a growth curve.

| Compound | Heparin | | | Potato galactan | | | Larch arabinogalactan | | |
|---|---|---|---|---|---|---|---|---|---|
| | *Lag phase* | *Growth rate* | *Max OD* | *Lag phase* | *Growth rate* | *Max OD* | *Lag phase* | *Growth rate* | *Max OD* |
| **1** | 0.0062 | 0.7343 | 0.4803 | 0.4383 | 0.0715 | 0.0395 | 0.0100 | 0.9391 | 0.4074 |
| **2** | ND | ND | ND | ND | ND | ND | ND | ND | ND |
| **3** | ND | ND | ND | 0.0173 | 0.7817 | 0.3000 | 0.0010 | 0.6453 | 0.2329 |
| **4** | ND | ND | ND | 0.0205 | 0.8193 | 0.2273 | 0.6737 | 0.1880 | 0.1510 |
| **5** | ND | ND | ND | 0.4428 | 0.0541 | 0.0518 | ND | ND | ND |
| **6** | 0.5380 | 0.1845 | 0.3459 | 0.5185 | 0.5235 | 0.8915 | 0.2234 | 0.5538 | 0.4971 |
| **7** | ND | ND | ND | 0.0148 | 0.0360 | 0.2612 | 0.2024 | 0.0517 | 0.0170 |
| **8** | ND | ND | ND | 0.0205 | 0.0348 | 0.08515 | ND | ND | ND |
| **9** | ND | ND | ND | 0.0759 | 0.2361 | 0.0146 | 0.0139 | 0.1083 | 0.0034 |
| **10** | ND | ND | ND | ND | ND | ND | ND | ND | ND |
| **11** | ND | ND | ND | ND | ND | ND | ND | ND | ND |
| **12** | ND | ND | ND | ND | ND | ND | ND | ND | ND |
| **13** | 0.0034 | 0.0629 | 0.0012 | ND | ND | ND | ND | ND | ND |
| **14** | ND | ND | ND | ND | ND | ND | ND | ND | ND |
| **15** | ND | ND | ND | ND | ND | ND | ND | ND | ND |
| **16** | ND | ND | ND | ND | ND | ND | ND | ND | ND |
| **17** | ND | ND | ND | ND | ND | ND | ND | ND | ND |
| **18** | 0.0023 | 0.3306 | 0.0419 | 0.0512 | 0.0044 | 0.0004 | ND | ND | ND |

**Supplementary table 2. Significant tests for heparin, larch arabinogalactan, and potato galactan growths.**
For growth curves that displayed the classical features of a growth curve, and were measurable, a two-tailed unpaired t-test was performed to check for significance at a threshold of $p<0.5$. Green indicates a significant difference at the chosen threshold, orange no significant difference, and red indicates an analysis was not done due to the lack of a growth curve.

| | PaAsta | HpSulf | BT3177 | BT4656 | Amuc1074 | Amuc1033 | BT1636 | BT1622 | Amuc0451 | Amuc0491 | BT3796 | Amuc1755 |
|---|---|---|---|---|---|---|---|---|---|---|---|---|
| **Control** | (-1.1 ± 0.02) x $10^{-4}$ | (-9.4 ± 0.1) x $10^{-6}$ | (2.7 ± 1.4) x $10^{-3}$ | (5.0 ± 8.8) x $10^{-5}$ | (4.2 ± 1.6) x $10^{-4}$ | (2.1 ± 0.09) x $10^{-4}$ | (3.2 ± 2.0) x $10^{-4}$ | (1.5 ± 2.0) x $10^{-4}$ | (5.4 ± 4.4) x $10^{-5}$ | (1.3 ± 0.3) x $10^{-4}$ | (4.1 ± 1.1) x $10^{-4}$ | (4.1 ± 0.06) x $10^{-4}$ |
| **WT** | (7.0 ± 0.1) x $10^{-3}$ | (5.8 ± 0.1) x $10^{-5}$ | (1.4 ± 0.05) x $10^{-3}$ | (3.3 ± 0.2) x $10^{-3}$ | (4.4 ± 0.2) x $10^{-3}$ | (1.5 ± 0.03) x $10^{-3}$ | (3.0 ± 0.1) x $10^{-3}$ | (7.0 ± 1.1) x $10^{-3}$ | (1.6 ± 0.01) x $10^{-3}$ | (2.5 ± 0.3) x $10^{-4}$ | (4.6 ± 0.2) x $10^{-3}$ | (2.9 ± 0.02) x $10^{-3}$ |
| **1** | NA | NA | (1.6 ± 0.01) x $10^{-3}$ | (3.9 ± 0.1) x $10^{-3}$ | (4.4 ± 0.1) x $10^{-3}$ | (1.5 ± 0.2) x $10^{-3}$ | (3.2 ± 0.2) x $10^{-3}$ | (7.4 ± 1.1) x $10^{-3}$ | (1.7 ± 0.08) x $10^{-3}$ | (2.1 ± 0.4) x $10^{-4}$ | (4.7 ± 0.3) x $10^{-3}$ | (2.5 ± 0.02) x $10^{-3}$ |
| **6** | (9.0 ± 0.01) x $10^{-3}$ | (6.7 ± 0.03) x $10^{-5}$ | (1.3 ± 0.09) x $10^{-3}$ | (3.7 ± 0.06) x $10^{-3}$ | (4.2 ± 0.5) x $10^{-3}$ | (1.6 ± 0.09) x $10^{-3}$ | (3.2 ± 0.2) x $10^{-3}$ | (6.9 ± 1.1) x $10^{-3}$ | (1.7 ± 0.09) x $10^{-3}$ | (2.6 ± 0.4) x $10^{-4}$ | (4.1 ± 0.2) x $10^{-3}$ | (2.9 ± 0.08) x $10^{-3}$ |
| **2** | NA | NA | (1.5 ± 0.02) x $10^{-3}$ | (4.0 ± 0.09) x $10^{-3}$ | (5.5 ± 0.6) x $10^{-3}$ | (1.7 ± 0.07) x $10^{-3}$ | (3.3 ± 0.2) x $10^{-3}$ | (7.5 ± 0.4) x $10^{-3}$ | (1.8 ± 0.07) x $10^{-3}$ | (2.6 ± 0.3) x $10^{-4}$ | (4.6 ± 0.3) x $10^{-3}$ | (2.8 ± 0.08) x $10^{-3}$ |
| **5** | NA | NA | (1.6 ± 0.1) x $10^{-3}$ | (3.8 ± 0.2) x $10^{-3}$ | (4.2 ± 0.3) x $10^{-3}$ | (1.7 ± 0.1) x $10^{-3}$ | (3.2 ± 0.2) x $10^{-3}$ | (6.9 ± 1.4) x $10^{-3}$ | (1.6 ± 0.09) x $10^{-3}$ | (2.5 ± 0.3) x $10^{-4}$ | (4.3 ± 0.3) x $10^{-3}$ | (2.9 ± 0.04) x $10^{-3}$ |
| **7** | (2.6 ± 0.05) x $10^{-3}$ | (5.0 ± 0.04) x $10^{-5}$ | (1.6 ± 0.2) x $10^{-3}$ | (3.8 ± 0.06) x $10^{-3}$ | (4.9 ± 0.6) x $10^{-3}$ | (1.6 ± 0.06) x $10^{-3}$ | (3.2 ± 0.3) x $10^{-3}$ | (7.4 ± 0.1) x $10^{-3}$ | (1.7 ± 0.09) x $10^{-3}$ | (2.2 ± 0.6) x $10^{-4}$ | (4.5 ± 0.1) x $10^{-3}$ | (2.7 ± 0.02) x $10^{-3}$ |
| **8** | (4.9 ± 0.3) x $10^{-3}$ | (4.7 ± 0.06) x $10^{-5}$ | (1.5 ± 0.01) x $10^{-3}$ | (3.8 ± 0.1) x $10^{-3}$ | (4.6 ± 0.3) x $10^{-3}$ | (1.9 ± 0.2) x $10^{-3}$ | (3.1 ± 0.1) x $10^{-3}$ | (7.7 ± 0.4) x $10^{-3}$ | (1.7 ± 0.07) x $10^{-3}$ | (3.0 ± 0.3) x $10^{-4}$ | (3.7 ± 0.2) x $10^{-3}$ | (2.8 ± 0.03) x $10^{-3}$ |
| **9** | NA | NA | (1.4 ± 0.04) x $10^{-3}$ | (3.7 ± 0.08) x $10^{-3}$ | (5.3 ± 0.1) x $10^{-3}$ | (1.8 ± 0.07) x $10^{-3}$ | (4.3 ± 0.9) x $10^{-3}$ | (8.7 ± 0.2) x $10^{-3}$ | (1.6 ± 0.04) x $10^{-3}$ | (2.8 ± 0.4) x $10^{-4}$ | (4.0 ± 0.5) x $10^{-3}$ | (2.7 ± 0.05) x $10^{-3}$ |
| **10** | NA | (0.3 ± 0.04) x $10^{-5}$ | (1.3 ± 0.04) x $10^{-3}$ | (3.1 ± 0.1) x $10^{-3}$ | (5.8 ± 0.2) x $10^{-3}$ | (2.3 ± 0.07) x $10^{-3}$ | (3.5 ± 0.2) x $10^{-3}$ | (8.9 ± 0.2) x $10^{-3}$ | (1.4 ± 0.01) x $10^{-3}$ | (2.5 ± 0.5) x $10^{-4}$ | (4.8 ± 0.3) x $10^{-3}$ | (2.4 ± 0.05) x $10^{-3}$ |
| **13** | NA | NA | (1.3 ± 0.05) x $10^{-3}$ | (4.1 ± 0.1) x $10^{-3}$ | (4.9 ± 0.2) x $10^{-3}$ | (1.9 ± 0.2) x $10^{-3}$ | (4.6 ± 1.0) x $10^{-3}$ | (8.0 ± 1.1) x $10^{-3}$ | (1.7 ± 0.01) x $10^{-3}$ | (2.3 ± 0.5) x $10^{-4}$ | (4.8 ± 0.3) x $10^{-3}$ | (3.1 ± 0.06) x $10^{-3}$ |
| **14** | NA | NA | (1.4 ± 0.05) x $10^{-3}$ | (4.0 ± 0.07) x $10^{-3}$ | (4.7 ± 0.07) x $10^{-3}$ | (1.4 ± 0.1) x $10^{-3}$ | (4.7 ± 0.6) x $10^{-3}$ | (8.4 ± 0.2) x $10^{-3}$ | (1.6 ± 0.05) x $10^{-3}$ | (2.7 ± 0.4) x $10^{-4}$ | (4.8 ± 0.2) x $10^{-3}$ | (2.9 ± 0.07) x $10^{-3}$ |
| **11** | NA | NA | (1.2 ± 0.01) x $10^{-3}$ | (4.0 ± 0.09) x $10^{-3}$ | (6.3 ± 1.0) x $10^{-3}$ | (2.0± 0.3) x $10^{-3}$ | (3.3 ± 0.2) x $10^{-3}$ | (7.3 ± 1.3) x $10^{-3}$ | (1.4 ± 0.05) x $10^{-3}$ | (2.6 ± 0.3) x $10^{-4}$ | (4.8 ± 0.2) x $10^{-3}$ | (2.6 ± 0.05) x $10^{-3}$ |
| **17** | NA | NA | (0.98 ± 0.04) x $10^{-3}$ | (4.4 ± 0.2) x $10^{-3}$ | (5.2 ± 0.6) x $10^{-3}$ | (2.1 ± 0.2) x $10^{-3}$ | (3.7 ± 0.7) x $10^{-3}$ | (9.0 ± 0.3) x $10^{-3}$ | (1.7 ± 0.05) x $10^{-3}$ | (2.6 ± 0.3) x $10^{-4}$ | (5.1 ± 0.2) x $10^{-3}$ | (3.5 ± 0.06) x $10^{-3}$ |
| **15** | NA | NA | NT | NT | NT | NT | NT | NT | (1.9 ± 0.05) x $10^{-3}$ | (2.2 ± 0.5) x $10^{-4}$ | (5.8 ± 0.4) x $10^{-3}$ | (2.9 ± 0.02) x $10^{-3}$ |
| **12** | NT | (2.4 ± 0.03) x $10^{-5}$ | NT | NT | NT | NT | NT | NT | NT | NT | NT | NT |
| **18** | NA | NA | (1.3 ± 0.02) x $10^{-3}$ | (4.3 ± 0.2) x $10^{-3}$ | (4.8 ± 0.3) x $10^{-3}$ | (1.6 ± 0.1) x $10^{-3}$ | (2.9 ± 0.2) x $10^{-3}$ | (9.4 ± 0.5) x $10^{-3}$ | (1.6 ± 0.03) x $10^{-3}$ | (2.8 ± 0.3) x $10^{-4}$ | (5.0 ± 0.2) x $10^{-3}$ | (2.7 ± 0.04) x $10^{-3}$ |
| **16** | NA | NA | (1.4 ± 0.03) x $10^{-3}$ | (4.5 ± 0.2) x $10^{-3}$ | (4.8 ± 0.4) x $10^{-3}$ | (1.7 ± 0.1) x $10^{-3}$ | (3.0 ± 0.4) x $10^{-3}$ | (9.2 ± 0.4) x $10^{-3}$ | (1.6 ± 0.06) x $10^{-3}$ | (2.2 ± 0.08) x $10^{-4}$ | (5.0 ± 0.4) x $10^{-3}$ | (4.6 ± 0.06) x $10^{-3}$ |
| **3** | *(3.8 ± 0.8) x $10^{-3}$ | NA | ( 1.2 ± 0.02) x $10^{-3}$ | (4.1 ± 0.1) x $10^{-3}$ | (4.8 ± 0.4) x $10^{-3}$ | (1.5 ± 0.1) x $10^{-3}$ | (3.2 ± 0.2) x $10^{-3}$ | (7.6 ± 0.1) x $10^{-3}$ | (1.6 ± 0.06) x $10^{-3}$ | (1.9± 0.07) x $10^{-4}$ | (4.6 ± 0.4) x $10^{-3}$ | (2.8 ± 0.3) x $10^{-3}$ |
| **4** | *(5.8 ± 0.4) x $10^{-3}$ | NA | (1.3 ± 0.03) x $10^{-3}$ | (4.0 ± 0.2) x $10^{-3}$ | (5.1 ± 0.3) x $10^{-3}$ | (1.5 ± 0.1) x $10^{-3}$ | (3.0 ± 0.1) x $10^{-3}$ | (8.2 ± 0.4) x $10^{-3}$ | (1.6 ± 0.04) x $10^{-3}$ | (2.1 ± 0.09) x $10^{-4}$ | (4.4 ± 0.3) x $10^{-3}$ | (2.9 ± 0.2) x $10^{-3}$ |

**Supplementary table 3. Kinetic rates of sulfatase when arylsulfamates were included in the assay conditions.**
The concentration of arylsulfamate was always 1 mM. For the steroid sulfatases the substrate *para*-nitrophenol sulfate was at a concentration of 1 mM, whilst for carbohydrate sulfatases substrate concentration was 1 μM. NA means no activity when compared to the control. NT means not tested. Numbers in the left most column refer to the arylsulfamate in Figure 1c added to the reaction; control represents substrate only whilst WT is substrate plus enzyme without an arylsulfamate inhibitor. Rates are μM product/min.

| | PaAsta | HpSulf | BT3177 | BT4656 | Amuc1074 | Amuc1033 | BT1636 | BT1622 | Amuc0451 | Amuc0491 | BT3796 | Amuc1755 |
|---|---|---|---|---|---|---|---|---|---|---|---|---|
| **Control** | $(-6.57 \pm 1.53) \times 10^{-5}$ | $(-7.9 \pm 2.7) \times 10^{-6}$ | $(1.9 \pm 1.4) \times 10^{-4}$ | $(2.3 \pm 1.9) \times 10^{-3}$ | $(2.8 \pm 0.2) \times 10^{-4}$ | $(0.6 \pm 1.0) \times 10^{-4}$ | $(8.8 \pm 2.7) \times 10^{-5}$ | $(1.2 \pm 1.7) \times 10^{-4}$ | $(7.6 \pm 3.2) \times 10^{-5}$ | $(7.6 \pm 2.5) \times 10^{-5}$ | $(5.3 \pm 0.5) \times 10^{-4}$ | $(2.9 \pm 0.6) \times 10^{-4}$ |
| **WT** | $(7.5 \pm 0.2) \times 10^{-3}$ | $(7.5 \pm 0.2) \times 10^{-4}$ | $(2.4 \pm 0.1) \times 10^{-3}$ | $(3.3 \pm 0.05) \times 10^{-3}$ | $(4.3 \pm 0.1) \times 10^{-3}$ | $(2.2 \pm 0.1) \times 10^{-3}$ | $(1.7 \pm 0.05) \times 10^{-3}$ | $(6.2 \pm 0.2) \times 10^{-3}$ | $(4.0 \pm 0.05) \times 10^{-3}$ | $(2.0 \pm 0.2) \times 10^{-4}$ | $(4.9 \pm 0.2) \times 10^{-3}$ | $(2.7 \pm 0.09) \times 10^{-3}$ |
| **1** | NA | NA | $(2.6 \pm 0.1) \times 10^{-3}$ | $(4.0 \pm 0.1) \times 10^{-3}$ | $(4.4 \pm 0.09) \times 10^{-3}$ | $(2.0 \pm 0.05) \times 10^{-3}$ | $(1.8 \pm 0.05) \times 10^{-3}$ | $(5.8 \pm 0.3) \times 10^{-3}$ | $(3.0 \pm 0.06) \times 10^{-3}$ | $(2.2 \pm 0.08) \times 10^{-4}$ | $(5.1 \pm 0.2) \times 10^{-3}$ | $(5.6 \pm 0.2) \times 10^{-3}$ |
| **6** | $(5.9 \pm 0.1) \times 10^{-3}$ | $(6.4 \pm 0.1) \times 10^{-4}$ | $(2.7 \pm 0.09) \times 10^{-3}$ | $(4.1 \pm 0.2) \times 10^{-3}$ | $(4.5 \pm 0.1) \times 10^{-3}$ | $(2.1 \pm 0.07) \times 10^{-3}$ | $(2.0 \pm 0.01) \times 10^{-3}$ | $(6.0 \pm 0.2) \times 10^{-3}$ | $(3.9 \pm 0.06) \times 10^{-3}$ | $(2.1 \pm 0.2) \times 10^{-4}$ | $(4.8 \pm 0.2) \times 10^{-3}$ | $(2.9 \pm 0.06) \times 10^{-3}$ |
| **2** | NA | NA | $(2.7 \pm 0.2) \times 10^{-3}$ | $(4.0 \pm 0.06) \times 10^{-3}$ | $(4.2 \pm 0.09) \times 10^{-3}$ | $(2.2 \pm 0.1) \times 10^{-3}$ | $(1.8 \pm 0.04) \times 10^{-3}$ | $(5.9 \pm 0.2) \times 10^{-3}$ | $(3.5 \pm 0.1) \times 10^{-3}$ | $(1.9 \pm 0.3) \times 10^{-4}$ | $(4.2 \pm 0.2) \times 10^{-3}$ | $(3.7 \pm 0.1) \times 10^{-3}$ |
| **5** | NA | NA | $(2.4 \pm 0.1) \times 10^{-3}$ | $(4.2 \pm 0.06) \times 10^{-3}$ | $(4.2 \pm 0.09) \times 10^{-3}$ | $(2.1 \pm 0.09) \times 10^{-3}$ | $(2.8 \pm 0.02) \times 10^{-3}$ | $(5.5 \pm 0.3) \times 10^{-3}$ | $(3.8 \pm 0.05) \times 10^{-3}$ | $(2.3 \pm 0.2) \times 10^{-4}$ | $(5.2 \pm 0.2) \times 10^{-3}$ | $(6.4 \pm 0.2) \times 10^{-3}$ |
| **7** | $(5.7 \pm 0.1) \times 10^{-3}$ | $(7.3 \pm 0.7) \times 10^{-4}$ | $(2.6 \pm 0.09) \times 10^{-3}$ | $(3.9 \pm 0.3) \times 10^{-3}$ | $(4.4 \pm 0.09) \times 10^{-3}$ | $(2.1 \pm 0.05) \times 10^{-3}$ | $(2.0 \pm 0.02) \times 10^{-3}$ | $(6.9 \pm 0.3) \times 10^{-3}$ | $(4.0 \pm 0.05) \times 10^{-3}$ | $(2.0 \pm 0.08) \times 10^{-4}$ | $(5.1 \pm 0.2) \times 10^{-3}$ | $(2.8 \pm 0.09) \times 10^{-3}$ |
| **8** | $(6.3 \pm 0.7) \times 10^{-3}$ | $(5.4 \pm 0.5) \times 10^{-4}$ | $(2.5 \pm 0.1) \times 10^{-3}$ | $(4.1 \pm 0.1) \times 10^{-3}$ | $(4.4 \pm 0.1) \times 10^{-3}$ | $(2.0 \pm 0.07) \times 10^{-3}$ | $(2.6 \pm 0.08) \times 10^{-3}$ | $(6.1 \pm 0.3) \times 10^{-3}$ | $(3.8 \pm 0.06) \times 10^{-3}$ | $(2.1 \pm 0.2) \times 10^{-4}$ | $(5.4 \pm 0.2) \times 10^{-3}$ | $(5.2 \pm 0.2) \times 10^{-3}$ |
| **9** | NA | NA | $(2.6 \pm 0.2) \times 10^{-3}$ | $(4.1 \pm 0.3) \times 10^{-3}$ | $(4.3 \pm 0.1) \times 10^{-3}$ | $(2.0 \pm 0.1) \times 10^{-3}$ | $(2.3 \pm 0.02) \times 10^{-3}$ | $(6.2 \pm 0.2) \times 10^{-3}$ | $(3.3 \pm 0.03) \times 10^{-3}$ | $(2.5 \pm 0.3) \times 10^{-4}$ | $(5.2 \pm 0.2) \times 10^{-3}$ | $(5.7 \pm 0.3) \times 10^{-3}$ |
| **10** | NA | NA | $(2.7 \pm 0.2) \times 10^{-3}$ | $(4.1 \pm 0.08) \times 10^{-3}$ | $(4.3 \pm 0.1) \times 10^{-3}$ | $(2.2 \pm 0.1) \times 10^{-3}$ | $(1.7 \pm 0.1) \times 10^{-3}$ | $(6.6 \pm 0.1) \times 10^{-3}$ | $(3.7 \pm 0.04) \times 10^{-3}$ | $(1.8 \pm 0.1) \times 10^{-4}$ | $(5.1 \pm 0.2) \times 10^{-3}$ | $(5.0 \pm 0.2) \times 10^{-3}$ |
| **13** | NA | NA | $(2.7 \pm 0.1) \times 10^{-3}$ | $(4.3 \pm 0.06) \times 10^{-3}$ | $(4.4 \pm 0.1) \times 10^{-3}$ | $(2.2 \pm 0.06) \times 10^{-3}$ | $(2.3 \pm 0.06) \times 10^{-3}$ | $(6.6 \pm 0.2) \times 10^{-3}$ | $(3.9 \pm 0.07) \times 10^{-3}$ | $(1.9 \pm 0.1) \times 10^{-4}$ | $(5.6 \pm 0.2) \times 10^{-3}$ | $(3.6 \pm 0.1) \times 10^{-3}$ |
| **14** | NA | NA | $(2.4 \pm 0.2) \times 10^{-3}$ | $(4.0 \pm 0.2) \times 10^{-3}$ | $(4.4 \pm 0.1) \times 10^{-3}$ | $(2.1 \pm 0.08) \times 10^{-3}$ | $(1.9 \pm 0.01) \times 10^{-3}$ | $(6.7 \pm 0.2) \times 10^{-3}$ | $(3.2 \pm 0.3) \times 10^{-3}$ | $(1.9 \pm 0.2) \times 10^{-4}$ | $(5.6 \pm 0.2) \times 10^{-3}$ | $(3.3 \pm 0.1) \times 10^{-3}$ |
| **11** | NA | NA | $(2.6 \pm 0.1) \times 10^{-3}$ | $(4.3 \pm 0.1) \times 10^{-3}$ | $(4.7 \pm 0.09) \times 10^{-3}$ | $(2.1 \pm 0.1) \times 10^{-3}$ | $(1.8 \pm 0.03) \times 10^{-3}$ | $(6.6 \pm 0.3) \times 10^{-3}$ | $(2.3 \pm 0.5) \times 10^{-3}$ | $(2.0 \pm 0.1) \times 10^{-4}$ | $(5.4 \pm 0.2) \times 10^{-3}$ | $(3.8 \pm 0.2) \times 10^{-3}$ |
| **17** | NA | NA | $(2.5 \pm 0.2) \times 10^{-3}$ | $(4.3 \pm 0.1) \times 10^{-3}$ | $(4.6 \pm 0.1) \times 10^{-3}$ | $(2.0 \pm 0.1) \times 10^{-3}$ | $(1.9 \pm 0.05) \times 10^{-3}$ | $(6.4 \pm 0.2) \times 10^{-3}$ | $(1.7 \pm 0.04) \times 10^{-3}$ | $(2.0 \pm 0.1) \times 10^{-4}$ | $(5.8 \pm 0.3) \times 10^{-3}$ | $(4.8 \pm 0.09) \times 10^{-3}$ |
| **15** | NA | NA | $(2.6 \pm 0.08) \times 10^{-3}$ | $(4.5 \pm 0.2) \times 10^{-3}$ | $(4.8 \pm 0.1) \times 10^{-3}$ | $(2.2 \pm 0.03) \times 10^{-3}$ | $(1.7 \pm 0.04) \times 10^{-3}$ | $(5.6 \pm 0.3) \times 10^{-3}$ | $(3.4 \pm 0.03) \times 10^{-3}$ | $(2.0 \pm 0.1) \times 10^{-4}$ | $(5.5 \pm 0.2) \times 10^{-3}$ | $(3.3 \pm 0.1) \times 10^{-3}$ |
| **12** | NT | NA | $(2.6 \pm 0.1) \times 10^{-3}$ | $(4.4 \pm 0.1) \times 10^{-3}$ | $(4.8 \pm 0.1) \times 10^{-3}$ | $(2.1 \pm 0.06) \times 10^{-3}$ | $(1.9 \pm 0.07) \times 10^{-3}$ | $(6.5 \pm 0.2) \times 10^{-3}$ | $(3.5 \pm 0.06) \times 10^{-3}$ | $(2.0 \pm 0.1) \times 10^{-4}$ | $(5.6 \pm 0.2) \times 10^{-3}$ | $(3.6 \pm 0.07) \times 10^{-3}$ |
| **18** | NA | NA | $(2.6 \pm 0.1) \times 10^{-3}$ | $(4.4 \pm 0.2) \times 10^{-3}$ | $(4.6 \pm 0.1) \times 10^{-3}$ | $(2.2 \pm 0.05) \times 10^{-3}$ | $(2.3 \pm 0.02) \times 10^{-3}$ | $(6.6 \pm 0.3) \times 10^{-3}$ | NT | NT | NT | NT |
| **16** | NA | NA | $(2.6 \pm 0.1) \times 10^{-3}$ | $(4.2 \pm 0.1) \times 10^{-3}$ | $(4.6 \pm 0.1) \times 10^{-3}$ | $(2.2 \pm 0.06) \times 10^{-3}$ | $(2.3 \pm 0.02) \times 10^{-3}$ | $(6.8 \pm 0.2) \times 10^{-3}$ | $(3.0 \pm 0.02) \times 10^{-3}$ | $(2.2 \pm 0.1) \times 10^{-4}$ | $(6.4 \pm 0.2) \times 10^{-3}$ | $(4.3 \pm 0.09) \times 10^{-3}$ |
| **3** | NA | NA | $(2.4 \pm 0.1) \times 10^{-3}$ | $(4.2 \pm 0.1) \times 10^{-3}$ | $(4.6 \pm 0.1) \times 10^{-3}$ | $(2.2 \pm 0.06) \times 10^{-3}$ | $(1.9 \pm 0.02) \times 10^{-3}$ | $(5.3 \pm 0.1) \times 10^{-3}$ | $(2.6 \pm 0.08) \times 10^{-3}$ | $(2.1 \pm 0.1) \times 10^{-4}$ | $(5.7 \pm 0.2) \times 10^{-3}$ | $(2.7 \pm 0.09) \times 10^{-3}$ |
| **4** | NA | NA | $(2.2 \pm 0.1) \times 10^{-3}$ | $(4.3 \pm 0.1) \times 10^{-3}$ | $(4.7 \pm 0.09) \times 10^{-3}$ | $(2.2 \pm 0.05) \times 10^{-3}$ | $(2.0 \pm 0.05) \times 10^{-3}$ | $(6.3 \pm 0.2) \times 10^{-3}$ | $(3.2 \pm 0.06) \times 10^{-3}$ | $(2.1 \pm 0.1) \times 10^{-4}$ | $(6.2 \pm 0.2) \times 10^{-3}$ | $(4.3 \pm 0.1) \times 10^{-3}$ |

**Supplementary table 4. Kinetic rates of sulfatase when sulfatases were pre-incubated with arylsulfamates then assayed.**
The concentration of arylsulfamate used for preincubation was always 1 mM and the enzyme concentration 25 - 50 μM. For the steroid sulfatases the substrate *para*-nitrophenol sulfate was at a concentration of 1 mM, whilst for carbohydrate sulfatases substrate concentration was 1 μM. NA means no activity when compared to the control. NT means not tested. Numbers in the left most column refer to the arylsulfamate in Figure 1c added to the reaction; control represents substrate only whilst WT is substrate plus enzyme without an arylsulfamate inhibitor. Rates are μM product/min.

| | BT1636 | |
|---|---|---|
| | **In assay** | **Pre-incubated** |
| **Control** | (2.7 ± 0.2) x $10^{-2}$ | (3.1 ± 2.3) x$10^{-2}$ |
| **WT** | 0.58 ± 0.07 | 1.4 ± 0.06 |
| **22** | 0.68 ± 0.02 | NT |
| **23** | NT | 0.49 ± 0.01 |
| **19** | 0.72 ± 0.09 | 1.3 ± 0.08 |
| **24** | 0.68 ± 0.03 | 0.60± 0.01 |
| **WT** | - | 1.2 ± 0.06 |
| **20** | 0.63 ± 0.03 | 1.2 ± 0.08 |
| **21** | 0.58 ± 0.03 | 1.2 ± 0.04 |
| **25** | 0.56 ± 0.02 | 1.2 ± 0.06 |

**Supplementary table 5. Kinetic rate of BT1636$^{3S\text{-}Gal}$ assayed against BODIPY labelled 3S-galactose sulfate with various carbohydrate and aryl- sulfonates/sulfamates.**
BT1636$^{3S\text{-}Gal}$ was assayed against 1 μM BODIPY labelled 3S-galactose with 1 mM of **with** various carbohydrate and aryl- sulfonates/sulfamates the assay, or after BT1636$^{3S\text{-}Gal}$ had been incubated with 1 mM of the same compounds; concentrations of 300 nM and 285 nM of BT1636$^{3S\text{-}Gal}$ were used, respectively. The assay was performed in 100 mM of MES pH 6.0 with 5 % DMSO, 150 mM NaCl, 0.02% (v/v) Brij-35 and 5 mM $CaCl_2$. Assays were performed in triplicate. Rates are fluorescence of product/min. The values in green indicate 5 % was not present.

| | 0.00 mM | 0.01 mM | 0.10 mM | 1.00 mM |
|---|---|---|---|---|
| **3S-Gal** | 54.6 ± 0.06 | 54.7 ± 0.03 | 55.6 ± 0.04 | 57.6 ± 0.04 |
| **20** | 54.6 ± 0.06 | 54.6 ± 0.03 | 54.6 ± 0.02 | 54.4 ± 0.03 |
| **21** | 54.6 ± 0.06 | 54.6 ± 0.07 | 54.5 ± 0.02 | 54.4 ± 0.01 |
| **25** | 54.6 ± 0.06 | 54.5 ± 0.01 | 54.6 ± 0.07 | 54.1 ± 0.07 |
| **19** | 53.4 ± 0.05 | 52.0 ± 0.03 | 51.4 ± 0.11 | 49.9 ± 0.12 |
| **24** | 53.4 ± 0.05 | 52.9 ± 0.05 | 52.7 ± 0.04 | 52.0 ± 0.07 |

**Supplementary table 6. Thermal melt values of BT1636$^{3S\text{-}Gal}$ in the absence and presence of various carbohydrate and aryl- sulfonates/sulfamates.**
BT1636$^{3S\text{-}Gal}$, at a concentration of 5 μM, was incubated with no compound, and varying concentrations and the effect on its melting temperature monitored, a shift in melting temperature is indicative of an interaction. The assays were performed in 100 mM of BTP pH 7.0 with 5 % DMSO, 150 mM NaCl. Assays were performed in triplicate.

| Description | Scientific name | Max Score | Total score | Query coverage | E-value | Identity (%) | Acc. Length | Accession no. |
|---|---|---|---|---|---|---|---|---|
| diacylglycerol kinase family protein | Bacteroides xylanisolvens XBA1 | 619 | 619 | 100% | 0.0 | 96.75% | 308 | WP_004316677.1 |
| YegS/Rv2252/BmrU family lipid kinase | Bacteroides ovatus ATCC 8438 | 615 | 615 | 100% | 0.0 | 96.10% | 308 | WP_004301802.1 |
| YegS/Rv2252/BmrU family lipid kinase | Bacteroides caccae ATCC 43185 | 610 | 610 | 100% | 0.0 | 95.13% | 308 | WP_005677430.1 |
| diacylglycerol kinase family protein | Bacteroides fragilis NCTC 9343 | 565 | 565 | 100% | 0.0 | 88.64% | 308 | WP_010992246.1 |
| diacylglycerol kinase family protein | Bacteroides intestinalis DSM 17393 | 548 | 548 | 100% | 0.0 | 84.42% | 308 | WP_007666434.1 |
| diacylglycerol kinase family protein | Bacteroides cellulolyticus DSM 14838 | 546 | 546 | 100% | 0.0 | 83.77% | 308 | WP_007212750.1 |
| diacylglycerol kinase family protein | Bacteroides oleiciplenus DSM 22535 | 545 | 545 | 100% | 0.0 | 83.77% | 308 | WP_009129050.1 |
| diacylglycerol kinase family protein | Bacteroides clarus DSM 22519 | 543 | 543 | 100% | 0.0 | 83.77% | 308 | WP_009122527.1 |
| YegS/Rv2252/BmrU family lipid kinase | Bacteroides salyersiae DSM 18765 | 537 | 537 | 100% | 0.0 | 83.44% | 308 | WP_007478527.1 |

**Supplementary table 7. Conservation of BT4322 in select Bacteroidota species of the human gut microbiota.**
BT4322 was used as the query against the genomes of the 9 other *Bacteroidoes* species from Figure 5 against which arylsulfamates 3 and 12 were tested. The BLASTp search was done using default settings with the blastp algorithm accessed through the NCBI interface.

**Dataset 1. Results output of the thermal proteome profiling analysis of compound 2 and 17**

**Dataset 2. Lipidomic analysis of *B. theta* grown in BHI in the presence of arylsulfamate 2.**
